# A biologically plausible decision-making model based on interacting neural populations

**DOI:** 10.1101/2023.02.28.530384

**Authors:** Emre Baspinar, Gloria Cecchini, Michael DePass, Marta Andujar, Pierpaolo Pani, Stefano Ferraina, Rubén Moreno-Bote, Ignasi Cos, Alain Destexhe

## Abstract

We present a novel decision-making model with two populations. Each population is composed of Regularly Spiking (excitatory) and Fast Spiking (inhibitory) cells in cortical layer 2/3. Each population votes for one of the two visual alternatives shown on a monitor in human and macaque experiments. The model is biophysically plausible since it is based on long-range cortico-cortical connections between the layer 2/3 populations. These connections are excitatory. They contact both Regularly Spiking and Fast Spiking cells. This long-range excitation is conflicted by an inhibition based on local connections within the populations. This configuration introduces a competition between the layer 2/3 populations, sufficient for making a decision to choose between two alternatives shown on the monitor. We integrate the model with a reward-driven learning mechanism. This allows the model to learn the optimal strategy maximizing the cumulative reward in the long term. We test the model on two decision-making tasks applied on human and macaque. This model elaborates certain biophysical details which were not considered by simpler phenomenological models proposed for similar decision-making tasks. Finally, the model can be embedded in a brain simulator such as The Virtual Brain to study decision-making in terms of large-scale brain dynamics.

## 1 Introduction

Decision-making refers to choosing one of the existing alternatives. A decision is often made by considering short and long term consequences of these alternatives. In many cases, making decision with a consideration of long term benefits is important for survival. This requires a resistance to immediate decisions bringing benefits in the short term but rather loss in the long term. Such resistance is based on a control over future planning to identify the optimal decision-making strategy bringing long-term, strong benefits.

In complex decision-making tasks, each choice is attributed to a reward. The brain builds a decision-making strategy according to these rewards. It aims to make the choices which maximize the reward. To build such a strategy is not straightforward since a choice might bring a high, immediate reward in the short term, but this reward can decrease in the long term. This happens when the consequences of the choice in the long term are not as beneficial as in the short term. Therefore, insisting on such choices amounts to a weak cumulative reward in the long term. For example, eating junk food gives strong, immediate pleasure, which is a high reward in the short term. However, if it is consumed systematically, it can cause several health problems in the long term, which is a weak reward or rather a loss. In such cases, maximization of cumulative reward requires a long-term planning. This can be provided via a decision-making strategy based on inhibitory self-control over the impulsive decisions.

Our interest is in such inhibitory self-control over the impulsive decisions. To study it, we focus on two experiments. These experiments are based on two similar reward-driven visual discrimination tasks: one applied on human (DePass et al., 2023; Cecchini et al., 2024) and the other one applied on macaque (Fontana et al., 2022); see Figures 1a and 1b. These tasks are composed of a certain number of units called episodes. The episodes are composed of subunits called trials. Each episode has the same fixed number of trials. In each trial, two objects are shown on a computer monitor to the subject. The subject chooses one of them. Then, a high reward or a low reward is provided to the subject according to the choice which the subject made. In every episode, both high and low rewards are increased or decreased at the end of each trial. Whether the rewards increase or decrease depends on the decision which the subject made in the trial. The goal is to obtain the maximum possible cumulative reward at the end of all trials in the episode (and in every episode), i.e., the maximum normalized sum of all rewards obtained through the trials.

**Fig 1.**
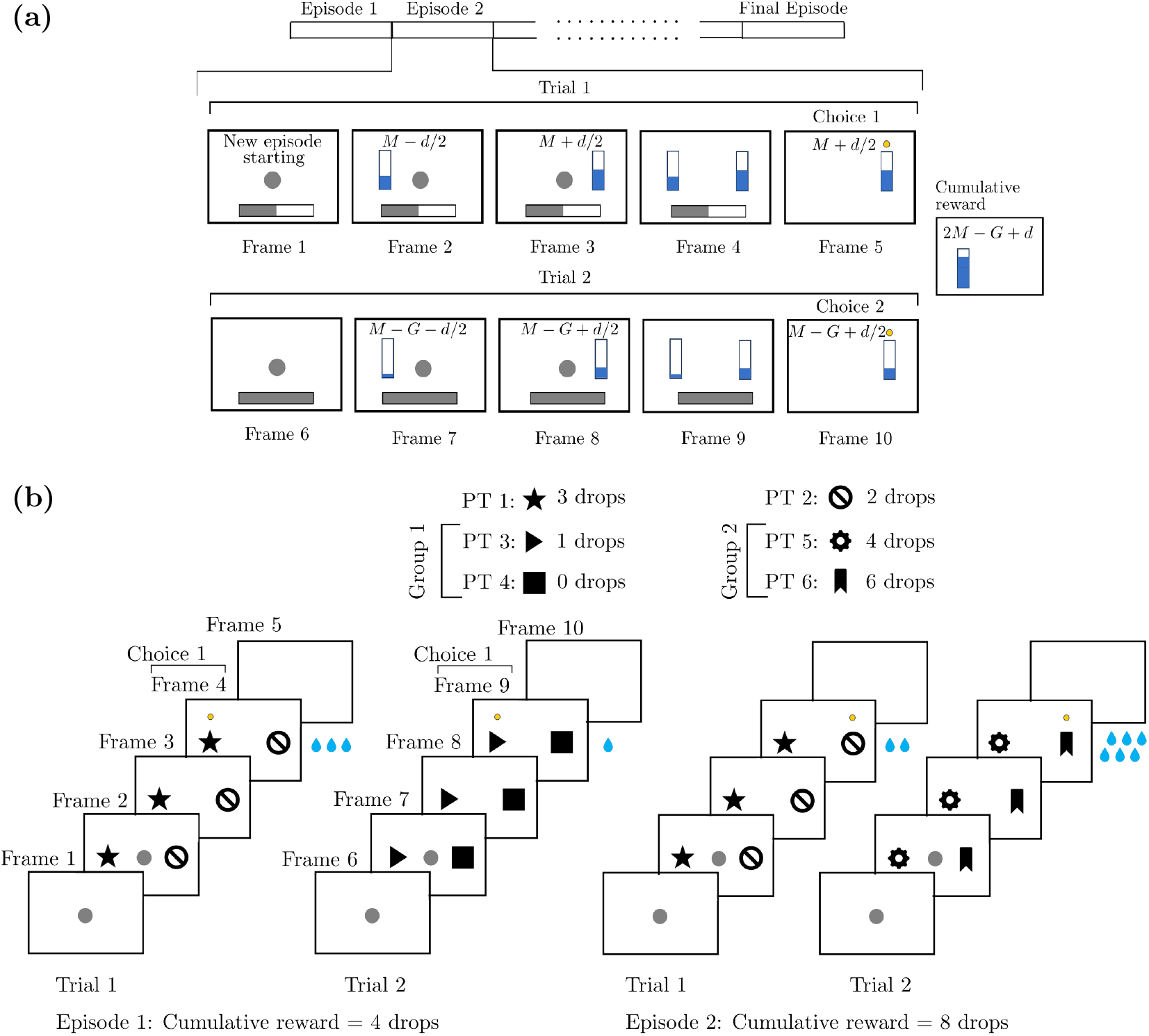
Example episodes from the Horizon 1 human and macaque experiments shown in (a) and (b), respectively. **(a)** A central target (CT) appears in Frame 1 to indicate the beginning of the episode. A progress bar at the bottom indicates in which trial the participant is. The stimuli with values *M* − *d/*2 and *M* + *d/*2 are shown separately in Frames 2 and 3, then together in Frame 4. The participant moves the pointer to the larger stimulus as highlighted by the yellow dot in Frame 5. The end of the episode is indicated in Frame 6 with a dot at the center. The stimuli are decreased in Episode 2, as shown separately in Frames 7 and 8. They are shown together in Frame 9 and the participant chooses the larger one as highlighted in Frame 10. The cumulative reward is the sum of the chosen stimuli. **(b)** A CT appears in Frame 1 to indicate the beginning of the episode. The subject touches the CT to initiate the episode. The peripheral targets (PTs) are shown together on the both sides of the CT in Frame 2. The CT disappears as the participant starts to move its hand to choose one of the PTs as shown in Frame 3. The chosen PT (PT 1) is highlighted by the yellow dot in Frame 4. The corresponding reward (3 water drops) is provided, and in Frame 5, a blank screen is shown to indicate the end of Trial 1. The same procedure is followed in Trial 2 but now with the PTs of Group 1. In the second scenario, the cumulative reward is higher since PT 2 is chosen in Trial 1 and hence the Group 2 PTs, which provide a higher reward, are shown on the monitor in Trial 2. The nomenclature and corresponding rewards to the PTs are given at the top. The cumulative reward is the sum of the provided water drops.

The particularity of the task is that choosing the high reward stimulus in every trial does not amount to the maximum cumulative reward. There is a specific choice pattern to follow throughout all the trials in an episode. This pattern was designed to provoke the self-control over impulsive decisions. In an example episode with two trials, if the high reward choice is made in the initial trial, the reward amounts (both high and low) will be smaller in the subsequent trial such that the cumulative reward cannot reach to the maximum. Therefore, the subject must follow a specific strategy to obtain the maximum cumulative reward: a strategy which promotes the choice bringing the low reward in the initial trial. This generates an increase in both reward values in the second trial. This is an increase which prevails the difference between the high and low rewards of the first trial. Subsequently, if the subject makes the choice bringing the high reward in the second trial, the cumulative reward achieves the maximum for that episode. This requires a self-control to avoid the decision which bring the high reward in the initial trial.

Our goal is to model the neural population dynamics induced by these experimental tasks. To achieve it, we consider one of the simplest and sufficient scenarios for a decision to be made: two layer 2/3 (L2/3) populations compete and the winning population determines which alternative to select (Desimone et al., 1995; Reynolds et al., 2004; Moore et al., 2017). In this scenario, the competition could be biased in favor of one of the two alternatives.

We propose a neural population model based on three modules for this scenario. The first one is the basic module. It generates the firing rates of the excitatory and inhibitory neurons by using a low dimensional approximation of the Adaptive Exponential (AdEx) neuronal network (Brette et al., 2005). In the basic module, there are two L2/3 populations. Each population is composed of one pair of excitatory-inhibitory subpopulations. Each excitatory subpopulation votes in favor of one of the two visual alternatives shown on the monitor. These excitatory subpopulations compete with each other to determine the decision. The one with the higher firing rate wins the competition and determines which visual alternative to choose. The second module is the regulatory module. It introduces the bias for either one of the two alternatives shown on the monitor. The final module is the reward-tracking module. It keeps track of the increase or decrease in the reward values, and feeds it into the regulatory module so that the regulatory module generates the bias accordingly: if the reward increases as the model proceeds to the next trials, then the bias favors the choices made in the previous trials, otherwise the bias disfavors them.

We use a mean-field approach to describe the subpopulation firing rates in the basic module. The mean-field equations provide a simple, low dimensional approximation of the averaged dynamics of the AdEx neuronal network (Brette et al., 2005), which is high dimensional and complex to analyze. This motivates our choice for the mean-field approach. In our model, the mean-field equations of the AdEx neuronal network represents a population of Regulary Spiking (RS, excitatory) and Fast Spiking (FS, inhibitory) cells in L2/3. The L2/3 populations are assumed to communicate via excitatory connections. This represents the network of functional columns found in the prefrontal cortex, as observed in macaque (Hirata et al., 2008). Our model examines if two of such interacting populations can implement decision-making mechanisms.

Several neural population models were previously proposed for similar decision-making problems. Some of them used attractor networks to model switching between two (or more) decisions. Such switching could occur due to perceptual bistability (Moreno-Bote et al., 2007), or due to further evidence accumulation to make a decision (Albantakis et al., 2011). Differently from these, a recurrent network model of object working memory was proposed in (Brunel et al., 2001). Thereafter, another network model with a similar architecture was proposed for a visual discrimination task in (Wang, 2018). This final architecture was adapted to a two-choice perceptual task, where the effect of memory-based experience on the decisions was included and modeled in terms of neural populations (Marcos et al., 2013; Cecchini et al., 2024). An alternative to these models was proposed in (Hayden et al., 2018). This framework considers the decision as the result of modulatory effects of attention, on the dynamics of a single population.

All these previous models are based on connectivity architectures which are not in accordance with the anatomy. In the basic module of our model, the L2/3 populations are connected to each other only via excitatory connections. Each excitatory connection originates from the excitatory subpopulation of one of these L2/3 populations, and it is exerted onto both excitatory and inhibitory subpopulations of the other L2/3 population (Figure 3a). This is similar to the network of functional columns in the prefrontal cortex, as found in macaque (Hirata et al., 2008). Therefore, our model is designed based on an anatomically coherent connectivity.

It is important to note that, the transfer functions in the AdEx mean-field equations were obtained by fitting membrane variables of a semi-analytical template to *in vitro* measurements (Zerlaut et al., 2016). Then, the free parameters of the same function were fitted to the network responses of RS and FS neurons which were bombarded by the spikes generated from a Poisson distribution (Zerlaut et al., 2016, 2018; di Volo et al., 2019). In the transfer functions, RS neurons display spike-frequency adaptation whereas FS neurons do not, as observed experimentally in the pyramidal neurons and interneurons, respectively. Finally, the AdEx mean-field framework simulates more precisely the inhibition and the excitatory/inhibitory balance through its spike-frequency adaptation compared to other frameworks.

All these points regarding the connectivity and the transfer functions, provide the biophysically plausible nature of our model. They make our model potentially applicable to neurophysiological recordings from the cortex. This distinguishes our framework from the previous models proposed for similar decision-making tasks.

Another important property of our model is at the modular level. Classical AdEx mean-field framework was proposed for only one population (Zerlaut et al., 2018; di Volo et al., 2019), which is not sufficient for a competition resulting in a decision. We extend these equations to two populations (i.e., two subpopulation pairs) so as to provoke a competition between them. Moreover, we integrate this extension with the regulatory and reward-tracking modules. This endows the AdEx mean-field framework with a reward-driven learning mechanism.

These novelties introduce several challenges at both mathematical and numerical levels. Firstly, the extension of the classical AdEx mean-field framework (Zerlaut et al., 2018; di Volo et al., 2019) to two L2/3 populations increases the dimension of the mean-field system from 6 to 16. Secondly, the connections between these L2/3 populations introduce additional derivatives of the cross-correlations between the subpopulation firing rates. These two points increase memory and computational load. Finally, two noise sources should be incorporated in the model. The first one is introduced to the basic module and it models the distortion effects of the extracellular matrix on the subpopulation firing rates. The second one is introduced to the regulatory module. It models hesitation, curiosity, perceptual difficulties or any other reason causing erroneous decisions. These two types of noise should be introduced in a balanced manner such that they do not dominate each other, neither the deterministic model dynamics.

We test our model on the experimental data in terms of behavioral metrics. Two parameters characterize the performance of the model with respect to these metrics: learning speed and flexibility parameter. The former determines how quickly the model captures the optimal strategy maximizing the cumulative reward. It can be thought of as the modulation effects of acetylcoline and dopamine on the learning dynamics (Brzosko et al., 2015, 2017). The latter determines how much the model is flexible to make mistakes, or to try different decision-making strategies to explore the consequences, sometimes even after that it learns the optimal strategy.

The implementation of the model is provided in Python as a Jupyter notebook (Baspinar et al., 2023), which can be simulated also online on the EBRAINS cloud system (Baspinar et al., 2023).

## 2 Experiment setup

The experiments were designed to study the behavioral background of the inhibitory self-control over impulsive decisions bringing immediate rewards, which nevertheless amount to low cumulative reward in the long term. These experiments were performed on human (Cecchini et al., 2024; DePass et al., 2023) and macaque (Fontana et al., 2022) by following a different protocol for each species. The difference is based on the type of the visual stimuli and rewards.

In both protocols, the experiment is composed of a number of episodes. At the beginning of each episode, the protocol is restarted with randomly generated visual stimuli. Each episode is composed of two successive trials. There is a preset strategy which was implemented in every episode of the experiment. This preset strategy refers to a unique pattern of choice sequence. In a single episode, this pattern is based on choosing the stimuli bringing the low reward in the first trial and the stimulus bringing the high reward in the second trial. The key point of the strategy is that, the rewards generated in the second trial are larger if the subject chooses the stimulus which brings the low reward in the first trial. They are larger, in comparison to the rewards generated in the second trial when the subject chooses the stimulus which brings the high reward in the first trial. This increase in the reward values is bigger than the difference between the high and low rewards of the first trial. Therefore, the maximum cumulative reward is achieved only if the subject follows the preset strategy, although the low reward of the first trial gives the opposite impression.

The goal of the subject is to obtain the maximum possible cumulative reward throughout every episode. This requires: (i) to identify the reward values and the correspondence of these values to the choices made, (ii) to capture the preset strategy in the task to make the sequence of decisions providing the maximum cumulative reward.

### 2.1 Human experiment

The human experiments were conducted at Universitat de Barcelona, Facultat de Matemàtiques i Informàtica. The task was run with 28 participants. We considered a 20-year-old female participant as an example to fit our model. Nevertheless, the model can be fitted to the other participants as well. Here we provide a summary of the task design. More details and relevant data can be found in (Cecchini et al., 2024) and on EBRAINS (DePass et al., 2023), respectively.

Two types of experiments are considered: Horizon 1 and Horizon 0. Each experiment is composed of *K* episodes, with *K* denoting the total number of episodes in the experiment. In Horizon 1, each episode is composed of two successive trials, which are named as Trial 1 and Trial 2 (Figure 1a). In Horizon 0, each episode is composed of a single trial. Therefore, Horizon 0 boils down to a visual discrimination task. Behavioral results are quantified in terms of performance indexes and also reaction times. Performance index provides a measure of the overlap between the choices made by the participant and the preset strategy in an episode. It is registered per episode. Reaction time shows how long it takes for the participant to make the decision in each trial. It is registered per trial.

It is important to note that the stimuli shown on the monitor are in the form of partially filled vertical bars (Figure 1a). Moreover, the reward in each trial is the amount of the filled part of the chosen vertical bar. The low reward is equal to the small stimulus, which is the less filled bar. The high reward is equal to the large stimulus, which is the more filled bar. The cumulative reward is the sum of the rewards, i.e., the sum of the filled parts of the chosen stimuli throughout these trials. The preset strategy in Horizon 1 experiments is to choose the smaller stimulus in Trial 1 and the larger stimulus in Trial 2. In the first trial of an Horizon 1 episode, if the choice of the participant overlaps with the preset strategy, the amounts of rewards (thus the filled parts of the stimuli) are increased equally in the second trial. Otherwise, they are decreased equally. The value of these increase and decrease is generated from a random set at the beginning of the experiment and it is kept fixed throughout the whole experiment.

Before the experiment, the participant is instructed to choose one of the two stimuli projected on the monitor in each trial. Furthermore, the participant is instructed to maximize the cumulative reward for each episode, thus for the whole experiment: the total cumulative reward for the whole experiment is maximum if and only if the participant makes the choices in complete coherence with the preset strategy in every episode of the experiment. In Horizon 0, the strategy boils down to choosing the larger stimulus in each episode, i.e., in each trial.

At the beginning of an episode, the amounts of the filled parts of the bars are generated as *M ± d/*2. These stimuli are shown at the beginning of Trial 1. Here *M* denotes the mean value of the stimuli, and *d* is the amount of difference between the two stimuli. In each episode, *M* is randomly and independently generated from a uniform distribution at the beginning of the episode. We use one of the five different values of *d* in each episode. In total, each one of these five values are used equally in *K/*5 episodes of the experiment. In Horizon 1, if the participant chooses the small stimulus in Trial 1, the gain *G > d* is added to both stimuli and *M* + *G ± d/*2 become the filled quantities of the two stimuli in Trial 2. Otherwise, the gain *G* is subtracted from the stimuli and the filled quantities become *M* −*G± d/*2 in Trial

2. The same procedure is applied in Trial 2 but now the choice according to the preset strategy is the large stimulus. Therefore, there are 4 possible choice patterns which the participant can follow. Each choice pattern corresponds to one of the following 4 cumulative reward values (from maximum to minimum): 2 *M* + *G*, 2 *M* + *G* −*d*, 2 *M* −*G* + *d*, and 2 *M* −*G*.

We provide an example scenario of an episode in Figure 1a. In Trial 1 of this scenario, the participant chooses the large stimulus with *M* + *d/*2 (i.e., the high reward). This does not overlap with the preset strategy. Therefore, the stimuli (i.e., the rewards) are decreased by *G*, resulting in *M* − *G* + *d/*2 and *M*− *G*− *d/*2 as large and small stimuli values, respectively. In Trial 2, the participant chooses the large stimuli (i.e., high reward), and this overlaps with the preset strategy. The resultant cumulative reward is 2− *M G* + *d*, which shows that the participant did not capture the preset strategy.

Finally, an important particularity of the human experiment is that the participant does not see at the end of Trial 2 the cumulative reward, or any explicit feedback. As a result of this lack of feedback at the end of the episode, the participant is forced to learn the consequence of the episode instead of learning a sequence of choices. See Figure 2 for some relevant example experimental results.

**Fig 2.**
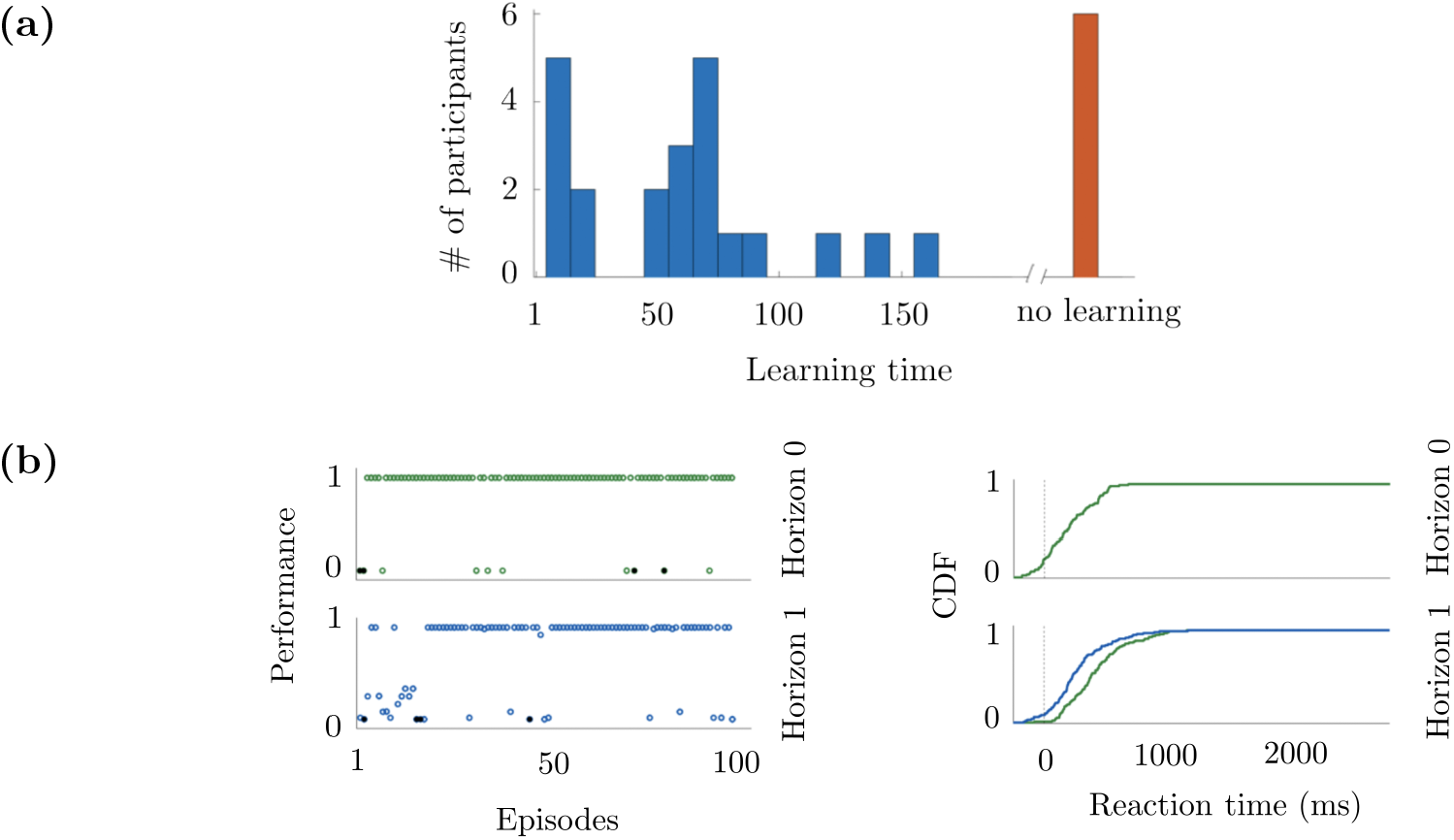
Some experiment results from (Cecchini et al., 2024). **(a)** The histogram of the learning times obtained from 28 human subjects. The learning time is in terms of episode numbers. The long learning time is classified as no learning. **(b)** The performance indexes of one of the participants, and the cumulative distribution function (CDF) of the reaction times of the same participant. The results are presented separately for Horizon 0 and Horizon 1, where in the latter, the CDFs of Trial 1 and Trial 2 are provided in green and blue, respectively.

**Fig 3.**
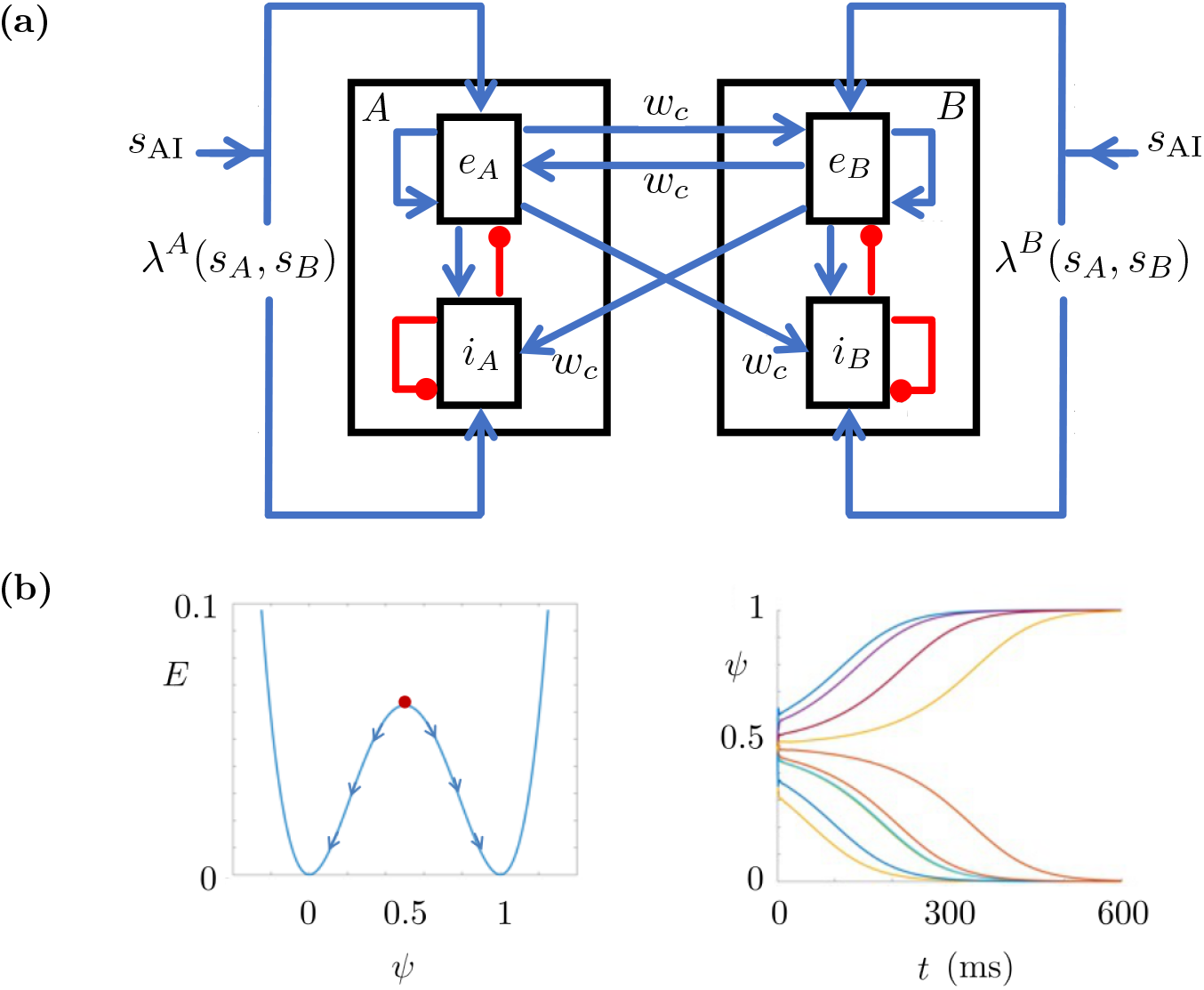
Model structures. **(a)** Basic module with two L2/3 populations composed of excitatory and inhibitory subpopulations. Populations *A* and *B* vote in favor of Stimulus *A* (*s*_*A*_) and Stimulus *B* (*s*_*B*_), respectively. The excitatory subpopulations are denoted by *e*_*A*_, *e*_*B*_ and the inhibitory subpopulations are represented with *i*_*A*_, *i*_*B*_. The excitatory and inhibitory connections are in blue and red, respectively. The weights of the connections between the L2/3 populations are denoted by *w*_*c*_ as in (4). Here *λ*^*A*^ and *λ*^*B*^ represent the outputs of the regulatory module introducing a bias to the input signals *s*_*A*_ and *s*_*B*_. Finally, *s*_AI_ is the base drive, which ensures that the model performs in the awake state. **(b)** Regulatory module. Left: Energy functional of *ψ*. The neutral initial condition is *ψ*(0) = 0.5. The increase and decrease in the rewards introduce a bias making *ψ*(0) *>* 0.5 or *ψ*(0) *<* 0.5 in the next episode. This results in *ψ*(*t*_*F*_) = 1 or *ψ*(*t*_*F*_) = 0 at the end of the corresponding trial of the next episode. The former introduces the bias favoring the large stimulus and the latter introduces the bias favoring the small stimulus. Right: Time evolution of *ψ* starting from different initial conditions. At the beginning, a small noise is introduced for modeling whatsover which might perturb the regulatory process, such as the hesitation, curiosity/exploratory behavior or perceptual difficulties.

### 2.2 Macaque experiment

The macaque experiments were conducted at Sapienza Università di Roma, Dipartimento di Fisiologia e Farmacologia. One 16-year-old, healthy, male macaque weighting 9.5 − 10.5 kg was trained for performing a Horizon 1 task similar to its counterpart in the human experiment. The relevant data can be found on EBRAINS (Fontana et al., 2022).

The task can be summarized based on the two possible scenarios provided in Figure 1b. A dataset of black and white stimuli were shown to the macaque in each trial. Six stimuli (peripheral targets; PTs) were extracted randomly from *Microsoft PowerPoint* shape library. Two of these six randomly extracted PTs were shown in Trial 1 and they were attributed to different rewards; PT 1: 3 water drops, PT 2: 2 drops. The remaining four stimuli were separated into two groups, each based on two PTs. In Group 1, we had PT 3 and PT 4; in Group 2, we had PT 5 and PT 6. Each of these four PTs corresponds to a different reward; PT 3: 1 drop, PT 4: 0 drop, PT 5: 4 drops and PT 6: 6 drops. The group which is shown in Trial 2 was determined by the choice made in Trial 1. More precisely, if the participant chooses PT 1 in Trial 1, Group 1 will be shown in Trial 2 as illustrated in Episode 1 of Figure 1b. Otherwise, Group 2 will be shown in Trial 2 as illustrated in Episode 2 of Figure 1b. At the beginning of each trial, a central target (CT) is shown at the center of a touchscreen monitor to indicate the beginning of the trial. The macaque is required to touch the CT to initialize the trial. Then, the stimuli are shown as the PTs appearing on the left and right hand sides of the CT. As the participant starts to move his hand, the CT disappears. This is considered as the Go signal initiating the reaction time, which is tracked for a maximum duration of 2000 ms. If a choice is made between two PTs within this 2000 ms, the reward is delivered to the subject. Once this protocol for Trial 1 is completed successfully, Trial 2 begins after an inter-trial interval of 1200 ms and it follows the same protocol.

Although both human and macaque tasks are based on perceptual distinguishing of the targets, there are three major differences between them. Firstly, the stimuli are partially filled bars in the human task, whereas they are some figures chosen randomly from the Powerpoint shapes library in the macaque task. Secondly, in the human task, the rewards are the same as the chosen stimuli. High and low rewards are the large and small stimuli, respectively. In the macaque task, the reward is provided as a certain number of water drops to the macaque. Finally, the reward is provided at the end of Trial 1 and Trial 2 in the macaque task. In the human task, there is an explicit feedback regarding the reward only at the end of Trial 1. There is no feedback at the end of Trial 2.

## 3 Model

We study the neural dynamics regarding the human and macaque experiment by using a neural population model. The model is composed of three layers: basic module, regulatory module, and reward-tracking module.

### 3.1 Basic module

The basic module generates the neural activity in terms of subpopulation firing rates. It describes the firing rates by using an extension of the classical AdEx mean-field equations (di Volo et al., 2019) to two L2/3 populations.

The classical AdEx mean-field equations were derived in (Zerlaut et al., 2018; di Volo et al., 2019) from the AdEx neuronal network (Brette et al., 2005). They describe the average dynamics of a pair of excitatory and inhibitory subpopulations which are coupled to a spike-frequency adaptation. We extend it to two pairs, where each pair is a L2/3 population composed of one excitatory and one inhibitory subpopulation (Figure 3a). In this extended setting, each L2/3 population votes for either one of the two stimuli: Population *A* votes for Stimulus A and Population *B* votes for Stimulus *B*. We denote Stimuli *A* and *B* with *s*_*A*_ and *s*_*B*_, respectively. One of them represents the choice bringing the high reward and the other one represents the choice bringing the low reward. Stimulus *A* and *B* may represent these choices interchangeably in different episodes. However, they are assigned to either one of the choices within a single episode, and it does not change throughout the episode. In the basic module, we denote the excitatory subpopulations by *e*_*A*_, *e*_*B*_ and the inhibitory subpopulations by *i*_*A*_, *i*_*B*_. The subindexes *A* and *B* indicate the corresponding L2/3 population. Each L2/3 population represents a functional column of *N*_*e*_ excitatory and *N*_*i*_ inhibitory neurons. The ratio *N*_*i*_*/N*_*e*_ is 0.2.

The basic module connectivity follows an anatomically coherent connectivity architecture. Across the L2/3 populations, there are only excitatory connections. This connectivity is similar to the network of functional columns in the prefrontal cortex, as found in macaque (Hirata et al., 2008). Each one of these excitatory connections between the L2/3 populations originates from the excitatory subpopulation of one L2/3 population and targets both the excitatory and inhibitory subpopulations of the other L2/3 population (Figure 3a). The weights of the connections between the L2/3 populations are denoted by *w*_*c*_. Local connections are within the populations. They are based on all-to-all connectivity within each population, with both recurrent and cross-subpopulation connections as shown in Figure 3a. All the local connections have unit weights.

Similarly to the classical AdEx mean-field setting (di Volo et al., 2019), the basic module can function in two modes: asynchronous irregular (AI) and up-down. Here AI mode refers to the awake state of the brain and up-down refers to the sleep (or under anesthesia) state of the brain. We consider only the AI state since decision-making requires that the subject is awake. Therefore, we keep the basic module in AI mode by feeding it with a constant base drive *s*_AI_. This is a DC input (=5 Hz) which keeps the basic module away from the up-down mode. The terms *λ*^*A*^(*s*_*A*_, *s*_*B*_) and *λ*^*B*^(*s*_*A*_, *s*_*B*_) in Figure 3a represent the inputs to Populations A and B, respectively. These are the biased versions of the stimuli *s*_*A*_, *s*_*B*_. The bias is introduced by a regulatory module that models plasticity as explained in Section 3.2. The biased inputs are simultaneously fed to Populations *A* and *B* through the functions *λ*^*A*^ and *λ*^*B*^ described in (7), respectively.

We denote the firing rates of the four subpopulations by 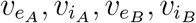.Each one of them represents the average spike frequency of one of the four subpopulations. We describe the neural interactions between these subpopulations via cross-correlations between the firing rates of interacting subpopulations. For simplicity of notation in the following, we use *α, β, ξ, η* to denote any of the four subpopulations *e*_*A*_, *e*_*B*_, *i*_*A*_, *i*_*B*_. Then, we represent the cross-correlation between *v*_*α*_ and *v*_*β*_ with *C*_*αβ*_. Finally, the variables 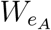 and 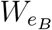 describe the spike-frequency adaptation of the RS cells, i.e., the excitatory subpopulations *e*_*A*_ and *e*_*B*_, respectively. The adaptation is only for the excitatory subpopulations as FS cells have no spike-frequency adaptation. Thus, we assume no adaptation for the inhibitory subpopulations, i.e., 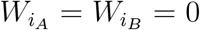.As a result, the basic module has 16 state variables.

The basic module equations read as:

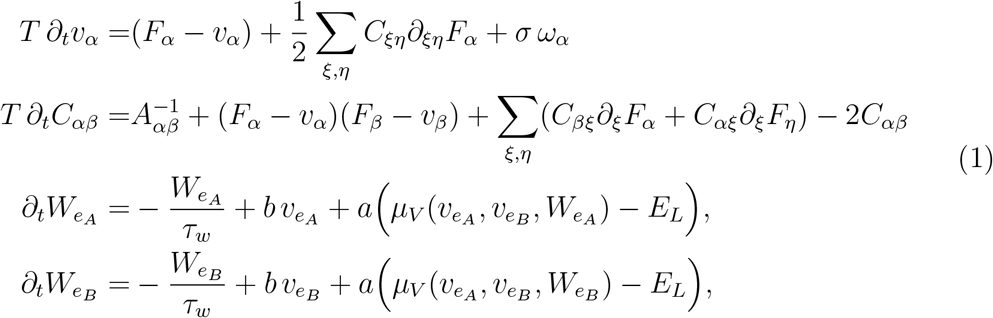

where we use the notation given by 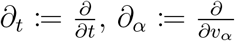 and 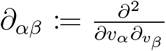 for the partial derivatives. Here *E*_*L*_ is a constant representing the reverse leakage potential, and *T, τ*_*w*_ are the time scale parameters. We denote by *ω*_*α*_ a white Gaussian noise, with *σ >* 0 denoting its intensity. This noise is sampled from a normal distribution independently for each subpopulation and at each instant. More precisely, 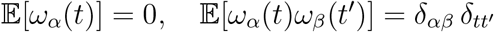.for all *α, β* ∈ {*e*_*A*_, *i*_*A*_, *e*_*B*_, *i*_*B*_} and for all *t, t*′ ≥ 0, where δ is the Dirac delta function. Here 𝔼 [·] represents the expected value. Function *A*_*αβ*_ appears from the mean-field derivation of the second order statistical moments and it is defined for *α, β* ∈ {*e*_*A*_, *i*_*A*_, *e*_*B*_, *i*_*B*_} as follows:

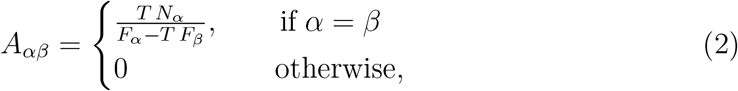

where *N*_*α*_ denotes the number of neurons represented in the subpopulation *α*. We choose 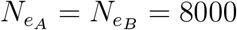 and 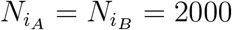.An important group of functions appearing in the mean-field equations (1) is the transfer functions, denoted by *F*_*α*_. A transfer function gives the output firing rate of a subpopulation according to the input that the subpopulation receives. This output firing rate is modulated by the spike-frequency adaptation in the excitatory subpopulations. The transfer functions appearing in (1) and (2) are written with their inputs as follows:

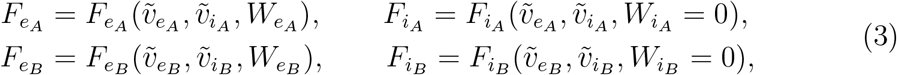

with subindexes indicating the subpopulation type. Each input to the transfer functions is composed of excitatory and inhibitory components 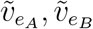 and 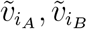,respectively. Each component has a local part and a part related to the interactions between the L2/3 populations. The latter is weighted by *w*_*c*_ since the interactions take place through the connections between the L2/3 populations. These excitatory and inhibitory components are as follows:

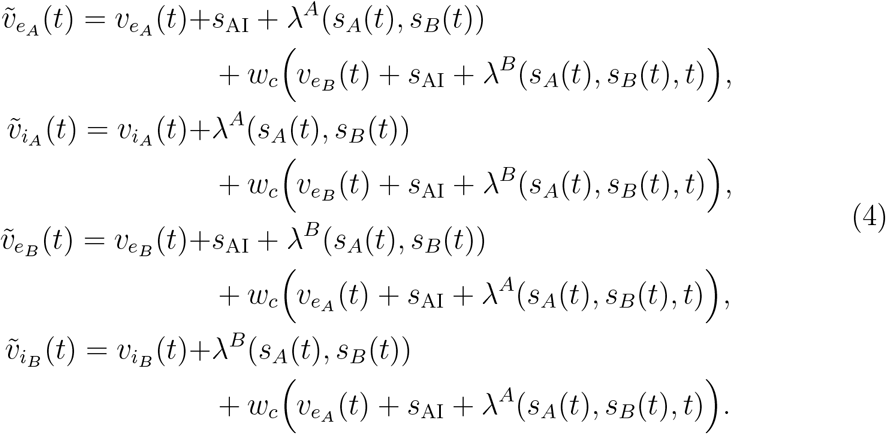

Here the time dependency is explicitly denoted and the terms with no explicit time variable are constants. The external stimuli *s*_*A*_, *s*_*B*_ are fed in (4) via *λ*^*A*^, *λ*^*B*^. It is to take into account the bias introduced through the regulatory module, which will be explained in Section 3.2. A common choice to model the external stimuli *s*_*A*_, *s*_*B*_ is Heaviside function in mean-field models. However, since the Heaviside function has a discontinuity due to the discrete jump, our model undergoes a transient phase which might obscure the instant when the model makes a decision. For this reason, we perform our simulations by using a sufficiently sharp sigmoid function as external stimulus; see Figure 10b. Finally, *s*_AI_ denotes the DC input which keeps the basic module in the AI state.

We use the same formula for the transfer functions as provided in (di Volo et al., 2019); see Appendix for the details. We report here the formula directly:

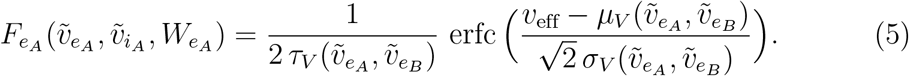

This formula is for *e*_*A*_ and it is the same for the other subpopulations once the subindexes are changed accordingly. Here, *µ*_*V*_ and *σ*_*V*_ represent the mean and standard deviation of the membrane voltages of the neurons found in the AdEx network, respectively. Their formulas can be found in (20) and (21). The term *τ*_*V*_ is the global autocorrelation time of the membrane voltage fluctuations of the neurons in the AdEx network; see (21) in Appendix. It gives a measure of the speed of the fluctuations. Finally, *v*_eff_ is the voltage-effective threshold and it is a second order polynomial of *µ*_*V*_, *σ*_*V*_, *τ*_*V*_ ; see (22). It was obtained by a fit according to the simulations of single neuron activity produced at multiple excitatory and inhibitory presynaptic frequencies (Zerlaut et al., 2018; di Volo et al., 2019). The transfer function (5) was derived via a semianalytical procedure based on the output spiking of an AdEx network of 10^4^ neurons, where each neuron was modeled by the AdEx integrate-and-fire equations given by (15). The network was bombarded with spikes generated from a Poisson distribution and a semi-analytical function was fitted to the outputs of the AdEx network (di Volo et al., 2019). See Appendix for more details.

The derivation of the AdEx mean-field equations from the AdEx neuronal network was provided in (di Volo et al., 2019). In the present paper, we use the same mean-field equations to describe the dynamics of each L2/3 population. The extension to two L2/3 populations is made symmetrically. That is, the L2/3 populations are set to the identical parameter values and they have the identical local connectivity architecture within the populations. Accross the L2/3 populations, the coupling is only excitatory and all the excitatory connections between the L2/3 populations have the same connection weight (= *w*_*c*_). This allows us to write the mean-field equations of our extended framework directly from the equations provided in (di Volo et al., 2019), avoiding a derivation from an AdEx network of four subpopulations.

### 3.2 Regulatory module

The basic module is not associated with any learning mechanism. Nevertheless, a learning mechanism is required to capture the preset decision-making strategy, which gives the maximum cumulative reward. To introduce a learning mechanism, we integrate the basic module with a regulatory module. The regulatory module identifies the reward quantities and how they change according to the choices made by the model. Then, it introduces the bias to the external stimuli in such a way that, the stimuli corresponding to the choices which bring large cumulative reward are promoted. This module can be thought of as a gating mechanism arranging the motor-plan flexibility for the response to the stimuli (or to the sensory inputs evoked by the stimuli). This mechanism might provide an explanation to the observations showing that relevant sensory inputs, distractors, and direct perturbations of decision-making circuits affect the behavior more strongly when they are introduced in the early phase of the stimulation (Seidemann et al., 1998; Kiani et al., 2008; Kopec et al., 2015; Van Ede et al., 2018; Zuo et al., 2019; Finkelstein et al., 2021).

The regulatory module was adapted from (Cecchini et al., 2024). It introduces the bias by weighting the stimuli fed into the L2/3 populations. The bias makes the population voting for the promoted stimulus to be more likely to have a higher excitatory firing rate than the other population. Consequently, the population voting for the promoted stimulus is more likely to win the competition and to make the decision; see Figures 4 and 10. We denote by 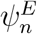 the regulatory function evolving during the *n*^th^ trial of the *E*^th^ episode. It is described via

**Fig 4.**
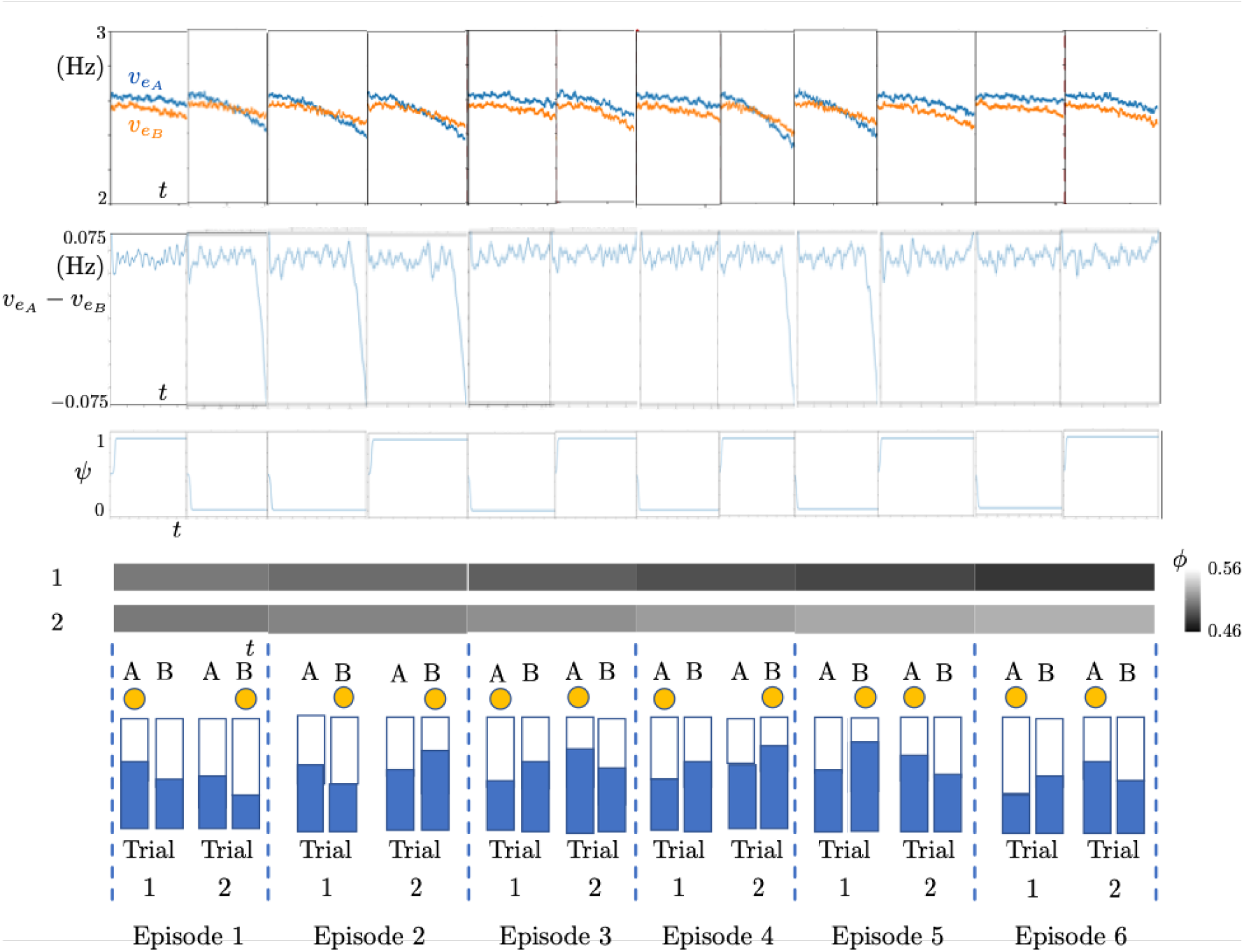
Simulation results of 6 episodes. First row: Time courses of the excitatory subpopulation firing rates. Second row: Time courses of the difference of the excitatory subpopulation firing rates. Third row: Time course of the regulatory function *ψ* for each trial of the corresponding episode given in the bottom row. Fourth row: Time course of the reward function φ for Trial 1 (top) and Trial 2 (bottom) of the corresponding episode. Bottom row: External stimuli. Chosen stimuli are highlighted by the yellow dot at the top. Trial and episode numbers are given at the bottom.

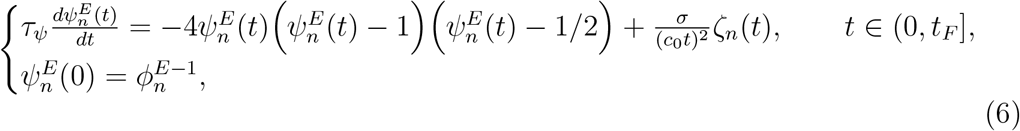

where *τ*_*ψ*_ is the time scale parameter. The regulatory function is restarted at the beginning of each trial. It evolves simultaneously with the basic module in time. Here *t*_*F*_ *>* 0 is the final time of the trial and it is the same for every trial. Moreover, ζ_*n*_ = ζ_*n*_(*t*) is a white Gaussian noise. Its intensity level *σ >* 0 is scaled by a constant *c*_0_ *>* 0 and time. Therefore, this noise introduces a strong stochastic behavior initially. Then, it decays in time with a rate depending on *c*_0_. It models the perceptual difficulties, hesitation and curiosity/exploratory behavior of the participant.

The decision in a trial depends on the evolution of the function 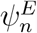 in time. This function converges to either 1 or 0 in the course of each trial. The convergence towards 1 pushes the model to choose the stimulus bringing high reward, e.g., the large stimulus in the human experiment. The convergence towards 0 pushes the model to choose the stimulus bringing low reward, e.g., the small stimulus in the human experiment. These dynamics of the regulatory module are continuously transmitted to Populations *A* and *B* via

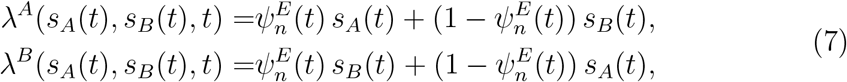

where *t*∈ [0, *t*_*F*_].

It is important to note that the process given by (6) is reinitialized at the beginning of each trial *n* of the *E*^th^ episode. It is reinitialized from the initial condition fixed to the value determined by the reward-tracking function 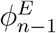, which will be explained in the next section. The function 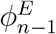 corresponds to the same trial number *n*, but of the (*E* − 1)^th^ episode. In this way, the reward-tracking function provides at the end of the (*E*− 1)^th^ episode, a separate feedback for each trial *n* of the *E*^th^ episode. This feedback is used as the initial condition for the regulatory function equations (6), at the beginning of each trial of the *E*^th^ episode. The feedback provides information about how optimal the decision-making strategy of the previous episode was, therefore about the accumulated experience. In this way, the feedback determines towards which decision the regulatory module will introduce the bias to the basic module in the next episode.

### 3.3 Reward-tracking module

The regulatory module introduces the bias depending on a feedback. This feedback is provided as the initial condition to the regulatory function equations given by (6). We should take into account the accumulated experience throughout the previous episodes to provide the feedback for the upcoming episodes. This feedback must be such that the model can identify which choices contribute more to the cumulative reward. We integrate our model with a reward-tracking module to endow the model with such feedback mechanism.

The reward-tracking module is based on a reinforcement learning equation. This equation motivates the choices which increase the reward amounts, and demotivates the choices which decrease the reward amounts. This is associated with the increasing dopamine signaling during predicted reward anticipation: motivation to repeat the same decision and the dopamine signaling increase, if the obtained reward is as predicted or more (Mirenowicz et al., 1994; Schultz, 2022). This results in an online learning. Once the preset strategy is learned, the system makes the decisions in almost complete coherence with the strategy.

We assume in the human experiments that the increase and decrease in the stimuli (i.e., reward) is predicted as profit and loss by the participants, respectively. In accordance with this assumption, the reward-tracking module is updated through a discrete evolution over the episode numbers. In other words, the reward-tracking function value remains constant during a trial and it is increased or decreased to another constant at the end of the trial. The updated value is provided to the regulatory function *ψ* in the next episode as the initial condition; see (6) and (8).

In the human case, we use the notation 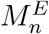 to denote the mean value of the stimuli corresponding to the *n*^th^ trial of the *E*^th^ episode. We consider here only Horizon 1 since Horizon 0 uses the same module but with *n* = 1. We write the evolution for the reward-tracking function φ as follows:

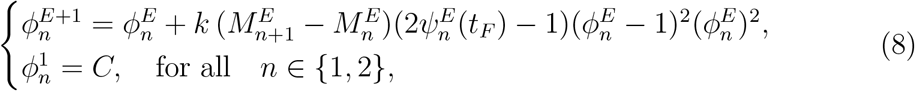

with *C* denoting a constant, which is fixed to 0.5 in our framework. Here *k* is the learning speed parameter. It determines how quickly the model captures the preset strategy throughout the episodes. In (8), 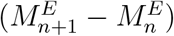 provides a coupling between the trials of an episode. It changes the sign of the polynomial. In this way, it determines how the bias provided by 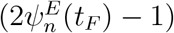 will be transmitted to the upcoming episode (i.e., to the episode *E* + 1): by favoring or by disfavoring the choice made in episode *E*? The same choice is favored in the upcoming episode if 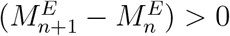, meaning that the stimuli (i.e., the rewards) increased as we pass from *n*^th^ to 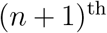 trial. It is disfavored if 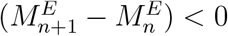, meaning that the stimuli (i.e, the rewards) decreased.

In the reward-tracking module for the macaque case, we use a similar idea to the human case, with a few differences. Let us recall that in the macaque experiment, the reward is provided to the subject as water drops. This is different from the human experiment, where the reward is the chosen stimulus itself. Therefore, we write the reward-tracking equations for the macaque case, in terms of the water drops provided for each choice. We replace the first line of (8) with

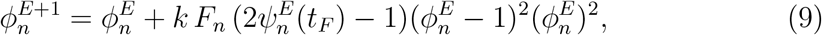

Where

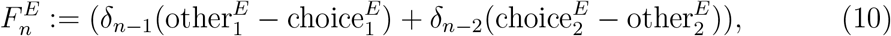

with *n* and δ denoting the trial number and the Dirac delta, respectively. Here choice 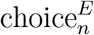 and other 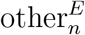 denote the numbers of water drops corresponding to the chosen stimulus and to the other stimulus, respectively, in the *n*^th^ trial of the *E*^th^ episode.

Here the reward is explicit. It is not represented in terms of the stimuli as in the human case.

### 3.4 Example simulation of the model

We provide here a small example simulation of the model to give an insight about the functionality of the three modules. In this example, we consider a human Horizon 1 simulation with 6 episodes.

In the human Horizon 1 experiments, the preset strategy requires choosing the smaller stimulus (the bar which is filled less) in Trial 1 and the larger stimulus (the bar which is filled more) in Trial 2 of each episode. This preset strategy is the optimal strategy which the model should learn so as to have the maximum cumulative reward. In Figure 4, we show a small example with 6 episodes of our Horizon 1 model simulations regarding the human case. The reward-tracking module φ is initiated from 0.5 for both trials in Episode 1, with a noise perturbing this initial value at the beginning of each trial. We observe that the model explores the strategy in Episode 1. Its decision is completely random with no bias since 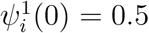 for both *i* = 1 and *i* = 2. In Episode 2, the reward-tracking module introduces a bias in the regulatory function *ψ* based on the feedback from Trial 1 and Trial 2 of Episode 1. The same procedure is repeated for Episode 3 but now based on the feedback from Episode 2, and so on. In Figure 4, we observe that the model learns the strategy in Episode 2. From Episode 2 onwards, it makes the choices in complete coherence with the preset strategy.

## 4 Noise sources

Noise is required in our model to consider different perturbative mechanisms. It appears at two levels in the model: in the basic module and in the regulatory module. The former models the distortion effects due to the extracellular media. The latter models the exploratory behavior of the subject which arises from curiosity, hesitation and/or perceptual difficulties.

We describe the noise in terms of Ornstein-Uhlenbeck (OU) processes. An OU process can be sampled directly from a Gaussian distribution once the initial condition of the process is chosen properly. This is similar to the previous framework (di Volo et al., 2019), however with one difference: we choose the initial condition of the OU process as a zero mean Gaussian, where its variance is scaled by the convergence rate of the OU process as given below. This allows to avoid explicit simulation of the OU process, reducing the computational load.

The noise is a stochastic process evolving in time, independently of the rest of the state variables of the model. For a white Gaussian noise, this evolution can be written as an OU process of the following type:

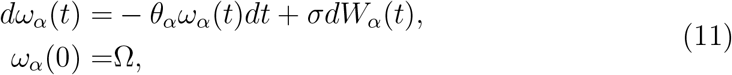

where *W*_*α*_(*t*) is a standard Brownian motion, *θ*_*α*_ is a positive constant associated to the convergence rate of the process to its long term mean, and Ω is the initial condition, which is a centered Gaussian noise with variance 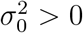.This is a linear stochastic differential equation and its unique solution (Arnold, 1974) is

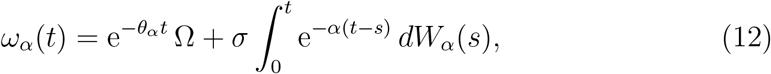

where the solution *ω*_*α*_(*t*) is a Gaussian process. It can be written for a small time step 0 *< h «* 1 as

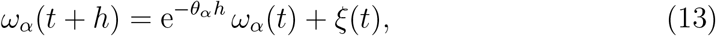

where *ξ*(*t*) is a centered Gaussian white noise with variance 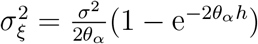, and which is sampled independently for each *t* and subpopulation *α*. We observe that *ω*_*α*_(*t* + *h*) is equivalent to a zero mean Gaussian with variance

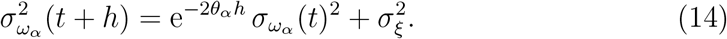

Once we choose 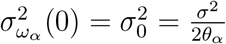,we observe that 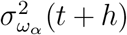 is time independent and it is equal to 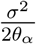.This allows us to generate *ω*_*α*_ (*t*) from the Gaussian distribution 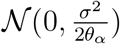 independently at each instant *t >* 0, avoiding explicit forward time simulations of the OU process given in (11). This is the idea behind introducing the noise terms *ω*_*α*_ in the mean-field system (1) and the noise term ζ_*i*_ appearing in the regulatory mechanism (6) as Gaussian white noise sampled directly from a Gaussian distribution at each time and for each subpopulation, but not as an explicit OU process evolving in time.

### 4.1 Extracellular media distortion effects

In the previous AdEx mean-field model (di Volo et al., 2019), the background noise was introduced as an additive noise to the base drive potential *s*_AI_. This noise was evolving as an OU process and it modeled dynamically the distortion effects appearing in the firing rates due to the extracellular medium and synaptic perturbations. In our model, we introduce the white Gaussian noise *ω*_*α*_, explicitly in the firing rate equations as shown in (1). In this way, we avoid that the noise undergoes the nonlinear effects of the transfer functions since now it is additive in the model equations. This noise models the distortion effects of the extracellular media on the neural dynamics.

### 4.2 Exploratory behavior

One of the typical behaviors in the decision-making task is that the participant makes decisions which do not comply with the preset strategy even after the participant learned the strategy. This behavior occurs due to two reasons: (i) perceptual difficulty in the human experiment, i.e., the difference between stimuli is small, thus the participant makes the wrong decision as a result of perceptual difficulty; (ii) the participant hesitates or is willing to explore whatsoever related to the experiment setup, and this might happen in both human and macaque experiments.

The perceptual difficulty is due to the stimuli, and it does not require any additional mechanism to be represented in the model. On the other side, the exploratory will of the participant is a part of the cognitive framework. We model it via the Gaussian white noise ζ_*n*_ introduced in the regulatory mechanism (6). This noise is sampled independently at each time instant. The noise level *σ* is scaled by (*c*_0_*t*^2^), where *c*_0_ *>* 0. Here *t* denotes the time instant and it is reinitialized at the beginning of each trial. This scaling term introduces a strong initial perturbation to the competition between the L2/3 populations. This perturbation decays in time. This decay is needed, since, otherwise the noise could dominate the whole competition instead of perturbing only the initial bias. The parameter *c*_0_ determines how strong the initial perturbation is. The regulatory mechanism therefore, is not only for the online reward-driven learning of the preset strategy but also for providing a natural behavior which is open to making wrong decisions. This is similar to what was observed in both human and macaque experiments. This exploratory behavior appears as irregular and rather sparse in-cluster deviations in the performance index plots; see Figure 11a.

## 5 Quantification of behavior

Quantification of behavior is required to analyze the results and to make a comparison between the simulation and experiment results. For a quantification of the results, we need adequate metrics. These metrics should provide a global characterization of the learning phase and of the erroneous decisions which the subjects and the model make.

Subject behavior in the experiments was quantified in terms of the coherence between the choices of the subject and the preset strategy. We follow the same approach to quantify the model behavior in the simulations. This coherence is measured by using a metric called performance index, which is defined in (29).

In each episode, the maximum performance index refers to the case in which the subject makes the choice in coherence with the preset strategy in every trial of the episode. The maximum performance index value is 1. The minimum performance index refers to the case in which the participant makes the incoherent choice in every trial of the episode. The minimum performance index is 0. All other cases with partial coherence are scaled between 0 and 1; see (29).

For the quantification of performance indexes, we use two objects: initial cluster episode and in-cluster deviation (Figure 11a). These objects characterize fully the behavioral pattern of a subject. The behavioral pattern is represented in terms of the performance indexes plotted with respect to their corresponding episodes; see Figure 11a. In the performance index plots, initial cluster episode gives a measure of how quickly the subject captured the preset strategy. It characterizes the learning phase. In-cluster deviation gives a measure of erroneous decisions that the subject made after learning the preset strategy. These erroneous decisions can be due to several reasons, e.g., perceptual difficulties and curiosity to explore. Therefore, in-cluster deviations characterize the behavior after the learning was completed. We use these two objects for quantification of behavior. However, we need first, the following three definitions to be able to describe initial cluster episode and in-cluster deviation:

**Definition 1** (*Performance deviation*): A performance deviation is a point with a performance index lower than the maximum performance index (= 1), except for the first episode.

**Definition 2** (*Deviation cluster*): A deviation cluster is a set of at least 3 successive performance deviations with respect to the episode numbers.

**Definition 3** (*Performance cluster*): Performance cluster is the set of performance indexes in which there is no deviation cluster and whose last performance index is also the last index of the whole experiment.

Now we can define initial cluster episode and in-cluster deviation as follows:

**Definition 4** (*Initial cluster episode*): The first episode number of a performance cluster.

**Definition 5** (*In-cluster deviation*): A performance deviation is called *in-cluster deviation* if it occurs in a performance cluster. It is called *out-cluster* otherwise.

A performance cluster starts at the episode where the participant begins to make decisions in complete coherence with the preset strategy. Within the cluster, the participant makes coherent decisions almost in each episode until the end of the cluster. Almost; because the participant might make decisions in the aforementioned exploratory manner, producing in-cluster deviations. This does not break the performance cluster as long as in-cluster deviations do not constitute a deviation cluster. We refer to Appendix for a characterization of the model behavior in terms of performance indexes and reaction times; see Figures 11 and 12.

## 6 Model predictions

### 6.1 Human performance

The results of the simulations and experiments of the human case are quantified based on the mean and standard deviation of the initial cluster episodes (Definition 4), as well as the mean and standard deviation of the in-cluster deviations (Definition 5). This quantification fully describes the learning pattern of the model, as well as the behavior of the model after it captures the preset strategy. We obtain the mean values and standard deviations for each parameter set from 10 realizations of the simulation with the corresponding parameter set. Each realization is composed of 100 episodes.

We vary two parameters to change the performance behavior of the model: *k* in the reward-tracking module given by (8) and *c*_0_ in the regulatory module given by (6). The former is the learning speed and it determines how quickly the model learns the preset strategy, thus the initial cluster episode. The latter is the flexibility parameter. It determines how much the model is flexible to make mistakes, even after learning the preset strategy. These mistakes model the erroneous choices arising from perceptual difficulties, hesitation and curiosity/exploratory behavior of the subject.

We present the results of Horizon 1 simulations in Figure 5a. We observe that as *k* or *c*_0_ increases, the initial episode number of performance clusters decreases as shown on the left column. This is the case also in Horizon 0 as seen in Figure 5b. However, differently from Horizon 0, the decreasing curves in Horizon 1 are concave, meaning that the decrease rate of the mean initial episode number of performance clusters increases. On the contrary, we observe an increasing pattern in the mean of in-cluster deviations as *k* or *c*_0_ increases. This can be due to that, as the initial episode of the performance cluster decreases, there are more episodes in which the model makes erroneous choices. This generates in-cluster deviations. Therefore, it is likely that there will be more in-cluster deviations. An interesting observation is that this is not the case in Horizon 0 simulations. We predict that it is due to the difference in the task complexity. Since the task complexity is low in Horizon 0, the model makes fewer erroneous choices in the performance clusters once it learns the preset strategy. In all Horizon 1 simulations, we observe that the model can produce close results to the experimental measurements once *k* and *c*_0_ are chosen properly, for example *k* = 0.05 and *c*_0_ = 2.

**Fig 5.**
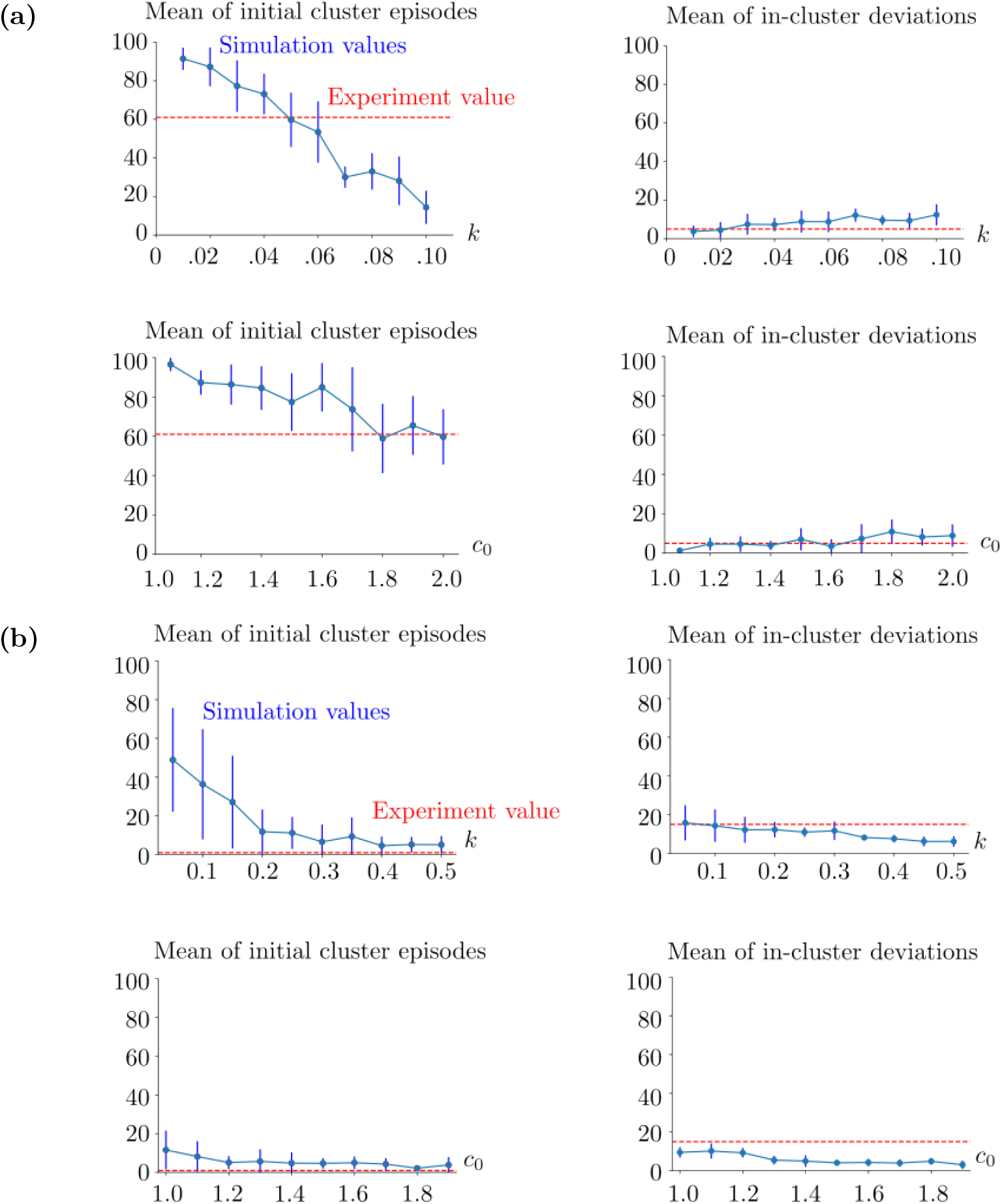
Simulation metrics with respect to the varied learning speed *k* and flexibility parameter *c*_0_. (a) Horizon 1 human case. On the top and bottom rows *k* and *c*_0_ are varied, respectively. In the top row, *c*_0_ = 2. In the bottom row, *k* = 0.05. The vertical blue lines show the standard deviations of the simulation results. The red horizontal lines indicate the initial cluster episode number (left column) and the number of in-cluster deviations (right column) obtained from the human experiment. (b) Horizon 0 human case. In the top row, *c*_0_ = 1.1 and *k* is varied. In the bottom row, *k* = 0.3 and *c*_0_ is varied. The blue vertical lines show the standard deviations of the corresponding statistical sample. The red horizontal lines indicate the experimental values as in (a). In both (a) and (b), the means and standard deviations are obtained from 10 realizations of the same simulation setup for each (*k, c*_0_) pair. The parameters *k* and *c*_0_ can be seen as rescaling constants, therefore they are unitless. See (23)-(25) for the rest of the parameters used in these simulations.

The results of Horizon 0 simulations are shown in Figure 5b. We observe in the left column that the learning speed decreases both with increasing *k* and with increasing *c*_0_, resulting in smaller values of the initial episode number of the performance clusters. The parameter *k* directly controls the learning speed, and *c*_0_ contributes to it by avoiding the deviation blocks which might break the performance clusters. In both cases, the decrease in the mean value profiles is convex. In all Horizon 0 simulations, we see that the model can reproduce close results once *k* and *c*_0_ are chosen properly, for example *k* = 0.3 and *c*_0_ = 1.1.

### 6.2 Human reaction times

In the experiments, the reaction time in one trial is measured as the time duration between the instant when the stimuli are shown simultaneously on the monitor, and the moment when the participant starts to move the pointer. In the simulations, we measure the reaction time as the time duration between the instant when the two stimuli are provided to the model and the time instant at which the decision is made, i.e., when the difference between the excitatory subpopulation firing rates 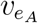 and 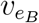 exceeds the decision threshold, which is fixed to 5 Hz.

In Figure 6a, we show the simulation and experiment histograms of the reaction times corresponding to Horizon 1, together with the fitted distributions for the best performance-providing *k* and *c*_0_ values among the tested ones. In each case, we fit the distributions of different types to both simulation and experiment histograms, and choose the distributions which give the smallest squared sum errors. We use the *Fitter* function from the Python *Fitter* package. We consider the histograms of the reaction times obtained from the first and second trials separately. In Trial 1, skewnorm distribution and hypergeometric distribution are fitted to the experiment and simulation histograms, respectively. Corresponding squared sum error values are 1.3e-5 and 9e-6 for the experiment and simulation histograms, respectively. Pearson correlation coefficient of these fitted distributions is 0.92. In Trial 2 histograms, skewnorm distribution and hypergeometric distribution are fitted to the experiment and simulation histograms, respectively. Corresponding squared sum error values are 1.7e-5 and 1.1e-5 for the experiment and simulation histograms, respectively. Pearson correlation coefficient of these fitted distributions is 0.63. Trial 2 has a weaker overlap between the experiment and simulation histograms compared to Trial 1.

**Fig 6.**
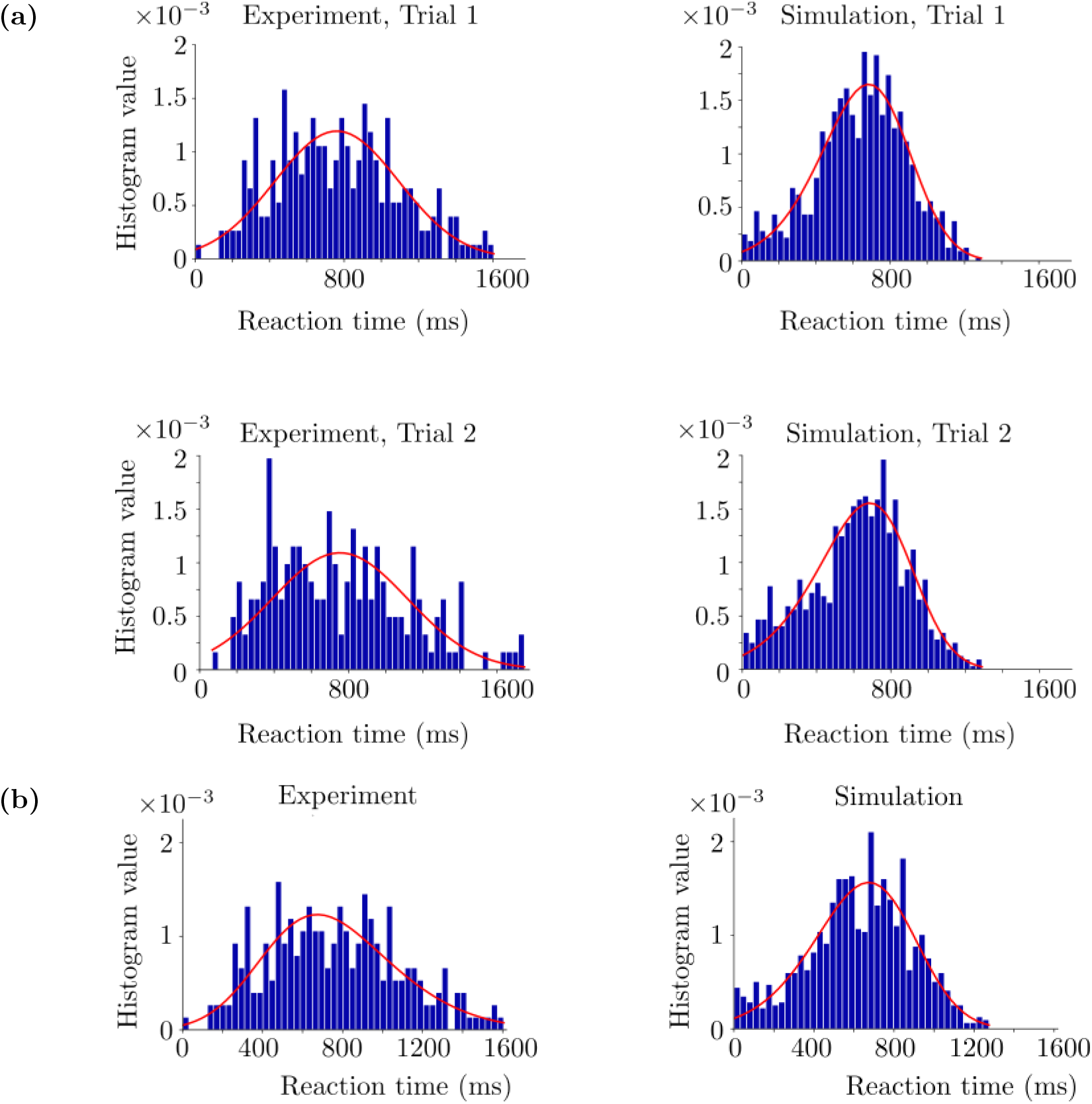
Reaction time histograms obtained from the human experiment and corresponding simulations. (a) Horizon 1. The red curves denote the fitted distributions via Python. Trial 1 and 2 histograms are in the top and bottom rows, respectively. The simulations are performed with *k* = 0.05, *c*_0_ = 2 and decision threshold = 5 Hz. (b) Horizon 0. The simulations were performed with *k* = 0.3, *c*_0_ = 1.1 and decision threshold = 5 Hz. See (23)-(25) for the rest of the parameters used in the simulations.

**Fig 7.**
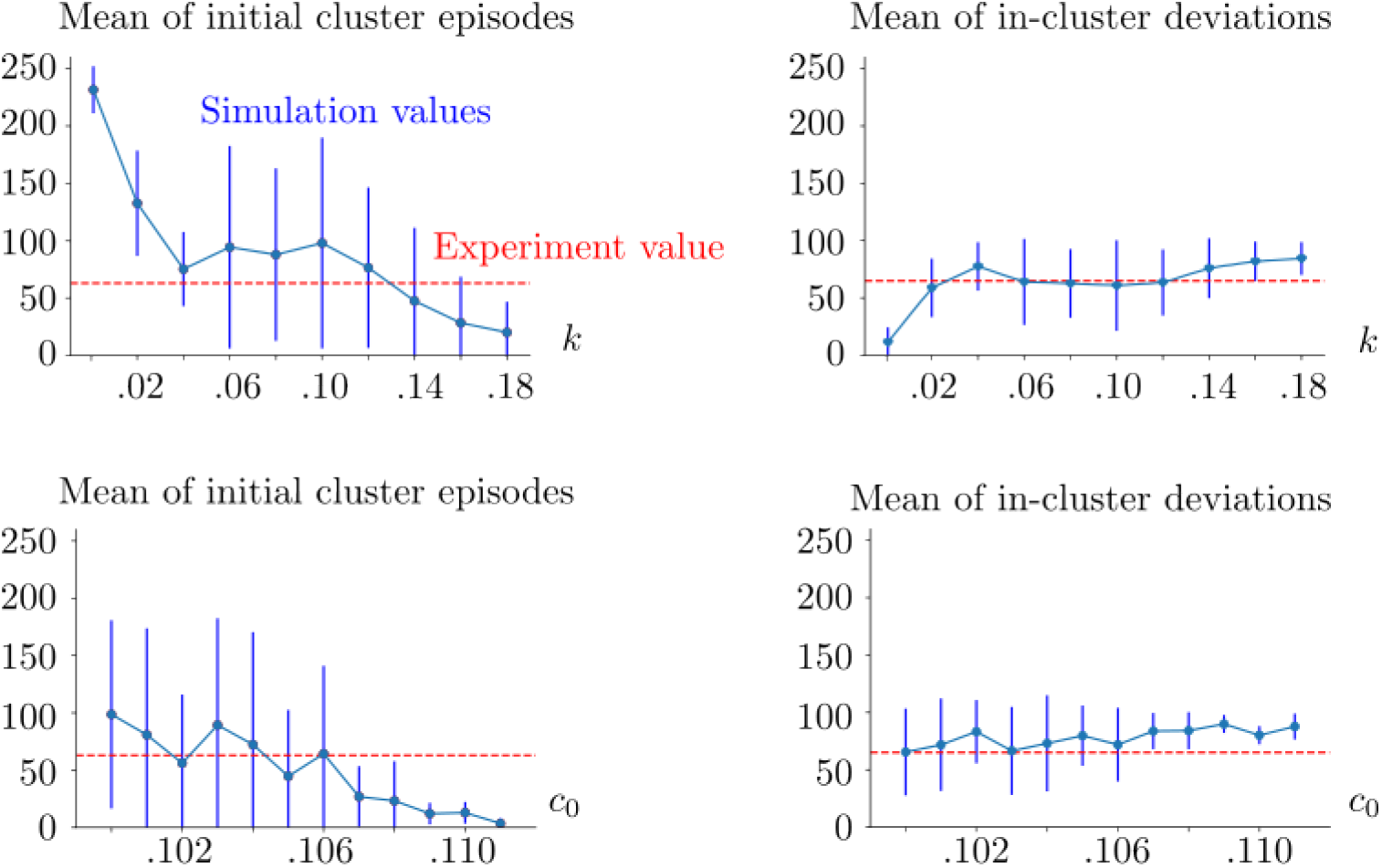
Simulation metrics of the macaque case. In the top row, *c*_0_ = 0.106, and *k* is varied. In the bottom row, *k* = 0.12, and *c*_0_ is varied. The vertical blue lines show the standard deviation of the corresponding statistical sample. The red dashed lines show the value obtained from the macaque experiment. The mean and standard deviations were obtained from 10 realizations of the same simulation setup for each (*k, c*_0_) pair. The decision threshold is 5 Hz. See (23)-(25) for the rest of the parameters used in the simulations.

In Figure 6b, we show the simulation and experiment histograms of the reaction times corresponding to Horizon 0 in the same fashion as the Horizon 1 results. The best fit is achieved with skewnorm distribution and hypergeometric distribution for the experiment and simulation histograms, respectively. Squared sum errors are 13e-6 and 12e-6 for the experiment and simulation histograms, respectively. Pearson correlation coefficient between the fitted distributions is 0.91.

The difference between the experimental and simulation reaction times could be due to several factors. In the experiments, reaction time includes the time between the instant when the participant makes the decision of which stimulus to choose and the instant when the participant starts to move the pointer towards the chosen stimulus. This adds a duration which may vary, in particular due to hesitation, fatigue, loss of motivation etc. The time duration between these two instants cannot be taken into account in the model, simply because there is no motor action in the model. Another reason could be the initial conditions. We do not have any access to the initial conditions of the neural dynamics of the participant. Therefore, we initialize the basic module from the same initial conditions at the beginning of each trial in our simulations. Nevertheless, it is likely that these conditions are different at the beginning of each trial in the experiments. These factors do not have strong effects on performance results, however they can change the results regarding the reaction times significantly.

### 6.3 Macaque performance

We observe that the trends in the macaque performance indexes are similar to the ones found in the Horizon 1 human task simulations. We see that the simulation results overlap with the macaque experiments for properly chosen parameters, for example *k* = 0.12, *c*_0_ = 0.106; see Figure 6a.

### 6.4 Macaque reaction times

In Figure 8, we show the simulation and experiment histograms of the reaction times corresponding to the macaque experiment, together with the fitted distributions. Here *k* and *c*_0_ are fixed to the values providing the best performance overlap. We fit the distributions to both simulation and experiment histograms by using the *Fitter* function from the Python *Fitter* package as in the case of the human experiment. In Trial 1, genlogistic distribution and powernorm distribution are fitted to the experiment and simulation histograms, respectively. Corresponding squared sum error values are 1e-5 and 9e-6 for the experiment and simulation histograms, respectively. Pearson correlation coefficient of these fitted distributions is 0.97. In Trial 2 histograms, burr distribution and kappa3 distribution are fitted to the experiment and simulation histograms, respectively. Corresponding squared sum error values are 9.9e-5 and 5.73e-4 for the experiment and simulation histograms, respectively. Pearson correlation coefficient of these fitted distributions is 0.31. Trial 2 has a weaker overlap between the experiment and simulation histograms compared to Trial 1 as in the case of the human task.

**Fig 8.**
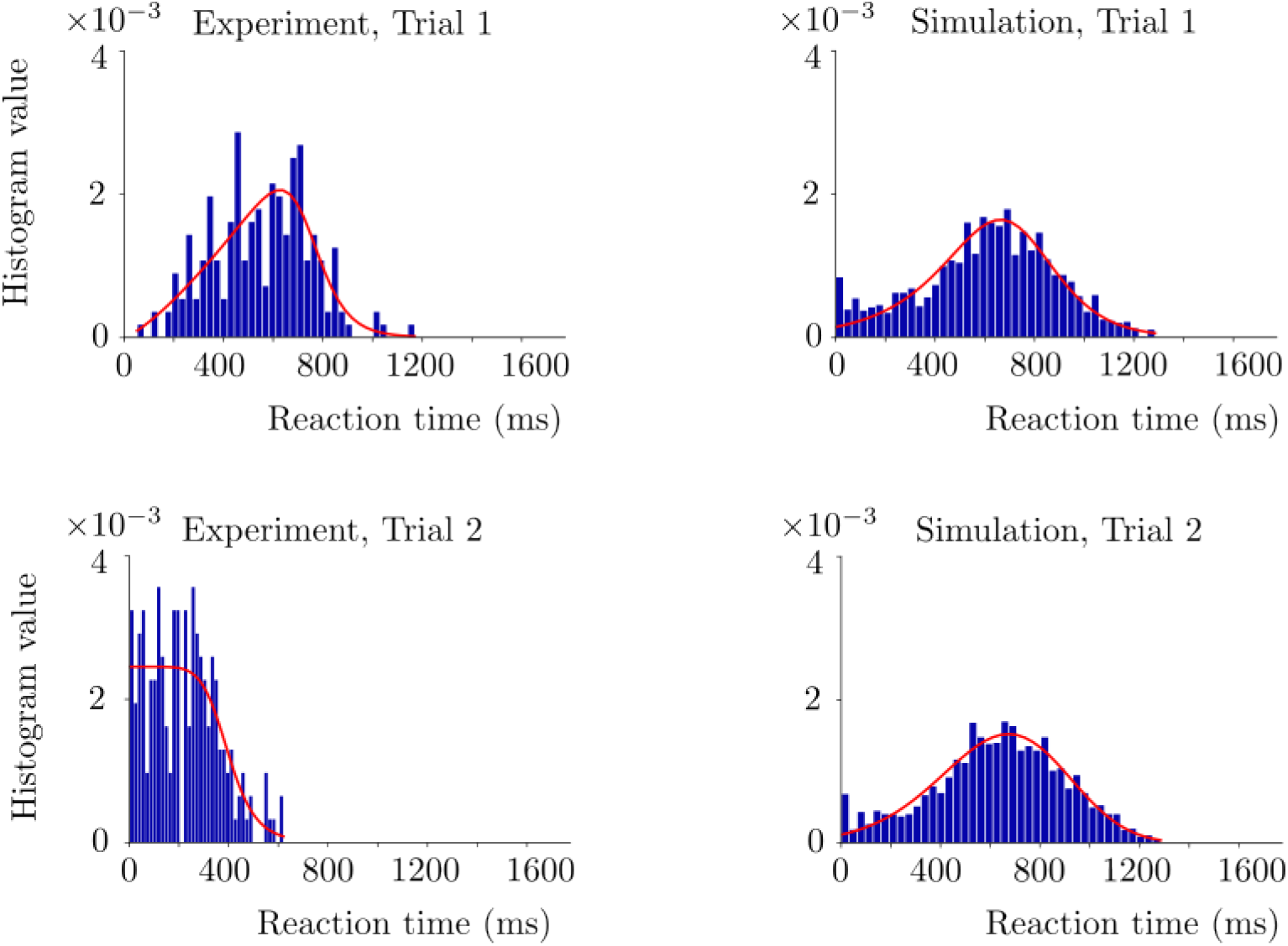
Reaction time histograms of the macaque case. The red curves denote the fitted distributions via Python. Trial 1 and 2 histograms are in the top and bottom rows, respectively. The simulations are performed with *k* = 0.12, *c*_0_ = 0.106 and decision threshold = 5. See (23)-(25) for the rest of the parameters used in the simulations.

The same factors regarding the reaction times and the initial conditions as in the human case are valid also in the macaque case. We predict that the difference between the experimental and simulation results regarding the reaction times arises from these factors; see Section 6.2.

## 7 Discussion

We presented a biologically plausible AdEx mean-field model based on two interacting L2/3 neural populations. The model is composed of three modules: basic, regulatory and reward-tracking modules. The basic module is based on the AdEx mean-field equations which we extended from one population (di Volo et al., 2019) to two populations. Each population is composed of a pair of excitatory-inhibitory subpopulations. The basic module generates the firing rates of the subpopulations. The excitatory subpopulations are in competition with each other to make a decision. The winning subpopulation makes the decision. The regulatory module introduces a bias into this competition such that one of the subpopulations becomes more likely to win the competition. Therefore, it makes one of the two stimuli more likely to be chosen. The bias depends on the increase or decrease in the reward amounts. The reward-tracking module keeps the track of these increases and decreases in the reward amounts. It transmits the information regarding these increases and decreases between successive episodes. We tested this model on two complex behavioral decision-making tasks, which were performed on human and macaque.

The model is biologically plausible since its connectivity architecture overlaps with an anatomically coherent case and the neural transfer functions are obtained based on *in vitro* measurements as well as neural responses obtained from the AdEx neuronal network. The novelties of the model are at both connectivity and structure levels.

Regarding the connectivity, the L2/3 populations are assumed to communicate via excitatory connections, as observed in macaque (Hirata et al., 2008). Hence, the connections between the L2/3 populations are only excitatory. Two such connections originate from the excitatory subpopulation of one of the L2/3 populations. The first one of these connections is onto the excitatory subpopulation of the other L2/3 population. The second one is onto the inhibitory subpopulation of the other L2/3 population.

These connections overlap with the long range cortico-cortical connections found in the prefrontal cortex (PFC). The long-range cortico-cortical connections can be found between the lateral PFC and the limbic areas related to dopamine neurons, such as the medial PFC. The former was suggested to be responsible for the choices related to long-term planning independent of the delay of the reward, and the latter was considered for the choices related to impulsive decisions with short-term rewards (McClure et al., 2004). In our model, inhibitory control over the excitatory subpopulation is triggered by the increasing activity of the excitatory subpopulation of the other L2/3 population. This is not the case in the previous models (Wang, 2018; Marcos et al., 2013). In (Wang, 2018), there is only one inhibitory subpopulation which is connected to all excitatory subpopulations via the long range connections. In (Marcos et al., 2013), the excitatory and inhibitory subpopulations are not explicit. They are intrinsically represented as one neural population. Two such populations are connected to each other via inhibitory connections, and recurrently connected to themselves via excitatory connections.

On the structure side, we model each L2/3 population as the mean-field limit of a pair of excitatory and inhibitory neural subpopulations. This allows us to provide the stimuli to both excitatory and inhibitory subpopulations, which is not the case in previous models (Wang, 2018; Marcos et al., 2013; Cecchini et al., 2024). As a result, our model can be extended to also decision-making with multiple choices. Moreover, the parameters are directly linked to the biophysical mechanisms of RS and FS cells. Therefore, our model can be applied to not only behavioral data but also to relevant neurophysiological data, in particular to multi-site array recordings of the macaque cortex. In this way, the model has the potential to link the phenomenological decision-making models (Wang, 2018; Marcos et al., 2013; Cecchini et al., 2024) to the neural dynamics observed at mesoscopic level.

Moreover, we adapted the regulatory module given in (Cecchini et al., 2024) to the AdEx system to introduce the plasticity which learns the preset strategy in the task. This machinery is essential for the model to find the optimal decision-making strategy. The regulatory module sustains the optimal strategy once this strategy is learned. This is due to the fact that once the introduced bias provides increase in the rewards, the regulatory module reinforces and retains the bias. This can happen successively over episodes, relating the regulatory module to working memory at phenomenological level.

The model was tested at the behavioral level by comparing its predictions to the experimental data obtained from the human and macaque participants. The comparison was made based on three statistical metrics: mean value of initial episode numbers of the performance clusters, mean value of in-cluster performance deviations and reaction time distributions. It was found that the model reproduces several characteristics of the experiment results. To begin with, the model predicts closely the performance indexes in the cases of both human and macaque for properly chosen parameter sets. It is possible to produce a good variety of behavioral patterns by varying the learning speed and the flexibility parameter. In reaction times, the model and experiment results have a mismatch, in particular in Trial 2 measurements of Horizon 1 experiments in both human and macaque. A possible reason is that the motor action is included in reaction time measurements in the experiments, whereas it is not possible to consider the motor action in the model setup. Another possible reason is that the decision threshold in the model is fixed. It is not dynamically adapted over the episodes as the strategy is learned.

Our work is linked to (di Volo et al., 2019; Cecchini et al., 2024). It uses the experimental data obtained from human and macaque subjects to provide the metrics based on performance indexes and reaction times. The data was given in (DePass et al., 2023; Fontana et al., 2022). Our work builds upon (di Volo et al., 2019) by extending the AdEx mean-field equations from a single pair, to two pairs of excitatory-inhibitory subpopulations. The extension relies on the excitatory connections between the L2/3 populations. Moreover, we integrate the regulatory and reward-tracking modules with this extended AdEx framework. These modules introduce a learning mechanism to the model, which was not considered previously in (di Volo et al., 2019). Our work builds upon (Cecchini et al., 2024) in terms of the basic module and the reward-tracking module. The basic module in our framework elaborates several biophysical details as a result of the use of AdEx equations. These details were not explicit in (Cecchini et al., 2024). Our model considers both human and macaque cases, with the two modes ((8) and (9)) in the reward-tracking module.

Our model was designed for the optimal decision-making. Yet, it has the potential to be extended to the sub-optimal decision-making. Optimal decision-making refers to making choices which provide the maximum possible reward (Doya et al., 2007). Albeit the absence of a precise definition, sub-optimal decision-making refers to the decision-making which deviates from the optimal decision-making in terms of performance indexes. The model already reproduces partially sub-optimal behavior in the performance clusters thanks to its exploratory behavior as shown, for example, in Figure 11d. This property can be improved: in the regulatory module, we can introduce a term which weights the flexibility parameter based on the changes in the reward amounts or in the motivation of the participant to conduct the task. This can be combined with the decision thresholds which are dynamically adapted throughout the episodes. This could project the changes in the motivation of the participant, on the performance indexes. This modification could provide a better understanding of the neural dynamics which modulate the behavioral strategies of the participant in accordance with the increase or decrease in the motivation. This has been an active research area in the experimental side (Leotti et al., 2010; Giamundo et al., 2021; Padmala et al., 2010), and it can benefit enormously from the computational side.

Another interesting future research direction could be related to the brain states.

The AdEx mean-field framework can simulate, in addition to the AI state, several other brain states such as sleeping, drowsiness or under anesthesia states (di Volo et al., 2019; Goldman et al., 2020). This property of the AdEx framework can be integrated into decision-making context as well. This can be interesting to study for example the effect of different levels of drowsiness on the capacity to perform cognitive tasks, see for example Figure 9. The figure shows the gradual deterioration of our model’s performance in a simulation of Horizon 1 decision-making task. In the simulation, our model goes through a transition from the awake state towards under anesthesia state. Studies in this direction might provide computational support for better understanding the effects of anesthesia or drugs on cognitive abilities.

**Fig 9.**
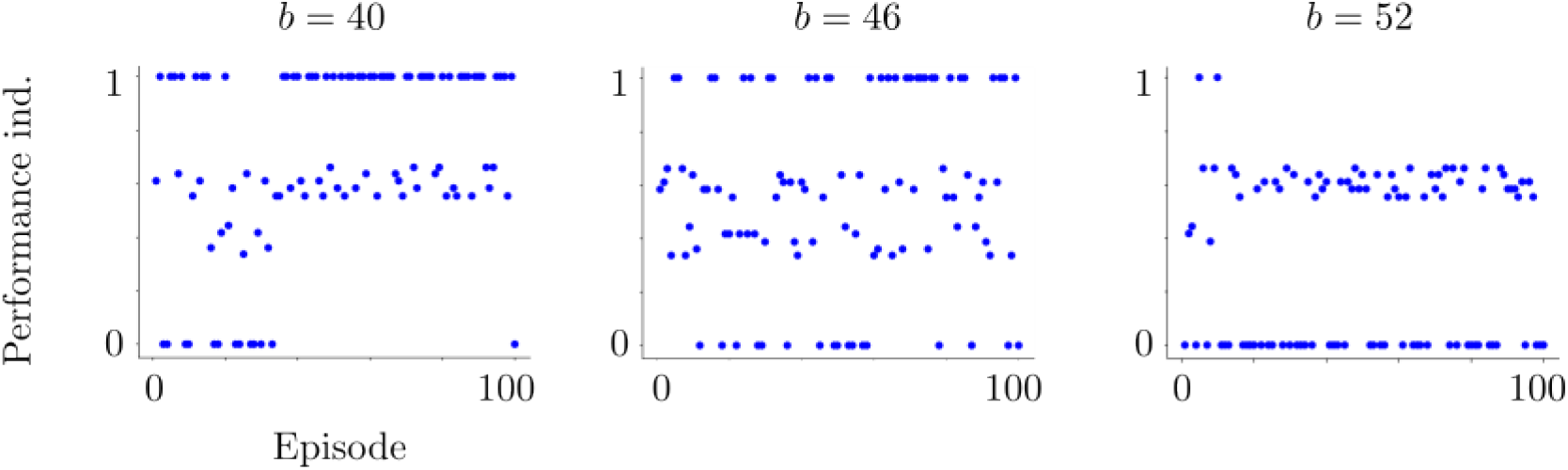
Performance index plots of the simulations of the model as the model goes through the transition from the awake state towards the sleeping state. The spike-frequency adaptation of the RS cells becomes stronger as we increase the parameter *b* found in the adaptation term of (1). This pushes the model towards the sleeping state. We observe that the model loses gradually the capacity to obtain maximum performance indexes, i.e., it loses gradually the ability to learn the preset strategy.

Finally, a further extension of the model can be towards the large-scale brain dynamics of decision-making. For this, our AdEx mean-field framework can be integrated in The Virtual Brain (Sanz Leon et al., 2013; Goldman et al., 2020) to model the decision-making brain dynamics of the PFC, which has been traditionally considered as the key area involved in decision-making (Bechara et al., 2000; Hampton et al., 2006; Rushworth et al., 2011; Shadlen et al., 2013; Domenech et al., 2015; Funahashi et al., 2017). In this way, the perceptual inputs can be provided directly from the visual (or other perceptual) areas to the PFC. Consequently, the outputs of our model can be provided as the outputs of the PFC to the motor areas. Some preliminary results in this direction can be found in (Turan et al., 2023).

## Supporting information

Tex files

## Appendix

### 1 Neural transfer functions

Neural transfer functions of AdEx mean-field equations were provided in (Zerlaut et al., 2018) and generalized to also spike-frequency adaptation in (di Volo et al., 2019). These functions were obtained by a semi-analytical fitting to the averaged output of the AdEx neuronal network. The output of the network was induced by random spike bombardments generated from a Poisson distribution. We use the same transfer functions provided in (di Volo et al., 2019). We summarize here how their explicit formulas were obtained. For details, we refer to (Zerlaut et al., 2018; di Volo et al., 2019).

#### AdEx neuronal network

The AdEx mean-field of a pair of excitatory-inhibitory subpopulations with spike-frequency adaptation approximates the average dynamics of the AdEx neuronal network. The network is composed of a certain number of neurons. Each neuron in the network is described by the AdEx integrate-and-fire model (Brette et al., 2005):

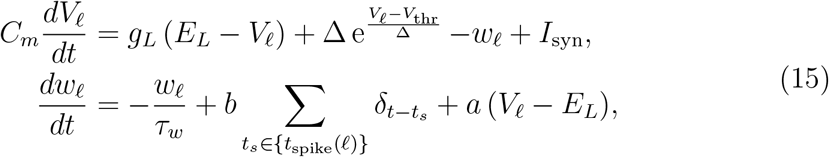

where *C*_*m*_ = 200 pF denotes the membrane capacity and *V*_𝓁_ is the membrane voltage of a neuron *𝓁* in the network. The voltage *V*_𝓁_ is reset to the resting voltage −65 mV at *t*_*s*_ ∈ {*t*_spike_(*𝓁*)}, which denote all the instants when the neuron generates a spike, i.e, when *V*_𝓁_ *> V*_thr_ = −50 mV. Here *g*_*L*_ = 10 nS and *E*_*L*_ =− 65 mV are the conductance and the leakage reversal of the leak term. Here Δ denotes the weight of the exponential term. It is 2 mV for RS cells and 5 mV for FS cells. Note that FS cells have no adaptation (*w*_𝓁_ = 0). Therefore, *a* = *b* = 0 for all inhibitory neurons. For RS cells, the adaptation is introduced with *a* = 4 and *b* = 40 in our simulations. Synaptic input to the neuron *𝓁* is defined as follows:

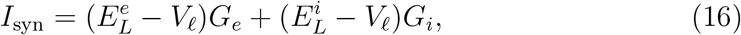

with 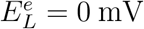 and 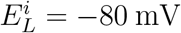 denoting the reversal potential of the excitatory and inhibitory presynaptic cells, respectively. The conductances are defined as

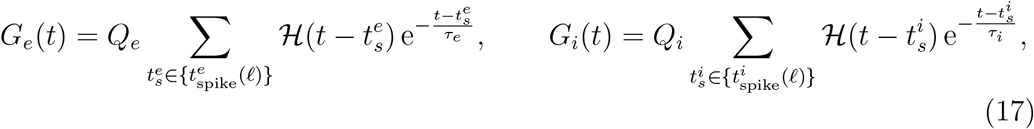

where ℋdenotes the Heaviside function and *τ*_*e*_ = *τ*_*i*_ = 5 ms are the decay rates of the excitatory and inhibitory presynaptic neurons. Here 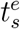 and 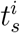 are the instants when the excitatory and inhibitory presynaptic neurons trigger spikes, respectively.

Finally, *Q*_*e*_ = 1.5 nS and *Q*_*i*_ = 5 nS are the excitatory and inhibitory quantal conductances, respectively.

#### Transfer functions

The classical AdEx mean-field equations were derived in (di Volo et al., 2019) from the AdEx network of neurons described by (15). The derivation of the mean-field equations was based on a master equation formalism which was provided in (El Boustani et al., 2009). The mean-field equations were for a single pair of excitatory-inhibitory subpopulations, with a spike-frequency adaptation. This is equivalent to decoupling the L2/3 populations of the basic module (1) of our model and considering only one of these decoupled populations. In that case, each L2/3 population reduces to a system of 6 state variables as in the classical mean-field system. Therefore, for the neural subpopulations in our model, we use the same transfer functions given for the classical AdEx mean-field system in (di Volo et al., 2019). The difference is that, the inputs to the transfer functions have not only the local components, but also the components related to the interactions between the L2/3 populations in our extended AdEx mean-field framework.

In the following, we will use the notation *e, i* to denote the excitatory and inhibitory subpopulations in either one of the decoupled L2/3 populations. The equations of the transfer functions are identical, independently of which L2/3 population we consider. These neural transfer functions were obtained via the semi-analytical derivation explained in (Zerlaut et al., 2018; di Volo et al., 2019).

The idea was based on the spike bombardment of a single neuron described by (15). The spikes were generated with various excitatory and inhibitory firing rates *v*_*e*_ and *v*_*i*_ which follow the Poisson statistics. This allowed to obtain the means 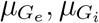 and the standard deviations 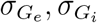 of the excitatory and inhibitory conductances in (17). These functions were found via Campbell-Hardy theorem (Papoulis et al., 2002) as follows:

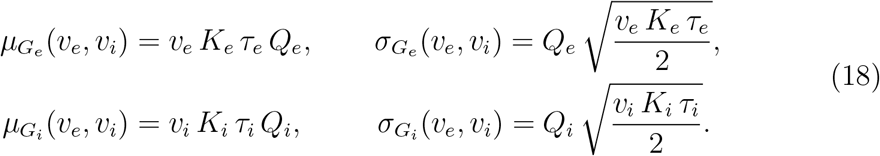

These functions control the input conductance *µ*_*G*_ and its effective membrane time constant *τ*_*m*_, which have the following expressions:

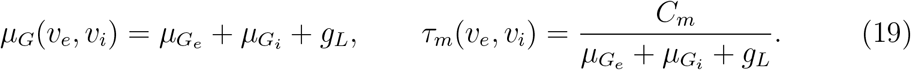

Consequently, mean membrane voltage of the neuron was obtained as:

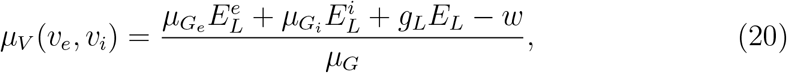

where *w* denotes the adaptation of the neuron. Finally, the standard deviation *σ*_*V*_ and the autocorrelation time *τ*_*V*_ of the membrane voltage fluctations were found from the power density of the fluctutations:

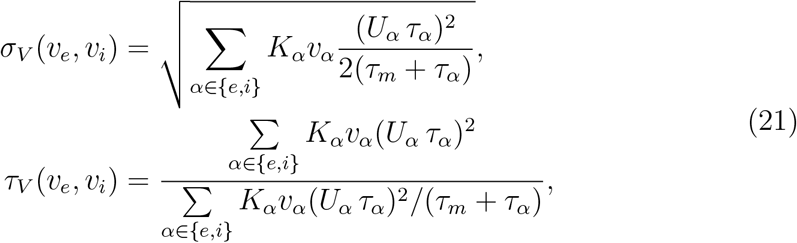

where 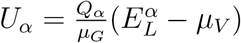.

In (21) and (22), we find the analytical expressions used in the transfer functions given by the formula (5). These analytical experessions are complemented by the voltage-effective threshold *v*_eff_, which was found by fitting the coefficients of the second order polynomial:

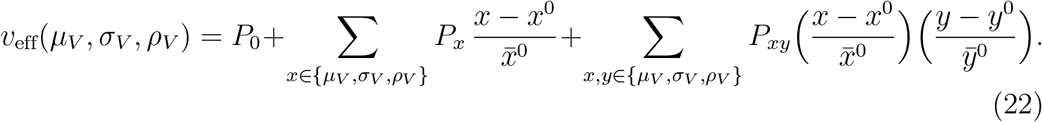

The fitting was made according to the simulations of single neuron activity described by (15). Here 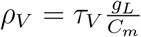 and 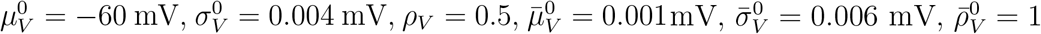. We use the same fitted coefficients as given in (di Volo et al., 2019). The fitting of the polynomial (22) and the functions in (20), (21) provide the transfer functions given by (5).

### 2 Simulation parameters

The parameters which are used in Figures 5-8 and Figures 11, 12 are as follows:

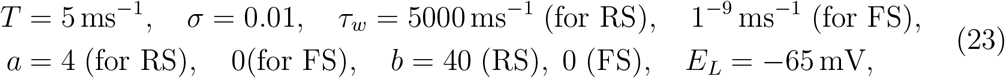

and it is given for (4) as

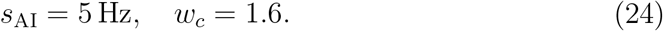

Finally, the parameters appearing in (6) and (7) are

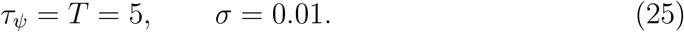

Each trial lasts 15 seconds in both Horizon 0 and 1 simulations. We set the decision threshold to 5 Hz in all the simulations.

### 3 Effects of varying the bias and the difference between the stimuli

The results in Figure 10 focus on the effects of varying the initial condition *ψ*(0) of the reuglatory module, therefore the effects of varying the bias. The decision threshold to measure the reaction time 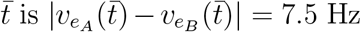.We use Euler-Maruyama scheme with time step Δ*t* = 0.5 and *t*_*f*_ = 4 for the presented results in Figure 10. Stimuli are applied at *t*_0_ = 2. The parameters in this framework are as follows:

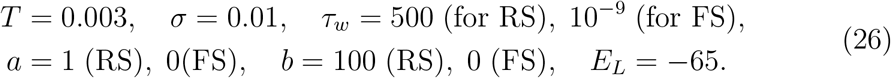

Moreover, in (4) we fix

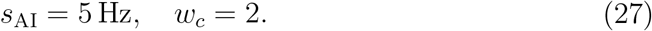

Finally, the parameters appearing in (6) are

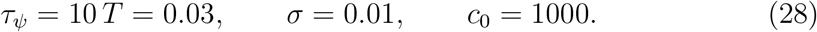

In Figure 10a, we provide the results of the cases with *ψ*(0) varied from 0 to 1, where the same stimuli were applied in each case. There is no clear difference in terms of reaction time between the cases with different initial conditions *ψ*(0) as long as the stimuli remain the same and *ψ* converges to the same value. It is due to the fact that, the convergence of the regulatory mechanism is rapid, therefore the competition is promoted towards the same L2/3 population. As we do not change the stimuli, the evolution of 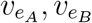 becomes different realizations of almost the same random processes. Consequently, the reaction times of those realizations fluctuate around the same value.

In Figure 10b, we show the effects of varying the difference between the stimuli on the model reaction time. The reaction time is measured in the same way as in Figure 10a. We set the amplitude of Stimuli *A* to 6 and vary the amplitude of Stimuli *B* between 0 and 6. We keep the initial value *ψ*(0) of the regulatory function constant (= 0.75) such that the system privileges always Stimulus *A*. We observe that the reaction time increases as the difference between the stimuli decreases. It is expected since it becomes more difficult to make a distinction between the stimuli as the difference between the filled quantities of the bars decreases.

**Fig 10.**
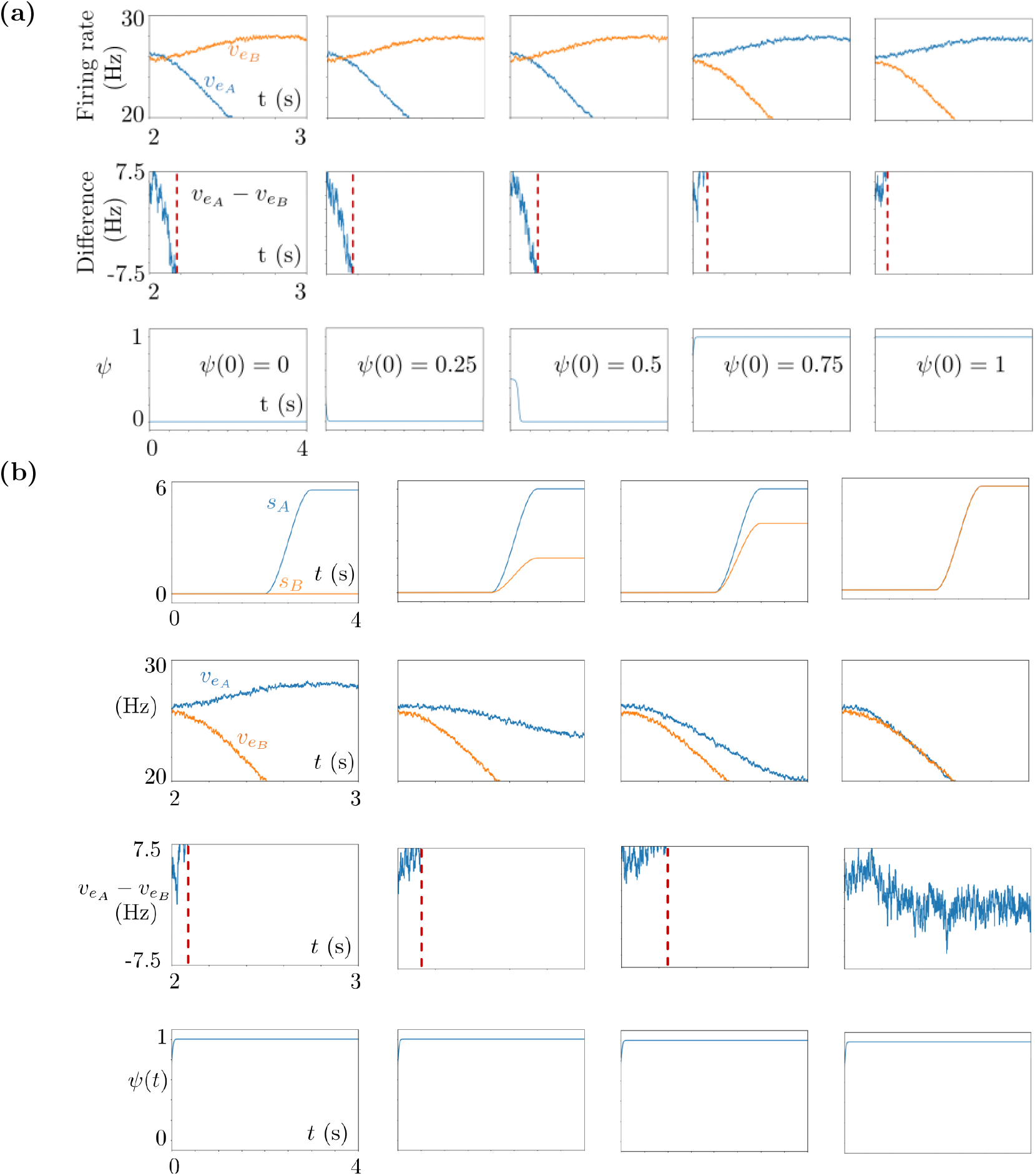
Simulation results of isolated trials, with varying regulatory module initial conditions *ψ*(0) in (a) and varying stimuli amplitudes in (b). The axes are specified only in the first columns in both panels, and they are identical in the other columns. **(a)** Top: Time courses of 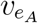 and 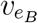.Middle: Difference of the firing rates where the red vertical line denotes the decision instant. Bottom: Time evolution of the regulatory function *ψ*(*t*). The initial values *ψ*(0) are 0, 0.25, 0.5, 0.75, 1 from left to right. **(c)** Top row: Time courses of the external stimuli *s*_*A*_ and *s*_*B*_. Second row: Time courses of 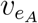 and 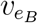.Third row: Time course of the difference between the membrane potentials where the red vertical line denotes the decision instant. Bottom row: Time course of the regulatory function *ψ*. The initial value *ψ*(0) is 0.75 in all plots.

### 4 Characterization of the model behavioral performance

Learning speed determines how quickly the model learns the preset strategy. The higher it is, the faster the model identifies the strategy. Flexibility parameter determines how much the model deviates from the preset strategy after it learns the strategy. It models the exploratory behavior and the deviations caused by perceptual difficulties. Finally, the gain rescales the learning speed. Here we provide the effects of these three parameters on the model behavior performance. We show in Figure 11b the performance index plots (human case) in which the learning speed *k* appearing in (8) varies from 0.1 to 0.3. We denote by 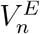 the value of the chosen stimulus in the *n*^th^ trial of the *E*^th^ episode. We denote the maximum and minimum cumulative reward values of episode *E* by 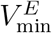 and 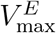, respectively. This is equivalent to maximum and minimum values that 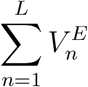 can attain among all possible scenarios of the *E*^th^ episode, with *L* denoting the number of trials in the episode. The performance index of the model in the *E*^th^ episode is computed as

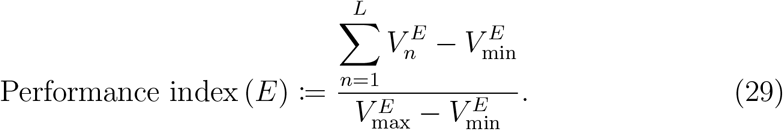

We observe in Figure 11b that as we increase *k*, the system captures the strategy earlier, and makes constantly right decisions thereafter, with very few deviations. The gain *G* is another factor affecting the learning speed. The learning speed decreases as the gain decreases since the gain *G* is equal to the multiplying factor 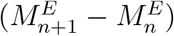 of the learning speed *k* in (8). As shown in Figure 11c, the model learns the preset strategy faster as we increase the gain.

Finally, the flexibility parameter *c*_0_ determines the number of deviations in the model performance results. The higher it is, the closer to the deterministic case the model is. This is due to the fact that the noise given in (6), and which produces the performance deviations, vanishes very quickly after that the regulatory variable *ψ* starts to evolve at the beginning of each trial; see Figure 11d. Moreover, *c*_0_ determines the beginning of the performance cluster together with the learning speed. As *c*_0_ increases, the performance cluster is more likely to begin from smaller episode numbers.

### 5 Characterization of the model reaction time

In Figure 12, we provide the histograms of the reaction time measurements obtained from the Horizon 1 simulations (human case). In these simulations, we varied the learning speed *k* and the flexibility parameter *c*_0_, separately in Trial 1 and Trial 2. We observe that there is no distinguishing change in the histograms of the simulation results, suggesting that these two parameters do not have noticeable effects on the reaction times. This is due to the fact that the reaction times are calculated by checking the threshold crossing of the difference between the firing rates 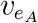 and 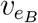.Here *k* and *c*_0_ do not enter in the equations that describe the firing rates, so they do not influence the reaction times. As the decision threshold increases, the competition between 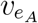 and 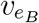 lasts longer, thus the reaction times increase as shown in the bottom row of Figure 12.

**Fig 11.**
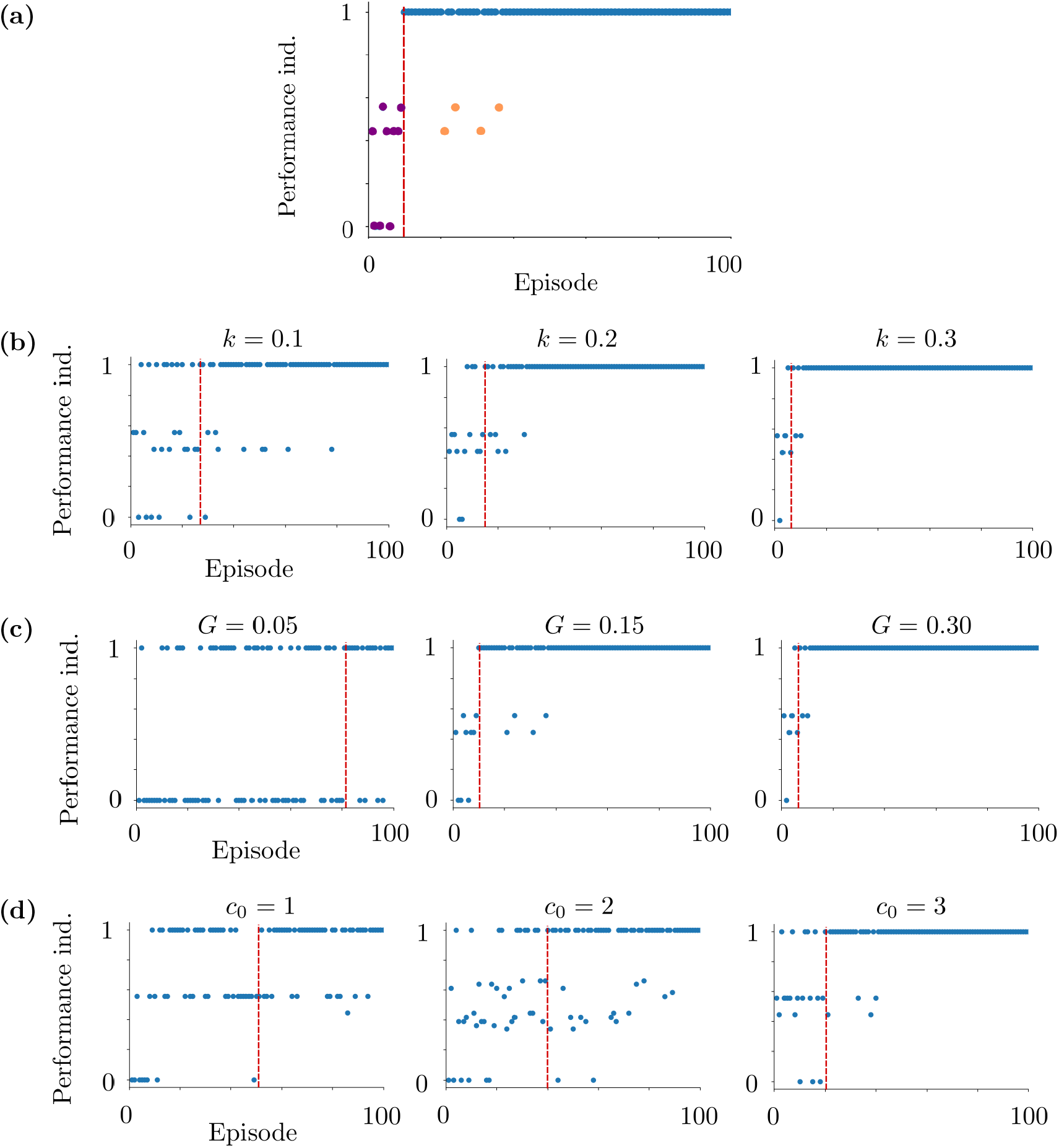
Objects used for quantification of behavior in (a), and performance indexes regarding the Horizon 1 human task simulations in (b), (c) and (d). **(a)** Purple samples correspond to a deviation block, and they are out-cluster deviations. Orange samples are in-cluster deviations, they do not form a deviation block. Both the purple and orange samples are performance deviations. Blue samples are maximum performance indexes and they form a performance cluster. The vertical red line marks the initial cluster episode, it corresponds to the beginning of the performance cluster. **(b)** Learning speed *k* varies. The flexibility parameter *c*_0_ is 2 and the gain *G* is 0.3. The initial episodes of the performance clusters are episodes 24, 15 and 5 from left to right. **(c)** *G* = 0.05, *G* = 0.15 and *G* = 0.3 from left to right. Here *k* = 0.3 and *c*_0_ = 2. The initial episodes of the performance clusters are episodes 81, 10 and 5 from left to right. **(d)** *c*_0_ = 1, *c*_0_ = 2 and *c*_0_ = 3 from left to right. Here *k* = 0.05 and the gain *G* = 0.3. The initial episodes of the performance clusters are episodes 52, 40 and 20; the number of in-cluster deviations are 12, 5 and 4 from left to right. The difficulty *d* is 0.2 in (b), (c) and (d).

**Fig 12.**
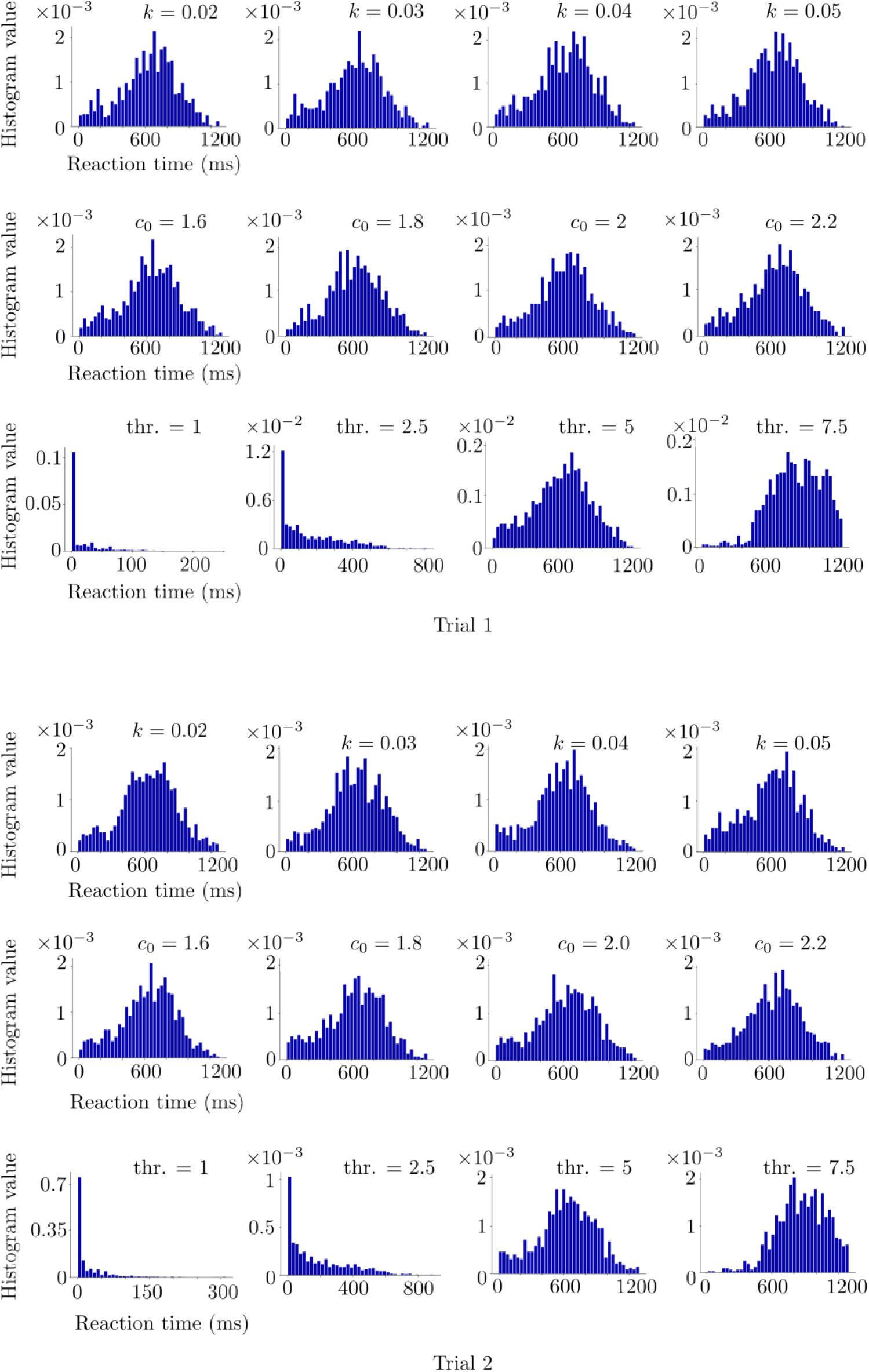
Simulation results regarding Horizon 1 reaction time histograms of Trials 1 and 2. Varying the learning speed *k* and the flexibility parameter *c*_0_ does not affect noticeably the distribution of the reaction times in terms of mean and standard deviation. Increasing the decision threshold increases the mean and the standard deviation of the distribution. The histograms are obtained from 10 realizations of the same simulation setup for each *k, c*_0_ and decision threshold.

## Acknowledgments

We hereby acknowledge that this research was supported by the Human Brain Project (European Union grant H2020-945539).

## Notes

https://doi.org/10.5281/zenodo.7682309

### Competing Interest Statement

The authors have declared no competing interest.

### Summary of Updates

- Major rewriting of the text - Introduction, Experiment setup, Model, Quantification of behavior sections are revised - "Neural transfer functions" section is added to Appendix

