## Supplementary material for "A biologically plausible decision-making model based on interacting neural populations": Tex files: main.pdf

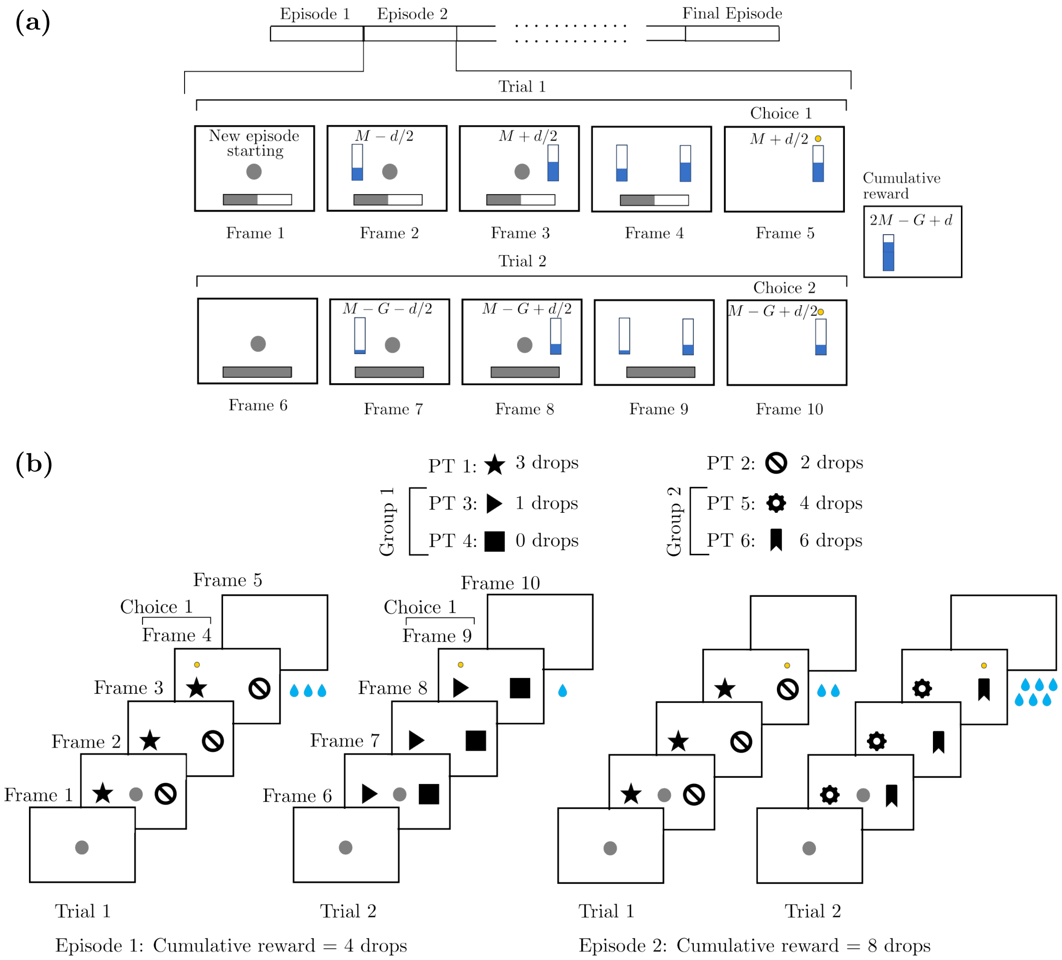


**Fig 1.** Example episodes from the Horizon 1 human and macaque experiments shown in (a) and (b), respectively. **(a)** A central target (CT) appears in Frame 1 to indicate the beginning of the episode. A progress bar at the bottom indicates in which trial the participant is. The stimuli with values $M - d/2$ and $M + d/2$ are shown separately in Frames 2 and 3, then together in Frame 4. The participant moves the pointer to the larger stimulus as highlighted by the yellow dot in Frame 5. The end of the episode is indicated in Frame 6 with a dot at the center. The stimuli are decreased in Episode 2, as shown separately in Frames 7 and 8. They are shown together in Frame 9 and the participant chooses the larger one as highlighted in Frame 10. The cumulative reward is the sum of the chosen stimuli. **(b)** A CT appears in Frame 1 to indicate the beginning of the episode. The subject touches the CT to initiate the episode. The peripheral targets (PTs) are shown together on the both sides of the CT in Frame 2. The CT disappears as the participant starts to move its hand to choose one of the PTs as shown in Frame 3. The chosen PT (PT 1) is highlighted by the yellow dot in Frame 4. The corresponding reward (3 water drops) is provided, and in Frame 5, a blank screen is shown to indicate the end of Trial 1. The same procedure is followed in Trial 2 but now with the PTs of Group 1. In the second scenario, the cumulative reward is higher since PT 2 is chosen in Trial 1 and hence the Group 2 PTs, which provide a higher reward, are shown on the monitor in Trial 2. The nomenclature and corresponding rewards to the PTs are given at the top. The cumulative reward is the sum of the provided water drops.

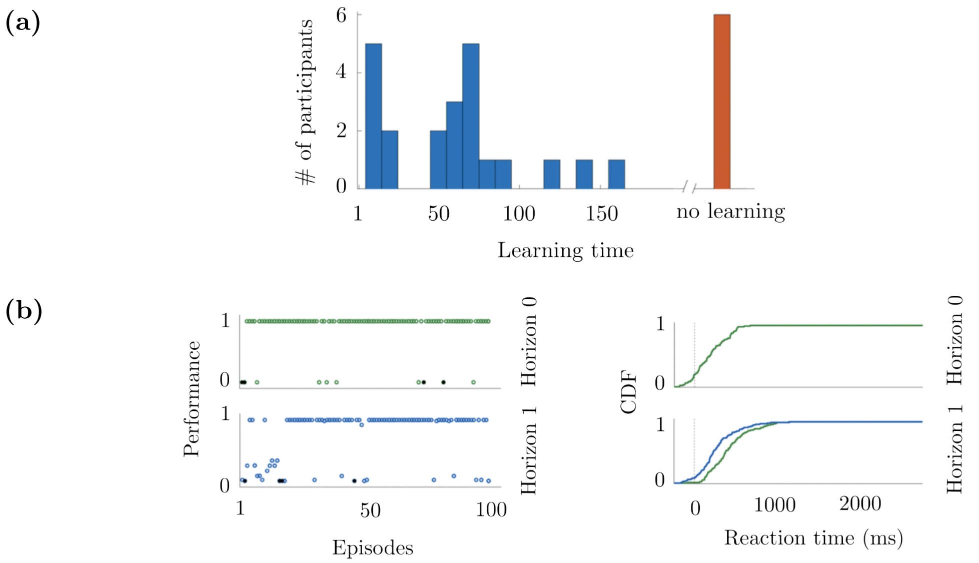


**Fig 2.** Some experiment results from (Cecchini et al., 2024). **(a)** The histogram of the learning times obtained from 28 human subjects. The learning time is in terms of episode numbers. The long learning time is classified as no learning. **(b)** The performance indexes of one of the participants, and the cumulative distribution function (CDF) of the reaction times of the same participant. The results are presented separately for Horizon 0 and Horizon 1, where in the latter, the CDFs of Trial 1 and Trial 2 are provided in green and blue, respectively.

We denote the firing rates of the four subpopulations by $v_{e_A}, v_{i_A}, v_{e_B}, v_{i_B}$. Each one of them represents the average spike frequency of one of the four subpopulations. We describe the neural interactions between these subpopulations via cross-correlations between the firing rates of interacting subpopulations. For simplicity of notation in the following, we use $\alpha, \beta, \xi, \eta$ to denote any of the four subpopulations $e_A, e_B, i_A, i_B$. Then, we represent the cross-correlation between $v_\alpha$ and $v_\beta$ with $C_{\alpha\beta}$. Finally, the variables $W_{e_A}$ and $W_{e_B}$ describe the spike-frequency adaptation of the RS cells, i.e., the excitatory subpopulations $e_A$ and $e_B$, respectively. The adaptation is only for the excitatory subpopulations as FS cells have no spike-frequency adaptation. Thus, we assume no adaptation for the inhibitory subpopulations, i.e., $W_{i_A} = W_{i_B} = 0$. As a result, the basic module has 16 state variables.

The basic module equations read as:

$$
\begin{aligned}
T\,\partial_t v_\alpha &= (F_\alpha - v_\alpha) + \frac{1}{2}\sum_{\xi,\eta} C_{\xi\eta}\partial_{\xi\eta}F_\alpha + \sigma\,\omega_\alpha \\
T\,\partial_t C_{\alpha\beta} &= A^{-1}_{\alpha\beta} + (F_\alpha - v_\alpha)(F_\beta - v_\beta) + \sum_{\xi,\eta}(C_{\beta\xi}\partial_\xi F_\alpha + C_{\alpha\xi}\partial_\xi F_\eta) - 2C_{\alpha\beta} \\
\partial_t W_{e_A} &= -\frac{W_{e_A}}{\tau_w} + b\,v_{e_A} + a\Big(\mu_V(v_{e_A}, v_{e_B}, W_{e_A}) - E_L\Big), \\
\partial_t W_{e_B} &= -\frac{W_{e_B}}{\tau_w} + b\,v_{e_B} + a\Big(\mu_V(v_{e_A}, v_{e_B}, W_{e_B}) - E_L\Big),
\end{aligned}
\tag{1}
$$

where we use the notation given by $\partial_t := \frac{\partial}{\partial t}$, $\partial_\alpha := \frac{\partial}{\partial v_\alpha}$ and $\partial_{\alpha\beta} := \frac{\partial^2}{\partial v_\alpha \partial_{v_\beta}}$ for the partial derivatives. Here $E_L$ is a constant representing the reverse leakage potential, and $T, \tau_w$ are the time scale parameters. We denote by $\omega_\alpha$ a white Gaussian noise, with $\sigma > 0$ denoting its intensity. This noise is sampled from a normal distribution independently for each subpopulation and at each instant. More precisely, $\mathbb{E}[\omega_\alpha(t)] = 0, \quad \mathbb{E}[\omega_\alpha(t)\omega_\beta(t')] = \delta_{\alpha\beta}\,\delta_{tt'}$ for all $\alpha, \beta \in \{e_A, i_A, e_B, i_B\}$ and for all $t, t' \geq 0$, where $\delta$ is the Dirac delta function. Here $\mathbb{E}[\,\cdot\,]$ represents the expected value. Function $A_{\alpha\beta}$ appears from the mean-field derivation of the second order statistical moments and it is defined for $\alpha, \beta \in \{e_A, i_A, e_B, i_B\}$ as follows:

$$
A_{\alpha\beta} = \begin{cases} \frac{T\,N_\alpha}{F_\alpha - T\,F_\beta}, & \text{if } \alpha = \beta \\ 0 & \text{otherwise,} \end{cases}
\tag{2}
$$

where $N_\alpha$ denotes the number of neurons represented in the subpopulation $\alpha$. We choose $N_{e_A} = N_{e_B} = 8000$ and $N_{i_A} = N_{i_B} = 2000$.

An important group of functions appearing in the mean-field equations (1) is the transfer functions, denoted by $F_\alpha$. A transfer function gives the output firing rate of a subpopulation according to the input that the subpopulation receives. This output firing rate is modulated by the spike-frequency adaptation in the excitatory subpopulations. The transfer functions appearing in (1) and (2) are written with their inputs as follows:

$$\begin{aligned} F_{e_A} &= F_{e_A}(\tilde{v}_{e_A}, \tilde{v}_{i_A}, W_{e_A}), \qquad & F_{i_A} &= F_{i_A}(\tilde{v}_{e_A}, \tilde{v}_{i_A}, W_{i_A} = 0), \\ F_{e_B} &= F_{e_B}(\tilde{v}_{e_B}, \tilde{v}_{i_B}, W_{e_B}), \qquad & F_{i_B} &= F_{i_B}(\tilde{v}_{e_B}, \tilde{v}_{i_B}, W_{i_B} = 0), \end{aligned} \tag{3}$$

with subindexes indicating the subpopulation type. Each input to the transfer functions is composed of excitatory and inhibitory components $\tilde{v}_{e_A}, \tilde{v}_{e_B}$ and $\tilde{v}_{i_A}, \tilde{v}_{i_B}$, respectively. Each component has a local part and a part related to the interactions between the L2/3 populations. The latter is weighted by $w_c$ since the interactions take place through the connections between the L2/3 populations. These excitatory and inhibitory components are as follows:

$$\begin{aligned} \tilde{v}_{e_A}(t) = v_{e_A}(t) &+ s_{\mathrm{AI}} + \lambda^A(s_A(t), s_B(t)) \\ &+ w_c\Big(v_{e_B}(t) + s_{\mathrm{AI}} + \lambda^B(s_A(t), s_B(t), t)\Big), \\ \tilde{v}_{i_A}(t) = v_{i_A}(t) &+ \lambda^A(s_A(t), s_B(t)) \\ &+ w_c\Big(v_{e_B}(t) + s_{\mathrm{AI}} + \lambda^B(s_A(t), s_B(t), t)\Big), \\ \tilde{v}_{e_B}(t) = v_{e_B}(t) &+ s_{\mathrm{AI}} + \lambda^B(s_A(t), s_B(t)) \\ &+ w_c\Big(v_{e_A}(t) + s_{\mathrm{AI}} + \lambda^A(s_A(t), s_B(t), t)\Big), \\ \tilde{v}_{i_B}(t) = v_{i_B}(t) &+ \lambda^B(s_A(t), s_B(t)) \\ &+ w_c\Big(v_{e_A}(t) + s_{\mathrm{AI}} + \lambda^A(s_A(t), s_B(t), t)\Big). \end{aligned} \tag{4}$$

Here the time dependency is explicitly denoted and the terms with no explicit time variable are constants. The external stimuli $s_A, s_B$ are fed in (4) via $\lambda^A, \lambda^B$. It is to take into account the bias introduced through the regulatory module, which will be explained in Section 3.2. A common choice to model the external stimuli $s_A, s_B$ is Heaviside function in mean-field models. However, since the Heaviside function has a discontinuity due to the discrete jump, our model undergoes a transient phase which might obscure the instant when the model makes a decision. For this reason, we perform our simulations by using a sufficiently sharp sigmoid function as external stimulus; see Figure 10b. Finally, $s_{\mathrm{AI}}$ denotes the DC input which keeps the basic module in the AI state.

We use the same formula for the transfer functions as provided in (di Volo et al., 2019); see Appendix for the details. We report here the formula directly:

$$F_{e_A}(\tilde{v}_{e_A}, \tilde{v}_{i_A}, W_{e_A}) = \frac{1}{2\,\tau_V(\tilde{v}_{e_A}, \tilde{v}_{e_B})}\ \mathrm{erfc}\Big(\frac{v_{\mathrm{eff}} - \mu_V(\tilde{v}_{e_A}, \tilde{v}_{e_B})}{\sqrt{2}\,\sigma_V(\tilde{v}_{e_A}, \tilde{v}_{e_B})}\Big). \tag{5}$$

$$\begin{cases} \tau_\psi \frac{d\psi_n^E(t)}{dt} = -4\psi_n^E(t)\Big(\psi_n^E(t) - 1\Big)\Big(\psi_n^E(t) - 1/2\Big) + \frac{\sigma}{(c_0 t)^2}\zeta_n(t), & t \in (0, t_F], \\ \psi_n^E(0) = \phi_n^{E-1}, \end{cases} \tag{6}$$

where $\tau_\psi$ is the time scale parameter. The regulatory function is restarted at the beginning of each trial. It evolves simultaneously with the basic module in time. Here $t_F > 0$ is the final time of the trial and it is the same for every trial. Moreover, $\zeta_n = \zeta_n(t)$ is a white Gaussian noise. Its intensity level $\sigma > 0$ is scaled by a constant $c_0 > 0$ and time. Therefore, this noise introduces a strong stochastic behavior initially. Then, it decays in time with a rate depending on $c_0$. It models the perceptual difficulties, hesitation and curiosity/exploratory behavior of the participant.

The decision in a trial depends on the evolution of the function $\psi_n^E$ in time. This function converges to either 1 or 0 in the course of each trial. The convergence towards 1 pushes the model to choose the stimulus bringing high reward, e.g., the large stimulus in the human experiment. The convergence towards 0 pushes the model to choose the stimulus bringing low reward, e.g., the small stimulus in the human experiment. These dynamics of the regulatory module are continuously transmitted to Populations $A$ and $B$ via

$$\begin{aligned} \lambda^A(s_A(t), s_B(t), t) =& \psi_n^E(t)\, s_A(t) + (1 - \psi_n^E(t))\, s_B(t), \\ \lambda^B(s_A(t), s_B(t), t) =& \psi_n^E(t)\, s_B(t) + (1 - \psi_n^E(t))\, s_A(t), \end{aligned} \tag{7}$$

where $t \in [0, t_F]$.

It is important to note that the process given by (6) is reinitialized at the beginning of each trial $n$ of the $E^{\text{th}}$ episode. It is reinitialized from the initial condition fixed to the value determined by the reward-tracking function $\phi_{n-1}^E$, which will be explained in the next section. The function $\phi_{n-1}^E$ corresponds to the same trial number $n$, but of the $(E-1)^{\text{th}}$ episode. In this way, the reward-tracking function provides at the end of the $(E-1)^{\text{th}}$ episode, a separate feedback for each trial $n$ of the $E^{\text{th}}$ episode. This feedback is used as the initial condition for the regulatory function equations (6), at the beginning of each trial of the $E^{\text{th}}$ episode. The feedback provides information about how optimal the decision-making strategy of the previous episode was, therefore about the accumulated experience. In this way, the feedback determines towards which decision the regulatory module will introduce the bias to the basic module in the next episode.

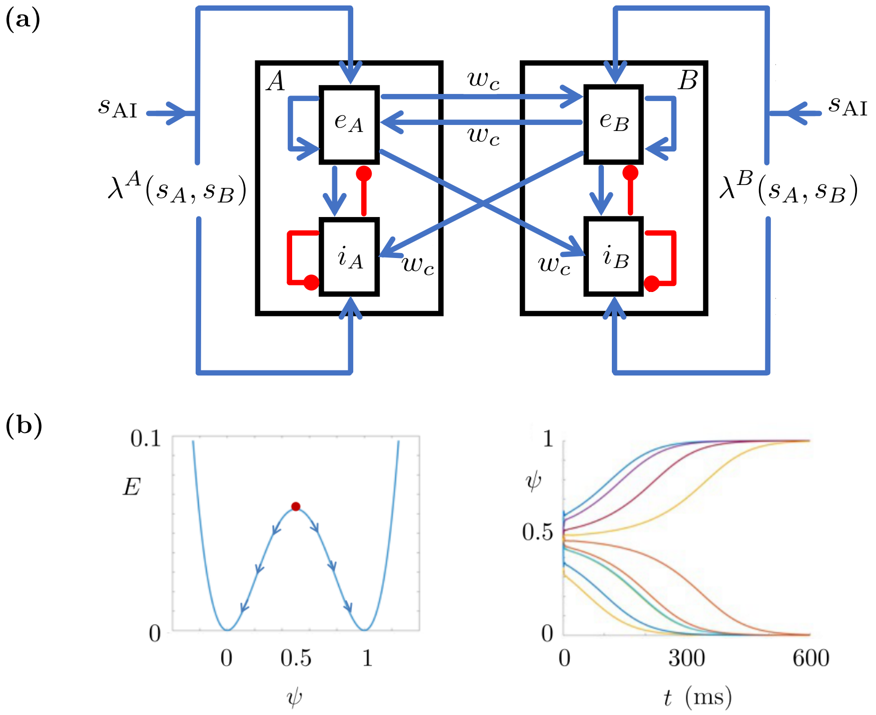


**Fig 3.** Model structures. **(a)** Basic module with two L2/3 populations composed of excitatory and inhibitory subpopulations. Populations $A$ and $B$ vote in favor of Stimulus $A$ ($s_A$) and Stimulus $B$ ($s_B$), respectively. The excitatory subpopulations are denoted by $e_A, e_B$ and the inhibitory subpopulations are represented with $i_A, i_B$. The excitatory and inhibitory connections are in blue and red, respectively. The weights of the connections between the L2/3 populations are denoted by $w_c$ as in (4). Here $\lambda^A$ and $\lambda^B$ represent the outputs of the regulatory module introducing a bias to the input signals $s_A$ and $s_B$. Finally, $s_{\mathrm{AI}}$ is the base drive, which ensures that the model performs in the awake state. **(b)** Regulatory module. Left: Energy functional of $\psi$. The neutral initial condition is $\psi(0) = 0.5$. The increase and decrease in the rewards introduce a bias making $\psi(0) > 0.5$ or $\psi(0) < 0.5$ in the next episode. This results in $\psi(t_F) = 1$ or $\psi(t_F) = 0$ at the end of the corresponding trial of the next episode. The former introduces the bias favoring the large stimulus and the latter introduces the bias favoring the small stimulus. Right: Time evolution of $\psi$ starting from different initial conditions. At the beginning, a small noise is introduced for modeling whatsover which might perturb the regulatory process, such as the hesitation, curiosity/exploratory behavior or perceptual difficulties.

##### 3.3 Reward-tracking module

The regulatory module introduces the bias depending on a feedback. This feedback is provided as the initial condition to the regulatory function equations given by (6). We should take into account the accumulated experience throughout the previous episodes to provide the feedback for the upcoming episodes. This feedback must be such that the model can identify which choices contribute more to the cumulative reward. We integrate our model with a reward-tracking module to endow the model with such feedback mechanism.

The reward-tracking module is based on a reinforcement learning equation. This equation motivates the choices which increase the reward amounts, and demotivates the choices which decrease the reward amounts. This is associated with the increasing dopamine signaling during predicted reward anticipation: motivation to repeat the same decision and the dopamine signaling increase, if the obtained reward is as predicted or more (Mirenowicz et al., 1994; Schultz, 2022). This results in an online learning. Once the preset strategy is learned, the system makes the decisions in almost complete coherence with the strategy.

We assume in the human experiments that the increase and decrease in the stimuli (i.e., reward) is predicted as profit and loss by the participants, respectively. In accordance with this assumption, the reward-tracking module is updated through a discrete evolution over the episode numbers. In other words, the reward-tracking function value remains constant during a trial and it is increased or decreased to another constant at the end of the trial. The updated value is provided to the regulatory function $\psi$ in the next episode as the initial condition; see (6) and (8).

In the human case, we use the notation $M_n^E$ to denote the mean value of the stimuli corresponding to the $n^{\text{th}}$ trial of the $E^{\text{th}}$ episode. We consider here only Horizon 1 since Horizon 0 uses the same module but with $n = 1$. We write the evolution for the reward-tracking function $\phi$ as follows:

$$\begin{cases} \phi_n^{E+1} = \phi_n^E + k\,(M_{n+1}^E - M_n^E)(2\psi_n^E(t_F) - 1)(\phi_n^E - 1)^2(\phi_n^E)^2, \\ \phi_n^1 = C, \quad \text{for all} \quad n \in \{1, 2\}, \end{cases} \tag{8}$$

with $C$ denoting a constant, which is fixed to 0.5 in our framework. Here $k$ is the learning speed parameter. It determines how quickly the model captures the preset strategy throughout the episodes. In (8), $(M_{n+1}^E - M_n^E)$ provides a coupling between the trials of an episode. It changes the sign of the polynomial. In this way, it determines how the bias provided by $(2\psi_n^E(t_F) - 1)$ will be transmitted to the upcoming episode (i.e., to the episode $E + 1$): by favoring or by disfavoring the choice made in episode $E$? The same choice is favored in the upcoming episode if $(M_{n+1}^E - M_n^E) > 0$, meaning that the stimuli (i.e., the rewards) increased as we pass from $n^{\text{th}}$ to $(n + 1)^{\text{th}}$ trial. It is disfavored if $(M_{n+1}^E - M_n^E) < 0$, meaning that the stimuli (i.e, the rewards) decreased.

In the reward-tracking module for the macaque case, we use a similar idea to the human case, with a few differences. Let us recall that in the macaque experiment, the reward is provided to the subject as water drops. This is different from the human experiment, where the reward is the chosen stimulus itself. Therefore, we write the reward-tracking equations for the macaque case, in terms of the water drops provided for each choice. We replace the first line of (8) with

$$\phi_n^{E+1} = \phi_n^E + k\,F_n\,(2\psi_n^E(t_F) - 1)(\phi_n^E - 1)^2(\phi_n^E)^2, \tag{9}$$

where

$$F_n^E := (\delta_{n-1}(\text{other}_1^E - \text{choice}_1^E) + \delta_{n-2}(\text{choice}_2^E - \text{other}_2^E)), \tag{10}$$

with $n$ and $\delta$ denoting the trial number and the Dirac delta, respectively. Here $\text{choice}_n^E$ and $\text{other}_n^E$ denote the numbers of water drops corresponding to the chosen stimulus and to the other stimulus, respectively, in the $n^{\text{th}}$ trial of the $E^{\text{th}}$ episode. Here the reward is explicit. It is not represented in terms of the stimuli as in the human case.

#### 3.4 Example simulation of the model

We provide here a small example simulation of the model to give an insight about the functionality of the three modules. In this example, we consider a human Horizon 1 simulation with 6 episodes.

In the human Horizon 1 experiments, the preset strategy requires choosing the smaller stimulus (the bar which is filled less) in Trial 1 and the larger stimulus (the bar which is filled more) in Trial 2 of each episode. This preset strategy is the optimal strategy which the model should learn so as to have the maximum cumulative reward. In Figure 4, we show a small example with 6 episodes of our Horizon 1 model simulations regarding the human case. The reward-tracking module $\phi$ is initiated from 0.5 for both trials in Episode 1, with a noise perturbing this initial value at the beginning of each trial. We observe that the model explores the strategy in Episode 1. Its decision is completely random with no bias since $\psi_i^1(0) = 0.5$ for both $i = 1$ and $i = 2$. In Episode 2, the reward-tracking module introduces a bias in the regulatory function $\psi$ based on the feedback from Trial 1 and Trial 2 of Episode 1. The same procedure is repeated for Episode 3 but now based on the feedback from Episode 2, and so on. In Figure 4, we observe that the model learns the strategy in Episode 2. From Episode 2 onwards, it makes the choices in complete coherence with the preset strategy.

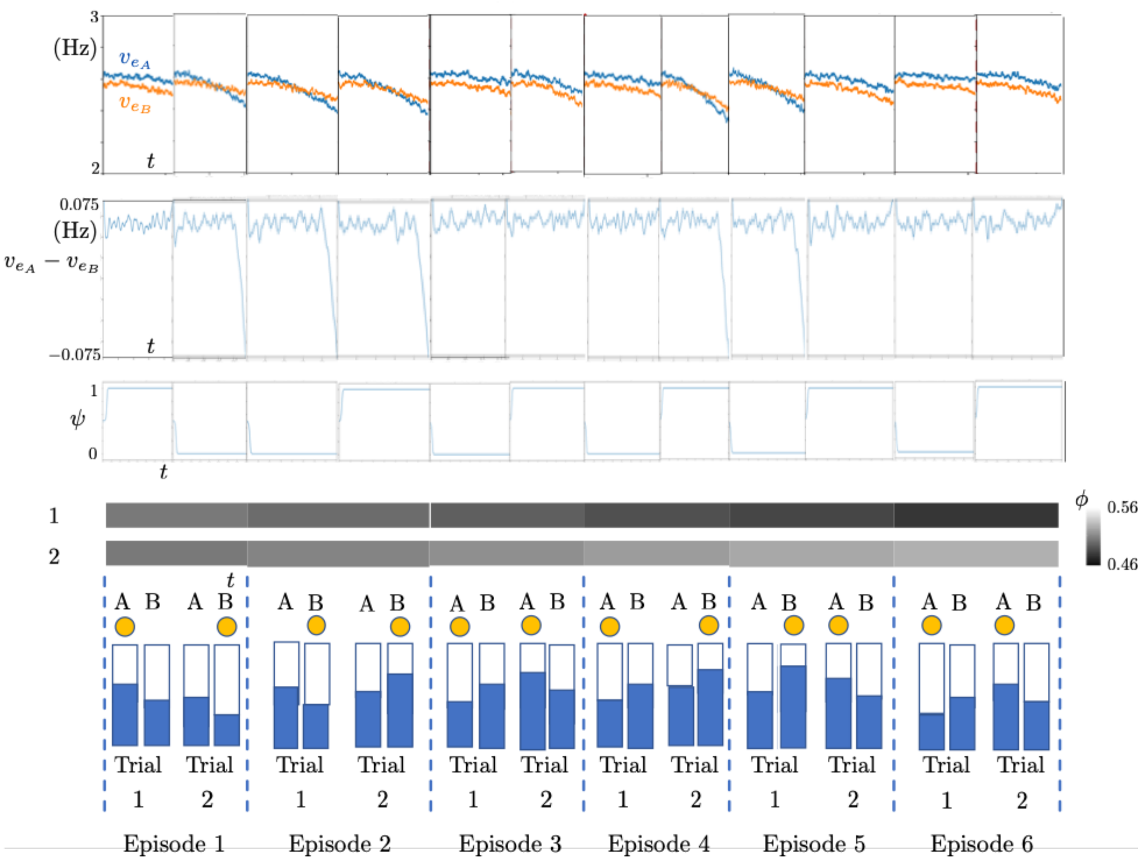


**Fig 4.** Simulation results of 6 episodes. First row: Time courses of the excitatory subpopulation firing rates. Second row: Time courses of the difference of the excitatory subpopulation firing rates. Third row: Time course of the regulatory function $\psi$ for each trial of the corresponding episode given in the bottom row. Fourth row: Time course of the reward function $\phi$ for Trial 1 (top) and Trial 2 (bottom) of the corresponding episode. Bottom row: External stimuli. Chosen stimuli are highlighted by the yellow dot at the top. Trial and episode numbers are given at the bottom.

### 4 Noise sources

Noise is required in our model to consider different perturbative mechanisms. It appears at two levels in the model: in the basic module and in the regulatory module. The former models the distortion effects due to the extracellular media. The latter models the exploratory behavior of the subject which arises from curiosity, hesitation and/or perceptual difficulties.

We describe the noise in terms of Ornstein-Uhlenbeck (OU) processes. An OU process can be sampled directly from a Gaussian distribution once the initial condition of the process is chosen properly. This is similar to the previous framework (di Volo et al., 2019), however with one difference: we choose the initial condition of the OU process as a zero mean Gaussian, where its variance is scaled by the convergence rate of the OU process as given below. This allows to avoid explicit simulation of the OU process, reducing the computational load.

The noise is a stochastic process evolving in time, independently of the rest of the state variables of the model. For a white Gaussian noise, this evolution can be written as an OU process of the following type:

$$\begin{aligned} d\omega_\alpha(t) =& -\theta_\alpha \omega_\alpha(t)dt + \sigma dW_\alpha(t), \\ \omega_\alpha(0) =& \Omega, \end{aligned} \tag{11}$$

where $W_\alpha(t)$ is a standard Brownian motion, $\theta_\alpha$ is a positive constant associated to the convergence rate of the process to its long term mean, and $\Omega$ is the initial condition, which is a centered Gaussian noise with variance $\sigma_0^2 > 0$. This is a linear stochastic differential equation and its unique solution (Arnold, 1974) is

$$\omega_\alpha(t) = \mathrm{e}^{-\theta_\alpha t}\,\Omega + \sigma \int_0^t \mathrm{e}^{-\alpha(t-s)}\, dW_\alpha(s), \tag{12}$$

where the solution $\omega_\alpha(t)$ is a Gaussian process. It can be written for a small time step $0 < h \ll 1$ as

$$\omega_\alpha(t+h) = \mathrm{e}^{-\theta_\alpha h}\,\omega_\alpha(t) + \xi(t), \tag{13}$$

where $\xi(t)$ is a centered Gaussian white noise with variance $\sigma_\xi^2 = \frac{\sigma^2}{2\theta_\alpha}(1 - \mathrm{e}^{-2\theta_\alpha h})$, and which is sampled independently for each $t$ and subpopulation $\alpha$. We observe that $\omega_\alpha(t+h)$ is equivalent to a zero mean Gaussian with variance

$$\sigma_{\omega_\alpha}^2(t+h) = \mathrm{e}^{-2\theta_\alpha h}\,\sigma_{\omega_\alpha}(t)^2 + \sigma_\xi^2. \tag{14}$$

Once we choose $\sigma_{\omega_\alpha}^2(0) = \sigma_0^2 = \frac{\sigma^2}{2\theta_\alpha}$, we observe that $\sigma_{\omega_\alpha}^2(t+h)$ is time independent and it is equal to $\frac{\sigma^2}{2\theta_\alpha}$. This allows us to generate $\omega_\alpha(t)$ from the Gaussian distribution $\mathcal{N}(0, \frac{\sigma^2}{2\theta_\alpha})$ independently at each instant $t > 0$, avoiding explicit forward time simulations of the OU process given in (11). This is the idea behind introducing the noise terms $\omega_\alpha$ in the mean-field system (1) and the noise term $\zeta_i$ appearing in the regulatory mechanism (6) as Gaussian white noise sampled directly from a Gaussian distribution at each time and for each subpopulation, but not as an explicit OU process evolving in time.

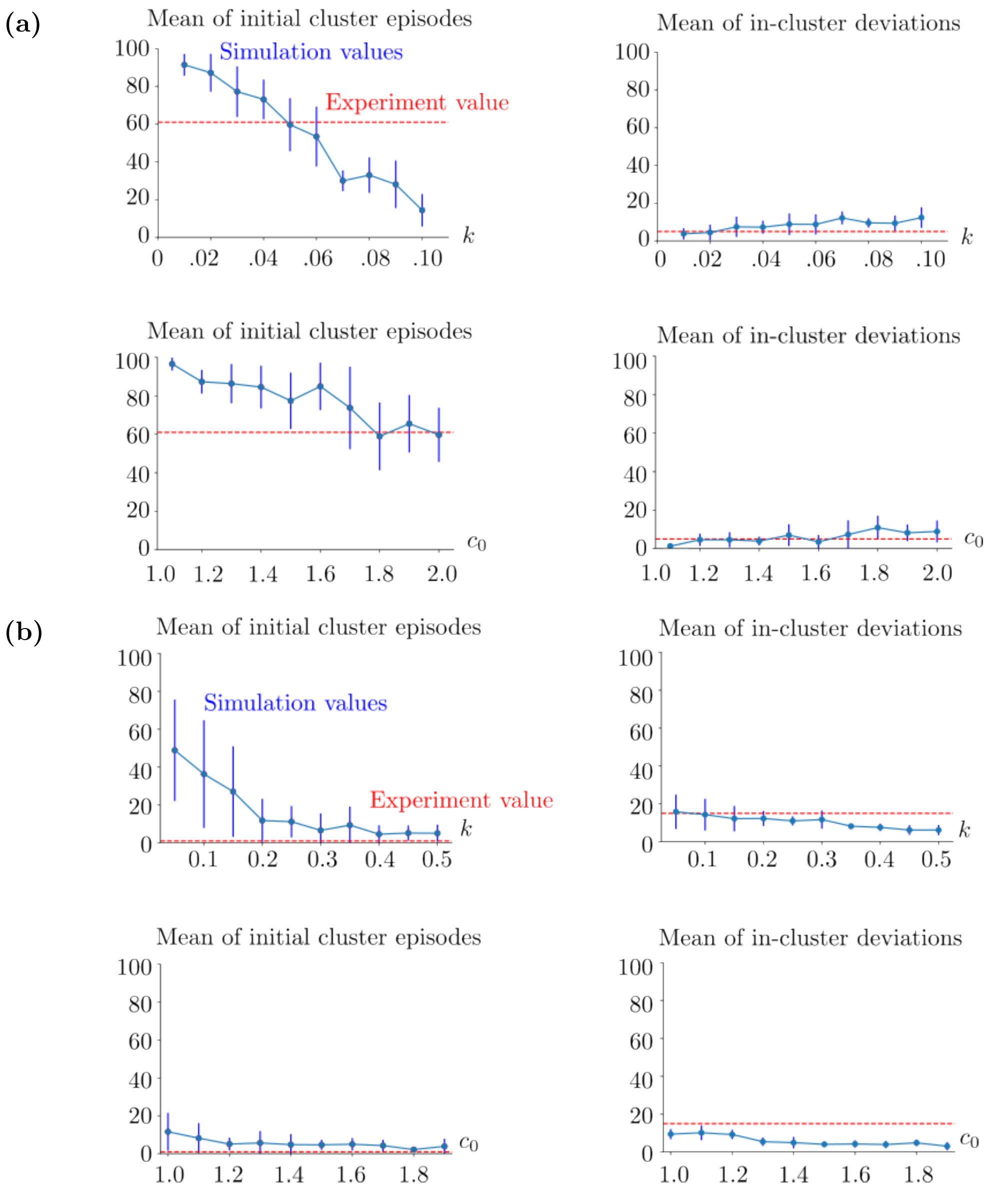


**Fig 5.** Simulation metrics with respect to the varied learning speed $k$ and flexibility parameter $c_0$. (a) Horizon 1 human case. On the top and bottom rows $k$ and $c_0$ are varied, respectively. In the top row, $c_0 = 2$. In the bottom row, $k = 0.05$. The vertical blue lines show the standard deviations of the simulation results. The red horizontal lines indicate the initial cluster episode number (left column) and the number of in-cluster deviations (right column) obtained from the human experiment. (b) Horizon 0 human case. In the top row, $c_0 = 1.1$ and $k$ is varied. In the bottom row, $k = 0.3$ and $c_0$ is varied. The blue vertical lines show the standard deviations of the corresponding statistical sample. The red horizontal lines indicate the experimental values as in (a). In both (a) and (b), the means and standard deviations are obtained from 10 realizations of the same simulation setup for each $(k, c_0)$ pair. The parameters $k$ and $c_0$ can be seen as rescaling constants, therefore they are unitless. See (23)-(25) for the rest of the parameters used in these simulations.

#### 6.2 Human reaction times

In the experiments, the reaction time in one trial is measured as the time duration between the instant when the stimuli are shown simultaneously on the monitor, and the moment when the participant starts to move the pointer. In the simulations, we measure the reaction time as the time duration between the instant when the two stimuli are provided to the model and the time instant at which the decision is made, i.e., when the difference between the excitatory subpopulation firing rates $v_{e_A}$ and $v_{e_B}$ exceeds the decision threshold, which is fixed to 5 Hz.

(a) (b)

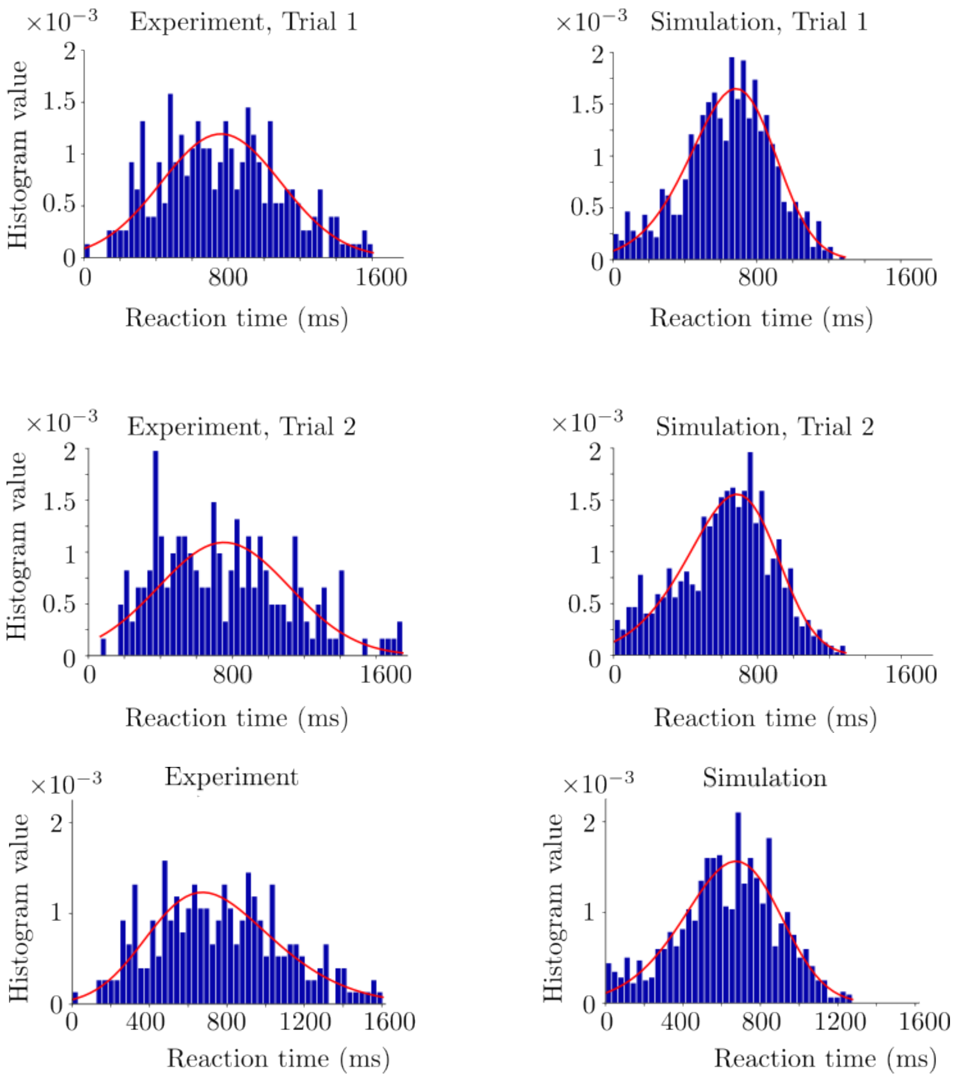


**Fig 6.** Reaction time histograms obtained from the human experiment and corresponding simulations. (a) Horizon 1. The red curves denote the fitted distributions via Python. Trial 1 and 2 histograms are in the top and bottom rows, respectively. The simulations are performed with $k = 0.05$, $c_0 = 2$ and decision threshold = 5 Hz. (b) Horizon 0. The simulations were performed with $k = 0.3$, $c_0 = 1.1$ and decision threshold = 5 Hz. See (23)-(25) for the rest of the parameters used in the simulations.

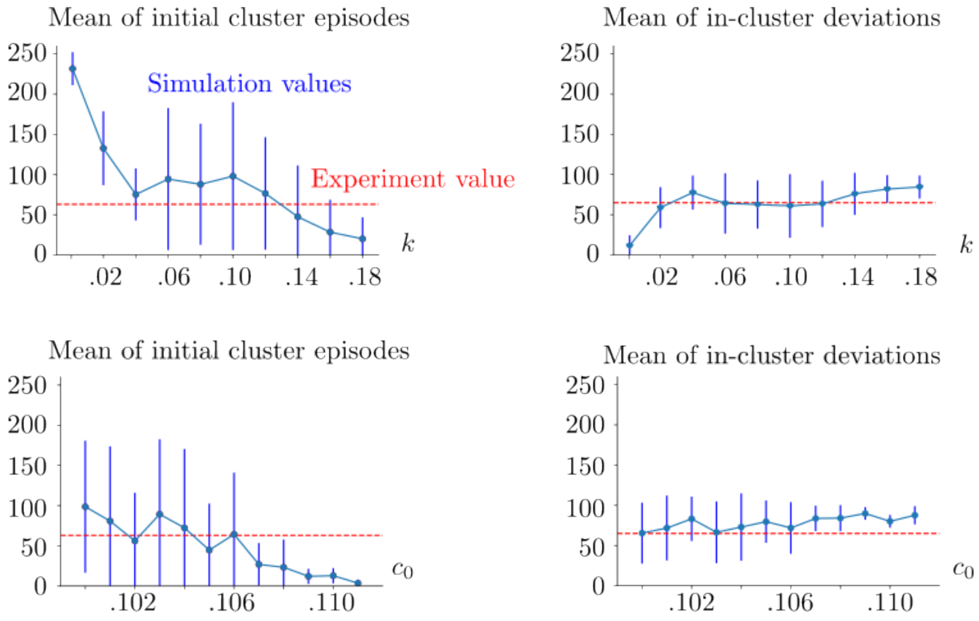


**Fig 7.** Simulation metrics of the macaque case. In the top row, $c_0 = 0.106$, and $k$ is varied. In the bottom row, $k = 0.12$, and $c_0$ is varied. The vertical blue lines show the standard deviation of the corresponding statistical sample. The red dashed lines show the value obtained from the macaque experiment. The mean and standard deviations were obtained from 10 realizations of the same simulation setup for each $(k, c_0)$ pair. The decision threshold is 5 Hz. See (23)-(25) for the rest of the parameters used in the simulations.

#### 6.4 Macaque reaction times

In Figure 8, we show the simulation and experiment histograms of the reaction times corresponding to the macaque experiment, together with the fitted distributions. Here $k$ and $c_0$ are fixed to the values providing the best performance overlap. We fit the distributions to both simulation and experiment histograms by using the *Fitter* function from the Python *Fitter* package as in the case of the human experiment. In Trial 1, genlogistic distribution and powernorm distribution are fitted to the experiment and simulation histograms, respectively. Corresponding squared sum error values are 1e-5 and 9e-6 for the experiment and simulation histograms, respectively. Pearson correlation coefficient of these fitted distributions is 0.97. In Trial 2 histograms, burr distribution and kappa3 distribution are fitted to the experiment and simulation histograms, respectively. Corresponding squared sum error values are 9.9e-5 and 5.73e-4 for the experiment and simulation histograms, respectively. Pearson correlation coefficient of these fitted distributions is 0.31. Trial 2 has a weaker overlap between the experiment and simulation histograms compared to Trial 1 as in the case of the human task.

The same factors regarding the reaction times and the initial conditions as in the human case are valid also in the macaque case. We predict that the difference between the experimental and simulation results regarding the reaction times arises from these factors; see Section 6.2.

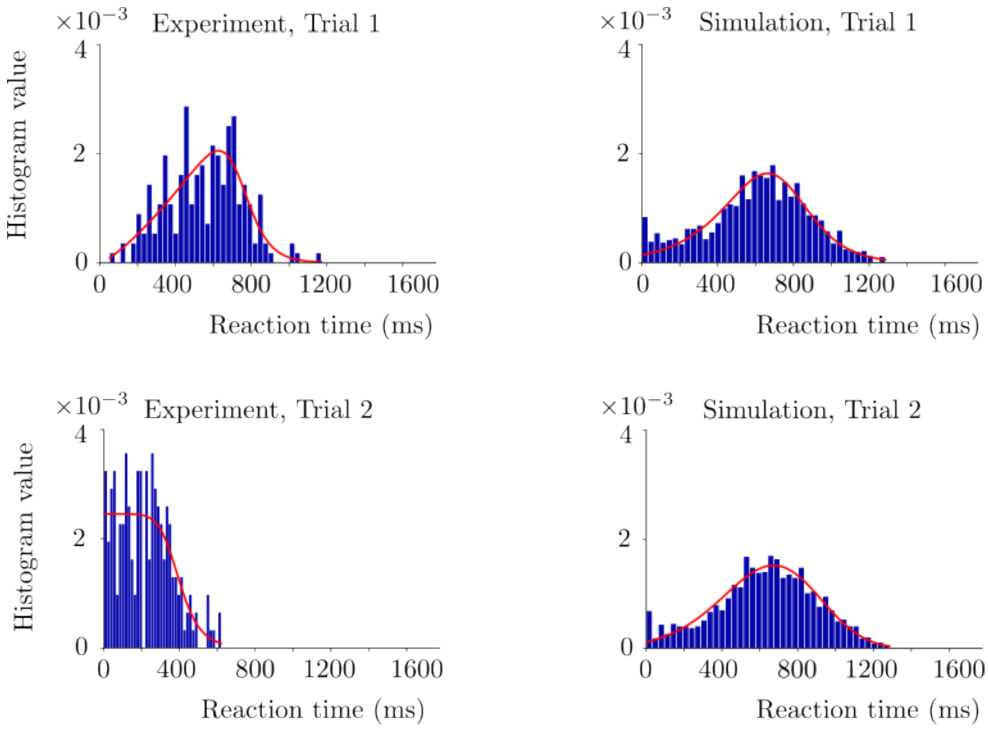


**Fig 8.** Reaction time histograms of the macaque case. The red curves denote the fitted distributions via Python. Trial 1 and 2 histograms are in the top and bottom rows, respectively. The simulations are performed with $k = 0.12$, $c_0 = 0.106$ and decision threshold $= 5$. See (23)-(25) for the rest of the parameters used in the simulations.

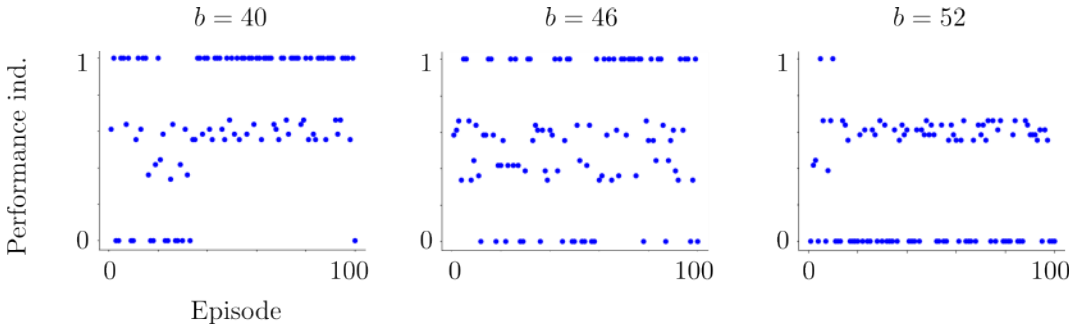


**Fig 9.** Performance index plots of the simulations of the model as the model goes through the transition from the awake state towards the sleeping state. The spike-frequency adaptation of the RS cells becomes stronger as we increase the parameter $b$ found in the adaptation term of (1). This pushes the model towards the sleeping state. We observe that the model loses gradually the capacity to obtain maximum performance indexes, i.e., it loses gradually the ability to learn the preset strategy.

Finally, a further extension of the model can be towards the large-scale brain dynamics of decision-making. For this, our AdEx mean-field framework can be integrated in The Virtual Brain (Sanz Leon et al., 2013; Goldman et al., 2020) to model the decision-making brain dynamics of the PFC, which has been traditionally considered as the key area involved in decision-making (Bechara et al., 2000; Hampton et al., 2006; Rushworth et al., 2011; Shadlen et al., 2013; Domenech et al., 2015; Funahashi et al., 2017). In this way, the perceptual inputs can be provided directly from the visual (or other perceptual) areas to the PFC. Consequently, the outputs of our model can be provided as the outputs of the PFC to the motor areas. Some preliminary results in this direction can be found in (Turan et al., 2023).

### Appendix

#### 1 Neural transfer functions

Neural transfer functions of AdEx mean-field equations were provided in (Zerlaut et al., 2018) and generalized to also spike-frequency adaptation in (di Volo et al., 2019). These functions were obtained by a semi-analytical fitting to the averaged output of the AdEx neuronal network. The output of the network was induced by random spike bombardments generated from a Poisson distribution. We use the same transfer functions provided in (di Volo et al., 2019). We summarize here how their explicit formulas were obtained. For details, we refer to (Zerlaut et al., 2018; di Volo et al., 2019).

##### AdEx neuronal network

The AdEx mean-field of a pair of excitatory-inhibitory subpopulations with spike-frequency adaptation approximates the average dynamics of the AdEx neuronal network. The network is composed of a certain number of neurons. Each neuron in the network is described by the AdEx integrate-and-fire model (Brette et al.,

2005):

$$C_m \frac{dV_\ell}{dt} = g_L\,(E_L - V_\ell) + \Delta\,\mathrm{e}^{\frac{V_\ell - V_{\rm thr}}{\Delta}} - w_\ell + I_{\rm syn},$$
$$\frac{dw_\ell}{dt} = -\frac{w_\ell}{\tau_w} + b \sum_{t_s \in \{t_{\rm spike}(\ell)\}} \delta_{t-t_s} + a\,(V_\ell - E_L), \tag{15}$$

where $C_m = 200$ pF denotes the membrane capacity and $V_\ell$ is the membrane voltage of a neuron $\ell$ in the network. The voltage $V_\ell$ is reset to the resting voltage $-65$ mV at $t_s \in \{t_{\rm spike}(\ell)\}$, which denote all the instants when the neuron generates a spike, i.e, when $V_\ell > V_{\rm thr} = -50$ mV. Here $g_L = 10$ nS and $E_L = -65$ mV are the conductance and the leakage reversal of the leak term. Here $\Delta$ denotes the weight of the exponential term. It is 2 mV for RS cells and 5 mV for FS cells. Note that FS cells have no adaptation ($w_\ell = 0$). Therefore, $a = b = 0$ for all inhibitory neurons. For RS cells, the adaptation is introduced with $a = 4$ and $b = 40$ in our simulations. Synaptic input to the neuron $\ell$ is defined as follows:

$$I_{\rm syn} = (E_L^e - V_\ell)G_e + (E_L^i - V_\ell)G_i, \tag{16}$$

with $E_L^e = 0$ mV and $E_L^i = -80$ mV denoting the reversal potential of the excitatory and inhibitory presynaptic cells, respectively. The conductances are defined as

$$G_e(t) = Q_e \sum_{t_s^e \in \{t^e_{\rm spike}(\ell)\}} \mathcal{H}(t - t_s^e)\,\mathrm{e}^{-\frac{t-t_s^e}{\tau_e}}, \qquad G_i(t) = Q_i \sum_{t_s^i \in \{t^i_{\rm spike}(\ell)\}} \mathcal{H}(t - t_s^i)\,\mathrm{e}^{-\frac{t-t_s^i}{\tau_i}}, \tag{17}$$

where $\mathcal{H}$ denotes the Heaviside function and $\tau_e = \tau_i = 5$ ms are the decay rates of the excitatory and inhibitory presynaptic neurons. Here $t_s^e$ and $t_s^i$ are the instants when the excitatory and inhibitory presynaptic neurons trigger spikes, respectively. Finally, $Q_e = 1.5$ nS and $Q_i = 5$ nS are the excitatory and inhibitory quantal conductances, respectively.

#### Transfer functions

The classical AdEx mean-field equations were derived in (di Volo et al., 2019) from the AdEx network of neurons described by (15). The derivation of the mean-field equations was based on a master equation formalism which was provided in (El Boustani et al., 2009). The mean-field equations were for a single pair of excitatory-inhibitory subpopulations, with a spike-frequency adaptation. This is equivalent to decoupling the L2/3 populations of the basic module (1) of our model and considering only one of these decoupled populations. In that case, each L2/3 population reduces to a system of 6 state variables as in the classical mean-field system. Therefore, for the neural subpopulations in our model, we use the same transfer functions given for the classical AdEx mean-field system in (di Volo et al., 2019). The difference is that, the inputs to the transfer functions have not only the local components, but also the components related to the interactions between the L2/3 populations in our extended AdEx mean-field framework.

In the following, we will use the notation $e, i$ to denote the excitatory and inhibitory subpopulations in either one of the decoupled L2/3 populations. The equations of the transfer functions are identical, independently of which L2/3 population we consider. These neural transfer functions were obtained via the semi-analytical derivation explained in (Zerlaut et al., 2018; di Volo et al., 2019).

The idea was based on the spike bombardment of a single neuron described by (15). The spikes were generated with various excitatory and inhibitory firing rates $v_e$ and $v_i$ which follow the Poisson statistics. This allowed to obtain the means $\mu_{G_e}, \mu_{G_i}$ and the standard deviations $\sigma_{G_e}, \sigma_{G_i}$ of the excitatory and inhibitory conductances in (17). These functions were found via Campbell-Hardy theorem (Papoulis et al., 2002) as follows:

$$\begin{aligned} \mu_{G_e}(v_e, v_i) &= v_e\, K_e\, \tau_e\, Q_e, \qquad \sigma_{G_e}(v_e, v_i) = Q_e \sqrt{\frac{v_e\, K_e\, \tau_e}{2}}, \\ \mu_{G_i}(v_e, v_i) &= v_i\, K_i\, \tau_i\, Q_i, \qquad \sigma_{G_i}(v_e, v_i) = Q_i \sqrt{\frac{v_i\, K_i\, \tau_i}{2}}. \end{aligned} \tag{18}$$

These functions control the input conductance $\mu_G$ and its effective membrane time constant $\tau_m$, which have the following expressions:

$$\mu_G(v_e, v_i) = \mu_{G_e} + \mu_{G_i} + g_L, \qquad \tau_m(v_e, v_i) = \frac{C_m}{\mu_{G_e} + \mu_{G_i} + g_L}. \tag{19}$$

Consequently, mean membrane voltage of the neuron was obtained as:

$$\mu_V(v_e, v_i) = \frac{\mu_{G_e} E_L^e + \mu_{G_i} E_L^i + g_L E_L - w}{\mu_G}, \tag{20}$$

where $w$ denotes the adaptation of the neuron. Finally, the standard deviation $\sigma_V$ and the autocorrelation time $\tau_V$ of the membrane voltage fluctations were found from the power density of the fluctutations:

$$\begin{aligned} \sigma_V(v_e, v_i) &= \sqrt{\sum_{\alpha \in \{e,i\}} K_\alpha v_\alpha \frac{(U_\alpha\, \tau_\alpha)^2}{2(\tau_m + \tau_\alpha)}}, \\ \tau_V(v_e, v_i) &= \frac{\sum_{\alpha \in \{e,i\}} K_\alpha v_\alpha (U_\alpha\, \tau_\alpha)^2}{\sum_{\alpha \in \{e,i\}} K_\alpha v_\alpha (U_\alpha\, \tau_\alpha)^2 / (\tau_m + \tau_\alpha)}, \end{aligned} \tag{21}$$

where $U_\alpha = \frac{Q_\alpha}{\mu_G}(E_L^\alpha - \mu_V)$.

In (21) and (22), we find the analytical expressions used in the transfer functions given by the formula (5). These analytical experessions are complemented by the voltage-effective threshold $v_{\text{eff}}$, which was found by fitting the coefficients of the second order polynomial:

$$v_{\text{eff}}(\mu_V, \sigma_V, \rho_V) = P_0 + \sum_{x \in \{\mu_V, \sigma_V, \rho_V\}} P_x \frac{x - x^0}{\bar{x}^0} + \sum_{x,y \in \{\mu_V, \sigma_V, \rho_V\}} P_{xy} \Big(\frac{x - x^0}{\bar{x}^0}\Big)\Big(\frac{y - y^0}{\bar{y}^0}\Big). \tag{22}$$

The fitting was made according to the simulations of single neuron activity described by (15). Here $\rho_V = \tau_V \frac{g_L}{C_m}$ and $\mu_V^0 = -60$ mV, $\sigma_V^0 = 0.004$ mV, $\rho_V = 0.5$, $\bar{\mu}_V^0 = 0.001$ mV, $\bar{\sigma}_V^0 = 0.006$ mV, $\bar{\rho}_V^0 = 1$. We use the same fitted coefficients as given in (di Volo et al., 2019). The fitting of the polynomial (22) and the functions in (20), (21) provide the transfer functions given by (5).

#### 2 Simulation parameters

The parameters which are used in Figures 5-8 and Figures 11, 12 are as follows:

$$\begin{aligned} T &= 5\,\mathrm{ms}^{-1}, \quad \sigma = 0.01, \quad \tau_w = 5000\,\mathrm{ms}^{-1} \text{ (for RS)}, \quad 1^{-9}\,\mathrm{ms}^{-1} \text{ (for FS)}, \\ a &= 4 \text{ (for RS)}, \quad 0\text{(for FS)}, \quad b = 40 \text{ (RS)},\ 0 \text{ (FS)}, \quad E_L = -65\,\mathrm{mV}, \end{aligned} \tag{23}$$

and it is given for (4) as

$$s_{\mathrm{AI}} = 5\,\mathrm{Hz}, \quad w_c = 1.6. \tag{24}$$

Finally, the parameters appearing in (6) and (7) are

$$\tau_\psi = T = 5, \qquad \sigma = 0.01. \tag{25}$$

Each trial lasts 15 seconds in both Horizon 0 and 1 simulations. We set the decision threshold to 5 Hz in all the simulations.

#### 3 Effects of varying the bias and the difference between the stimuli

The results in Figure 10 focus on the effects of varying the initial condition $\psi(0)$ of the reuglatory module, therefore the effects of varying the bias. The decision threshold to measure the reaction time $\bar{t}$ is $|v_{e_A}(\bar{t}) - v_{e_B}(\bar{t})| = 7.5$ Hz. We use Euler-Maruyama scheme with time step $\Delta t = 0.5$ and $t_f = 4$ for the presented results in Figure 10. Stimuli are applied at $t_0 = 2$. The parameters in this framework are as follows:

$$\begin{aligned} T &= 0.003, \quad \sigma = 0.01, \quad \tau_w = 500 \text{ (for RS)},\ 10^{-9} \text{ (for FS)}, \\ a &= 1 \text{ (RS)},\ 0\text{(FS)}, \quad b = 100 \text{ (RS)},\ 0 \text{ (FS)}, \quad E_L = -65. \end{aligned} \tag{26}$$

Moreover, in (4) we fix

$$s_{\mathrm{AI}} = 5\,\mathrm{Hz}, \quad w_c = 2. \tag{27}$$

Finally, the parameters appearing in (6) are

$$\tau_\psi = 10\,T = 0.03, \qquad \sigma = 0.01, \qquad c_0 = 1000. \tag{28}$$

In Figure 10a, we provide the results of the cases with $\psi(0)$ varied from 0 to 1, where the same stimuli were applied in each case. There is no clear difference in terms of reaction time between the cases with different initial conditions $\psi(0)$ as long as the stimuli remain the same and $\psi$ converges to the same value. It is due to the fact that, the convergence of the regulatory mechanism is rapid, therefore the competition is promoted towards the same L2/3 population. As we do not change the stimuli, the evolution of $v_{e_A}, v_{e_B}$ becomes different realizations of almost the same random processes. Consequently, the reaction times of those realizations fluctuate around the same value.

In Figure 10b, we show the effects of varying the difference between the stimuli on the model reaction time. The reaction time is measured in the same way as in Figure 10a. We set the amplitude of Stimuli $A$ to 6 and vary the amplitude of Stimuli $B$ between 0 and 6. We keep the initial value $\psi(0)$ of the regulatory function constant ($= 0.75$) such that the system privileges always Stimulus $A$. We observe that the reaction time increases as the difference between the stimuli decreases. It is expected since it becomes more difficult to make a distinction between the stimuli as the difference between the filled quantities of the bars decreases.

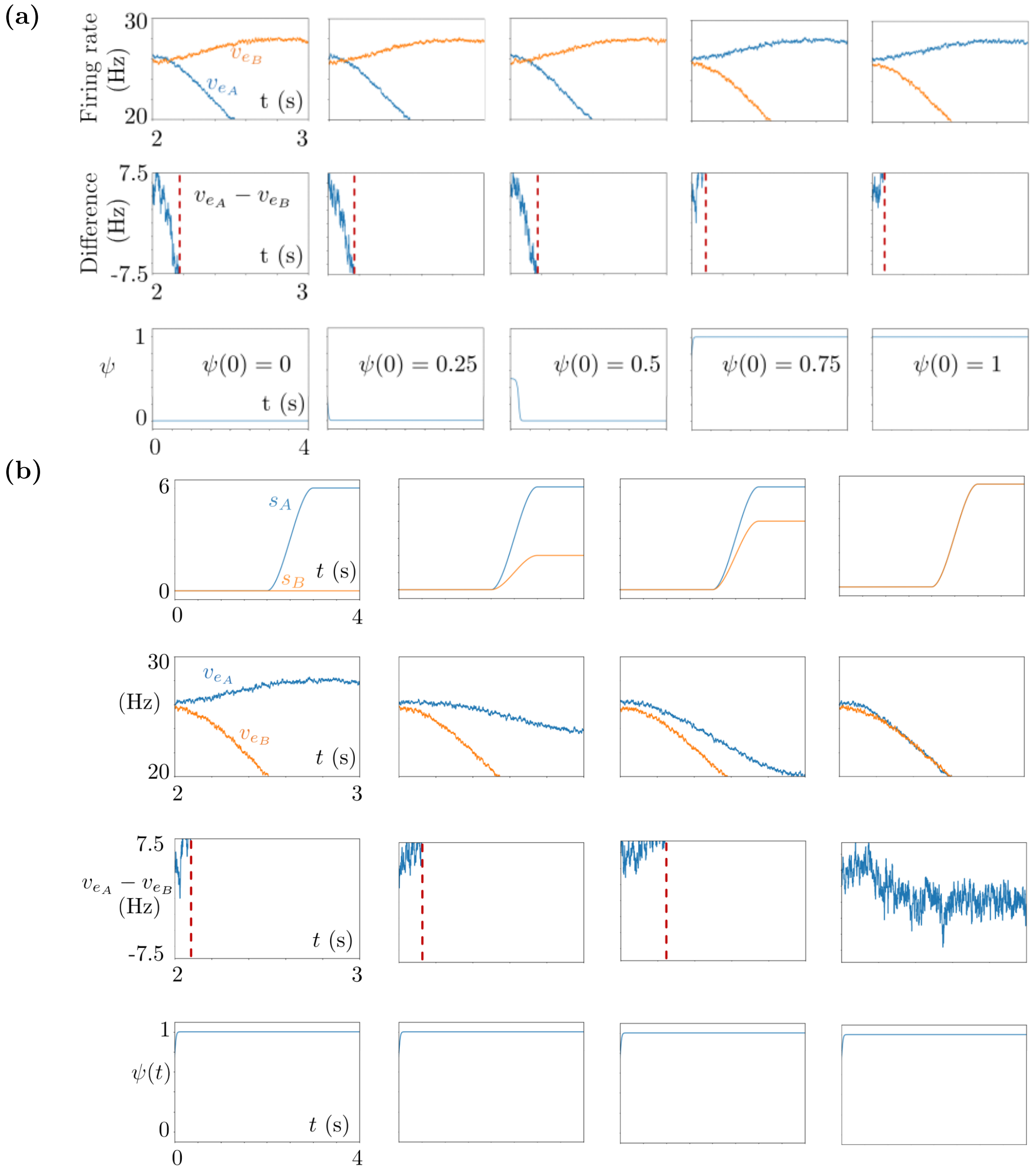


**Fig 10.** Simulation results of isolated trials, with varying regulatory module initial conditions $\psi(0)$ in (a) and varying stimuli amplitudes in (b). The axes are specified only in the first columns in both panels, and they are identical in the other columns. **(a)** Top: Time courses of $v_{e_A}$ and $v_{e_B}$. Middle: Difference of the firing rates where the red vertical line denotes the decision instant. Bottom: Time evolution of the regulatory function $\psi(t)$. The initial values $\psi(0)$ are 0, 0.25, 0.5, 0.75, 1 from left to right. **(c)** Top row: Time courses of the external stimuli $s_A$ and $s_B$. Second row: Time courses of $v_{e_A}$ and $v_{e_B}$. Third row: Time course of the difference between the membrane potentials where the red vertical line denotes the decision instant. Bottom row: Time course of the regulatory function $\psi$. The initial value $\psi(0)$ is 0.75 in all plots.

#### 4 Characterization of the model behavioral performance

Learning speed determines how quickly the model learns the preset strategy. The higher it is, the faster the model identifies the strategy. Flexibility parameter determines how much the model deviates from the preset strategy after it learns the strategy. It models the exploratory behavior and the deviations caused by perceptual difficulties. Finally, the gain rescales the learning speed. Here we provide the effects of these three parameters on the model behavior performance.

We show in Figure 11b the performance index plots (human case) in which the learning speed $k$ appearing in (8) varies from 0.1 to 0.3. We denote by $V_n^E$ the value of the chosen stimulus in the $n^{\text{th}}$ trial of the $E^{\text{th}}$ episode. We denote the maximum and minimum cumulative reward values of episode $E$ by $V_{\min}^E$ and $V_{\max}^E$, respectively. This is equivalent to maximum and minimum values that $\sum_{n=1}^{L} V_n^E$ can attain among all possible scenarios of the $E^{\text{th}}$ episode, with $L$ denoting the number of trials in the episode. The performance index of the model in the $E^{\text{th}}$ episode is computed as

$$\text{Performance index}\,(E) := \frac{\sum_{n=1}^{L} V_n^E - V_{\min}^E}{V_{\max}^E - V_{\min}^E}. \tag{29}$$

We observe in Figure 11b that as we increase $k$, the system captures the strategy earlier, and makes constantly right decisions thereafter, with very few deviations.

The gain $G$ is another factor affecting the learning speed. The learning speed decreases as the gain decreases since the gain $G$ is equal to the multiplying factor $(M_{n+1}^E - M_n^E)$ of the learning speed $k$ in (8). As shown in Figure 11c, the model learns the preset strategy faster as we increase the gain.

Finally, the flexibility parameter $c_0$ determines the number of deviations in the model performance results. The higher it is, the closer to the deterministic case the model is. This is due to the fact that the noise given in (6), and which produces the performance deviations, vanishes very quickly after that the regulatory variable $\psi$ starts to evolve at the beginning of each trial; see Figure 11d. Moreover, $c_0$ determines the beginning of the performance cluster together with the learning speed. As $c_0$ increases, the performance cluster is more likely to begin from smaller episode numbers.

#### 5 Characterization of the model reaction time

In Figure 12, we provide the histograms of the reaction time measurements obtained from the Horizon 1 simulations (human case). In these simulations, we varied the learning speed $k$ and the flexibility parameter $c_0$, separately in Trial 1 and Trial 2. We observe that there is no distinguishing change in the histograms of the simulation results, suggesting that these two parameters do not have noticeable effects on the reaction times. This is due to the fact that the reaction times are calculated by checking the threshold crossing of the difference between the firing rates $v_{e_A}$ and $v_{e_B}$. Here $k$ and $c_0$ do not enter in the equations that describe the firing rates, so they do not influence the reaction times. As the decision threshold increases, the competition between $v_{e_A}$ and $v_{e_B}$ lasts longer, thus the reaction times increase as shown in the bottom row of Figure 12.

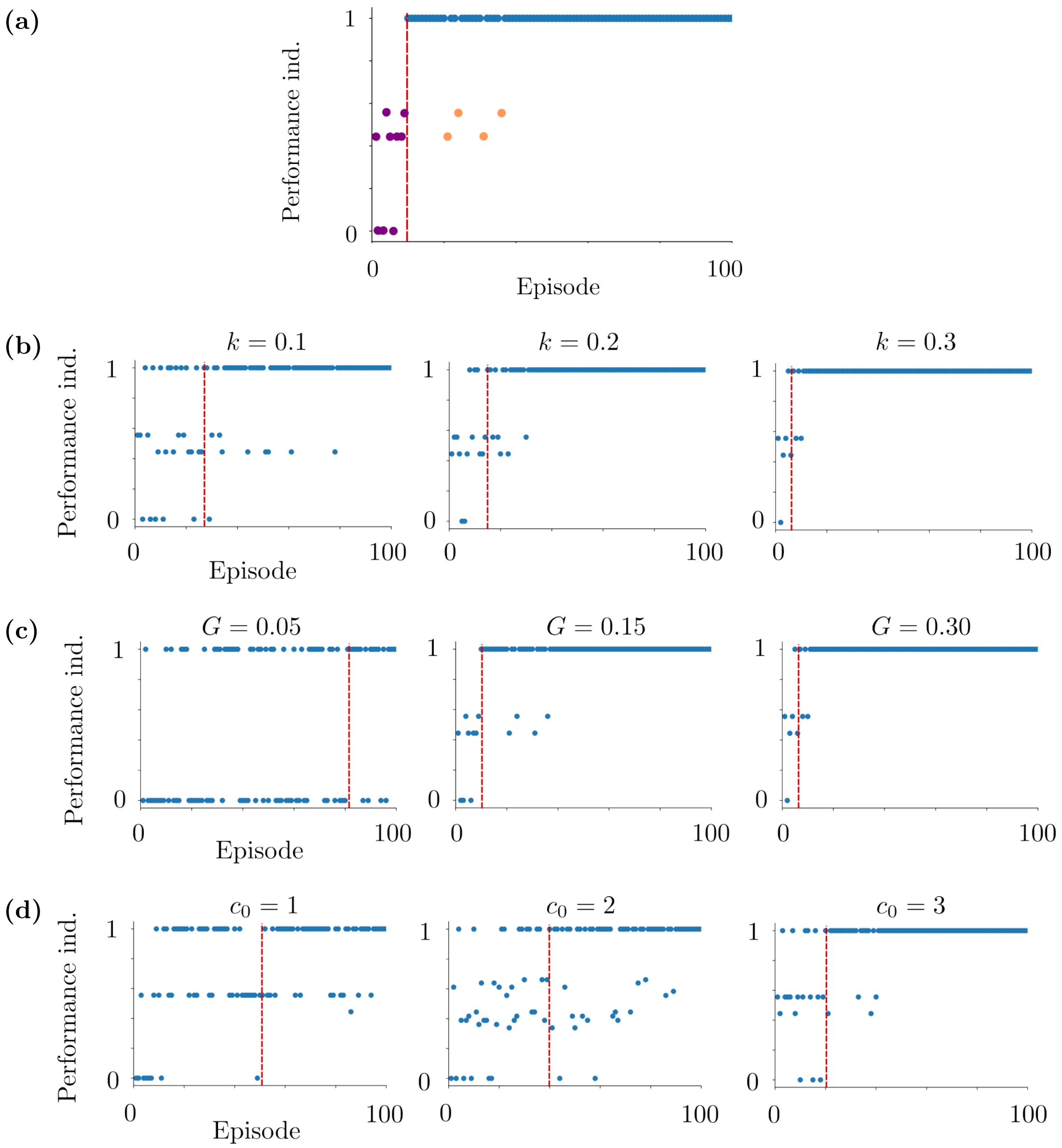


**Fig 11.** Objects used for quantification of behavior in (a), and performance indexes regarding the Horizon 1 human task simulations in (b), (c) and (d). **(a)** Purple samples correspond to a deviation block, and they are out-cluster deviations. Orange samples are in-cluster deviations, they do not form a deviation block. Both the purple and orange samples are performance deviations. Blue samples are maximum performance indexes and they form a performance cluster. The vertical red line marks the initial cluster episode, it corresponds to the beginning of the performance cluster. **(b)** Learning speed $k$ varies. The flexibility parameter $c_0$ is 2 and the gain $G$ is 0.3. The initial episodes of the performance clusters are episodes 24, 15 and 5 from left to right. **(c)** $G = 0.05$, $G = 0.15$ and $G = 0.3$ from left to right. Here $k = 0.3$ and $c_0 = 2$. The initial episodes of the performance clusters are episodes 81, 10 and 5 from left to right. **(d)** $c_0 = 1$, $c_0 = 2$ and $c_0 = 3$ from left to right. Here $k = 0.05$ and the gain $G = 0.3$. The initial episodes of the performance clusters are episodes 52, 40 and 20; the number of in-cluster deviations are 12, 5 and 4 from left to right. The difficulty $d$ is 0.2 in (b), (c) and (d).

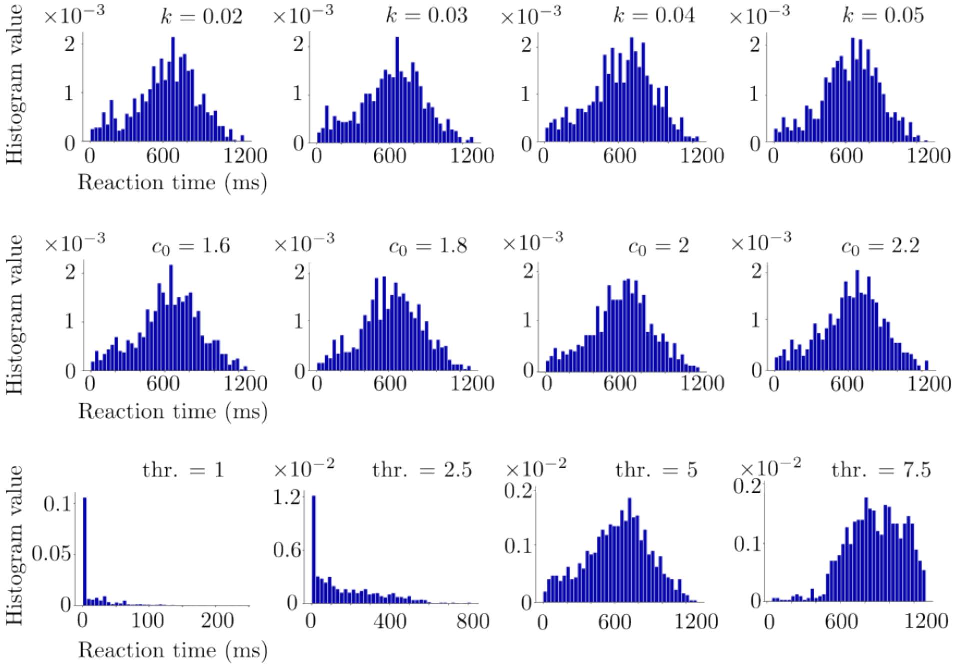


Trial 1

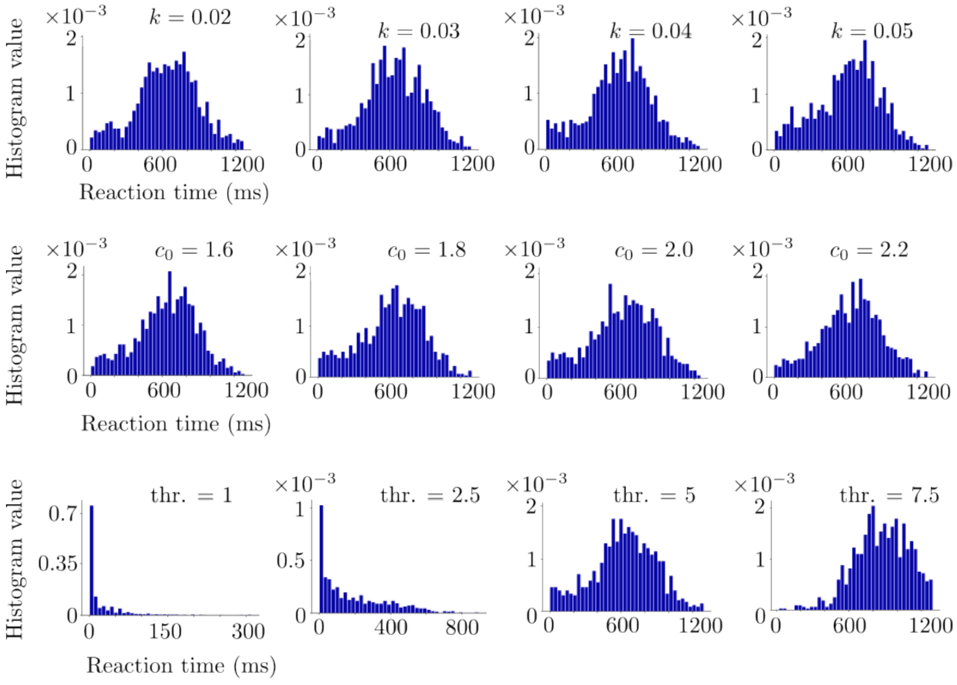


Trial 2

**Fig 12.** Simulation results regarding Horizon 1 reaction time histograms of Trials 1 and 2. Varying the learning speed $k$ and the flexibility parameter $c_0$ does not affect noticeably the distribution of the reaction times in terms of mean and standard deviation. Increasing the decision threshold increases the mean and the standard deviation of the distribution. The histograms are obtained from 10 realizations of the same simulation setup for each $k$, $c_0$ and decision threshold.

### Acknowledgments

We hereby acknowledge that this research was supported by the Human Brain Project (European Union grant H2020-945539).
