## Supplementary material for "A biologically plausible decision-making model based on interacting neural populations": Tex files: main.tex

% Template for PLoS
% Version 3.6 Aug 2022
%
% % % % % % % % % % % % % % % % % % % % % %
%
% -- IMPORTANT NOTE
%
% This template contains comments intended 
% to minimize problems and delays during our production 
% process. Please follow the template instructions
% whenever possible.
%
% % % % % % % % % % % % % % % % % % % % % % % 
%
% Once your paper is accepted for publication, 
% PLEASE REMOVE ALL TRACKED CHANGES in this file 
% and leave only the final text of your manuscript. 
% PLOS recommends the use of latexdiff to track changes during review, as this will help to maintain a clean tex file.
% Visit https://www.ctan.org/pkg/latexdiff?lang=en for info or contact us at.
%
%
% There are no restrictions on package use within the LaTeX files except that no packages listed in the template may be deleted.
%
% Please do not include colors or graphics in the text.
%
% The manuscript LaTeX source should be contained within a single file (do not use \input, \externaldocument, or similar commands).
%
% % % % % % % % % % % % % % % % % % % % % % %
%
% -- FIGURES AND TABLES
%
% Please include tables/figure captions directly after the paragraph where they are first cited in the text.
%
% DO NOT INCLUDE GRAPHICS IN YOUR MANUSCRIPT
% - Figures should be uploaded separately from your manuscript file. 
% - Figures generated using LaTeX should be extracted and removed from the PDF before submission. 
% - Figures containing multiple panels/subfigures must be combined into one image file before submission.
% For figure citations, please use "Fig" instead of "Figure".
% See http://journals.plos.org/plosone/s/figures for PLOS figure guidelines.
%
% Tables should be cell-based and may not contain:
% - spacing/line breaks within cells to alter layout or alignment
% - do not nest tabular environments (no tabular environments within tabular environments)
% - no graphics or colored text (cell background color/shading OK)
% See http://journals.plos.org/plosone/s/tables for table guidelines.
%
% For tables that exceed the width of the text column, use the adjustwidth environment as illustrated in the example table in text below.
%
% % % % % % % % % % % % % % % % % % % % % % % %
%
% -- EQUATIONS, MATH SYMBOLS, SUBSCRIPTS, AND SUPERSCRIPTS
%
% IMPORTANT
% Below are a few tips to help format your equations and other special characters according to our specifications. For more tips to help reduce the possibility of formatting errors during conversion, please see our LaTeX guidelines at http://journals.plos.org/plosone/s/latex
%
% For inline equations, please be sure to include all portions of an equation in the math environment.  For example, x$^2$ is incorrect; this should be formatted as $x^2$ (or $\mathrm{x}^2$ if the romanized font is desired).
%
% Do not include text that is not math in the math environment. For example, CO2 should be written as CO\textsubscript{2} instead of CO$_2$.
%
% Please add line breaks to long display equations when possible in order to fit size of the column. 
%
% For inline equations, please do not include punctuation (commas, etc) within the math environment unless this is part of the equation.
%
% When adding superscript or subscripts outside of brackets/braces, please group using {}.  For example, change "[U(D,E,\gamma)]^2" to "{[U(D,E,\gamma)]}^2". 
%
% Do not use \cal for caligraphic font.  Instead, use \mathcal{}
%
% % % % % % % % % % % % % % % % % % % % % % % % 
%
% Please contact with any questions.
%
% % % % % % % % % % % % % % % % % % % % % % % %

\documentclass[12pt,a4paper]{article}
%\usepackage[top=0.85in,left=2.75in,footskip=0.75in]{geometry}
%\usepackage{geometry}
\usepackage[top=0.7in, bottom=0.95in]{geometry}
%\usepackage[a4paper, left=0.5in, right=0.5in, top=0.7in, bottom=1in, footskip=.25in]{geometry}

% amsmath and amssymb packages, useful for mathematical formulas and symbols
\usepackage{amsmath,amssymb}

% Use adjustwidth environment to exceed column width (see example table in text)
\usepackage{changepage}

% textcomp package and marvosym package for additional characters
\usepackage{textcomp,marvosym}

%% cite package, to clean up citations in the main text. Do not remove.
%\usepackage{cite}

% Use nameref to cite supporting information files (see Supporting Information section for more info)
\usepackage{nameref,hyperref}

% line numbers
%\usepackage[right]{lineno}
%\nolinenumbers

%% ligatures disabled
%\usepackage[nopatch=eqnum]{microtype}
%\DisableLigatures[f]{encoding = *, family = * }
% ligatures disabled
\usepackage{microtype}
\DisableLigatures[f]{encoding = *, family = * }

% color can be used to apply background shading to table cells only
\usepackage[table]{xcolor}

% array package and thick rules for tables
\usepackage{array}

% My packages
\usepackage{floatrow}
\floatsetup[figure]{subcapbesideposition=top}
\usepackage[caption=false]{subfig}
\usepackage{placeins}
\usepackage{mathtools}
\usepackage[authoryear]{natbib}

\usepackage[colorinlistoftodos]{todonotes}
\usepackage{color,soul}
%\usepackage[font=small,labelfont=bf]{caption}

% create "+" rule type for thick vertical lines
\newcolumntype{+}{!{\vrule width 2pt}}

% create \thickcline for thick horizontal lines of variable length
\newlength\savedwidth
\newcommand\thickcline[1]{%
  \noalign{\global\savedwidth\arrayrulewidth\global\arrayrulewidth 2pt}%
  \cline{#1}%
  \noalign{\vskip\arrayrulewidth}%
  \noalign{\global\arrayrulewidth\savedwidth}%
}

% \thickhline command for thick horizontal lines that span the table
\newcommand\thickhline{\noalign{\global\savedwidth\arrayrulewidth\global\arrayrulewidth 2pt}%
\hline
\noalign{\global\arrayrulewidth\savedwidth}}


% Remove comment for double spacing
%\usepackage{setspace} 
%\doublespacing

%% Text layout
%\raggedright
%\setlength{\parindent}{0.5cm}
%\textwidth 5.25in 
%\textheight 8.75in

% Bold the 'Figure #' in the caption and separate it from the title/caption with a period
% Captions will be left justified
\usepackage[aboveskip=1pt,labelfont=bf,labelsep=period,
justification=justified, singlelinecheck=off]{caption}
\renewcommand{\figurename}{Fig}

% Use the PLoS provided BiBTeX style
%\bibliographystyle{plos2015}

% Remove brackets from numbering in List of References
\makeatletter
\renewcommand{\@biblabel}[1]{\quad#1.}
\makeatother

%% margins
%\textheight 23.4cm
%\textwidth 14.65cm
%\oddsidemargin 0.375in
%\evensidemargin 0.375in
%\topmargin  -0.55in

%% Header and Footer with logo
%\usepackage{lastpage,fancyhdr,graphicx}
%\usepackage{epstopdf}
%%\pagestyle{myheadings}
%\pagestyle{fancy}
%\fancyhf{}
%%\setlength{\headheight}{27.023pt}
%%\lhead{\includegraphics[width=2.0in]{PLOS-submission.eps}}
%\rfoot{\thepage/\pageref{LastPage}}
%\renewcommand{\headrulewidth}{0pt}
%\renewcommand{\footrule}{\hrule height 2pt \vspace{2mm}}
%\fancyheadoffset[L]{2.25in}
%\fancyfootoffset[L]{2.25in}
%\lfoot{\today}

%% Include all macros below

\newcommand{\lorem}{{\bf LOREM}}
\newcommand{\ipsum}{{\bf IPSUM}}

%% END MACROS SECTION


\begin{document}
\vspace*{0.2in}

% Title must be 250 characters or less.
\begin{flushleft}
{\Large
\textbf\newline{A biologically plausible decision-making model based on interacting neural populations} % Please use "sentence case" for title and headings (capitalize only the first word in a title (or heading), the first word in a subtitle (or subheading), and any proper nouns).
}
\newline
% Insert author names, affiliations and corresponding author email (do not include titles, positions, or degrees).
\\
Emre Baspinar\textsuperscript{1,*},
Gloria Cecchini\textsuperscript{2,3},
Michael DePass\textsuperscript{3},
Marta Andujar\textsuperscript{4},
Pierpaolo Pani\textsuperscript{4},
Stefano Ferraina\textsuperscript{4},
Rub\'{e}n Moreno-Bote\textsuperscript{3,5},
Ignasi Cos\textsuperscript{2,5},
Alain Destexhe\textsuperscript{1}
\\
\bigskip
\textbf{1} Paris-Saclay University, CNRS, NeuroPSI, Saclay, France
\\
\textbf{2} Facultat de Matem\`{a}tiques i Inform\`{a}tica, Universitat de Barcelona, Barcelona, Catalonia, Spain
\\
\textbf{3} Center for Brain and Cognition, DTIC, Universitat Pompeu Fabra, Barcelona, Catalonia, Spain
\\
\textbf{4} Department of Physiology and Pharmacology, Sapienza University of Rome, Rome, Italy
\\
\textbf{5} Serra-Hunter Fellow Programme, Barcelona, Catalonia, Spain
\\
\bigskip

%% Insert additional author notes using the symbols described below. Insert symbol callouts after author names as necessary.
%% 
%% Remove or comment out the author notes below if they aren't used.
%%
%% Primary Equal Contribution Note
%\Yinyang These authors contributed equally to this work.
%
%% Additional Equal Contribution Note
%% Also use this double-dagger symbol for special authorship notes, such as senior authorship.
%\ddag These authors also contributed equally to this work.
%
%% Current address notes
%\textcurrency Current Address: Dept/Program/Center, Institution Name, City, State, Country % change symbol to "\textcurrency a" if more than one current address note
%% \textcurrency b Insert second current address 
%% \textcurrency c Insert third current address
%
%% Deceased author note
%\dag Deceased
%
%% Group/Consortium Author Note
%\textpilcrow Membership list can be found in the Acknowledgments section.

% Use the asterisk to denote corresponding authorship and provide email address in note below.

\end{flushleft}


\section*{Abstract}
We present a novel decision-making model with two populations. Each population is composed of Regularly Spiking (excitatory) and Fast Spiking (inhibitory) cells in cortical layer 2/3. Each population votes for one of the two visual alternatives shown on a monitor in human and macaque experiments. The model is biophysically plausible since it is based on long-range cortico-cortical connections between the layer 2/3 populations. These connections are excitatory. They contact both Regularly Spiking and Fast Spiking cells. This long-range excitation is conflicted by an inhibition based on local connections within the populations. This configuration introduces a competition between the layer 2/3 populations, sufficient for making a decision to choose between two alternatives shown on the monitor. We integrate the model with a reward-driven learning mechanism. This allows the model to learn the optimal strategy maximizing the cumulative reward in the long term. We test the model on two decision-making tasks applied on human and macaque. This model elaborates certain biophysical details which were not considered by simpler phenomenological models proposed for similar decision-making tasks. Finally, the model can be embedded in a brain simulator such as The Virtual Brain to study decision-making in terms of large-scale brain dynamics.

% Please keep the Author Summary between 150 and 200 words
% Use first person. PLOS ONE authors please skip this step. 
% Author Summary not valid for PLOS ONE submissions.   

%\section*{Author summary}
%Decision-making refers to choosing one of the two alternatives by calculating the benefits and costs of each alternative. The choice is determined via a competition between the neural populations in the brain. Visual discrimination experiments on human and macaque were done to study the behavioral background of these competitive neural mechanisms privileging strong, long term benefits rather than weak but immediate benefits \citep{cecchini2024cognitive, humanEbrains, macaqueEbrains}. We complement these experiments by proposing a novel biophysically plausible neural population model taking into account a control over future planning to maximize the benefits in the long term. The model represents two distant cortical columns consisting of excitatory-inhibitory neural populations by using AdEx mean-field equations. Each column votes in favor of one of the two alternatives. Bicolumnar competition is introduced by using intercolumnar excitation which is conflicted by intracolumnar inhibition, in coherence with the anatomy. This framework is integrated with a reward-driven learning, which endows the model with a cognitive context. This learning mechanism captures the optimal strategy to increase the long term benefits. We fit the model to the behavioral data of the human and macaque experiments. The model has the potential to be extended towards the large-scale brain dynamics~\citep{goldman2020brain, turan2023whole} to study decision-making dynamics at the whole-brain level.

%\linenumbers


% Use "Eq" instead of "Equation" for equation citations.
\section{Introduction}

Decision-making refers to choosing one of the existing alternatives. A decision is often made by considering short and long term consequences of these alternatives. In many cases, making decision with a consideration of long term benefits is important for survival. This requires a resistance to immediate decisions bringing benefits in the short term but rather loss in the long term. Such resistance is based on a control over future planning to identify the optimal decision-making strategy bringing long-term, strong benefits.

In complex decision-making tasks, each choice is attributed to a reward. The brain builds a decision-making strategy according to these rewards. It aims to make the choices which maximize the reward. To build such a strategy is not straightforward since a choice might bring a high, immediate reward in the short term, but this reward can decrease in the long term. This happens when the consequences of the choice in the long term are not as beneficial as in the short term. Therefore, insisting on such choices amounts to a weak cumulative reward in the long term. For example, eating junk food gives strong, immediate pleasure, which is a high reward in the short term. However, if it is consumed systematically, it can cause several health problems in the long term, which is a weak reward or rather a loss. In such cases, maximization of cumulative reward requires a long-term planning. This can be provided via a decision-making strategy based on inhibitory self-control over the impulsive decisions.  

Our interest is in such inhibitory self-control over the impulsive decisions. To study it, we focus on two experiments. These experiments are based on two similar reward-driven visual discrimination tasks: one applied on human~\citep{humanEbrains, cecchini2024cognitive} and the other one applied on macaque~\citep{macaqueEbrains}; see Figures~\ref{fig:humanExpScenario} and~\ref{fig:monkeyExpScenario}. These tasks are composed of a certain number of units called episodes. The episodes are composed of subunits called trials. Each episode has the same fixed number of trials. In each trial, two objects are shown on a computer monitor to the subject. The subject chooses one of them. Then, a high reward or a low reward is provided to the subject according to the choice which the subject made. In every episode, both high and low rewards are increased or decreased at the end of each trial. Whether the rewards increase or decrease depends on the decision which the subject made in the trial. The goal is to obtain the maximum possible cumulative reward at the end of all trials in the episode (and in every episode), i.e., the maximum normalized sum of all rewards obtained through the trials.

The particularity of the task is that choosing the high reward stimulus in every trial does not amount to the maximum cumulative reward. There is a specific choice pattern to follow throughout all the trials in an episode. This pattern was designed to provoke the self-control over impulsive decisions. In an example episode with two trials, if the high reward choice is made in the initial trial, the reward amounts (both high and low) will be smaller in the subsequent trial such that the cumulative reward cannot reach to the maximum. Therefore, the subject must follow a specific strategy to obtain the maximum cumulative reward: a strategy which promotes the choice bringing the low reward in the initial trial. This generates an increase in both reward values in the second trial. This is an increase which prevails the difference between the high and low rewards of the first trial. Subsequently, if the subject makes the choice bringing the high reward in the second trial, the cumulative reward achieves the maximum for that episode. This requires a self-control to avoid the decision which bring the high reward in the initial trial.

Our goal is to model the neural population dynamics induced by these experimental tasks. To achieve it, we consider one of the simplest and sufficient scenarios for a decision to be made: two layer 2/3 (L2/3) populations compete and the winning population determines which alternative to select~\citep{desimone1995neural, reynolds2004attentional, moore2017neural}. In this scenario, the competition could be biased in favor of one of the two alternatives.

We propose a neural population model based on three modules for this scenario. The first one is the basic module. It generates the firing rates of the excitatory and inhibitory neurons by using a low dimensional approximation of the Adaptive Exponential (AdEx) neuronal network~\citep{brette2005adaptive}. In the basic module, there are two L2/3 populations. Each population is composed of one pair of excitatory-inhibitory subpopulations. Each excitatory subpopulation votes in favor of one of the two visual alternatives shown on the monitor. These excitatory subpopulations compete with each other to determine the decision. The one with the higher firing rate wins the competition and determines which visual alternative to choose. The second module is the regulatory module. It introduces the bias for either one of the two alternatives shown on the monitor. The final module is the reward-tracking module. It keeps track of the increase or decrease in the reward values, and feeds it into the regulatory module so that the regulatory module generates the bias accordingly: if the reward increases as the model proceeds to the next trials, then the bias favors the choices made in the previous trials, otherwise the bias disfavors them.

We use a mean-field approach to describe the subpopulation firing rates in the basic module. The mean-field equations provide a simple, low dimensional approximation of the averaged dynamics of the AdEx neuronal network~\citep{brette2005adaptive}, which is high dimensional and complex to analyze. This motivates our choice for the mean-field approach. In our model, the mean-field equations of the AdEx neuronal network represents a population of Regulary Spiking (RS, excitatory) and Fast Spiking (FS, inhibitory) cells in L2/3. The L2/3 populations are assumed to communicate via excitatory connections. This represents the network of functional columns found in the prefrontal cortex, as observed in macaque~\citep{hirata2008functional}. Our model examines if two of such interacting populations can implement decision-making mechanisms.  

Several neural population models were previously proposed for similar decision-making problems. Some of them used attractor networks to model switching between two (or more) decisions. Such switching could occur due to perceptual bistability~\citep{moreno2007noise}, or due to further evidence accumulation to make a decision~\citep{albantakis2011changes}. Differently from these, a recurrent network model of object working memory was proposed in~\citep{brunel2001effects}. Thereafter, another network model with a similar architecture was proposed for a visual discrimination task in~\citep{wang2002probabilistic}. This final architecture was adapted to a two-choice perceptual task, where the effect of memory-based experience on the decisions was included and modeled in terms of neural populations~\citep{marcos2013neural, cecchini2024cognitive}. An alternative to these models was proposed in~\citep{hayden2018neuronal}. This framework considers the decision as the result of modulatory effects of attention, on the dynamics of a single population. 

All these previous models are based on connectivity architectures which are not in accordance with the anatomy. In the basic module of our model, the L2/3 populations are connected to each other only via excitatory connections. Each excitatory connection originates from the excitatory subpopulation of one of these L2/3 populations, and it is exerted onto both excitatory and inhibitory subpopulations of the other L2/3 population (Figure~\ref{fig:basicModule}). This is similar to the network of functional columns in the prefrontal cortex, as found in macaque~\citep{hirata2008functional}. Therefore, our model is designed based on an anatomically coherent connectivity. 


It is important to note that, the transfer functions in the AdEx mean-field equations were obtained by fitting membrane variables of a semi-analytical template to \textit{in vitro} measurements~\citep{zerlaut2016heterogeneous}. Then, the free parameters of the same function were fitted to the network responses of RS and FS neurons which were bombarded by the spikes generated from a Poisson distribution~\citep{zerlaut2016heterogeneous, zerlaut2018modeling, di2019biologically}. In the transfer functions, RS neurons display spike-frequency adaptation whereas FS neurons do not, as observed experimentally in the pyramidal neurons and interneurons, respectively. Finally, the AdEx mean-field framework simulates more precisely the inhibition and the excitatory/inhibitory balance through its spike-frequency adaptation compared to other frameworks. 

All these points regarding the connectivity and the transfer functions, provide the biophysically plausible nature of our model. They make our model potentially applicable to neurophysiological recordings from the cortex. This distinguishes our framework from the previous models proposed for similar decision-making tasks.

Another important property of our model is at the modular level. Classical AdEx mean-field framework was proposed for only one population~\citep{zerlaut2018modeling, di2019biologically}, which is not sufficient for a competition resulting in a decision. We extend these equations to two populations (i.e., two subpopulation pairs) so as to provoke a competition between them. Moreover, we integrate this extension with the regulatory and reward-tracking modules. This endows the AdEx mean-field framework with a reward-driven learning mechanism.

These novelties introduce several challenges at both mathematical and numerical levels. Firstly, the extension of the classical AdEx mean-field framework~\citep{zerlaut2018modeling, di2019biologically} to two L2/3 populations increases the dimension of the mean-field system from 6 to 16. Secondly, the connections between these L2/3 populations introduce additional derivatives of the cross-correlations between the subpopulation firing rates. These two points increase memory and computational load. Finally, two noise sources should be incorporated in the model. The first one is introduced to the basic module and it models the distortion effects of the extracellular matrix on the subpopulation firing rates. The second one is introduced to the regulatory module. It models hesitation, curiosity, perceptual difficulties or any other reason causing erroneous decisions. These two types of noise should be introduced in a balanced manner such that they do not dominate each other, neither the deterministic model dynamics.

We test our model on the experimental data in terms of behavioral metrics. Two parameters characterize the performance of the model with respect to these metrics: learning speed and flexibility parameter. The former determines how quickly the model captures the optimal strategy maximizing the cumulative reward. It can be thought of as the modulation effects of acetylcoline and dopamine on the learning dynamics~\citep{brzosko2015, brzosko2017}. The latter determines how much the model is flexible to make mistakes, or to try different decision-making strategies to explore the consequences, sometimes even after that it learns the optimal strategy. 

The implementation of the model is provided in Python as a Jupyter notebook \citep{baspinarJupyter2023}, which can be simulated also online on the EBRAINS cloud system~\citep{baspinarJupyterEbrains2023}.


%{\color{orange}WE MIGHT USE THESE GRAY PARAGRAPHS ELSEWHERE, OR SIMPLY GET RID OF THEM! :)} 

%{\color{gray}
%Finally, the AdEx mean-field framework simulates more precisely the inhibition and the excitatory/inhibitory balance through its spike-frequency adaptation.
%
%In a general decision-making task, the brain encodes and interprets the sensory stimuli corresponding to the choice alternatives. Then, it weights the evidence corresponding to each choice so as to select one of the alternatives. Finally, it generates the neural activity for oriented motor actions in accordance with the selection.
%
%We compare our model simulations to the experiments in terms of two behavioral metrics: performance index and decision time.
%
%We test our model on the decision-making experiments.
%
%
%Moreover, the neural transfer functions which were used in the neural populations were analytical functions in these models, e.g., sigmoid. Their parameters were chosen ad hoc, without any calibration concerning biological measurements of the excitatory and inhibitory neurons found in the cortex.
%}


\section{Experiment setup}\label{sec:experimentalSetup}


The experiments were designed to study the behavioral background of the inhibitory self-control over impulsive decisions bringing immediate rewards, which nevertheless amount to low cumulative reward in the long term. These experiments were performed on human~\citep{cecchini2024cognitive, humanEbrains} and macaque~\citep{macaqueEbrains} by following a different protocol for each species. The difference is based on the type of the visual stimuli and rewards. 

In both protocols, the experiment is composed of a number of episodes. At the beginning of each episode, the protocol is restarted with randomly generated visual stimuli. Each episode is composed of two successive trials. There is a preset strategy which was implemented in every episode of the experiment. This preset strategy refers to a unique pattern of choice sequence. In a single episode, this pattern is based on choosing the stimuli bringing the low reward in the first trial and the stimulus bringing the high reward in the second trial. The key point of the strategy is that, the rewards generated in the second trial are larger if the subject chooses the stimulus which brings the low reward in the first trial. They are larger, in comparison to the rewards generated in the second trial when the subject chooses the stimulus which brings the high reward in the first trial. This increase in the reward values is bigger than the difference between the high and low rewards of the first trial. Therefore, the maximum cumulative reward is achieved only if the subject follows the preset strategy, although the low reward of the first trial gives the opposite impression. 

The goal of the subject is to obtain the maximum possible cumulative reward throughout every episode. This requires: (i) to identify the reward values and the correspondence of these values to the choices made, (ii) to capture the preset strategy in the task to make the sequence of decisions providing the maximum cumulative reward.


\subsection{Human experiment}

The human experiments were conducted at Universitat de Barcelona, Facultat de Matem\`{a}tiques i Inform\`{a}tica. The task was run with 28 participants. We considered a 20-year-old female participant as an example to fit our model. Nevertheless, the model can be fitted to the other participants as well. Here we provide a summary of the task design. More details and relevant data can be found in~\citep{cecchini2024cognitive} and on EBRAINS~\citep{humanEbrains}, respectively. 

Two types of experiments are considered: Horizon 1 and Horizon 0. Each experiment is composed of $K$ episodes, with $K$ denoting the total number of episodes in the experiment. In Horizon 1, each episode is composed of two successive trials, which are named as Trial 1 and Trial 2 (Figure~\ref{fig:humanExpScenario}). In Horizon 0, each episode is composed of a single trial. Therefore, Horizon 0 boils down to a visual discrimination task. Behavioral results are quantified in terms of performance indexes and also reaction times. Performance index provides a measure of the overlap between the choices made by the participant and the preset strategy in an episode. It is registered per episode. Reaction time shows how long it takes for the participant to make the decision in each trial. It is registered per trial. 

It is important to note that the stimuli shown on the monitor are in the form of partially filled vertical bars (Figure~\ref{fig:humanExpScenario}). Moreover, the reward in each trial is the amount of the filled part of the chosen vertical bar. The low reward is equal to the small stimulus, which is the less filled bar. The high reward is equal to the large stimulus, which is the more filled bar. The cumulative reward is the sum of the rewards, i.e., the sum of the filled parts of the chosen stimuli throughout these trials. The preset strategy in Horizon 1 experiments is to choose the smaller stimulus in Trial 1 and the larger stimulus in Trial 2. In the first trial of an Horizon 1 episode, if the choice of the participant overlaps with the preset strategy, the amounts of rewards (thus the filled parts of the stimuli) are increased equally in the second trial. Otherwise, they are decreased equally. The value of these increase and decrease is generated from a random set at the beginning of the experiment and it is kept fixed throughout the whole experiment. 

Before the experiment, the participant is instructed to choose one of the two stimuli projected on the monitor in each trial. Furthermore, the participant is instructed to maximize the cumulative reward for each episode, thus for the whole experiment: the total cumulative reward for the whole experiment is maximum if and only if the participant makes the choices in complete coherence with the preset strategy in every episode of the experiment. In Horizon 0, the strategy boils down to choosing the larger stimulus in each episode, i.e., in each trial.


At the beginning of an episode, the amounts of the filled parts of the bars are generated as $M\pm d/2$. These stimuli are shown at the beginning of Trial 1. Here $M$ denotes the mean value of the stimuli, and $d$ is the amount of difference between the two stimuli. In each episode, $M$ is randomly and independently generated from a uniform distribution at the beginning of the episode. We use one of the five different values of $d$ in each episode. In total, each one of these five values are used equally in $K/5$ episodes of the experiment. In Horizon 1, if the participant chooses the small stimulus in Trial 1, the gain $G>d$ is added to both stimuli and $M+G\pm d/2$ become the filled quantities of the two stimuli in Trial 2. Otherwise, the gain $G$ is subtracted from the stimuli and the filled quantities become $M-G\pm d/2$ in Trial 2. The same procedure is applied in Trial 2 but now the choice according to the preset strategy is the large stimulus. Therefore, there are 4 possible choice patterns which the participant can follow. Each choice pattern corresponds to one of the following 4 cumulative reward values (from maximum to minimum): $2\, M+G$, $2\, M+G-d$, $2\, M-G+d$, and $2\, M-G$. 


We provide an example scenario of an episode in Figure~\ref{fig:humanExpScenario}. In Trial 1 of this scenario, the participant chooses the large stimulus with $M+d/2$ (i.e., the high reward). This does not overlap with the preset strategy. Therefore, the stimuli (i.e., the rewards) are decreased by $G$, resulting in $M-G+ d/2$ and $M-G-d/2$ as large and small stimuli values, respectively. In Trial 2, the participant chooses the large stimuli (i.e., high reward), and this overlaps with the preset strategy. The resultant cumulative reward is $2M-G+d$, which shows that the participant did not capture the preset strategy. 

Finally, an important particularity of the human experiment is that the participant does not see at the end of Trial 2 the cumulative reward, or any explicit feedback. As a result of this lack of feedback at the end of the episode, the participant is forced to learn the consequence of the episode instead of learning a sequence of choices. See Figure~\ref{fig:exampleExperimentResults} for some relevant example experimental results.


%In Figure~\ref{fig:humanExpScenario}, we provide an example scenario of one episode of Horizon 1. In this example, the amounts of the filled parts of the bars are $M\pm d/2$ at the beginning of Trial 1. Here $M$ is randomly and independently generated from a uniform distribution at the beginning of each episode. Five different values of $d$ are used in the experiments. Each one of them is fixed during $K/5$ episodes of the experiment. Here $M$ denotes the mean value of the stimuli, and $d$ determines the difference between the two  stimuli. In Horizon 1, if the participant chooses the small stimulus in Trial 1, the gain $G>d$ is added to both stimuli and $M+G\pm d/2$ become the filled quantities of the two stimuli in Trial 2. Otherwise, the gain $G$ is subtracted from the stimuli and the filled quantities become $M-G\pm d/2$ in Trial 2, as in the case of Figure~\ref{fig:humanExpScenario}. The same procedure is applied in Trial 2 but now the choice according to the preset strategy is the large stimulus. The participant can obtain one of the 4 different cumulative reward values: $2\,M + G$, $2\, M + G-d$, $2\, M-G + d$, and $2\, M-G$ (ordered from the best to the worst). Moreover, the participant does not see at the end of Trial 2 any added or subtracted gain since it is the end of the episode. As a result of this lack of feedback at the end of the episode, the participant is forced to learn the consequence of the episode instead of learning a sequence of choices. See Figure~\ref{fig:exampleExperimentResults} for some relevant example experimental results.
       
\begin{figure}[!htbp]
\begin{minipage}[c]{\textwidth}
%\centering
\sidesubfloat[(a)]{\hspace{1.4cm}
\includegraphics[clip,width=0.7\columnwidth]{humanExpScenario.png}
\label{fig:humanExpScenario}
}

\bigskip
%\bigskip
%\bigskip
  
\sidesubfloat[(b)]{\hspace{-1cm}
  \includegraphics[clip,width=0.95\columnwidth]{monkeyExpScenario.png}%
\label{fig:monkeyExpScenario}
}
\end{minipage}

\caption[\labelsep=space]{Example episodes from the Horizon 1 human and macaque experiments shown in (a) and (b), respectively. \textbf{(a)} A central target (CT) appears in Frame 1 to indicate the beginning of the episode. A progress bar at the bottom indicates in which trial the participant is. The stimuli with values $M-d/2$ and $M+d/2$ are shown separately in Frames 2 and 3, then together in Frame 4. The participant moves the pointer to the larger stimulus as highlighted by the yellow dot in Frame 5. The end of the episode is indicated in Frame 6 with a dot at the center. The stimuli are decreased in Episode 2, as shown separately in Frames 7 and 8. They are shown together in Frame 9 and the participant chooses the larger one as highlighted in Frame 10. The cumulative reward is the sum of the chosen stimuli. \textbf{(b)} A CT appears in Frame 1 to indicate the beginning of the episode. The subject touches the CT to initiate the episode. The peripheral targets (PTs) are shown together on the both sides of the CT in Frame 2. The CT disappears as the participant starts to move its hand to choose one of the PTs as shown in Frame 3. The chosen PT (PT 1) is highlighted by the yellow dot in Frame 4. The corresponding reward (3 water drops) is provided, and in Frame 5, a blank screen is shown to indicate the end of Trial 1. The same procedure is followed in Trial 2 but now with the PTs of Group 1. In the second scenario, the cumulative reward is higher since PT 2 is chosen in Trial 1 and hence the Group 2 PTs, which provide a higher reward, are shown on the monitor in Trial 2. The nomenclature and corresponding rewards to the PTs are given at the top. The cumulative reward is the sum of the provided water drops.}
\end{figure}


\begin{figure}[!htbp]
\begin{minipage}[c]{\textwidth}
\sidesubfloat[(a)]{\hspace{3cm}
\includegraphics[clip,width=0.44\columnwidth]{learningTime.png}
\label{fig:expHuman1}
}

\bigskip


\sidesubfloat[(b)]{\hspace{1.5cm}
\includegraphics[clip,width=0.7\columnwidth]{performanceCDF.png}
\label{fig:expHuman2}
}
\end{minipage}

\caption[\labelsep=space]{Some experiment results from~\citep{cecchini2024cognitive}. \textbf{(a)} The histogram of the learning times obtained from 28 human subjects. The learning time is in terms of episode numbers. The long learning time is classified as no learning. \textbf{(b)} The performance indexes of one of the participants, and the cumulative distribution function (CDF) of the reaction times of the same participant. The results are presented separately for Horizon 0 and Horizon 1, where in the latter, the CDFs of Trial 1 and Trial 2 are provided in green and blue, respectively.}
 \label{fig:exampleExperimentResults}
\end{figure}


\FloatBarrier

\subsection{Macaque experiment}

The macaque experiments were conducted at Sapienza Universit\`{a} di Roma, Dipartimento di Fisiologia e Farmacologia.  One 16-year-old, healthy, male macaque weighting $9.5 - 10.5$ kg was trained for performing a Horizon 1 task similar to its counterpart in the human experiment. The relevant data can be found on EBRAINS~\citep{macaqueEbrains}. 

The task can be summarized based on the two possible scenarios provided in Figure~\ref{fig:monkeyExpScenario}. A dataset of black and white stimuli were shown to the macaque in each trial. Six stimuli (peripheral targets; PTs) were extracted randomly from \textit{Microsoft PowerPoint} shape library. Two of these six randomly extracted PTs were shown in Trial 1 and they were attributed to different rewards; PT 1: 3 water drops, PT 2: 2 drops. The remaining four stimuli were separated into two groups, each based on two PTs. In Group 1, we had PT 3 and PT 4; in Group 2, we had PT 5 and PT 6. Each of these four PTs corresponds to a different reward; PT 3: 1 drop, PT 4: 0 drop, PT 5: 4 drops and PT 6: 6 drops. The group which is shown in Trial 2 was determined by the choice made in Trial 1. More precisely, if the participant chooses PT 1 in Trial 1, Group 1 will be shown in Trial 2 as illustrated in Episode 1 of Figure~\ref{fig:monkeyExpScenario}. Otherwise, Group 2 will be shown in Trial 2 as illustrated in Episode 2 of Figure~\ref{fig:monkeyExpScenario}. At the beginning of each trial, a central target (CT) is shown at the center of a touchscreen monitor to indicate the beginning of the trial. The macaque is required to touch the CT to initialize the trial. Then, the stimuli are shown as the PTs appearing on the left and right hand sides of the CT. As the participant starts to move his hand, the CT disappears. This is considered as the Go signal initiating the reaction time, which is tracked for a maximum duration of 2000 ms. If a choice is made between two PTs within this 2000 ms, the reward is delivered to the subject. Once this protocol for Trial 1 is completed successfully, Trial 2 begins after an inter-trial interval of 1200 ms and it follows the same protocol.


Although both human and macaque tasks are based on perceptual distinguishing of the targets, there are three major differences between them. Firstly, the stimuli are partially filled bars in the human task, whereas they are some figures chosen randomly from the Powerpoint shapes library in the macaque task. Secondly, in the human task, the rewards are the same as the chosen stimuli. High and low rewards are the large and small stimuli, respectively. In the macaque task, the reward is provided as a certain number of water drops to the macaque. Finally, the reward is provided at the end of Trial 1 and Trial 2 in the macaque task. In the human task, there is an explicit feedback regarding the reward only at the end of Trial 1. There is no feedback at the end of Trial 2. 


\section{Model}\label{sec:model}

We study the neural dynamics regarding the human and macaque experiment by using a neural population model. The model is composed of three layers: basic module, regulatory module, and reward-tracking module.

\subsection{Basic module}

 
The basic module generates the neural activity in terms of subpopulation firing rates. It describes the firing rates by using an extension of the classical AdEx mean-field equations~\citep{di2019biologically} to two L2/3 populations. 

The classical AdEx mean-field equations were derived in~\citep{zerlaut2018modeling, di2019biologically} from the AdEx neuronal network~\citep{brette2005adaptive}. They describe the average dynamics of a pair of excitatory and inhibitory subpopulations which are coupled to a spike-frequency adaptation. We extend it to two pairs, where each pair is a L2/3 population composed of one excitatory and one inhibitory subpopulation (Figure~\ref{fig:basicModule}). In this extended setting, each L2/3 population votes for either one of the two stimuli: Population $A$ votes for Stimulus A and Population $B$ votes for Stimulus $B$. We denote Stimuli $A$ and $B$ with $s_A$ and $s_B$, respectively. One of them represents the choice bringing the high reward and the other one represents the choice bringing the low reward. Stimulus $A$ and $B$ may represent these choices interchangeably in different episodes. However, they are assigned to either one of the choices within a single episode, and it does not change throughout the episode. In the basic module, we denote the excitatory subpopulations by $e_A, e_B$ and the inhibitory subpopulations by $i_A, i_B$. The subindexes $A$ and $B$ indicate the corresponding L2/3 population. Each L2/3 population represents a functional column of $N_e$ excitatory and $N_i$ inhibitory neurons. The ratio $N_i/N_e$ is $0.2$.

The basic module connectivity follows an anatomically coherent connectivity architecture. Across the L2/3 populations, there are only excitatory connections. This connectivity is similar to the network of functional columns in the prefrontal cortex, as found in macaque~\citep{hirata2008functional}. Each one of these excitatory connections between the L2/3 populations originates from the excitatory subpopulation of one L2/3 population and targets both the excitatory and inhibitory subpopulations of the other L2/3 population (Figure~\ref{fig:basicModule}). The weights of the connections between the L2/3 populations are denoted by $w_c$. Local connections are within the populations. They are based on all-to-all connectivity within each population, with both recurrent and cross-subpopulation connections as shown in Figure~\ref{fig:basicModule}. All the local connections have unit weights.


Similarly to the classical AdEx mean-field setting~\citep{di2019biologically}, the basic module can function in two modes: asynchronous irregular (AI) and up-down. Here AI mode refers to the awake state of the brain and up-down refers to the sleep (or under anesthesia) state of the brain. We consider only the AI state since decision-making requires that the subject is awake. Therefore, we keep the basic module in AI mode by feeding it with a constant base drive $s_{\text{AI}}$. This is a DC input (=5 Hz) which keeps the basic module away from the up-down mode. The terms $\lambda^A(s_A, s_B)$ and $\lambda^B(s_A, s_B)$ in Figure~\ref{fig:basicModule} represent the inputs to Populations A and B, respectively. These are the biased versions of the stimuli $s_A, s_B$. The bias is introduced by a regulatory module that models plasticity as explained in Section~\ref{sec:regulatoryModule}. The biased inputs are simultaneously fed to Populations $A$ and $B$ through the functions $\lambda^A$ and $\lambda^B$ described in~\eqref{eq:lambdaExpression}, respectively.

We denote the firing rates of the four subpopulations by $v_{e_A}, v_{i_A}, v_{e_B}, v_{i_B}$. Each one of them represents the average spike frequency of one of the four subpopulations. We describe the neural interactions between these subpopulations via cross-correlations between the firing rates of interacting subpopulations. For simplicity of notation in the following, we use $\alpha, \beta, \xi, \eta$ to denote any of the four subpopulations $e_A, e_B, i_A, i_B$. Then, we represent the cross-correlation between $v_\alpha$ and $v_\beta$ with $C_{\alpha\beta}$. Finally, the variables $W_{e_A}$ and $W_{e_B}$ describe the spike-frequency adaptation of the RS cells, i.e., the excitatory subpopulations $e_A$ and $e_B$, respectively. The adaptation is only for the excitatory subpopulations as FS cells have no spike-frequency adaptation. Thus, we assume no adaptation for the inhibitory subpopulations, i.e., $W_{i_A}=W_{i_B}=0$. As a result, the basic module has 16 state variables.


The basic module equations read as:
\begin{equation}\label{eq:mfEqns}
\begin{split}
T\,\partial_t v_{\alpha} = & (F_\alpha - v_\alpha) + \frac{1}{2}\sum_{\xi,\eta}C_{\xi\eta}\partial_{\xi\eta}F_\alpha + \sigma\,\omega_\alpha\\
T\, \partial_t C_{\alpha\beta} = & A_{\alpha\beta}^{-1} + (F_\alpha-v_\alpha)(F_\beta-v_\beta)+\sum_{\xi, \eta}(C_{\beta\xi}\partial_\xi F_\alpha + C_{\alpha\xi}\partial_\xi F_\eta) - 2C_{\alpha\beta}\\
\partial_t W_{e_A} = & -\frac{W_{e_A}}{\tau_w} +  b\, v_{e_A} + a\Big ( \mu_V(v_{e_A}, v_{e_B}, W_{e_A} ) - E_L \Big ),\\
\partial_t W_{e_B} = & -\frac{W_{e_B}}{\tau_w} + b\, v_{e_B} + a\Big ( \mu_V(v_{e_A}, v_{e_B}, W_{e_B} ) - E_L \Big ),
\end{split}
\end{equation}
where we use the notation given by $\partial_t\coloneqq\frac{\partial}{\partial t}$, $\partial_\alpha \coloneqq \frac{\partial}{\partial v_\alpha}$ and $\partial_{\alpha\beta} \coloneqq \frac{\partial^2}{\partial {v_\alpha}\partial_{v_\beta}}$ for the partial derivatives. Here $E_L$ is a constant representing the reverse leakage potential, and $T,\tau_w$ are the time scale parameters. We denote by $\omega_\alpha$ a white Gaussian noise, with $\sigma>0$ denoting its intensity. This noise is sampled from a normal distribution independently for each subpopulation and at each instant. More precisely, $\mathbb{E}[\omega_\alpha(t)]=0,\quad\mathbb{E}[\omega_\alpha(t) \omega_\beta(t^\prime)]=\delta_{\alpha\beta}\, \delta_{tt^\prime}$ for all $\alpha,\beta\in \{e_A, i_A, e_B, i_B\}$ and for all $t,t^\prime\geq 0$, where $\delta$ is the Dirac delta function. Here $\mathbb{E}[ \;\cdot\; ]$ represents the expected value. Function $A_{\alpha\beta}$ appears from the mean-field derivation of the second order statistical moments and it is defined for $\alpha,\beta\in \{e_A, i_A, e_B, i_B\}$ as follows:
\begin{equation}
\label{eq:Aalpha}
A_{\alpha\beta}=\begin{cases}\frac{T\,N_\alpha}{F_\alpha-T\,F_\beta},\quad\quad \text{if\;} \alpha=\beta\\
0\quad\quad\quad\quad\quad \text{otherwise,} 
\end{cases}
\end{equation}
where $N_\alpha$ denotes the number of neurons represented in the subpopulation $\alpha$. We choose $N_{e_A}=N_{e_B}=8000$ and $N_{i_A}=N_{i_B}=2000$.

% It should be chosen properly not to violate Markovian assumption required in the derivation of the mean-field approximation of the AdEx neuronal network ~\citep{di2019biologically}.

An important group of functions appearing in the mean-field equations~\eqref{eq:mfEqns} is the transfer functions, denoted by $F_\alpha$. A transfer function gives the output firing rate of a subpopulation according to the input that the subpopulation receives. This output firing rate is modulated by the spike-frequency adaptation in the excitatory subpopulations. The transfer functions appearing in~\eqref{eq:mfEqns} and~\eqref{eq:Aalpha} are written with their inputs as follows:
\begin{equation}\label{eq:transferFcnsRegulated}
\begin{split}
F_{e_A} & =  F_{e_A}(\tilde{v}_{e_A}, \tilde{v}_{i_A}, W_{e_A}),  \quad\quad
F_{i_A}= F_{i_A}(\tilde{v}_{e_A}, \tilde{v}_{i_A}, W_{i_A}=0),\\
F_{e_B} & =  F_{e_B}(\tilde{v}_{e_B}, \tilde{v}_{i_B}, W_{e_B}), \quad\quad
F_{i_B}= F_{i_B}(\tilde{v}_{e_B}, \tilde{v}_{i_B}, W_{i_B}=0),
\end{split}
\end{equation}
with subindexes indicating the subpopulation type. Each input to the transfer functions is composed of excitatory and inhibitory components $\tilde{v}_{e_A}, \tilde{v}_{e_B}$ and $\tilde{v}_{i_A}, \tilde{v}_{i_B}$, respectively. Each component has a local part and a part related to the interactions between the L2/3 populations. The latter is weighted by $w_c$ since the interactions take place through the connections between the L2/3 populations. These excitatory and inhibitory components are as follows:
\begin{equation}\label{eq:inputTF_noRegulation}
\begin{split}
\tilde{v}_{e_A}(t)=  v_{e_A}(t)+ &  s_{\text{AI}}+  \lambda^A(s_A(t), s_B(t))\\
	& +w_c \Big ( v_{e_B}(t) + s_{\text{AI}}+\lambda^B(s_A(t), s_B(t), t) \Big ),\\ 
\tilde{v}_{i_A}(t)=  v_{i_A}(t)+ & \lambda^A(s_A(t), s_B(t))\\
 & + w_c \Big ( v_{e_B}(t) + s_{\text{AI}}+\lambda^B(s_A(t), s_B(t), t) \Big ),\\
\tilde{v}_{e_B}(t) =  v_{e_B}(t) + & s_{\text{AI}}+\lambda^B(s_A(t), s_B(t)) \\ 
  & + w_c \Big ( v_{e_A}(t) + s_{\text{AI}}+\lambda^A(s_A(t), s_B(t), t) \Big ),\\
\tilde{v}_{i_B}(t)=  v_{i_B}(t)+ & \lambda^B(s_A(t), s_B(t)) \\
   & + w_c \Big ( v_{e_A}(t)+ s_{\text{AI}}+\lambda^A(s_A(t), s_B(t), t) \Big ).
\end{split}
\end{equation}
Here the time dependency is explicitly denoted and the terms with no explicit time variable are constants. The external stimuli $s_A, s_B$ are fed in~\eqref{eq:inputTF_noRegulation} via $\lambda^A, \lambda^B$. It is to take into account the bias introduced through the regulatory module, which will be explained in Section~\ref{sec:regulatoryModule}. A common choice to model the external stimuli $s_A,s_B$ is Heaviside function in mean-field models. However, since the Heaviside function has a discontinuity due to the discrete jump, our model undergoes a transient phase which might obscure the instant when the model makes a decision. For this reason, we perform our simulations by using a sufficiently sharp sigmoid function as external stimulus; see Figure~\ref{fig:differentA_B_Stimuli}. Finally, $s_\text{AI}$ denotes the DC input which keeps the basic module in the AI state. 

We use the same formula for the transfer functions as provided in~\citep{di2019biologically}; see Appendix for the details. We report here the formula directly:
\begin{equation}\label{eq:transferFunction}
F_{e_A}(\tilde{v}_{e_A}, \tilde{v}_{i_A},W_{e_A})=\frac{1}{2\,\tau_V(\tilde{v}_{e_A}, \tilde{v}_{e_B})}\,\operatorname{erfc}\Big (\frac{v_{\text{eff}}-\mu_V(\tilde{v}_{e_A}, \tilde{v}_{e_B})}{\sqrt{2}\, \sigma_V(\tilde{v}_{e_A}, \tilde{v}_{e_B})} \Big ).
\end{equation}
This formula is for $e_A$ and it is the same for the other subpopulations once the subindexes are changed accordingly. Here, $\mu_V$ and $\sigma_V$ represent the mean and standard deviation of the membrane voltages of the neurons found in the AdEx network, respectively. Their formulas can be found in~\eqref{eq:forTF1} and~\eqref{eq:forTF2}. The term $\tau_V$ is the global autocorrelation time of the membrane voltage fluctuations of the neurons in the AdEx network; see~\eqref{eq:forTF2} in Appendix. It gives a measure of the speed of the fluctuations. Finally, $v_{\text{eff}}$ is the voltage-effective threshold and it is a second order polynomial of $\mu_V, \sigma_V, \tau_V$; see~\eqref{eq:forTF3}. It was obtained by a fit according to the simulations of single neuron activity produced at multiple excitatory and inhibitory presynaptic frequencies~\citep{zerlaut2018modeling, di2019biologically}. The transfer function~\eqref{eq:transferFunction} was derived via a semianalytical procedure based on the output spiking of an AdEx network of $10^4$ neurons, where each neuron was modeled by the AdEx integrate-and-fire equations given by~\eqref{eq:AdExNetwork}. The network was bombarded with spikes generated from a Poisson distribution and a semi-analytical function was fitted to the outputs of the AdEx network~\citep{di2019biologically}. See Appendix for more details.

The derivation of the AdEx mean-field equations from the AdEx neuronal network was provided in~\citep{di2019biologically}. In the present paper, we use the same mean-field equations to describe the dynamics of each L2/3 population. The extension to two L2/3 populations is made symmetrically. That is, the L2/3 populations are set to the identical parameter values and they have the identical local connectivity architecture within the populations. Accross the L2/3 populations, the coupling is only excitatory and all the excitatory connections between the L2/3 populations have the same connection weight ($=w_c$). This allows us to write the mean-field equations of our extended framework directly from the equations provided in~\citep{di2019biologically}, avoiding a derivation from an AdEx network of four subpopulations.

%across the pools an Therefore, the interactions among the four populations can be described by the   This allows us to derive the equations for four populations by starting from   
%{\color{orange}--WE SHOULD THINK ABOUT HOW TO DESCRIBE THIS TERM AND ALSO THE TRANSFER FUNCTIONS. WE MIGHT PARAPHRASE THE CORRESPONDING PARTS FROM DI VOLO PAPER!!!----}The term $\mu_V$ in~\eqref{eq:mfEqns} is the average membrane potential of the population{\color{red}---We should explain the term $\mu_V$ quickly here or in the appendix!!!!---or see the orange colored Section 3.1.1 just below!!!}. It is the same term as the one given in~\citep[Section 2.3.1]{di2019biologically}, so we provide there the final formula directly:
%\begin{equation}
%\mu_V(v_e, v_I, W_{e_A})\coloneqq \frac{\mu_{G_{e_A}}E_{e_A}+\mu_{G_{}}}{}
%\end{equation}
%
%{\color{orange}
%\subsubsection{Transfer functions and average membrane potential}
%
%WE MIGHT ADD A SMALL SECTION HERE TO EXPLAIN THE TRANSFER FUNTION CALCULATION BY PARAPHRASING FROM diVOLO, or WE CAN ADD SUCH A SECTION TO THE APPENDIX !}

\subsection{Regulatory module}
\label{sec:regulatoryModule}

The basic module is not associated with any learning mechanism. Nevertheless, a learning mechanism is required to capture the preset decision-making strategy, which gives the maximum cumulative reward. To introduce a learning mechanism, we integrate the basic module with a regulatory module. The regulatory module identifies the reward quantities and how they change according to the choices made by the model. Then, it introduces the bias to the external stimuli in such a way that, the stimuli corresponding to the choices which bring large cumulative reward are promoted. This module can be thought of as a gating mechanism arranging the motor-plan flexibility for the response to the stimuli (or to the sensory inputs evoked by the stimuli). This mechanism might provide an explanation to the observations showing that relevant sensory inputs, distractors, and direct perturbations of decision-making circuits affect the behavior more strongly when they are introduced in the early phase of the stimulation \citep{seidemann1998temporal, kiani2008bounded, kopec2015cortical, van2018decoding, zuo2019rats, finkelstein2021attractor}.

The regulatory module was adapted from~\citep{cecchini2024cognitive}. It introduces the bias by weighting the stimuli fed into the L2/3 populations. The bias makes the population voting for the promoted stimulus to be more likely to have a higher excitatory firing rate than the other population. Consequently, the population voting for the promoted stimulus is more likely to win the competition and to make the decision; see Figures~\ref{fig:withRewardFullSys1} and~\ref{fig:psiAllTypes}. We denote by $\psi_n^E$ the regulatory function evolving during the $n^{\text{th}}$ trial of the $E^{\text{th}}$ episode. It is described via
\begin{equation}\label{eq:regulatory_Cauchy}
\begin{cases}
\tau_\psi \frac{d\psi^E_n(t)}{dt} = -4 \psi^E_n(t) \Big ( \psi^E_n(t)-1 \Big )\Big ( \psi^E_n(t)-1/2 \Big )+\frac{\sigma}{(c_0 t)^2}\zeta_n(t),\quad\quad  t\in (0,t_F],\\
\psi_n^E(0)= \phi^{E-1}_n,
\end{cases}
\end{equation}
where $\tau_\psi$ is the time scale parameter. The regulatory function is restarted at the beginning of each trial. It evolves simultaneously with the basic module in time. Here $t_F>0$ is the final time of the trial and it is the same for every trial. Moreover, $\zeta_n = \zeta_n(t)$ is a white Gaussian noise. Its intensity level $\sigma>0$ is scaled by a constant $c_0>0$ and time. Therefore, this noise introduces a strong stochastic behavior initially. Then, it decays in time with a rate depending on $c_0$. It models the perceptual difficulties, hesitation and curiosity/exploratory behavior of the participant.

%In an episode $E$, at each trial $i$, both stimuli are shown to the participant. The stimuli have an excitatory effect on all the populations.
The decision in a trial depends on the evolution of the function $\psi_n^E$ in time. This function converges to either 1 or 0 in the course of each trial. The convergence towards 1 pushes the model to choose the stimulus bringing high reward, e.g., the large stimulus in the human experiment. The convergence towards 0 pushes the model to choose the stimulus bringing low reward, e.g., the small stimulus in the human experiment. These dynamics of the regulatory module are continuously transmitted to Populations $A$ and $B$ via
\begin{equation}\label{eq:lambdaExpression}
\begin{split}
\lambda^A(s_A(t), s_B(t), t) = & \psi_n^E(t)\,s_A(t) + (1-\psi_n^E(t))\,s_B(t),\\
\lambda^B(s_A(t), s_B(t), t) =  & \psi_n^E(t)\,s_B(t) + (1-\psi_n^E(t))\,s_A(t),
\end{split}
\end{equation}
where $t\in [0, t_F]$.

It is important to note that the process given by~\eqref{eq:regulatory_Cauchy} is reinitialized at the beginning of each trial $n$ of the $E^{\text{th}}$ episode. It is reinitialized from the initial condition fixed to the value determined by the reward-tracking function $\phi^E_{n-1}$, which will be explained in the next section. The function $\phi^E_{n-1}$ corresponds to the same trial number $n$, but of the $(E-1)^{\text{th}}$ episode. In this way, the reward-tracking function provides at the end of the $(E-1)^{\text{th}}$ episode, a separate feedback for each trial $n$ of the $E^{\text{th}}$ episode. This feedback is used as the initial condition for the regulatory function equations~\eqref{eq:regulatory_Cauchy}, at the beginning of each trial of the $E^{\text{th}}$ episode. The feedback provides information about how optimal the decision-making strategy of the previous episode was, therefore about the accumulated experience. In this way, the feedback determines towards which decision the regulatory module will introduce the bias to the basic module in the next episode.


 
%\begin{figure} 
%  \centering
%  \includegraphics[scale=1.25]{energyFunctionalConverge.png}
%  \caption{Regulatory module. Left: Energy functional of $\psi$. The neutral initial condition is $\psi(0)=0.5$. The high and low rewards introduce a bias making $\psi(0)>0.5$ or $\psi(0)<0.5$ in the next episode. This results in $\psi=1$ or $\psi=0$ at the end of the corresponding trial of the next episode. The former introduces the bias favoring the large stimulus and the latter introduces the bias favoring the small stimulus. Right: Time evolution of $\psi$ starting from different initial conditions. At the beginning, a small noise is introduced for modeling whatsover which might perturb the regulatory process, such as the exploratory behavior or perceptual difficulties.}
%  \label{fig:energyFunctionalConverge}
%\end{figure}


 


\begin{figure}[!htbp]
\begin{minipage}[c]{\textwidth}
\sidesubfloat[(a)]{\includegraphics[clip,scale=0.5]{basicModule}\label{fig:basicModule}
}

\bigskip


\sidesubfloat[(b)]{\hspace{0.6cm}\includegraphics[clip,scale=1.25]{energyFunctionalConverge}\label{fig:energyFunctionalConverge}
}

\end{minipage}
\caption{Model structures. \textbf{(a)} Basic module with two L2/3 populations composed of excitatory and inhibitory subpopulations. Populations $A$ and $B$ vote in favor of Stimulus $A$ ($s_A$) and Stimulus $B$ ($s_B$), respectively. The excitatory subpopulations are denoted by $e_A, e_B$ and the inhibitory subpopulations are represented with $i_A, i_B$. The excitatory and inhibitory connections are in blue and red, respectively. The weights of the connections between the L2/3 populations are denoted by $w_c$ as in~\eqref{eq:inputTF_noRegulation}. Here $\lambda^A$ and $\lambda^B$ represent the outputs of the regulatory module introducing a bias to the input signals $s_A$ and $s_B$. Finally, $s_{\text{AI}}$ is the base drive, which ensures that the model performs in the awake state. \textbf{(b)} Regulatory module. Left: Energy functional of $\psi$. The neutral initial condition is $\psi(0)=0.5$. The increase and decrease in the rewards introduce a bias making $\psi(0)>0.5$ or $\psi(0)<0.5$ in the next episode. This results in $\psi(t_F)=1$ or $\psi(t_F)=0$ at the end of the corresponding trial of the next episode. The former introduces the bias favoring the large stimulus and the latter introduces the bias favoring the small stimulus. Right: Time evolution of $\psi$ starting from different initial conditions. At the beginning, a small noise is introduced for modeling whatsover which might perturb the regulatory process, such as the hesitation, curiosity/exploratory behavior or perceptual difficulties.}
\label{fig:modelStructures}
\end{figure}

\FloatBarrier


\subsection{Reward-tracking module}

The regulatory module introduces the bias depending on a feedback. This feedback is provided as the initial condition to the regulatory function equations given by~\eqref{eq:regulatory_Cauchy}. We should take into account the accumulated experience throughout the previous episodes to provide the feedback for the upcoming episodes. This feedback must be such that the model can identify which choices contribute more to the cumulative reward. We integrate our model with a reward-tracking module to endow the model with such feedback mechanism.

The reward-tracking module is based on a reinforcement learning equation. This equation motivates the choices which increase the reward amounts, and demotivates the choices which decrease the reward amounts. This is associated with the increasing dopamine signaling during predicted reward anticipation: motivation to repeat the same decision and the dopamine signaling increase, if the obtained reward is as predicted or more~\citep{mirenowicz1994importance, schultz2022dopamine}. This results in an online learning. Once the preset strategy is learned, the system makes the decisions in almost complete coherence with the strategy.

We assume in the human experiments that the increase and decrease in the stimuli (i.e., reward) is predicted as profit and loss by the participants, respectively. In accordance with this assumption, the reward-tracking module is updated through a discrete evolution over the episode numbers. In other words, the reward-tracking function value remains constant during a trial and it is increased or decreased to another constant at the end of the trial. The updated value is provided to the regulatory function $\psi$ in the next episode as the initial condition; see~\eqref{eq:regulatory_Cauchy} and~\eqref{eq:rewardMechanismEqn}.

In the human case, we use the notation $M^E_n$ to denote the mean value of the stimuli corresponding to the $n^{\text{th}}$ trial of the $E^{\text{th}}$ episode. We consider here only Horizon 1 since Horizon 0 uses the same module but with $n=1$. We write the evolution for the reward-tracking function $\phi$ as follows:
\begin{equation}\label{eq:rewardMechanismEqn}
\begin{cases}
\phi^{E+1}_n =   \phi^E_n + k\,  ( M^E_{n+1} - M^E_n ) (2\psi_n^E(t_{F})-1)(\phi^E_n  -1 )^2(\phi_n^E)^2,\\
\phi^1_n = C,\quad \text{for all}\quad n\in \{1, 2\},
\end{cases}
\end{equation}
with $C$ denoting a constant, which is fixed to $0.5$ in our framework. Here $k$ is the learning speed parameter. It determines how quickly the model captures the preset strategy throughout the episodes. In~\eqref{eq:rewardMechanismEqn}, $(M^E_{n+1} - M^E_{n})$ provides a coupling between the trials of an episode. It changes the sign of the polynomial. In this way, it determines how the bias provided by $(2\psi_n^E(t_{F})-1)$ will be transmitted to the upcoming episode (i.e., to the episode $E+1$): by favoring or by disfavoring the choice made in episode $E$? The same choice is favored in the upcoming episode if $(M^E_{n+1} - M^E_{n})>0$, meaning that the stimuli (i.e., the rewards) increased as we pass from $n^{\text{th}}$ to $(n+1)^\text{th}$ trial. It is disfavored if $(M^E_{n+1} - M^E_{n})<0$, meaning that the stimuli (i.e, the rewards) decreased.

In the reward-tracking module for the macaque case, we use a similar idea to the human case, with a few differences. Let us recall that in the macaque experiment, the reward is provided to the subject as water drops. This is different from the human experiment, where the reward is the chosen stimulus itself. Therefore, we write the reward-tracking equations for the macaque case, in terms of the water drops provided for each choice. We replace the first line of \eqref{eq:rewardMechanismEqn} with
\begin{equation}\label{eq:macaqueRewardTracking}
\phi^{E+1}_n =   \phi^E_n + k\,  F_n\, (2\psi_n^E(t_F)-1)(\phi^E_n  -1 )^2(\phi_n^E)^2,
\end{equation}
where
\begin{equation}
F_n^E:=(\delta_{n-1}(\text{other}^E_1-\text{choice}^E_1) +  \delta_{n-2}(\text{choice}^E_2 - \text{other}^E_2)),
\end{equation}
with $n$ and $\delta$ denoting the trial number and the Dirac delta, respectively. Here $\text{choice}_n^E$ and $\text{other}^E_n$ denote the numbers of water drops corresponding to the chosen stimulus and to the other stimulus, respectively, in the $n^{\text{th}}$ trial of the $E^{\text{th}}$ episode. Here the reward is explicit. It is not represented in terms of the stimuli as in the human case.


\subsection{Example simulation of the model}

We provide here a small example simulation of the model to give an insight about the functionality of the three modules. In this example, we consider a human Horizon 1 simulation with 6 episodes. 

In the human Horizon 1 experiments, the preset strategy requires choosing the smaller stimulus (the bar which is filled less) in Trial 1 and the larger stimulus (the bar which is filled more) in Trial 2 of each episode. This preset strategy is the optimal strategy which the model should learn so as to have the maximum cumulative reward. In~Figure~\ref{fig:withRewardFullSys1}, we show a small example with 6 episodes of our Horizon 1 model simulations regarding the human case. The reward-tracking module $\phi$ is initiated from 0.5 for both trials in Episode 1, with a noise perturbing this initial value at the beginning of each trial. We observe that the model explores the strategy in Episode 1. Its decision is completely random with no bias since $\psi^1_i(0) = 0.5$ for both $i=1$ and $i=2$. In Episode 2, the reward-tracking module introduces a bias in the regulatory function $\psi$ based on the feedback from Trial 1 and Trial 2 of Episode 1. The same procedure is repeated for Episode 3 but now based on the feedback from Episode 2, and so on. In Figure~\ref{fig:withRewardFullSys1}, we observe that the model learns the strategy in Episode 2. From Episode 2 onwards, it makes the choices in complete coherence with the preset strategy. 

%We use the same parameters given in~\eqref{eq:parametersSmoothtim1}-\eqref{eq:parametersSmoothStim3} for these simulations.

%The speed of learning the preset strategy depends on the parameter $k$ given in~\eqref{eq:rewardMechanismEqn}. We name $k$ learning speed. In Figure~\ref{fig:withRewardFullSys1}, we chose $k = 0.1$ and we fix the gain $G$ to $0.75$. The gain is added to the stimuli if the correct decision according to the preset strategy is made, otherwise, it is subtracted from the stimuli; see Figure~\ref{fig:humanExpScenario}.

\begin{figure}[!h] 
  \includegraphics[width=1.02\textwidth]{withRewardFullSys1.png} 
  \caption{Simulation results of $6$ episodes. First row: Time courses of the excitatory subpopulation firing rates. Second row: Time courses of the difference of the excitatory subpopulation firing rates. Third row: Time course of the regulatory function $\psi$ for each trial of the corresponding episode given in the bottom row. Fourth row: Time course of the reward function $\phi$ for Trial 1 (top) and Trial 2 (bottom) of the corresponding episode. Bottom row: External stimuli. Chosen stimuli are highlighted by the yellow dot at the top. Trial and episode numbers are given at the bottom.}
  \label{fig:withRewardFullSys1}
\end{figure}

\FloatBarrier

\section{Noise sources}\label{sec:noiseSources}

Noise is required in our model to consider different perturbative mechanisms. It appears at two levels in the model: in the basic module and in the regulatory module. The former models the distortion effects due to the extracellular media. The latter models the exploratory behavior of the subject which arises from curiosity, hesitation and/or perceptual difficulties. 

We describe the noise in terms of Ornstein-Uhlenbeck (OU) processes. An OU process can be sampled directly from a Gaussian distribution once the initial condition of the process is chosen properly. This is similar to the previous framework \citep{di2019biologically}, however with one difference: we choose the initial condition of the OU process as a zero mean Gaussian, where its variance is scaled by the convergence rate of the OU process as given below. This allows to avoid explicit simulation of the OU process, reducing the computational load.

The noise is a stochastic process evolving in time, independently of the rest of the state variables of the model. For a white Gaussian noise, this evolution can be written as an OU process of the following type:
\begin{equation}\label{eq:OUprocess}
\begin{split}
d\omega_\alpha (t) = & -\theta_\alpha \omega_\alpha(t) dt + \sigma dW_\alpha(t),\\
\omega_\alpha(0) = & \Omega, 
\end{split}
\end{equation}
where $W_\alpha (t)$ is a standard Brownian motion, $\theta_\alpha$ is a positive constant associated to the convergence rate of the process to its long term mean, and $\Omega$ is the initial condition, which is a centered Gaussian noise with variance $\sigma_0^2>0$. This is a linear stochastic differential equation and its unique solution \citep{arnold1974stochastic}  is
\begin{equation}
\omega_\alpha (t) = \operatorname{e}^{-\theta_\alpha t}\Omega + \sigma \int_0^t \operatorname{e}^{-\alpha(t-s)}dW_\alpha(s),
\end{equation}
where the solution $\omega_\alpha(t)$ is a Gaussian process. It can be written for a small time step $0<h\ll 1$ as
\begin{equation}
\omega_\alpha(t+h) = \operatorname{e}^{-\theta_\alpha h } \omega_\alpha(t) + \xi(t),
\end{equation}
where $\xi(t)$ is a centered Gaussian white noise with variance $\sigma_\xi^2 = \frac{\sigma^2}{2\theta_\alpha}(1-\operatorname{e}^{-2\theta_\alpha h})$, and which is sampled independently for each $t$ and subpopulation $\alpha$. We observe that $\omega_\alpha(t+h)$ is equivalent to a zero mean Gaussian with variance
\begin{equation}
\sigma_{\omega_\alpha}^2(t+h) =\operatorname{e}^{-2\theta_\alpha h}\sigma_{\omega_\alpha}(t)^2+ \sigma_\xi^2. 
\end{equation}
Once we choose $\sigma_{\omega_\alpha}^2(0) = \sigma_0^2= \frac{\sigma^2}{2\theta_\alpha}$, we observe that $\sigma^2_{\omega_\alpha}(t+h)$ is time independent and it is equal to $\frac{\sigma^2}{2\theta_\alpha}$. This allows us to generate $\omega_{\alpha}(t)$ from the Gaussian distribution $\mathcal{N}(0, \frac{\sigma^2}{2\theta_\alpha})$ independently at each instant $t>0$, avoiding explicit forward time simulations of the OU process given in \eqref{eq:OUprocess}. This is the idea behind introducing the noise terms $\omega_\alpha$ in the mean-field system \eqref{eq:mfEqns} and the noise term $\zeta_i$ appearing in the regulatory mechanism \eqref{eq:regulatory_Cauchy} as Gaussian white noise sampled directly from a Gaussian distribution at each time and for each subpopulation, but not as an explicit OU process evolving in time.


\subsection{Extracellular media distortion effects}

In the previous AdEx mean-field model \citep{di2019biologically}, the background noise was introduced as an additive noise to the base drive potential $s_{\text{AI}}$. This noise was evolving as an OU process and it modeled dynamically the distortion effects appearing in the firing rates due to the extracellular medium and synaptic perturbations. In our model, we introduce the white Gaussian noise $\omega_\alpha$, explicitly in the firing rate equations as shown in~\eqref{eq:mfEqns}. In this way, we avoid that the noise undergoes the nonlinear effects of the transfer functions since now it is additive in the model equations. This noise models the distortion effects of the extracellular media on the neural dynamics.

\subsection{Exploratory behavior}
One of the typical behaviors in the decision-making task is that the participant makes decisions which do not comply with the preset strategy even after the participant learned the strategy. This behavior occurs due to two reasons: (i) perceptual difficulty in the human experiment, i.e., the difference between stimuli is small, thus the participant makes the wrong decision as a result of perceptual difficulty; (ii) the participant hesitates or is willing to explore whatsoever related to the experiment setup, and this might happen in both human and macaque experiments.

The perceptual difficulty is due to the stimuli, and it does not require any additional mechanism to be represented in the model. On the other side, the exploratory will of the participant is a part of the cognitive framework. We model it via the Gaussian white noise $\zeta_n$ introduced in the regulatory mechanism \eqref{eq:regulatory_Cauchy}. This noise is sampled independently at each time instant. The noise level $\sigma$ is scaled by $(c_0 t^2)$, where $c_0>0$. Here $t$ denotes the time instant and it is reinitialized at the beginning of each trial. This scaling term introduces a strong initial perturbation to the competition between the L2/3 populations. This perturbation decays in time. This decay is needed, since, otherwise the noise could dominate the whole competition instead of perturbing only the initial bias. The parameter $c_0$ determines how strong the initial perturbation is. The regulatory mechanism therefore, is not only for the online reward-driven learning of the preset strategy but also for providing a natural behavior which is open to making wrong decisions. This is similar to what was observed in both human and macaque experiments. This exploratory behavior appears as irregular and rather sparse in-cluster deviations in the performance index plots; see Figure~\ref{fig:globalMeasures}.

\section{Quantification of behavior}\label{sec:quantification}

Quantification of behavior is required to analyze the results and to make a comparison between the simulation and experiment results. For a quantification of the results, we need adequate metrics. These metrics should provide a global characterization of the learning phase and of the erroneous decisions which the subjects and the model make.
 
Subject behavior in the experiments was quantified in terms of the coherence between the choices of the subject and the preset strategy. We follow the same approach to quantify the model behavior in the simulations. This coherence is measured by using a metric called performance index, which is defined in~\eqref{eq:scoreMeasure}. 

In each episode, the maximum performance index refers to the case in which the subject makes the choice in coherence with the preset strategy in every trial of the episode. The maximum performance index value is 1. The minimum performance index refers to the case in which the participant makes the incoherent choice in every trial of the episode. The minimum performance index is 0. All other cases with partial coherence are scaled between 0 and 1; see~\eqref{eq:scoreMeasure}.

For the quantification of performance indexes, we use two objects: initial cluster episode and in-cluster deviation (Figure~\ref{fig:globalMeasures}). These objects characterize fully the behavioral pattern of a subject. The behavioral pattern is represented in terms of the performance indexes plotted with respect to their corresponding episodes; see Figure~\ref{fig:globalMeasures}. In the performance index plots, initial cluster episode gives a measure of how quickly the subject captured the preset strategy. It characterizes the learning phase. In-cluster deviation gives a measure of erroneous decisions that the subject made after learning the preset strategy. These erroneous decisions can be due to several reasons, e.g., perceptual difficulties and curiosity to explore. Therefore, in-cluster deviations characterize the behavior after the learning was completed. We use these two objects for quantification of behavior. However, we need first, the following three definitions to be able to describe initial cluster episode and in-cluster deviation:

\medskip

\noindent\textbf{Definition 1} ({\textit{Performance deviation}): A performance deviation is a point with a performance index lower than the maximum performance index ($=1$), except for the first episode. 

\medskip
\noindent\textbf{Definition 2} (\textit{Deviation cluster}): A deviation cluster is a set of at least 3 successive performance deviations with respect to the episode numbers. 

\medskip
\noindent\textbf{Definition 3} (\textit{Performance cluster}): Performance cluster is the set of performance indexes in which there is no deviation cluster and whose last performance index is also the last index of the whole experiment.

\medskip

\noindent Now we can define initial cluster episode and in-cluster deviation as follows:

\medskip
\noindent\textbf{Definition 4} (\textit{Initial cluster episode}): The first episode number of a performance cluster.


\medskip
\noindent\textbf{Definition 5} (\textit{In-cluster deviation}): A performance deviation is called \textit{in-cluster deviation} if it occurs in a performance cluster. It is called \textit{out-cluster} otherwise.  

\medskip

A performance cluster starts at the episode where the participant begins to make decisions in complete coherence with the preset strategy. Within the cluster, the participant makes coherent decisions almost in each episode until the end of the cluster. Almost; because the participant might make decisions in the aforementioned exploratory manner, producing in-cluster deviations. This does not break the performance cluster as long as in-cluster deviations do not constitute a deviation cluster. We refer to Appendix for a characterization of the model behavior in terms of performance indexes and reaction times; see Figures~\ref{fig:FigPerformanceSimulation} and~\ref{fig:RT_H1_T1_T2}.

%In particular, performance deviations arising from the exploratory behavior and perceptual difficulties are momentary, they are locally concentrated around certain episodes appearing after the participant learned the preset strategy. Therefore, the quantification of the behavioral performance requires global metrics which are robust to such local deviations and which can quantify these deviations at the same time. 


\section{Model predictions}\label{sec:results}

\subsection{Human performance}\label{sec:H1_humanPerformance}

The results of the simulations and experiments of the human case are quantified based on the mean and standard deviation of the initial cluster episodes (Definition 4), as well as the mean and standard deviation of the in-cluster deviations (Definition 5). This quantification fully describes the learning pattern of the model, as well as the behavior of the model after it captures the preset strategy. We obtain the mean values and standard deviations for each parameter set from 10 realizations of the simulation with the corresponding parameter set. Each realization is composed of 100 episodes.

We vary two parameters to change the performance behavior of the model: $k$ in the reward-tracking module given by~\eqref{eq:rewardMechanismEqn} and $c_0$ in the regulatory module given by~\eqref{eq:regulatory_Cauchy}. The former is the learning speed and it determines how quickly the model learns the preset strategy, thus the initial cluster episode. The latter is the flexibility parameter. It determines how much the model is flexible to make mistakes, even after learning the preset strategy. These mistakes model the erroneous choices arising from perceptual difficulties, hesitation and curiosity/exploratory behavior of the subject.

We present the results of Horizon 1 simulations in Figure~\ref{fig:statisticalH1}. We observe that as $k$ or $c_0$ increases, the initial episode number of performance clusters decreases as shown on the left column. This is the case also in Horizon 0 as seen in Figure~\ref{fig:statisticalH0}. However, differently from Horizon 0, the decreasing curves in Horizon 1 are concave, meaning that the decrease rate of the mean initial episode number of performance clusters increases. On the contrary, we observe an increasing pattern in the mean of in-cluster deviations as $k$ or $c_0$ increases. This can be due to that, as the initial episode of the performance cluster decreases, there are more episodes in which the model makes erroneous choices. This generates in-cluster deviations. Therefore, it is likely that there will be more in-cluster deviations. An interesting observation is that this is not the case in Horizon 0 simulations. We predict that it is due to the difference in the task complexity. Since the task complexity is low in Horizon 0, the model makes fewer erroneous choices in the performance clusters once it learns the preset strategy. In all Horizon 1 simulations, we observe that the model can produce close results to the experimental measurements once $k$ and $c_0$ are chosen properly, for example $k=0.05$ and $c_0=2$. 

The results of Horizon 0 simulations are shown in Figure~\ref{fig:statisticalH0}. We observe in the left column that the learning speed decreases both with increasing $k$ and with increasing $c_0$, resulting in smaller values of the initial episode number of the performance clusters. The parameter $k$ directly controls the learning speed, and $c_0$ contributes to it by avoiding the deviation blocks which might break the performance clusters. In both cases, the decrease in the mean value profiles is convex. In all Horizon 0 simulations, we see that the model can reproduce close results once $k$ and $c_0$ are chosen properly, for example $k=0.3$ and $c_0=1.1$.

\begin{figure}[!htbp]
\begin{minipage}[c]{\textwidth}
\sidesubfloat[(a)]{ \hspace{0.5cm} \includegraphics[clip,scale=0.9]{statisticalH1}\label{fig:statisticalH1}
}

\bigskip

\sidesubfloat[(b)]{ \hspace{0.5cm} \includegraphics[clip,scale=0.9]{statisticalH0}\label{fig:statisticalH0}
}

\end{minipage}
\caption{Simulation metrics with respect to the varied learning speed $k$ and flexibility parameter $c_0$. (a) Horizon 1 human case. On the top and bottom rows $k$ and $c_0$ are varied, respectively. In the top row, $c_0=2$. In the bottom row, $k=0.05$. The vertical blue lines show the standard deviations of the simulation results. The red horizontal lines indicate the initial cluster episode number (left column) and the number of in-cluster deviations (right column) obtained from the human experiment. (b) Horizon 0 human case. In the top row, $c_0=1.1$ and $k$ is varied. In the bottom row, $k=0.3$ and $c_0$ is varied. The blue vertical lines show the standard deviations of the corresponding statistical sample. The red horizontal lines indicate the experimental values as in (a). In both (a) and (b), the means and standard deviations are obtained from 10 realizations of the same simulation setup for each $(k, c_0)$ pair. The parameters $k$ and $c_0$ can be seen as rescaling constants, therefore they are unitless. See~\eqref{eq:parametersIn}-\eqref{eq:parametersFin} for the rest of the parameters used in these simulations.}
\label{fig:statisticalPerformance}
\end{figure}


%\begin{figure}[!htbp] 
%\noindent\makebox[\textwidth]{
%  \includegraphics{statisticalH1}
%  }
%  \caption{Horizon 1 human case. Simulation statistics with respect to the varied learning speed $k$ in the top row, and with respect to the varied flexibility parameter $c_0$ in the bottom row. In the top row, $c_0=2$. In the bottom row, $k=0.05$. The vertical blue lines show the standard deviations of the simulation results. The red horizontal lines indicate the initial cluster episode number and the number of in-cluster deviations obtained from the human experiment. The mean and standard deviation are obtained from 10 realizations of the same simulation setup for each $(k, c_0)$ pair. The parameters $k$ and $c_0$ can be seen as rescaling constants, therefore they are unitless. See Supporting information for the rest of the parameters.}
%\label{fig:statisticalH1}
%\end{figure}
%
%
%
%\begin{figure}[!htbp]
%\noindent\makebox[\textwidth]{
%  \includegraphics{statisticalH0}
%  }
%  \caption{Horizon 0 human case. Simulation statistics with respect to the varied learning speed $k$ in the top row, and with respect to the varied flexibility parameter $c_0$ in the bottom row. In the top row, $c_0=1.1$. In the bottom row, $k=0.3$. The blue vertical lines show the standard deviations of the corresponding statistical sample. The red dashed lines are the experimental values. The mean and standard deviations are obtained via repeating the simulation 10 times for each $(k, c_0)$ pair. See \eqref{eq:parametersIn}-\eqref{eq:parametersFin} for the rest of the parameters.}
%  \label{fig:statisticalH0}
%\end{figure}


\FloatBarrier


\subsection{Human reaction times}
\label{sec:HumanReactionTimes}

In the experiments, the reaction time in one trial is measured as the time duration between the instant when the stimuli are shown simultaneously on the monitor, and the moment when the participant starts to move the pointer. In the simulations, we measure the reaction time as the time duration between the instant when the two stimuli are provided to the model and the time instant at which the decision is made, i.e., when the difference between the excitatory subpopulation firing rates $v_{e_A}$ and $v_{e_B}$ exceeds the decision threshold, which is fixed to 5 Hz.


\begin{figure}[!htbp]
\begin{minipage}[c]{\textwidth}
\sidesubfloat[(a)]{ \hspace{0.5cm} \includegraphics[clip,scale=1]{H1_RT_Histogram}\label{fig:H1_RT_Histogram}
}

\bigskip

\sidesubfloat[(b)]{ \hspace{0.5cm} \includegraphics[clip,scale=1]{H0_RT_Histogram}\label{fig:H0_RT_Histogram}
}

\end{minipage}
\caption{Reaction time histograms obtained from the human experiment and corresponding simulations. (a) Horizon 1. The red curves denote the fitted distributions via Python. Trial 1 and 2 histograms are in the top and bottom rows, respectively. The simulations are performed with $k=0.05$, $c_0=2$ and $\text{decision threshold}=5$ Hz. (b) Horizon 0. The simulations were performed with $k=0.3$, $c_0=1.1$ and $\text{decision threshold}=5$ Hz. See~\eqref{eq:parametersIn}-\eqref{eq:parametersFin} for the rest of the parameters used in the simulations.}
\label{fig:RT_Histograms}
\end{figure}

In Figure~\ref{fig:H1_RT_Histogram}, we show the simulation and experiment histograms of the reaction times corresponding to Horizon 1, together with the fitted distributions for the best performance-providing $k$ and $c_0$ values among the tested ones. In each case, we fit the distributions of different types to both simulation and experiment histograms, and choose the distributions which give the smallest squared sum errors. We use the \textit{Fitter} function from the Python \textit{Fitter} package. We consider the histograms of the reaction times obtained from the first and second trials separately. In Trial 1, skewnorm distribution and hypergeometric distribution are fitted to the experiment and simulation histograms, respectively. Corresponding squared sum error values are 1.3e-5 and 9e-6 for the experiment and simulation histograms, respectively. Pearson correlation coefficient of these fitted distributions is 0.92. In Trial 2 histograms, skewnorm distribution and hypergeometric distribution are fitted to the experiment and simulation histograms, respectively. Corresponding squared sum error values are 1.7e-5 and 1.1e-5 for the experiment and simulation histograms, respectively. Pearson correlation coefficient of these fitted distributions is 0.63. Trial 2 has a weaker overlap between the experiment and simulation histograms compared to Trial 1.

In Figure~\ref{fig:H0_RT_Histogram}, we show the simulation and experiment histograms of the reaction times corresponding to Horizon 0 in the same fashion as the Horizon 1 results. The best fit is achieved with skewnorm distribution and hypergeometric distribution for the experiment and simulation histograms, respectively. Squared sum errors are 13e-6 and 12e-6 for the experiment and simulation histograms, respectively. Pearson correlation coefficient between the fitted distributions is 0.91.


The difference between the experimental and simulation reaction times could be due to several factors. In the experiments, reaction time includes the time between the instant when the participant makes the decision of which stimulus to choose and the instant when the participant starts to move the pointer towards the chosen stimulus. This adds a duration which may vary, in particular due to hesitation, fatigue, loss of motivation etc. The time duration between these two instants cannot be taken into account in the model, simply because there is no motor action in the model. Another reason could be the initial conditions. We do not have any access to the initial conditions of the neural dynamics of the participant. Therefore, we initialize the basic module from the same initial conditions at the beginning of each trial in our simulations. Nevertheless, it is likely that these conditions are different at the beginning of each trial in the experiments. These factors do not have strong effects on performance results, however they can change the results regarding the reaction times significantly.
 


\FloatBarrier


\subsection{Macaque performance}

We observe that the trends in the macaque performance indexes are similar to the ones found in the Horizon 1 human task simulations. We see that the simulation results overlap with the macaque experiments for properly chosen parameters, for example $k=0.12$, $c_0=0.106$; see Figure~\ref{fig:H1_RT_Histogram}.

\begin{figure}[!htbp] 
\noindent\makebox[\textwidth]{
  \includegraphics[clip, scale=1]{statisticalH1Monkey}
  }
  \caption{Simulation metrics of the macaque case. In the top row, $c_0=0.106$, and $k$ is varied. In the bottom row, $k=0.12$, and $c_0$ is varied. The vertical blue lines show the standard deviation of the corresponding statistical sample. The red dashed lines show the value obtained from the macaque experiment. The mean and standard deviations were obtained from 10 realizations of the same simulation setup for each $(k, c_0)$ pair. The decision threshold is 5 Hz. See~\eqref{eq:parametersIn}-\eqref{eq:parametersFin} for the rest of the parameters used in the simulations.}
  \label{fig:statisticalH1Monkey}
\end{figure}


\FloatBarrier


\subsection{Macaque reaction times}


\begin{figure}[!htbp]
\noindent\makebox[\textwidth]{
  \includegraphics[scale=1]{H1_Monkey_RT_Histogram}
  }
  \caption{Reaction time histograms of the macaque case. The red curves denote the fitted distributions via Python. Trial 1 and 2 histograms are in the top and bottom rows, respectively. The simulations are performed with $k=0.12$, $c_0=0.106$ and $\text{decision threshold}=5$. See~\eqref{eq:parametersIn}-\eqref{eq:parametersFin} for the rest of the parameters used in the simulations.}
  \label{fig:H1_Monkey_RT_Histogram}
\end{figure}

% Similarly to the simulations of the human experiment, as the decision threshold increases, the competition between $v_{e_A}$ and $v_{e_B}$ lasts longer, thus the reaction times increase as shown in Figure~\ref{fig:RT_Monkey_H1_T1_T2} separately for Trials 1 and 2. Those results support that the reaction time is determined rather by the decision threshold than by $k$ and $c_0$.


In Figure~\ref{fig:H1_Monkey_RT_Histogram}, we show the simulation and experiment histograms of the reaction times corresponding to the macaque experiment, together with the fitted distributions. Here $k$ and $c_0$ are fixed to the values providing the best performance overlap. We fit the distributions to both simulation and experiment histograms by using the \textit{Fitter} function from the Python \textit{Fitter} package as in the case of the human experiment. In Trial 1, genlogistic distribution and powernorm distribution are fitted to the experiment and simulation histograms, respectively. Corresponding squared sum error values are 1e-5 and 9e-6 for the experiment and simulation histograms, respectively. Pearson correlation coefficient of these fitted distributions is 0.97. In Trial 2 histograms, burr distribution and kappa3 distribution are fitted to the experiment and simulation histograms, respectively. Corresponding squared sum error values are 9.9e-5 and 5.73e-4 for the experiment and simulation histograms, respectively. Pearson correlation coefficient of these fitted distributions is 0.31. Trial 2 has a weaker overlap between the experiment and simulation histograms compared to Trial 1 as in the case of the human task.

The same factors regarding the reaction times and the initial conditions as in the human case are valid also in the macaque case. We predict that the difference between the experimental and simulation results regarding the reaction times arises from these factors; see Section~\ref{sec:HumanReactionTimes}.

\FloatBarrier


\section{Discussion}\label{sec:discussion}

We presented a biologically plausible AdEx mean-field model based on two interacting L2/3 neural populations. The model is composed of three modules: basic, regulatory and reward-tracking modules. The basic module is based on the AdEx mean-field equations which we extended from one population~\citep{di2019biologically} to two populations. Each population is composed of a pair of excitatory-inhibitory subpopulations. The basic module generates the firing rates of the subpopulations. The excitatory subpopulations are in competition with each other to make a decision. The winning subpopulation makes the decision. The regulatory module introduces a bias into this competition such that one of the subpopulations becomes more likely to win the competition. Therefore, it makes one of the two stimuli more likely to be chosen. The bias depends on the increase or decrease in the reward amounts. The reward-tracking module keeps the track of these increases and decreases in the reward amounts. It transmits the information regarding these increases and decreases between successive episodes. We tested this model on two complex behavioral decision-making tasks, which were performed on human and macaque. 

The model is biologically plausible since its connectivity architecture overlaps with an anatomically coherent case and the neural transfer functions are obtained based on \textit{in vitro} measurements as well as neural responses obtained from the AdEx neuronal network. The novelties of the model are at both connectivity and structure levels.

Regarding the connectivity, the L2/3 populations are assumed to communicate via excitatory connections, as observed in macaque~\citep{hirata2008functional}. Hence, the connections between the L2/3 populations are only excitatory. Two such connections originate from the excitatory subpopulation of one of the L2/3 populations. The first one of these connections is onto the excitatory subpopulation of the other L2/3 population. The second one is onto the inhibitory subpopulation of the other L2/3 population. 

These connections overlap with the long range cortico-cortical connections found in the prefrontal cortex (PFC). The long-range cortico-cortical connections can be found between the lateral PFC and the limbic areas related to dopamine neurons, such as the medial PFC. The former was suggested to be responsible for the choices related to long-term planning independent of the delay of the reward, and the latter was considered for the choices related to impulsive decisions with short-term rewards~\citep{mcclure2004}. In our model, inhibitory control over the excitatory subpopulation is triggered by the increasing activity of the excitatory subpopulation of the other L2/3 population. This is not the case in the previous models \citep{wang2002probabilistic, marcos2013neural}. In \citep{wang2002probabilistic}, there is only one inhibitory subpopulation which is connected to all excitatory subpopulations via the long range connections. In \citep{marcos2013neural}, the excitatory and inhibitory subpopulations are not explicit. They are intrinsically represented as one neural population. Two such populations are connected to each other via inhibitory connections, and recurrently connected to themselves via excitatory connections.  


%These connections are necessary to provoke the bicolumnar competition in a biophysically realistic way. To achieve it, we extended rigorously the AdEx mean-field model which was proposed in \citep{di2019biologically} for an AdEx network of an isolated single column with adaptation. This extension provides the bicolumnar architecture in which the columns are connected to each other via the long range cortico-cortical connections. This allows the model to provoke the bicolumnar competition. The model chooses one of the two alternatives in accordance with the choice represented by the column which wins the competition.

On the structure side, we model each L2/3 population as the mean-field limit of a pair of excitatory and inhibitory neural subpopulations. This allows us to provide the stimuli to both excitatory and inhibitory subpopulations, which is not the case in previous models~\citep{wang2002probabilistic,marcos2013neural, cecchini2024cognitive}. As a result, our model can be extended to also decision-making with multiple choices. Moreover, the parameters are directly linked to the biophysical mechanisms of RS and FS cells. Therefore, our model can be applied to not only behavioral data but also to relevant neurophysiological data, in particular to multi-site array recordings of the macaque cortex. In this way, the model has the potential to link the phenomenological decision-making models \citep{wang2002probabilistic, marcos2013neural, cecchini2024cognitive} to the neural dynamics observed at mesoscopic level.

Moreover, we adapted the regulatory module given in~\citep{cecchini2024cognitive} to the AdEx system to introduce the plasticity which learns the preset strategy in the task. This machinery is essential for the model to find the optimal decision-making strategy. The regulatory module sustains the optimal strategy once this strategy is learned. This is due to the fact that once the introduced bias provides increase in the rewards, the regulatory module reinforces and retains the bias. This can happen successively over episodes, relating the regulatory module to working memory at phenomenological level.


%Finally, the AdEx mean-field framework was never used for a cognitive task. We show here that it can reproduce the output of the whole-brain scale activity represented on a behavioral scale, and in the context of the considered decision-making tasks. This supports the idea of that the AdEx mean-field framework can be embedded in a whole-brain simulator such as The Virtual Brain (TVB) \citep{sanz2013virtual, goldman2020brain} and can reproduce the whole-brain dynamics induced by cognitive tasks, in particular by decision-making tasks. Some preliminary results in this direction can be found in~\citep{turan2023whole}. 


The model was tested at the behavioral level by comparing its predictions to the experimental data obtained from the human and macaque participants. The comparison was made based on three statistical metrics: mean value of initial episode numbers of the performance clusters, mean value of in-cluster performance deviations and reaction time distributions. It was found that the model reproduces several characteristics of the experiment results. To begin with, the model predicts closely the performance indexes in the cases of both human and macaque for properly chosen parameter sets. It is possible to produce a good variety of behavioral patterns by varying the learning speed and the flexibility parameter. In reaction times, the model and experiment results have a mismatch, in particular in Trial 2 measurements of Horizon 1 experiments in both human and macaque. A possible reason is that the motor action is included in reaction time measurements in the experiments, whereas it is not possible to consider the motor action in the model setup. Another possible reason is that the decision threshold in the model is fixed. It is not dynamically adapted over the episodes as the strategy is learned.

Our work is linked to~\citep{di2019biologically, cecchini2024cognitive}. It uses the experimental data obtained from human and macaque subjects to provide the metrics based on performance indexes and reaction times. The data was given in~\citep{humanEbrains, macaqueEbrains}. Our work builds upon~\citep{di2019biologically} by extending the AdEx mean-field equations from a single pair, to two pairs of excitatory-inhibitory subpopulations. The extension relies on the excitatory connections between the L2/3 populations. Moreover, we integrate the regulatory and reward-tracking modules with this extended AdEx framework. These modules introduce a learning mechanism to the model, which was not considered previously in~\citep{di2019biologically}. Our work builds upon~\citep{cecchini2024cognitive} in terms of the basic module and the reward-tracking module. The basic module in our framework elaborates several biophysical details as a result of the use of  AdEx equations. These details were not explicit in~\citep{cecchini2024cognitive}. Our model considers both human and macaque cases, with the two modes (\eqref{eq:rewardMechanismEqn} and~\eqref{eq:macaqueRewardTracking}) in the reward-tracking module.


Our model was designed for the optimal decision-making. Yet, it has the potential to be extended to the sub-optimal decision-making. Optimal decision-making refers to making choices which provide the maximum possible reward \citep{doya2007bayesian}. Albeit the absence of a precise definition, sub-optimal decision-making refers to the decision-making which deviates from the optimal decision-making in terms of performance indexes. The model already reproduces partially sub-optimal behavior in the performance clusters thanks to its exploratory behavior as shown, for example, in Figure~\ref{fig:c0EffectV}. This property can be improved: in the regulatory module, we can introduce a term which weights the flexibility parameter based on the changes in the reward amounts or in the motivation of the participant to conduct the task. This can be combined with the decision thresholds which are dynamically adapted throughout the episodes. This could project the changes in the motivation of the participant, on the performance indexes. This modification could provide a better understanding of the neural dynamics which modulate the behavioral strategies of the participant in accordance with the increase or decrease in the motivation. This has been an active research area in the experimental side \citep{leotti2010motivational, giamundo2021neuronal, padmala2010interactions}, and it can benefit enormously from the computational side.


Another interesting future research direction could be related to the brain states. The AdEx mean-field framework can simulate, in addition to the AI state, several other brain states such as sleeping, drowsiness or under anesthesia states~\citep{di2019biologically, goldman2020brain}. This property of the AdEx framework can be integrated into decision-making context as well. This can be interesting to study for example the effect of different levels of drowsiness on the capacity to perform cognitive tasks, see for example Figure~\ref{fig:awakeToSleep}. The figure shows the gradual deterioration of our model's performance in a simulation of Horizon 1 decision-making task. In the simulation, our model goes through a transition from the awake state towards under anesthesia state. Studies in this direction might provide computational support for better understanding the effects of anesthesia or drugs on cognitive abilities.

\begin{figure}[!htbp] 
\noindent\makebox[\textwidth]{
  \includegraphics{awakeToSleep}
  }
\caption{Performance index plots of the simulations of the model as the model goes through the transition from the awake state towards the sleeping state. The spike-frequency adaptation of the RS cells becomes stronger as we increase the parameter $b$ found in the adaptation term of~\eqref{eq:mfEqns}. This pushes the model towards the sleeping state. We observe that the model loses gradually the capacity to obtain maximum performance indexes, i.e., it loses gradually the ability to learn the preset strategy.}
\label{fig:awakeToSleep}
\end{figure}


Finally, a further extension of the model can be towards the large-scale brain dynamics of decision-making. For this, our AdEx mean-field framework can be integrated in The Virtual Brain~\citep{sanz2013virtual, goldman2020brain} to model the decision-making brain dynamics of the PFC, which has been traditionally considered as the key area involved in decision-making \citep{bechara2000characterization, hampton2006role, rushworth2011frontal, shadlen2013decision, domenech2015executive, funahashi2017prefrontal}. In this way, the perceptual inputs can be provided directly from the visual (or other perceptual) areas to the PFC. Consequently, the outputs of our model can be provided as the outputs of the PFC to the motor areas. Some preliminary results in this direction can be found in~\citep{turan2023whole}.

%Among these areas, the prefrontal cortex (PFC) has been traditionally considered as the key area involved in decision-making \citep{bechara2000characterization, hampton2006role, rushworth2011frontal, shadlen2013decision, domenech2015executive, funahashi2017prefrontal}. Our model focuses on the PFC, it does not take into account the subcortical regions. It considers one of the simplest and sufficient scenarios generating a selection: Two excitatory neural populations compete and the winning population determines which alternative to select~\citep{desimone1995neural, reynolds2004attentional, moore2017neural}. Therefore, its embedding to TVB must be considered at the cortical level.


%Indeed, decision-making is a complex task involving several cortical and subcortical regions~\citep{paulus2003increased, sanfey2006neuroeconomics, guo2013brain, broche2016neural}. Among these areas, the prefrontal cortex (PFC) has been traditionally considered as the key area involved in decision-making \citep{bechara2000characterization, hampton2006role, rushworth2011frontal, shadlen2013decision, domenech2015executive, funahashi2017prefrontal}. Our model focuses on the PFC, it does not take into account the subcortical regions. It considers one of the simplest and sufficient scenarios generating a selection: Two excitatory neural populations compete and the winning population determines which alternative to select~\citep{desimone1995neural, reynolds2004attentional, moore2017neural}. Therefore, its embedding to TVB must be considered at the cortical level.


\section*{Appendix}
\setcounter{section}{0}
\renewcommand*\thesubsection{\arabic{subsection}}

\subsection{Neural transfer functions}


Neural transfer functions of AdEx mean-field equations were provided in~\citep{zerlaut2018modeling} and generalized to also spike-frequency adaptation in~\citep{di2019biologically}. These functions were obtained by a semi-analytical fitting to the averaged output of the AdEx neuronal network. The output of the network was induced by random spike bombardments generated from a Poisson distribution. We use the same transfer functions provided in~\citep{di2019biologically}. We summarize here how their explicit formulas were obtained. For details, we refer to~\citep{zerlaut2018modeling, di2019biologically}.

\subsubsection*{AdEx neuronal network}

The AdEx mean-field of a pair of excitatory-inhibitory subpopulations with spike-frequency adaptation approximates the average dynamics of the AdEx neuronal network. The network is composed of a certain number of neurons. Each neuron in the network is described by the AdEx integrate-and-fire model~\citep{brette2005adaptive}:
\begin{equation}\label{eq:AdExNetwork}
\begin{split}
C_m\frac{dV_\ell}{dt} & = g_L\,(E_L-V_\ell)+\Delta \operatorname{e}^{\frac{V_\ell-V_{\text{thr}}}{\Delta}}-w_\ell +I_\text{syn},\\
\frac{dw_\ell}{dt} & = -\frac{w_\ell}{\tau_w} + b\sum_{t_s\in\{t_\text{spike}(\ell) \}} \delta_{t-t_s}+a\,(V_\ell-E_L),
\end{split}
\end{equation}
where $C_m=200$ pF denotes the membrane capacity and $V_\ell$ is the membrane voltage of a neuron $\ell$ in the network. The voltage $V_\ell$ is reset to the resting voltage $-65$ mV at $t_s\in \{t_\text{spike}(\ell) \}$, which denote all the instants when the neuron generates a spike, i.e, when $V_\ell>V_{\text{thr}}=-50$ mV. Here $g_L=10$ nS and $E_L= -65$ mV are the conductance and the leakage reversal of the leak term. Here $\Delta$ denotes the weight of the exponential term. It is 2 mV for RS cells and 5 mV for FS cells. Note that FS cells have no adaptation ($w_\ell=0$). Therefore, $a=b=0$ for all inhibitory neurons. For RS cells, the adaptation is introduced with $a=4$ and $b=40$ in our simulations. Synaptic input to the neuron $\ell$ is defined as follows:
\begin{equation}
I_{\text{syn}}=(E_L^e-V_\ell)G_e + (E_L^i-V_\ell)G_i,
\end{equation}
with $E_L^e = 0$ mV and $E_L^i=-80$ mV denoting the reversal potential of the excitatory and inhibitory presynaptic cells, respectively. The conductances are defined as
\begin{equation}\label{eq:conductances}
G_e(t)= Q_e \sum_{t_s^e\in\{t_\text{spike}^e(\ell) \}}\mathcal{H}(t-t_s^e)\operatorname{e}^{-\frac{t-t_s^e}{\tau_e}},\quad\quad G_i(t)= Q_i \sum_{t_s^i\in\{t_\text{spike}^i(\ell) \}}\mathcal{H}(t-t_s^i)\operatorname{e}^{-\frac{t-t_s^i}{\tau_i}},
\end{equation}
where $\mathcal{H}$ denotes the Heaviside function and $\tau_e=\tau_i=5$ ms are the decay rates of the excitatory and inhibitory presynaptic neurons. Here $t_s^e$ and $t_s^i$ are the instants when the excitatory and inhibitory presynaptic neurons trigger spikes, respectively. Finally, $Q_e=1.5$ nS and $Q_i=5$ nS are the excitatory and inhibitory quantal conductances, respectively.

\subsubsection*{Transfer functions}

The classical AdEx mean-field equations were derived in~\citep{di2019biologically} from the AdEx network of neurons described by~\eqref{eq:AdExNetwork}. The derivation of the mean-field equations was based on a master equation formalism which was provided in~\citep{el2009master}. The mean-field equations were for a single pair of excitatory-inhibitory subpopulations, with a spike-frequency adaptation. This is equivalent to decoupling the L2/3 populations of the basic module~\eqref{eq:mfEqns} of our model and considering only one of these decoupled populations. In that case, each L2/3 population reduces to a system of 6 state variables as in the classical mean-field system. Therefore, for the neural subpopulations in our model, we use the same transfer functions given for the classical AdEx mean-field system in~\citep{di2019biologically}. The difference is that, the inputs to the  transfer functions have not only the local components, but also the components related to the interactions between the L2/3 populations in our extended AdEx mean-field framework.

In the following, we will use the notation $e,i$ to denote the excitatory and inhibitory subpopulations in either one of the decoupled L2/3 populations. The equations of the transfer functions are identical, independently of which L2/3 population we consider. These neural transfer functions were obtained via the semi-analytical derivation explained in~\citep{zerlaut2018modeling, di2019biologically}. The idea was based on the spike bombardment of a single neuron described by~\eqref{eq:AdExNetwork}. The spikes were generated with various excitatory and inhibitory firing rates $v_e$ and $v_i$ which follow the Poisson statistics. This allowed to obtain the means $\mu_{G_e}, \mu_{G_i}$  and the standard deviations $\sigma_{G_e}, \sigma_{G_i}$ of the excitatory and inhibitory conductances in~\eqref{eq:conductances}. These functions were found via Campbell-Hardy theorem~\citep{papoulis2002Probability} as follows:
\begin{equation}
\begin{split}
\mu_{G_e}(v_e, v_i) & = v_e\, K_e\, \tau_e\, Q_e,\quad\quad \sigma_{G_e}(v_e, v_i)=Q_e\,\sqrt{\frac{v_e\, K_e\,\tau_e}{2}},\\
\mu_{G_i}(v_e, v_i) & = v_i\, K_i\, \tau_i\, Q_i,\quad\quad\;\, \sigma_{G_i}(v_e, v_i)=Q_i\,\sqrt{\frac{v_i\, K_i\, \tau_i}{2}}.
\end{split}
\end{equation}
These functions control the input conductance $\mu_G$ and its effective membrane time constant $\tau_m$, which have the following expressions:
\begin{equation}
\mu_G(v_e, v_i)=\mu_{G_e} + \mu_{G_i} + g_L,\quad\quad \tau_m(v_e, v_i)=\frac{C_m}{\mu_{G_e}+\mu_{G_i}+g_L}.
\end{equation}
Consequently, mean membrane voltage of the neuron was obtained as:
\begin{equation}\label{eq:forTF1}
\mu_V(v_e, v_i)=\frac{\mu_{G_e}E^e_L+\mu_{G_i}E^i_L+g_L E_L-w}{\mu_G},
\end{equation}
where $w$ denotes the adaptation of the neuron. Finally, the standard deviation $\sigma_V$ and the autocorrelation time $\tau_V$ of the membrane voltage fluctations were found from the power density of the fluctutations:
\begin{equation}\label{eq:forTF2}
\begin{split}
\sigma_V (v_e, v_i)& =\sqrt{\sum_{\alpha\in\{e, i\}} K_\alpha v_\alpha \frac{(U_\alpha\,\tau_\alpha)^2}{2(\tau_m+\tau_\alpha)}},\\
\tau_V(v_e, v_i)& = \frac{\sum\limits_{\alpha\in \{e,i \}} K_\alpha v_\alpha(U_\alpha\,\tau_\alpha)^2}{\sum\limits_{\alpha\in \{e, i \}} K_\alpha v_\alpha (U_\alpha\,\tau_\alpha)^2/(\tau_m +\tau_\alpha)},
\end{split}
\end{equation}
where $U_\alpha = \frac{Q_\alpha}{\mu_G}(E_L^\alpha-\mu_V)$.

In~\eqref{eq:forTF2} and~\eqref{eq:forTF3}, we find the analytical expressions used in the transfer functions given by the formula~\eqref{eq:transferFunction}. These analytical experessions are complemented by the voltage-effective threshold $v_{\text{eff}}$, which was found by fitting the coefficients of the second order polynomial:
\begin{equation}\label{eq:forTF3}
v_{\text{eff}}(\mu_V, \sigma_V, \rho_V)= P_0 + \sum\limits_{x\in\{\mu_V, \sigma_V, \rho_V\}} P_x\,\frac{x-x^0}{\bar{x}^0} + \sum\limits_{x,y\in \{\mu_V, \sigma_V, \rho_V  \}} P_{xy} \Big (\frac{x-x^0}{\bar{x}^0} \Big ) \Big (\frac{y-y^0}{\bar{y}^0}   \Big ).
\end{equation}
The fitting was made according to the simulations of single neuron activity described by~\eqref{eq:AdExNetwork}. Here $\rho_V = \tau_V\frac{g_L}{C_m}$ and $\mu_V^0=-60$ mV, $\sigma_V^0=0.004$ mV, $\rho_V = 0.5$, $\bar{\mu}_V^0=0.001$ mV, $\bar{\sigma}^0_V=0.006$ mV, $\bar{\rho}_V^0 = 1$. We use the same fitted coefficients as given in~\citep{di2019biologically}. The fitting of the polynomial~\eqref{eq:forTF3} and the functions in~\eqref{eq:forTF1},~\eqref{eq:forTF2} provide the transfer functions given by~\eqref{eq:transferFunction}. 


\subsection{Simulation parameters}

The parameters which are used in Figures ~\ref{fig:statisticalPerformance}-\ref{fig:H1_Monkey_RT_Histogram} and Figures~\ref{fig:FigPerformanceSimulation}, \ref{fig:RT_H1_T1_T2} are as follows:

\begin{equation}\label{eq:parametersIn}
\begin{split}
T &= 5\, \text{ms}^{-1},\quad \sigma=0.01, \quad \tau_w = 5000\, \text{ms}^{-1}\; (\text{for RS}),\quad 1^{-9}\, \text{ms}^{-1}\;(\text{for FS}), \\
a & = 4\; (\text{for RS}),\quad 0 (\text{for FS}),\quad b= 40\; (\text{RS}),\; 0\; (\text{FS}),\quad E_L = -65\,\text{mV},
\end{split}
\end{equation}
and it is given for~\eqref{eq:inputTF_noRegulation} as
\begin{equation}
\begin{split}
s_{\text{AI}}= 5\, \text{Hz},\quad w_c=1.6.
\end{split}
\end{equation}
Finally, the parameters appearing in~\eqref{eq:regulatory_Cauchy} and~\eqref{eq:lambdaExpression} are
\begin{equation}\label{eq:parametersFin}
\tau_\psi = T = 5,\quad\quad \sigma = 0.01.
\end{equation}
Each trial lasts 15 seconds in both Horizon 0 and 1 simulations. We set the decision threshold to 5 Hz in all the simulations.

\FloatBarrier


\subsection{Effects of varying the bias and the difference between the stimuli}

The results in Figure~\ref{fig:psiAllTypes} focus on the effects of varying the initial condition $\psi(0)$ of the reuglatory module, therefore the effects of varying the bias. The decision threshold to measure the reaction time $\bar{t}$ is $\lvert v_{e_A}(\bar{t}) - v_{e_B}(\bar{t})  \rvert = 7.5\; \text{Hz}$. We use Euler-Maruyama scheme with time step $\Delta t = 0.5$ and $t_f = 4$ for the presented results in Figure~\ref{fig:psiAllTypes}. Stimuli are applied at $t_0=2$. The parameters in this framework are as follows:
\begin{equation}\label{eq:parametersSmoothStim1}
\begin{split}
T &= 0.003,\quad \sigma=0.01, \quad \tau_w = 500\; (\text{for RS}),\; 10^{-9}\;(\text{for FS}), \\
a & = 1\; (\text{RS}),\; 0 (\text{FS}),\quad b= 100\; (\text{RS}),\; 0\; (\text{FS}),\quad E_L = -65.
\end{split}
\end{equation}\label{eq:parametersSmoothStim2}
Moreover, in~\eqref{eq:inputTF_noRegulation} we fix
\begin{equation}
\begin{split}
s_{\text{AI}}= 5\, \text{Hz},\quad w_c=2.
\end{split}
\end{equation}
Finally, the parameters appearing in~\eqref{eq:regulatory_Cauchy}  are
\begin{equation}\label{eq:parametersSmoothStim3}
\tau_\psi = 10\,T = 0.03,\quad\quad \sigma = 0.01,\quad\quad c_0=1000.
\end{equation}

%In Figure~\ref{fig:examplePsiEffect}, we show an example of the applied stimuli and the results of the case with $\psi(0)=0$. In Figure~\ref{fig:PsiEffectFullSys}, we provide the results of the cases with $\psi(0)$ varied from $0$ to $1$, where the same stimuli shown in Figure~\ref{fig:examplePsiEffect} were applied. In the cases where $\psi$ converges to $1$, i.e., where Stimulus $A$ is chosen, the reaction time is lower ($\approx 0.1$ s) than the cases where $\psi$ converges to $0$, i.e., where Stimulus $B$ is chosen ($\approx 0.18$ s). This is due to that we chose $s_A(0)>s_B(0)$ in the trials given in Figure~\ref{fig:examplePsiEffect}, and therefore $v_{e_A}(\bar{t})-v_{e_B}(\bar{t})=7.5$ is reached in a shorter time duration compared to $v_{e_A}(\bar{t})-v_{e_B}(\bar{t})=-7.5$.

In Figure~\ref{fig:PsiEffectFullSys}, we provide the results of the cases with $\psi(0)$ varied from $0$ to $1$, where the same stimuli were applied in each case. There is no clear difference in terms of reaction time between the cases with different initial conditions $\psi(0)$ as long as the stimuli remain the same and $\psi$ converges to the same value. It is due to the fact that, the convergence of the regulatory mechanism is rapid, therefore the competition is promoted towards the same L2/3 population. As we do not change the stimuli, the evolution of $v_{e_A}, v_{e_B}$ becomes different realizations of almost the same random processes. Consequently, the reaction times of those realizations fluctuate around the same value. 

%In the cases where $\psi$ converges to $1$, i.e., where Stimulus $A$ is chosen, the reaction time is lower ($\approx 0.1$ s) than the cases where $\psi$ converges to $0$, i.e., where Stimulus $B$ is chosen ($\approx 0.18$ s). This is due to that Stimulus A is stronger initially in each case, i.e., $s_A(0)>s_B(0)$. Therefore, $v_{e_A}(\bar{t})-v_{e_B}(\bar{t})=7.5$ is reached in a shorter time duration compared to $v_{e_A}(\bar{t})-v_{e_B}(\bar{t})=-7.5$. 


In Figure~\ref{fig:differentA_B_Stimuli}, we show the effects of varying the difference between the stimuli on the model reaction time. The reaction time is measured in the same way as in Figure~\ref{fig:PsiEffectFullSys}. We set the amplitude of Stimuli $A$ to $6$ and vary the amplitude of Stimuli $B$ between 0 and 6. We keep the initial value $\psi(0)$ of the regulatory function constant ($=0.75$) such that the system privileges always Stimulus $A$. We observe that the reaction time increases as the difference between the stimuli decreases. It is expected since it becomes more difficult to make a distinction between the stimuli as the difference between the filled quantities of the bars decreases.


\begin{figure}[!htbp]
\begin{minipage}[c]{\textwidth}
%\sidesubfloat[(a)]{\hspace{1.1cm}
%  \includegraphics[clip,width=0.8\columnwidth]{examplePsiEffect.png}\label{fig:examplePsiEffect}
%}
  
\sidesubfloat[(a)]{\hspace{0.2cm} \includegraphics[clip,width=0.9\columnwidth]{PsiEffectFullSys.png}\label{fig:PsiEffectFullSys}
}

\sidesubfloat[(b)]{\hspace{0.2cm}
  \includegraphics[clip,width=0.9\columnwidth]{differentA_B_Stimuli.png}\label{fig:differentA_B_Stimuli}
}


\end{minipage}

\caption[labelsep=space]{Simulation results of isolated trials, with varying regulatory module initial conditions $\psi(0)$ in (a) and varying stimuli amplitudes in (b). The axes are specified only in the first columns in both panels, and they are identical in the other columns. \textbf{(a)} Top: Time courses of $v_{e_A}$ and $v_{e_B}$. Middle: Difference of the firing rates where the red vertical line denotes the decision instant. Bottom: Time evolution of the regulatory function $\psi(t)$. The initial values $\psi(0)$ are 0, 0.25, 0.5, 0.75, 1 from left to right. \textbf{(c)} Top row: Time courses of the external stimuli $s_A$ and $s_B$. Second row: Time courses of $v_{e_A}$ and $v_{e_B}$. Third row: Time course of the difference between the membrane potentials where the red vertical line denotes the decision instant. Bottom row: Time course of the regulatory function $\psi$. The initial value $\psi(0)$ is 0.75 in all plots.} \label{fig:psiAllTypes}
\end{figure}

\FloatBarrier


\subsection{Characterization of the model behavioral performance}

Learning speed determines how quickly the model learns the preset strategy. The higher it is, the faster the model identifies the strategy. Flexibility parameter determines how much the model deviates from the preset strategy after it learns the strategy. It models the exploratory behavior and the deviations caused by perceptual difficulties. Finally, the gain rescales the learning speed. Here we provide the effects of these three parameters on the model behavior performance.

%Initial stimuli in the model are $M_1^E\pm \frac{d}{2}$, where $M_1^E$ is generated from a uniform distribution independently for each episode and $d$ is a constant. Here $d$ determines the difficulty of the trial. If $d$ is small, then it is more difficult to make a distinction between two stimuli. The value of $d$ remains constant throughout one episode, however, this value need not necessarily be the same for different episodes.

We show in Figure~\ref{fig:learningSpeedEffect} the performance index plots (human case) in which the learning speed $k$ appearing in~\eqref{eq:rewardMechanismEqn} varies from 0.1 to 0.3. We denote by $V^E_n$ the value of the chosen stimulus in the $n^{\text{th}}$ trial of the $E^{\text{th}}$ episode. We denote the maximum and minimum cumulative reward values of episode $E$ by $V_{\text{min}}^E$ and $V_{\text{max}}^E$, respectively. This is equivalent to maximum and minimum values that $\displaystyle\sum_{n=1}^{L} V^E_n$ can attain among all possible scenarios of the $E^{\text{th}}$ episode, with $L$ denoting the number of trials in the episode. The performance index of the model in the $E^{\text{th}}$ episode is computed as
\begin{equation}\label{eq:scoreMeasure}
\text{Performance index}\,(E) \coloneqq \frac{\displaystyle\sum_{n=1}^{L} V^E_n-V_\text{min}^E}{V_\text{max}^E - V_\text{min}^E}.
\end{equation}
We observe in Figure~\ref{fig:learningSpeedEffect} that as we increase $k$, the system captures the strategy earlier, and makes constantly right decisions thereafter, with very few deviations.

The gain $G$ is another factor affecting the learning speed. The learning speed decreases as the gain decreases since the gain $G$ is equal to the multiplying factor $(M^E_{n+1}-M_n^E)$ of the learning speed $k$ in~\eqref{eq:rewardMechanismEqn}. As shown in Figure~\ref{fig:gainEffect}, the model learns the preset strategy faster as we increase the gain.

Finally, the flexibility parameter $c_0$ determines the number of deviations in the model performance results. The higher it is, the closer to the deterministic case the model is. This is due to the fact that the noise given in~\eqref{eq:regulatory_Cauchy}, and which produces the performance deviations, vanishes very quickly after that the regulatory variable $\psi$ starts to evolve at the beginning of each trial; see Figure~\ref{fig:c0EffectV}. Moreover, $c_0$ determines the beginning of the performance cluster together with the learning speed. As $c_0$ increases, the performance cluster is more likely to begin from smaller episode numbers.


%\begin{figure}[!htbp]
%  \includegraphics[scale=0.8]{globalMeasures}
%  \caption{}
%  \label{fig:globalMeasures}
%\end{figure}


\begin{figure}[!htbp]
\begin{minipage}[c]{\textwidth}
\sidesubfloat[(a)]{\hspace{3.1cm}\includegraphics[clip,scale=0.95]{globalMeasures}\label{fig:globalMeasures}
}

\bigskip

\sidesubfloat[(b)]{\includegraphics[clip,scale=1
]{learningSpeedEffect}\label{fig:learningSpeedEffect}
}

\bigskip

\sidesubfloat[(c)]{  \includegraphics[clip,scale=1]{gainEffect}\label{fig:gainEffect}
}

\bigskip
  
\sidesubfloat[(d)]{\includegraphics[clip,scale=1]{c0EffectV}\label{fig:c0EffectV}
}
\end{minipage}
\caption{Objects used for quantification of behavior in (a), and performance indexes regarding the Horizon 1 human task simulations in (b), (c) and (d). \textbf{(a)} Purple samples correspond to a deviation block, and they are out-cluster deviations. Orange samples are in-cluster deviations, they do not form a deviation block. Both the purple and orange samples are performance deviations. Blue samples are maximum performance indexes and they form a performance cluster. The vertical red line marks the initial cluster episode, it corresponds to the beginning of the performance cluster. \textbf{(b)} Learning speed $k$ varies. The flexibility parameter $c_0$ is $2$ and the gain $G$ is 0.3. The initial episodes of the performance clusters are episodes 24, 15 and 5 from left to right. \textbf{(c)} $G=0.05$, $G=0.15$ and $G=0.3$ from left to right. Here $k=0.3$ and $c_0=2$. The initial episodes of the performance clusters are episodes 81, 10 and 5 from left to right. \textbf{(d)} $c_0=1$, $c_0=2$ and $c_0=3$ from left to right. Here $k=0.05$ and the gain $G=0.3$. The initial episodes of the performance clusters are episodes 52, 40 and 20; the number of in-cluster deviations are 12, 5 and 4 from left to right. The difficulty $d$ is 0.2 in (b), (c) and (d).}
\label{fig:FigPerformanceSimulation}
\end{figure}


\subsection{Characterization of the model reaction time}
In Figure~\ref{fig:RT_H1_T1_T2}, we provide the histograms of the reaction time measurements obtained from the Horizon 1 simulations (human case). In these simulations, we varied the learning speed $k$ and the flexibility parameter $c_0$, separately in Trial 1 and Trial 2. We observe that there is no distinguishing change in the histograms of the simulation results, suggesting that these two parameters do not have noticeable effects on the reaction times. This is due to the fact that the reaction times are calculated by checking the threshold crossing of the difference between the firing rates $v_{e_A}$ and $v_{e_B}$. Here $k$ and $c_0$ do not enter in the equations that describe the firing rates, so they do not influence the reaction times. As the decision threshold increases, the competition between $v_{e_A}$ and $v_{e_B}$ lasts longer, thus the reaction times increase as shown in the bottom row of Figure~\ref{fig:RT_H1_T1_T2}.

\begin{figure}[!htbp]
\noindent\makebox[\textwidth]{
  \includegraphics[scale=0.85]{RT_H1_T1_T2}
  }\caption{Simulation results regarding Horizon 1 reaction time histograms of Trials 1 and 2. Varying the learning speed $k$ and the flexibility parameter $c_0$ does not affect noticeably the distribution of the reaction times in terms of mean and standard deviation. Increasing the decision threshold increases the mean and the standard deviation of the distribution. The  histograms are obtained from 10 realizations of the same simulation setup for each $k$, $c_0$ and decision threshold.}
  \label{fig:RT_H1_T1_T2}
\end{figure}

\FloatBarrier


%\subsection*{Effect of varying the difference between stimuli}
%
%In Figure~\ref{fig:differentA_B_Stimuli}, we observe the effects of varying the difference between the stimuli on the model reaction time. The reaction time is measured in the same way as in Figure~\ref{fig:PsiEffectFullSys}. We set the amplitude of Stimuli $A$ to $6$ and vary the amplitude of Stimuli $B$ between 0 and 6. We keep constant ($=0.75$) the initial value $\psi(0)$ of the regulatory function $\psi$ such that the system privileges always Stimulus $A$. We observe that the reaction time increases as the difference between the stimuli decreases. It is expected since it becomes more difficult to make a distinction between the stimuli as the difference between the filled quantities of the bars decreases.

%\begin{figure}[!htbp]
%\noindent\makebox[\textwidth]{
%  \includegraphics{differentA_B_Stimuli.png}
%  }
%  \caption{Simulation results with changing $s_B$. Top row: Time courses of the external stimuli $s_A$ and $s_B$. Second row: Time courses of $v_{e_A}$ and $v_{e_B}$. Third row: Time course of the difference between the membrane potentials where the red vertical line denotes the decision instant. Bottom row: Time course of the regulatory function $\psi$. The initial value $\psi(0)$ is 0.75 in all plots.}
%  \label{fig:differentA_B_Stimuli}
%\end{figure}
%
%\FloatBarrier

%\subsection*{Horizon 0 human performance}
%In Figure~\ref{fig:statisticalH0}, we provide the results of the Horizon 0 simulations similarly to the Horizon 1 results presented in Figure~\ref{fig:statisticalH1}. The red horizontal lines in Figure~\ref{fig:statisticalH0} show the experimental values. We observe in the left column that the learning speed decreases both with increasing $k$ and with increasing $c_0$, resulting in smaller values of the initial episode number of the performance clusters. The parameter $k$ directly controls the learning speed, and $c_0$ contributes to it by avoiding the deviation blocks which might break the performance clusters. In both cases, the decrease in the mean value profiles is convex. We see that in both cases, the model can reproduce close statistics once $k$ and $c_0$ are chosen properly, for example $k=0.3$ and $c_0=1.1$.


%TASIDIK BUNU YUKARIYA!!!!\begin{figure}[!htbp]
%\noindent\makebox[\textwidth]{
%  \includegraphics{statisticalH0}
%  }
%  \caption{Horizon 0 human case. Simulation statistics with respect to the varied learning speed $k$ in the top row, and with respect to the varied flexibility parameter $c_0$ in the bottom row. In the top row, $c_0=1.1$. In the bottom row, $k=0.3$. The blue vertical lines show the standard deviations of the corresponding statistical sample. The red dashed lines are the experimental values. The mean and standard deviations are obtained via repeating the simulation 10 times for each $(k, c_0)$ pair. See \eqref{eq:parametersIn}-\eqref{eq:parametersFin} for the rest of the parameters.}
%  \label{fig:statisticalH0}
%\end{figure}

\FloatBarrier

%\subsection*{Horizon 0 human reaction times}
%{\color{red}REMOVE THIS, iIn Figure~\ref{fig:RT_H0}, we provide the histograms of the reaction times obtained from Horizon 0 simulations with the varied learning speed $k$ and flexibility parameter $c_0$. We observe that there is no distinguishing change in the histograms as we vary $k$ and $c_0$, suggesting that these two parameters do not have noticeable effects on the reaction times.}
%
%%SILDIK BUNU!!!!\begin{figure}[!htbp]
%%\noindent\makebox[\textwidth]{
%%  \includegraphics{RT_H0}
%%  }
%%\caption{Horizon 0 reaction time histograms. Top: The histograms of the varied $k$, with $c_0=1.1$, $\text{decision threshold}=5\;\text{Hz}$. Middle: The histograms of the varied $c_0$, with $k=0.3$, $\text{decision threshold}=5 \;\text{Hz}$. Bottom: The histograms of the varied decision threshold, where $k=0.3$, $c_0=1.1$. The histograms are obtained for each parameter from 10 realizations of the same simulation setup. The parameters are the same as those of Figure~\ref{fig:statisticalH0}.}
%%  \label{fig:RT_H0}
%%\end{figure}
%
%In Figure~\ref{fig:H0_RT_Histogram}, we show the simulation and experiment histograms of the reaction times corresponding to Horizon 0, with the fitted distributions. Here $k$ and $c_0$ are fixed to the values providing the best performance overlap as in the Horizon 1 case. The best fit is achieved with skewnorm distribution and hypergeometric distribution for the experiment and simulation histograms, respectively. Squared sum errors are 13e-6 and 12e-6 for the experiment and simulation histograms, respectively. Pearson correlation coefficient between the fitted distributions is 0.91.
%
%%TASIDIK BUNU YUKARI\begin{figure}[!htbp] 
%%\noindent\makebox[\textwidth]{
%%  \includegraphics{H0_RT_Histogram}
%%  }
%%  \caption{Horizon 0 human case. Histograms obtained from the human experiment and simulations for comparison. The red curves denote the fitted distributions via Python. The simulations are performed with $k=0.3$, $c_0=1.1$ and $\text{decision threshold}=0.5$.}
%%  \label{fig:H0_RT_Histogram}
%%\end{figure}

\FloatBarrier

\section*{Acknowledgments}
We hereby acknowledge that this research was supported by the Human Brain Project (European Union grant H2020-945539).

%\nolinenumbers

% Either type in your references using
% \begin{thebibliography}{}
% \bibitem{}
% Text
% \end{thebibliography}
%
% or
%
% Compile your BiBTeX database using our plos2015.bst
% style file and paste the contents of your .bbl file
% here. See http://journals.plos.org/plosone/s/latex for 
% step-by-step instructions.
% 


%\begin{thebibliography}{10}
%
%
%\bibitem{albantakis2011changes}
%{\sc L.~Albantakis and G.~Deco}, {\em Changes of mind in an attractor network
%  of decision-making}, PLoS Comput Biol, 7 (2011), p.~e1002086.
%
%\bibitem{amari1977dynamics}
%{\sc S.-i. Amari}, {\em Dynamics of pattern formation in lateral-inhibition
%  type neural fields}, Biological Cybernetics, 27 (1977), pp.~77--87.
%
%\bibitem{arnold1974stochastic}
%{\sc L.~Arnold}, {\em Stochastic differential equations}, New York,  (1974).
%
%\bibitem{baspinarJupyterEbrains2023}
%{\sc E.~Baspinar, G.~Cecchini, R.~Moreno-Bote, I.~Cos, and A.~Destexhe}, {\em
%  {Double columnar Adaptive Exponential mean-field model for decision-making
%  }}, \textit{EBRAINS}, 2023.
%
%
%\bibitem{baspinarJupyter2023}
%\leavevmode\vrule height 2pt depth -1.6pt width 23pt, {\em {Jupyter notebook of
%  a biophysically plausible decision-making model based on interacting cortical
%  columns}}, \textit{Zenodo}, 2023.
%
%\bibitem{bechara2000characterization}
%{\sc A.~Bechara, D.~Tranel, and H.~Damasio}, {\em Characterization of the
%  decision-making deficit of patients with ventromedial prefrontal cortex
%  lesions}, Brain, 123 (2000), pp.~2189--2202.
%
%\bibitem{brette2005adaptive}
%{\sc R.~Brette and W.~Gerstner}, {\em Adaptive exponential integrate-and-fire
%  model as an effective description of neuronal activity}, Journal of
%  Neurophysiology, 94 (2005), pp.~3637--3642.
%
%\bibitem{broche2016neural}
%{\sc Y.~Broche-P{\'e}rez, L.~H. Jim{\'e}nez, and E.~Omar-Mart{\'\i}nez}, {\em
%  Neural substrates of decision-making}, Neurolog{\'\i}a (English Edition), 31
%  (2016), pp.~319--325.
%
%\bibitem{brunel2001effects}
%{\sc N.~Brunel and X.-J. Wang}, {\em Effects of neuromodulation in a cortical
%  network model of object working memory dominated by recurrent inhibition},
%  Journal of Computational Neuroscience, 11 (2001), pp.~63--85.
%
%\bibitem{brzosko2015}
%{\sc Z.~Brzosko, W.~Schultz, and O.~Paulsen}, {\em Retroactive modulation of
%  spike timing-dependent plasticity by dopamine}, eLife, 4 (2015), p.~e09685.
%
%\bibitem{brzosko2017}
%{\sc Z.~Brzosko, S.~Zannone, W.~Schultz, C.~Clopath, and O.~Paulsen}, {\em
%  Sequential neuromodulation of hebbian plasticity offers mechanism for
%  effective reward-based navigation}, eLife, 6 (2017), p.~e27756.
%
%\bibitem{cecchini2024cognitive}
%{\sc G.~Cecchini, M.~DePass, E.~Baspinar, M.~Andujar, S.~Ramawat, P.~Pani,
%  S.~Ferraina, A.~Destexhe, R.~Moreno-Bote, and I.~Cos}, {\em Consequence
%  assessment and behavioral patterns of inhibition in decision-making:
%  modelling its underlying mechanisms}, bioRxiv,  (2023).
%
%\bibitem{churchland2008decision}
%{\sc A.~K. Churchland, R.~Kiani, and M.~N. Shadlen}, {\em Decision-making with
%  multiple alternatives}, Nature Neuroscience, 11 (2008), pp.~693--702.
%
%\bibitem{humanEbrains}
%{\sc M.~DePass, G.~Cecchini, and I.~Cos}, {\em Consequence assessment and
%  behavioral patterns of inhibition in decision-making (v1)}, EBRAINS dataset,
%  2023.
%  
%\bibitem{desimone1995neural}
%{\sc R.~Desimone and J.~Duncan}, {\em Neural mechanisms of selective visual attention}, Annual Rreview of Neuroscience, 18 (1995), pp.~193--222.
%  
%
%\bibitem{di2019biologically}
%{\sc M.~Di~Volo, A.~Romagnoni, C.~Capone, and A.~Destexhe}, {\em Biologically
%  realistic mean-field models of conductance-based networks of spiking neurons
%  with adaptation}, Neural Computation, 31 (2019), pp.~653--680.
%
%\bibitem{domenech2015executive}
%{\sc P.~Domenech and E.~Koechlin}, {\em Executive control and decision-making
%  in the prefrontal cortex}, Current Opinion in Behavioral Sciences, 1 (2015),
%  pp.~101--106.
%
%\bibitem{doya2007bayesian}
%{\sc K.~Doya, S.~Ishii, A.~Pouget, and R.~P. Rao}, {\em Bayesian brain:
%  Probabilistic approaches to neural coding}, MIT press, 2007.
%
%\bibitem{drugowitsch2012cost}
%{\sc J.~Drugowitsch, R.~Moreno-Bote, A.~K. Churchland, M.~N. Shadlen, and
%  A.~Pouget}, {\em The cost of accumulating evidence in perceptual decision
%  making}, Journal of Neuroscience, 32 (2012), pp.~3612--3628.
%
%\bibitem{el2009master}
%{\sc S.~El~Boustani and A.~Destexhe}, {\em A master equation formalism for
%  macroscopic modeling of asynchronous irregular activity states}, Neural
%  Computation, 21.
%
%\bibitem{finkelstein2021attractor}
%{\sc A.~Finkelstein, L.~Fontolan, M.~N. Economo, N.~Li, S.~Romani, and
%  K.~Svoboda}, {\em Attractor dynamics gate cortical information flow during
%  decision-making}, Nature Neuroscience,  (2021), pp.~1--8.
%
%\bibitem{macaqueEbrains}
%{\sc R.~Fontana, M.~Andujar, I.~B. Marc, V.~Giuffrida, S.~Ramawat, G.~Bardella,
%  E.~Brunamonti, P.~Pani, and S.~Ferraina}, {\em Evaluating consequences in
%  decision-making: behavioral data and neural recordings from monkey premotor
%  and prefrontal cortex (v1)}, EBRAINS dataset, 2022.
%
%\bibitem{funahashi2017prefrontal}
%{\sc S.~Funahashi}, {\em Prefrontal contribution to decision-making under
%  free-choice conditions}, Frontiers in neuroscience, 11 (2017), p.~431.
%
%\bibitem{giamundo2021neuronal}
%{\sc M.~Giamundo, F.~Giarrocco, E.~Brunamonti, F.~Fabbrini, P.~Pani, and
%  S.~Ferraina}, {\em Neuronal activity in the premotor cortex of monkeys
%  reflects both cue salience and motivation for action generation and
%  inhibition}, Journal of Neuroscience, 41 (2021), pp.~7591--7606.
%
%\bibitem{gilbert1979morphology}
%{\sc C.~D. Gilbert and T.~N. Wiesel}, {\em Morphology and intracortical
%  projections of functionally characterised neurones in the cat visual cortex},
%  Nature, 280 (1979), pp.~120--125.
%
%\bibitem{goldman2020brain}
%{\sc J.~S. Goldman, L.~Kusch, B.~H. Yalcinkaya, D.~Depannemaecker, T.-A.~E.
%  Nghiem, V.~Jirsa, and A.~Destexhe}, {\em Brain-scale emergence of slow-wave
%  synchrony and highly responsive asynchronous states based on biologically
%  realistic population models simulated in the virtual brain}, bioRxiv,
%  (2020).
%
%\bibitem{guo2013brain}
%{\sc Z.~Guo, J.~Chen, S.~Liu, Y.~Li, B.~Sun, and Z.~Gao}, {\em Brain areas
%  activated by uncertain reward-based decision-making in healthy volunteers},
%  Neural Regeneration Research, 8 (2013), p.~3344.
%
%\bibitem{hampton2006role}
%{\sc A.~N. Hampton, P.~Bossaerts, and J.~P. O’doherty}, {\em The role of the
%  ventromedial prefrontal cortex in abstract state-based inference during
%  decision making in humans}, Journal of Neuroscience, 26 (2006),
%  pp.~8360--8367.
%
%\bibitem{hayden2018neuronal}
%{\sc B.~Y. Hayden and R.~Moreno-Bote}, {\em A neuronal theory of sequential
%  economic choice}, Brain and Neuroscience Advances, 2 (2018),
%  p.~2398212818766675.
%
%\bibitem{kiani2008bounded}
%{\sc R.~Kiani, T.~D. Hanks, and M.~N. Shadlen}, {\em Bounded integration in
%  parietal cortex underlies decisions even when viewing duration is dictated by
%  the environment}, Journal of Neuroscience, 28 (2008), pp.~3017--3029.
%
%\bibitem{kisvarday1986synaptic}
%{\sc Z.~F. Kisv{\'a}rday, K.~A. Martin, T.~Freund, Z.~Magloczky,
%  D.~Whitteridge, and P.~Somogyi}, {\em Synaptic targets of hrp-filled layer
%  iii pyramidal cells in the cat striate cortex}, Experimental Brain Research,
%  64 (1986), pp.~541--552.
%
%\bibitem{kopec2015cortical}
%{\sc C.~D. Kopec, J.~C. Erlich, B.~W. Brunton, K.~Deisseroth, and C.~D. Brody},
%  {\em Cortical and subcortical contributions to short-term memory for
%  orienting movements}, Neuron, 88 (2015), pp.~367--377.
%
%\bibitem{leotti2010motivational}
%{\sc L.~A. Leotti and T.~D. Wager}, {\em Motivational influences on response
%  inhibition measures.}, Journal of Experimental Psychology: Human Perception
%  and Performance, 36 (2010), p.~430.
%
%\bibitem{marcos2013neural}
%{\sc E.~Marcos, P.~Pani, E.~Brunamonti, G.~Deco, S.~Ferraina, and
%  P.~Verschure}, {\em Neural variability in premotor cortex is modulated by
%  trial history and predicts behavioral performance}, Neuron, 78 (2013),
%  pp.~249--255.
%
%\bibitem{martin1984form}
%{\sc K.~Martin and D.~Whitteridge}, {\em Form, function and intracortical
%  projections of spiny neurones in the striate visual cortex of the cat.}, The
%  Journal of Physiology, 353 (1984), pp.~463--504.
%
%\bibitem{mcguire1991targets}
%{\sc B.~A. McGuire, C.~D. Gilbert, P.~K. Rivlin, and T.~N. Wiesel}, {\em
%  Targets of horizontal connections in macaque primary visual cortex}, Journal
%  of Comparative Neurology, 305 (1991), pp.~370--392.
%  
%\bibitem{mcclure2004}
%{\sc S.~M. McClure, D.~I. Laibson, G. Loewenstein, and J.~D. Cohen}, {\em
%  Separate neural systems value immediate and delayed monetary rewards}, 306 (2004), pp.~503--507.  
%  
% 
%  
%  
%\bibitem{mirenowicz1994importance}
%{\sc J.~Mirenowicz and W.~Schultz}, {\em Importance of unpredictability for
%  reward responses in primate dopamine neurons}, Journal of neurophysiology, 72
%  (1994), pp.~1024--1027.
%
%\bibitem{mochol2021prefrontal}
%{\sc G.~Mochol, R.~Kiani, and R.~Moreno-Bote}, {\em Prefrontal cortex
%  represents heuristics that shape choice bias and its integration into future
%  behavior}, Current Biology, 31 (2021), pp.~1234--1244.
%
%\bibitem{monosov2012regionally}
%{\sc I.~E. Monosov and O.~Hikosaka}, {\em Regionally distinct processing of
%  rewards and punishments by the primate ventromedial prefrontal cortex},
%  Journal of Neuroscience, 32 (2012), pp.~10318--10330.
%  
%\bibitem{moore2017neural}
%{\sc T.~Moore and M.~Zirnsak}, {\em Neural mechanisms of selective visual attention},
%  Annual Review of Psychology, 68 (2017), pp.~47--72.
%
%\bibitem{moreno2007noise}
%{\sc R.~Moreno-Bote, J.~Rinzel, and N.~Rubin}, {\em Noise-induced alternations
%  in an attractor network model of perceptual bistability}, Journal of
%  Neurophysiology, 98 (2007), pp.~1125--1139.
%
%\bibitem{padmala2010interactions}
%{\sc S.~Padmala and L.~Pessoa}, {\em Interactions between cognition and
%  motivation during response inhibition}, Neuropsychologia, 48 (2010),
%  pp.~558--565.
%
%\bibitem{parker1998sense}
%{\sc A.~J. Parker and W.~T. Newsome}, {\em Sense and the single neuron: probing
%  the physiology of perception}, Annual Review of Neuroscience, 21 (1998),
%  pp.~227--277.
%
%\bibitem{paulus2003increased}
%{\sc M.~P. Paulus, C.~Rogalsky, A.~Simmons, J.~S. Feinstein, and M.~B. Stein},
%  {\em Increased activation in the right insula during risk-taking decision
%  making is related to harm avoidance and neuroticism}, Neuroimage, 19 (2003),
%  pp.~1439--1448.
%  
%\bibitem{reynolds2004attentional}
%{\sc J.~H. Reynolds and L.~Chelazzi},
%  {\em Attentional modulation of visual processing}, Annual Review of Neuroscience, 27 (2004),
%  pp.~611--647.  
%
%\bibitem{romo2001touch}
%{\sc R.~Romo and E.~Salinas}, {\em Touch and go: decision-making mechanisms in
%  somatosensation}, Annual Review of Neuroscience, 24 (2001), pp.~107--137.
%
%\bibitem{rushworth2011frontal}
%{\sc M.~F. Rushworth, M.~P. Noonan, E.~D. Boorman, M.~E. Walton, and T.~E.
%  Behrens}, {\em Frontal cortex and reward-guided learning and
%  decision-making}, Neuron, 70 (2011), pp.~1054--1069.
%
%\bibitem{sanfey2006neuroeconomics}
%{\sc A.~G. Sanfey, G.~Loewenstein, S.~M. McClure, and J.~D. Cohen}, {\em
%  Neuroeconomics: cross-currents in research on decision-making}, Trends in
%  Cognitive Sciences, 10 (2006), pp.~108--116.
%
%\bibitem{sanz2013virtual}
%{\sc P.~Sanz~Leon, S.~A. Knock, M.~M. Woodman, L.~Domide, J.~Mersmann, A.~R.
%  McIntosh, and V.~Jirsa}, {\em The virtual brain: a simulator of primate brain
%  network dynamics}, Frontiers in neuroinformatics, 7 (2013), p.~10.
%
%\bibitem{schall2001neural}
%{\sc J.~D. Schall}, {\em Neural basis of deciding, choosing and acting}, Nature
%  Reviews Neuroscience, 2 (2001), pp.~33--42.
%
%\bibitem{schultz2022dopamine}
%{\sc W.~Schultz}, {\em Dopamine reward prediction error coding}, Dialogues in
%  clinical neuroscience,  (2022).
%
%\bibitem{seidemann1998temporal}
%{\sc E.~Seidemann, E.~Zohary, and W.~T. Newsome}, {\em Temporal gating of
%  neural signals during performance of a visual discrimination task}, Nature,
%  394 (1998), pp.~72--75.
%
%\bibitem{shadlen2013decision}
%{\sc M.~N. Shadlen and R.~Kiani}, {\em Decision making as a window on
%  cognition}, Neuron, 80 (2013), pp.~791--806.
%
%\bibitem{sirigu2022reward}
%{\sc A.~Sirigu and J.-R. Duhamel}, {\em Reward and decision processes in the
%  brains of humans and nonhuman primates}, Dialogues in Clinical Neuroscience,
%  (2022).
%
%\bibitem{suzuki2021distinct}
%{\sc S.~Suzuki, V.~M. Lawlor, J.~A. Cooper, A.~R. Arulpragasam, and M.~T.
%  Treadway}, {\em Distinct regions of the striatum underlying effort, movement
%  initiation and effort discounting}, Nature human behaviour, 5 (2021),
%  pp.~378--388.
%
%\bibitem{turan2023whole}
%{\sc A.~Turan, E.~Baspinar, and A.~Destexhe}, {\em A whole-brain model of
%  auditory discrimination}, bioRxiv,  (2023), pp.~2023--09.
%
%\bibitem{van2018decoding}
%{\sc F.~Van~Ede, S.~R. Chekroud, M.~G. Stokes, and A.~C. Nobre}, {\em Decoding
%  the influence of anticipatory states on visual perception in the presence of
%  temporal distractors}, Nature Communications, 9 (2018), pp.~1--12.
%
%\bibitem{wang2002probabilistic}
%{\sc X.-J. Wang}, {\em Probabilistic decision making by slow reverberation in
%  cortical circuits}, Neuron, 36 (2002), pp.~955--968.
%
%\bibitem{wilson1972excitatory}
%{\sc H.~R. Wilson and J.~D. Cowan}, {\em Excitatory and inhibitory interactions
%  in localized populations of model neurons}, Biophysical Journal, 12 (1972),
%  pp.~1--24.
%
%\bibitem{xiao2021data}
%{\sc Z.-C. Xiao, K.~K. Lin, and L.-S. Young}, {\em A data-informed mean-field
%  approach to mapping of cortical parameter landscapes}, PLoS Computational
%  Biology, 17 (2021), p.~e1009718.
%  
%\bibitem{zerlaut2016heterogeneous}
%{\sc Y.~Zerlaut, B.~Tele{\'n}czuk, C.~Deleuze, T.~Bal, G.~Ouanounou, and A.~Destexhe}, {\em Heterogeneous firing rate response of mouse layer V pyramidal neurons in the fluctuation-driven regime}, The Journal of Physiology, 13 (2016), pp.~3791--3808.
%
%\bibitem{zerlaut2018modeling}
%{\sc Y.~Zerlaut, S.~Chemla, F.~Chavane, and A.~Destexhe}, {\em Modeling
%  mesoscopic cortical dynamics using a mean-field model of conductance-based
%  networks of adaptive exponential integrate-and-fire neurons}, Journal of
%  Computational Neuroscience, 44 (2018), pp.~45--61.
%
%\bibitem{zuo2019rats}
%{\sc Y.~Zuo and M.~E. Diamond}, {\em Rats generate vibrissal sensory evidence
%  until boundary crossing triggers a decision}, Current Biology, 29 (2019),
%  pp.~1415--1424.
%
%
%\end{thebibliography}


\bibliographystyle{APA}
\begin{thebibliography}{100}
\providecommand{\natexlab}[1]{#1}
\expandafter\ifx\csname urlstyle\endcsname\relax
  \providecommand{\doi}[1]{doi:\discretionary{}{}{}#1}\else
  \providecommand{\doi}{doi:\discretionary{}{}{}\begingroup
  \urlstyle{rm}\Url}\fi


\bibitem[{Albantakis et~al.(2011)Albantakis, \& Deco}]{albantakis2011changes}
Albantakis, L., \& Deco, G. (2011).
\newblock Changes of mind in an attractor network of decision-making.
\newblock \emph{PLoS Comput. Biol.}, \emph{7}, e1002086

\bibitem[{Amari(1977)}]{amari1977dynamics}
Amari, S. I. (1977).
\newblock Dynamics of pattern formation in lateral-inhibition type neural fields.
\newblock \emph{Biological Cybernetics}, \emph{27}, 77 -- 87.

\bibitem[{Arnold(1974)}]{arnold1974stochastic}
Arnold, L. (1974).
\newblock Stochastic differential equations: theory and applications.
\newblock \emph{Wiley Interscience}\emph{}.

\bibitem[{Baspinar et~al.(2023)Baspinar, Cecchini, Moreno-Bote, Cos, \& Destexhe}]{baspinarJupyterEbrains2023}
Baspinar, E., Cecchini, G., Moreno-Bote, R., Cos, I., \& Destexhe, A. (2023).
\newblock Double columnar Adaptive Exponential mean-field model for decision-making.
\newblock \emph{EBRAINS} \emph{}.

\bibitem[{Baspinar et~al.(2023)Baspinar, Cecchini, Moreno-Bote, Cos, \& Destexhe}]{baspinarJupyter2023}
Baspinar, E., Cecchini, G., Moreno-Bote, R., Cos, I., \& Destexhe, A. (2023).
\newblock Jupyter notebook of a biophysically plausible decision-making model based on interacting cortical columns.
\newblock \emph{Zenodo} \emph{}.

\bibitem[{Bechara et~al.(2000)Bechara, Tranel, \& Damasio}]{bechara2000characterization}
Bechara, A., Tranel, D., \& Damasio, H. (2000).
\newblock Characterization of the decision-making deficit of patients with ventromedial prefrontal cortex lesions.
\newblock \emph{Brain}, \emph{123}, 2189 -- 2202.


\bibitem[{Brette et~al.(2005)Brette, \& Gerstner}]{brette2005adaptive}
Brette, R., \& Gerstner, W. (2005).
\newblock Adaptive exponential integrate-and-fire model as an effective description of neuronal activity.
\newblock \emph{Journal of
  Neurophysiology}, \emph{94}, 3637 -- 3642.
  
  
\bibitem[{Broche-P{\'e}rez et~al.(2016)Broche-P{\'e}rez, Jim{\'e}nez, \& Omar-Mart{\'\i}nez}]{broche2016neural}
Broche-P{\'e}rez, Y., Jim{\'e}nez, L. H., \& Omar-Mart{\'\i}nez, E. (2016).
\newblock Neural substrates of decision-making.
\newblock \emph{Neurolog{\'\i}a (English Edition)}, \emph{31}, 319 -- 325.

\bibitem[{Brunel et~al.(2001)Brunel, \& Wang}]{brunel2001effects}
Brunel, N., \& Wang, X. J. (2001).
\newblock Effects of neuromodulation in a cortical network model of object working memory dominated by recurrent inhibition.
\newblock \emph{Neurolog{\'i}a (English Edition)}, \emph{31}, 319 -- 325.

\bibitem[{Brzosko et~al.(2015)Brzosko, Schultz, \& Paulsen}]{brzosko2015}
Brzosko, Z., Schultz, W., \& Paulsen, O. (2015).
\newblock Retroactive modulation of spike timing-dependent plasticity by dopamine.
\newblock \emph{eLife}, \emph{4}, e09685.

\bibitem[{Brzosko et~al.(2017)Brzosko, Zannone, Schultz, Clopath, \& Paulsen}]{brzosko2017}
Brzosko, Z., Zannone, S., Schultz, W., Clopath, C., \& Paulsen, O. (2017).
\newblock Sequential neuromodulation of Hebbian plasticity offers mechanism for effective reward-based navigation.
\newblock \emph{eLife}, \emph{6}, e27756.

%\bibitem[{Cecchini et~al.(2023)Cecchini, DePass, Baspinar, Andujar, Ramawat, Pani, Ferraina, Destexhe, Moreno-Bote, \& Cos}]{cecchiniPreprint}
%Cecchini, G., DePass, M., Baspinar, E., Andujar, M., Ramawat, S., Pani, P., Ferraina, S., Destexhe, A., Moreno-Bote, R., \& Cos, I. (2023).
%\newblock Consequence assessment and behavioral patterns of inhibition in decision-making: modelling its underlying mechanisms.
%\newblock \emph{bioRxiv} \emph{}.

\bibitem[{Cecchini et~al.(2024)Cecchini, DePass, Baspinar, Andujar, Ramawat, Pani, Ferraina, Destexhe, Moreno-Bote, \& Cos}]{cecchini2024cognitive}
Cecchini, G., DePass, M., Baspinar, E., Andujar, M., Ramawat, S., Pani, P., Ferraina, S., Destexhe, A., Moreno-Bote, R., \& Cos, I. (2024).
\newblock Cognitive mechanisms of learning in sequential decision-making under uncertainty: an experimental and theoretical approach.
\newblock \emph{Front. Behavioral Neuroscience}, \emph{18}.


\bibitem[{Churchland et~al.(2023)Churchland, Kiani, \& Shadlen}]{churchland2008decision}
Churchland, A. K., Kiani, R., \& Shadlen, M.N. (2023).
\newblock Decision-making with multiple alternatives.
\newblock \emph{Nature Neuroscience}, \emph{11}, 693 -- 702.

\bibitem[{DePass et~al.(2023)DePass, Cecchini, \& Cos}]{humanEbrains}
DePass, M., Cecchini, G., \& Cos, I. (2023).
\newblock Consequence assessment and
  behavioral patterns of inhibition in decision-making.
\newblock \emph{EBRAINS dataset} \emph{}.

\bibitem[{Desimone et~al.(1995)Desimone, \& Duncan}]{desimone1995neural}
Desimone, R., \& Duncan, J. (1995).
\newblock Neural mechanisms of selective visual attention. 
\newblock \emph{Annual Review of Neuroscience}, \emph{18}, 193 -- 222.

\bibitem[{di Volo et~al.(2019)di Volo, Romagnoni, Capone, \& Destexhe}]{di2019biologically}
di Volo, M., Romagnoni, A., Capone, C., \& Destexhe, A. (2019).
\newblock Biologically realistic mean-field models of conductance-based networks of spiking neurons with adaptation.
\newblock \emph{Neural Computation}, \emph{31}, 653 -- 680.

\bibitem[{Domenech et~al.(2015)Domenech, \& Koechlin}]{domenech2015executive}
Domenech, P., \& Koechlin, E. (2015).
\newblock Executive control and decision-making in the prefrontal cortex. 
\newblock \emph{Current Opinion in Behavioral Sciences}, \emph{1}, 101 -- 106.

\bibitem[{Doya et~al.(2007)Doya, Ishii, Pouget, \& Rao}]{doya2007bayesian}
Doya, K., Ishii, S., Pouget, A., \& Rao, R. P. (2007).
\newblock Bayesian brain: probabilistic approaches to neural coding. 
\newblock \emph{MIT press} \emph{}.

\bibitem[{Drugowitsch et~al.(2012)Drugowitsch, Moreno-Bote, Churchland, Shadlen, \& Pouget}]{drugowitsch2012cost}
Drugowitsch, J., Moreno-Bote, R., Churchland, A. K., Shadlen, M. N., \& Pouget, A. (2012).
\newblock The cost of accumulating evidence in perceptual decision making. 
\newblock \emph{Journal of Neuroscience}, \emph{32}, 3612 -- 3628.

\bibitem[{El Boustani et~al.(2009)El Boustani, \& Destexhe}]{el2009master}
El Boustani, S., \& Destexhe, A. (2009).
\newblock A master equation formalism for macroscopic modeling of asynchronous irregular activity states. 
\newblock \emph{Neural Computation}, \emph{21}, 46 -- 100.

\bibitem[{Finkelstein et~al.(2021)Finkelstein, Fontolan, Economo, Li, Romani, \& Svoboda}]{finkelstein2021attractor}
Finkelstein, A., Fontolan, L., Economo, M. N., Li, N., Romani, S., \& Svoboda, K. (2021).
\newblock Attractor dynamics gate cortical information flow during decision-making. 
\newblock \emph{Nature Neuroscience}, \emph{24}, 843 -- 850.

\bibitem[{Fontana et~al.(2022)Fontana, Andujar, Marc, Giuffrida, Ramawat, Bardella, Brunamonti, Pani, \& Ferraina}]{macaqueEbrains}
Fontana, R., Andujar, M., Marc, I. B., Giuffrida, V., Ramawat, S., Bardella, G., Brunamonti, E., Pani, P., \& Ferraina, S. (2022).
\newblock Evaluating consequences in decision-making: behavioral data and neural recordings from monkey premotor and prefrontal cortex.
\newblock \emph{EBRAINS dataset} \emph{}.


\bibitem[{Funahashi et~al.(2017)Funahashi}]{funahashi2017prefrontal}
Funahashi, S. (2017).
\newblock Prefrontal contribution to decision-making under
  free-choice conditions. 
\newblock \emph{Frontiers in Neuroscience}, \emph{11}, 431.

\bibitem[{Giamundo et~al.(2021)Giamundo, Giarrocco, Brunamonti, Fabbrini, Pani, \& Ferraina}]{giamundo2021neuronal}
Giamundo, M., Giarrocco, F., Brunamonti, E., Fabbrini, F., Pani, P., \& Ferraina, S. (2021).
\newblock Neuronal activity in the premotor cortex of monkeys reflects both cue salience and motivation for action generation and inhibition. 
\newblock \emph{Journal of Neuroscience}, \emph{41}, 7591 -- 7606.

%\bibitem[{Gilbert et~al.(1979)Gilbert, \& Wiesel}]{gilbert1979morphology}
%Gilbert, C. D., \& Wiesel, T. N. (1979).
%\newblock Morphology and intracortical projections of functionally characterised neurones in the cat visual cortex. 
%\newblock \emph{Nature}, \emph{280}, 120 -- 125.

\bibitem[{Goldman et~al.(2020)Goldman, Kusch, Yalcinkaya, Depannemaecker, Nghiem, Jirsa, \& Destexhe}]{goldman2020brain}
Goldman, J. S., Kusch, L., Yalcinkaya, B. H., Depannemaecker, D., Nghiem, T. A. E., Jirsa, V., \& Destexhe, A. (2020).
\newblock Brain-scale emergence of slow-wave synchrony and highly responsive asynchronous states based on biologically realistic population models simulated in the virtual brain. 
\newblock \emph{bioRxiv} \emph{}.


\bibitem[{Guo et~al.(2013)Guo, Chen, Liu, Li, Sun, \& Gao}]{guo2013brain}
Guo, Z., Chen, J., Liu, S., Li, Y., Sun, B., \& Gao, Z. (2013).
\newblock Brain areas activated by uncertain reward-based decision-making in healthy volunteers. 
\newblock \emph{Neural Regeneration Research}, \emph{8}, 3344.

\bibitem[{Hampton et~al.(2006)Hampton, Bossaerts, \& O’doherty}]{hampton2006role}
Hampton, A. N., Bossaerts, P., \& O’doherty, J. P. (2006).
\newblock The role of the ventromedial prefrontal cortex in abstract state-based inference during decision making in humans. 
\newblock \emph{Journal of Neuroscience}, \emph{26}, 8360 -- 8367.


\bibitem[{Hayden et~al.(2018)Hayden, \& Moreno-Bote}]{hayden2018neuronal}
Hayden, B. Y., \& Moreno-Bote, R. (2018).
\newblock A neuronal theory of sequential
  economic choice. 
\newblock \emph{Brain and Neuroscience Advances}, \emph{2}, 2398212818766675.


\bibitem[{Hirata et~al.(2008)Hirata, \& Sawaguchi}]{hirata2008functional}
Hirata, Y., \& Sawaguchi, Toshiyuki. (2008).
\newblock Functional columns in the primate prefrontal cortex revealed by optical imaging in vitro. 
\newblock \emph{Neuroscience Research}, \emph{61}, 1 -- 10.


\bibitem[{Kiani et~al.(2008)Kiani, Hanks, \& Shadlen}]{kiani2008bounded}
Kiani, R., Hanks, T. D., \& Shadlen, M. N. (2008).
\newblock Bounded integration in parietal cortex underlies decisions even when viewing duration is dictated by the environment. 
\newblock \emph{Journal of Neuroscience}, \emph{28}, 3017 -- 3029.

%\bibitem[{Kisv{\'a}rday et~al.(1986)Kisv{\'a}rday, Martin, Freund, Magloczky, Whitteridge, \& Somogyi}]{kisvarday1986synaptic}
%Kisv{\'a}rday, Z. G., Martin, K. A., Freund, T., Magloczky, Z., Whitteridge, D., \& Somogyi, P. (1986).
%\newblock Synaptic targets of HRP-filled layer III pyramidal cells in the cat striate cortex. 
%\newblock \emph{Experimental Brain Research}, \emph{64}, 541 -- 552.

\bibitem[{Kopec et~al.(2015)Kopec, Erlich, Brunton, Deisseroth, \& Brody}]{kopec2015cortical}
Kopec, C. D., Erlich, J. C., Brunton, B. W., Deisseroth, K., \& Brody, C. D.  (2015).
\newblock Cortical and subcortical contributions to short-term memory for orienting movements. 
\newblock \emph{Neuron}, \emph{88}, 367 -- 377.

\bibitem[{Leotti et~al.(2010)Leotti, \& Wager}]{leotti2010motivational}
Leotti, L. A., \& Wager, T. D. (2010).
\newblock Motivational influences on response inhibition measures. 
\newblock \emph{Journal of Experimental Psychology: Human Perception
  and Performance}, \emph{36}, 430.
  
\bibitem[{Marcos et~al.(2013)Marcos, Pani, Brunamonti, Deco, Ferraina, \& Verschure}]{marcos2013neural}
Marcos, E., Pani, P., Brunamonti, E., Deco, G., Ferraina, S., \& Verschure, P. (2013).
\newblock Neural variability in premotor cortex is modulated by trial history and predicts behavioral performance. 
\newblock \emph{Neuron}, \emph{78}, 249 -- 255.

%\bibitem[{Martin et~al.(1984)Martin, \& Whitteridge}]{martin1984form}
%Martin, K., \& Whitteridge, D. (1984).
%\newblock Form, function and intracortical projections of spiny neurones in the striate visual cortex of the cat. 
%\newblock \emph{The Journal of Physiology}, \emph{353}, 463 -- 504.

\bibitem[{McClure et~al.(2004)McClure, Laibson, David, Loewenstein, \& Cohen}]{mcclure2004}
McClure, S. M., Laibson, D. I., Loewenstein, G., \& Cohen, J. D. (2004).
\newblock Separate neural systems value immediate and delayed monetary rewards. 
\newblock \emph{Science}, \emph{306}, 503 -- 507.

%\bibitem[{McGuire et~al.(1991)McGuire, Gilbert, Rivlin, \& Wiesel}]{mcguire1991targets}
%McGuire, B. A., Gilbert, C. D., Rivlin, P. K., \& Wiesel, T. N. (1991).
%\newblock Targets of horizontal connections in macaque primary visual cortex. 
%\newblock \emph{Journal of Comparative Neurology}, \emph{305}, 370 -- 392.

\bibitem[{Mirenowicz et~al.(1994)Mirenowicz, \& Schultz}]{mirenowicz1994importance}
Mirenowicz, J., \& Schultz, W. (1994).
\newblock Importance of unpredictability for reward responses in primate dopamine neurons. 
\newblock \emph{Journal of Neurophysiology}, \emph{72}, 1024 -- 1027.

\bibitem[{Mochol et~al.(2021)Mochol, Kiani, \& Moreno-Bote}]{mochol2021prefrontal}
Mochol, G., Kiani, R., \& Moreno-Bote, R. (2021).
\newblock Prefrontal cortex represents heuristics that shape choice bias and its integration into future behavior. 
\newblock \emph{Current Biology}, \emph{31}, 1234 -- 1244.

\bibitem[{Monosov et~al.(2012)Monosov, \& Hikosaka}]{monosov2012regionally}
Monosov, I. E., \& Hikosaka, O. (2012).
\newblock Regionally distinct processing of rewards and punishments by the primate ventromedial prefrontal cortex. 
\newblock \emph{Journal of Neuroscience}, \emph{32}, 10318 -- 10330.

\bibitem[{Moore et~al.(2017)Moore, \& Zirnsak}]{moore2017neural}
Moore, T., \& Zirnsak, M. (2017).
\newblock Neural mechanisms of selective visual attention. 
\newblock \emph{Annual Review of Psychology}, \emph{68}, 47 -- 72.

\bibitem[{Moreno-Bote et~al.(2007)Moreno-Bote, Rinzel, \& Rubin}]{moreno2007noise}
Moreno-Bote, R., Rinzel, J., \& Rubin, N. (2007).
\newblock Noise-induced alternations in an attractor network model of perceptual bistability. 
\newblock \emph{Journal of Neurophysiology}, \emph{98}, 1125 -- 1139.

\bibitem[{Padmala et~al.(2010)Padmala, \& Pessoa}]{padmala2010interactions}
Padmala, S., \& Pessoa, L. (2010).
\newblock Interactions between cognition and
  motivation during response inhibition. 
\newblock \emph{Neuropsychologia}, \emph{48}, 558 -- 565.

\bibitem[{Papoulis et~al.(2002)Papoulis \& Unnikrishna}]{papoulis2002Probability}
Papoulis, A., \& Pillai, S. U. (2002).
\newblock Probability, random variables, and stochastic processes. 
\newblock \emph{McGraw-Hill Europe: New York, NY, USA} \emph{}.

\bibitem[{Parker et~al.(1998)Parker, \& Newsome}]{parker1998sense}
Parker, A. J., \& Newsome, W. T. (1998).
\newblock Sense and the single neuron: probing
  the physiology of perception. 
\newblock \emph{Annual Review of Neuroscience}, \emph{21}, 227 -- 277.

\bibitem[{Paulus et~al.(2003)Paulus, Rogalsky, Simmons, Feinstein, \& Stein}]{paulus2003increased}
Paulus, M. P., Rogalsky, C., Simmons, A., Feinstein, J. S., \& Stein, M. (2003).
\newblock Increased activation in the right insula during risk-taking decision making is related to harm avoidance and neuroticism. 
\newblock \emph{Neuroimage}, \emph{19}, 1439 -- 1448.

\bibitem[{Reynolds et~al.(2004)Reynolds, \& Chelazzi}]{reynolds2004attentional}
Reynolds, J. H., \& Chelazzi, L. (2004).
\newblock Attentional modulation of visual processing. 
\newblock \emph{Annual Review of Neuroscience}, \emph{27}, 611 -- 647.


\bibitem[{Romo et~al.(2001)Romo, \& Salinas}]{romo2001touch}
Romo, R., \& Salinas, E. (2001).
\newblock Touch and go: decision-making mechanisms in
somatosensation. 
\newblock \emph{Annual Review of Neuroscience}, \emph{24}, 107 -- 137.

\bibitem[{Rushworth et~al.(2011)Rushworth, Noonan, Boorman, Walton, \& Behrens}]{rushworth2011frontal}
Rushworth, M. F., Noonan, M. P., Boorman, E. D., Walton, M. E., \& Behrens, T. E. (2011).
\newblock Frontal cortex and reward-guided learning and decision-making. 
\newblock \emph{Neuron}, \emph{70}, 1054 -- 1069.

\bibitem[{Sanfey et~al.(2006)Sanfey, Loewenstein, McClure, \& Cohen}]{sanfey2006neuroeconomics}
Sanfey, A. G., Loewenstein, G., McClure, S. M., \& Cohen, J. D. (2006).
\newblock Neuroeconomics: cross-currents in research on decision-making. 
\newblock \emph{Trends in Cognitive Sciences}, \emph{10}, 108 -- 116.

\bibitem[{Sanz~Leon et~al.(2013)Sanz~Leon, Knock, Woodman, Domide, Mersmann, McIntosh, \& Jirsa}]{sanz2013virtual}
Sanz~Leon, P., Knock, S. A., Woodman, M. M., Domide, L., Mersmann, J., McIntosh, A. R., \& Jirsa, V. (2013).
\newblock The Virtual Brain: a simulator of primate brain network dynamics. 
\newblock \emph{Frontiers in Neuroinformatics}, \emph{7}, 10.

\bibitem[{Schall(2001)Schall}]{schall2001neural}
Schall, J. D. (2001).
\newblock Neural basis of deciding, choosing and acting. 
\newblock \emph{Nature Reviews Neuroscience}, \emph{2}, 33 -- 42.

\bibitem[{Schultz(2022)Schultz}]{schultz2022dopamine}
Schultz, W. (2022).
\newblock Dopamine reward prediction error coding. 
\newblock \emph{Dialogues in
  Clinical Neuroscience} \emph{}.


\bibitem[{Seidemann et~al.(1998)Seidemann, Zohary, \& Newsome}]{seidemann1998temporal}
Seidemann, E., Zohary, E., \& Newsome, W. T. (1998).
\newblock Temporal gating of neural signals during performance of a visual discrimination task. 
\newblock \emph{Nature}, \emph{394}, 72 -- 75.

\bibitem[{Shadlen et~al.(2013)Shadlen, \& Kiani}]{shadlen2013decision}
Shadlen, M. N., \& Kiani, R. (2013).
\newblock Decision making as a window on cognition. 
\newblock \emph{Neuron}, \emph{80}, 791 -- 806.

\bibitem[{Sirigu et~al.(2022)Sirigu, \& Duhamel}]{sirigu2022reward}
Sirigu, A., \& Duhamel, J. R. (2022).
\newblock Reward and decision processes in the brains of humans and nonhuman primates. 
\newblock \emph{Dialogues in Clinical Neuroscience} \emph{}.

\bibitem[{Suzuki et~al.(2021)Suzuki, Lawlor, Cooper, Arulpragasam, \& Treadway}]{suzuki2021distinct}
Suzuki, S., Lawlor, V. M., Cooper, J. A., Arulpragasam, A. R., \& Treadway, M. T. (2021).
\newblock Distinct regions of the striatum underlying effort, movement initiation and effort discounting. 
\newblock \emph{Nature Human Behaviour}, \emph{5}, 378 -- 388.

\bibitem[{Turan et~al.(2023)Turan, Baspinar, \& Destexhe}]{turan2023whole}
Turan, A., Baspinar, E., \& Destexhe, A. (2023).
\newblock A whole-brain model of auditory discrimination. 
\newblock \emph{bioRxiv} \emph{}.

\bibitem[{Van~Ede et~al.(2018)Van~Ede, Chekroud, Stokes \& Nobre}]{van2018decoding}
Van~Ede, F., Chekroud, S. R., Stokes, M. G., \& Nobre, A. C. (2018).
\newblock Decoding the influence of anticipatory states on visual perception in the presence of temporal distractors. 
\newblock \emph{Nature Communications}, \emph{9}, 1 -- 12.

\bibitem[{Wang(2018)Wang}]{wang2002probabilistic}
Wang, X. J. (2002).
\newblock Probabilistic decision making by slow reverberation in
  cortical circuits. 
\newblock \emph{Neuron}, \emph{36}, 955 -- 968.


\bibitem[{Wilson et~al.(1972)Wilson, \& Cowan}]{wilson1972excitatory}
Wilson, H. R., \& Cowan, J. D. (1972).
\newblock Excitatory and inhibitory interactions in localized populations of model neurons. 
\newblock \emph{Biophysical Journal}, \emph{12}, 1 -- 24.

\bibitem[{Xiao et~al.(2021)Xiao, Lin, \& Young}]{xiao2021data}
Xiao, Z. C., Lin, K. K., \& Young, L. S. (2021).
\newblock A data-informed mean-field approach to mapping of cortical parameter landscapes. 
\newblock \emph{PLoS Computational
  Biology}, \emph{17}, e1009718.
  
\bibitem[{Zerlaut et~al.(2016)Zerlaut, Tele{\'n}czuk, Deleuze, Bal, Ouanounou, \& Destexhe}]{zerlaut2016heterogeneous}
Zerlaut, Y., Tele{\'n}czuk, B., Deleuze, C., Bal, T., Ouanounou, G., \& Destexhe, A. (2016).
\newblock Heterogeneous firing rate response of mouse layer V pyramidal neurons in the fluctuation-driven regime. 
\newblock \emph{The Journal of Physiology}, \emph{13}, 3791 -- 3808.
  

\bibitem[{Zerlaut et~al.(2018)Zerlaut, Chemla, Chavane, \& Destexhe}]{zerlaut2018modeling}
Zerlaut, Y., Chemla, S., Chavan, F., \& Destexhe, A. (2018).
\newblock Modeling mesoscopic cortical dynamics using a mean-field model of conductance-based networks of adaptive exponential integrate-and-fire neurons. 
\newblock \emph{Journal of
  Computational Neuroscience}, \emph{44}, 45 -- 61.

\bibitem[{Zuo et~al.(2019)Zuo, \& Diamond}]{zuo2019rats}
Zuo, Y., \& Diamond, M. E. (2018).
\newblock Rats generate vibrissal sensory evidence until boundary crossing triggers a decision. 
\newblock \emph{Current Biology}, \emph{29}, 1415 -- 1424.


\end{thebibliography}


\end{document}
