## Supplementary material for "A biologically plausible decision-making model based on interacting neural populations": Tex files: references.bib

\begin{thebibliography}{1}


@article{di2019biologically,
  title={Biologically realistic mean-field models of conductance-based networks of spiking neurons with adaptation},
  author={Di Volo, Matteo and Romagnoni, Alberto and Capone, Cristiano and Destexhe, Alain},
  journal={Neural Computation},
  volume={31},
  number={4},
  pages={653--680},
  year={2019},
  publisher={MIT Press}
}

@article{el2009master,
  title={A master equation formalism for macroscopic modeling of asynchronous irregular activity states},
  author={El Boustani, Sami and Destexhe, Alain},
  journal={Neural Computation},
  volume={21},
  number={1},@article {10.7554/eLife.27756,
article_type = {journal},
title = {Sequential neuromodulation of Hebbian plasticity offers mechanism for effective reward-based navigation},
author = {Brzosko, Zuzanna and Zannone, Sara and Schultz, Wolfram and Clopath, Claudia and Paulsen, Ole},
editor = {Bartos, Marlene},
volume = 6,
year = 2017,
month = {jul},
pub_date = {2017-07-10},
pages = {e27756},
citation = {eLife 2017;6:e27756},
doi = {10.7554/eLife.27756},
url = {https://doi.org/10.7554/eLife.27756},
abstract = {Spike timing-dependent plasticity (STDP) is under neuromodulatory control, which is correlated with distinct behavioral states. Previously, we reported that dopamine, a reward signal, broadens the time window for synaptic potentiation and modulates the outcome of hippocampal STDP even when applied after the plasticity induction protocol (Brzosko et al., 2015). Here, we demonstrate that sequential neuromodulation of STDP by acetylcholine and dopamine offers an efficacious model of reward-based navigation. Specifically, our experimental data in mouse hippocampal slices show that acetylcholine biases STDP toward synaptic depression, whilst subsequent application of dopamine converts this depression into potentiation. Incorporating this bidirectional neuromodulation-enabled correlational synaptic learning rule into a computational model yields effective navigation toward changing reward locations, as in natural foraging behavior. Thus, temporally sequenced neuromodulation of STDP enables associations to be made between actions and outcomes and also provides a possible mechanism for aligning the time scales of cellular and behavioral learning.},
keywords = {synaptic plasticity, neuromodulation, hippocampus, reward, navigation},
journal = {eLife},
issn = {2050-084X},
publisher = {eLife Sciences Publications, Ltd},
}
  pages={46--100},
  year={2009},
  publisher={MIT Press}
}

@article{zerlaut2018modeling,
  title={Modeling mesoscopic cortical dynamics using a mean-field model of conductance-based networks of adaptive exponential integrate-and-fire neurons},
  author={Zerlaut, Yann and Chemla, Sandrine and Chavane, Frederic and Destexhe, Alain},
  journal={Journal of Computational Neuroscience},
  volume={44},
  number={1},
  pages={45--61},
  year={2018},
  publisher={Springer}
}

@article{brette2005adaptive,
  title={Adaptive exponential integrate-and-fire model as an effective description of neuronal activity},
  author={Brette, Romain and Gerstner, Wulfram},
  journal={Journal of Neurophysiology},
  volume={94},
  number={5},
  pages={3637--3642},
  year={2005},
  publisher={American Physiological Society}
}

@article{finkelstein2021attractor,
  title={Attractor dynamics gate cortical information flow during decision-making},
  author={Finkelstein, Arseny and Fontolan, Lorenzo and Economo, Michael N and Li, Nuo and Romani, Sandro and Svoboda, Karel},
  journal={Nature Neuroscience},
  pages={1--8},
  year={2021},
  publisher={Nature Publishing Group}
}

@article{seidemann1998temporal,
  title={Temporal gating of neural signals during performance of a visual discrimination task},
  author={Seidemann, Eyal and Zohary, Ehud and Newsome, William T},
  journal={Nature},
  volume={394},
  number={6688},
  pages={72--75},
  year={1998},
  publisher={Nature Publishing Group}
}

@article{kiani2008bounded,
  title={Bounded integration in parietal cortex underlies decisions even when viewing duration is dictated by the environment},
  author={Kiani, Roozbeh and Hanks, Timothy D and Shadlen, Michael N},
  journal={Journal of Neuroscience},
  volume={28},
  number={12},
  pages={3017--3029},
  year={2008},
  publisher={Soc Neuroscience}
}

@article{kopec2015cortical,
  title={Cortical and subcortical contributions to short-term memory for orienting movements},
  author={Kopec, Charles D and Erlich, Jeffrey C and Brunton, Bingni W and Deisseroth, Karl and Brody, Carlos D},
  journal={Neuron},
  volume={88},
  number={2},
  pages={367--377},
  year={2015},
  publisher={Elsevier}
}

@article{van2018decoding,
  title={Decoding the influence of anticipatory states on visual perception in the presence of temporal distractors},
  author={Van Ede, Freek and Chekroud, Sammi R and Stokes, Mark G and Nobre, Anna C},
  journal={Nature Communications},
  volume={9},
  number={1},
  pages={1--12},
  year={2018},
  publisher={Nature Publishing Group}
}

@article{zuo2019rats,
  title={Rats generate vibrissal sensory evidence until boundary crossing triggers a decision},
  author={Zuo, Yanfang and Diamond, Mathew E},
  journal={Current Biology},
  volume={29},
  number={9},
  pages={1415--1424},
  year={2019},
  publisher={Elsevier}
}

@article{parker1998sense,
  title={Sense and the single neuron: probing the physiology of perception},
  author={Parker, Andrew J and Newsome, William T},
  journal={Annual Review of Neuroscience},
  volume={21},
  number={1},
  pages={227--277},
  year={1998},
  publisher={Annual Reviews 4139 El Camino Way, PO Box 10139, Palo Alto, CA 94303-0139, USA}
}

@article{romo2001touch,
  title={Touch and go: decision-making mechanisms in somatosensation},
  author={Romo, Ranulfo and Salinas, Emilio},
  journal={Annual Review of Neuroscience},
  volume={24},
  number={1},
  pages={107--137},
  year={2001},
  publisher={Annual Reviews 4139 El Camino Way, PO Box 10139, Palo Alto, CA 94303-0139, USA}
}

@article{schall2001neural,
  title={Neural basis of deciding, choosing and acting},
  author={Schall, Jeffrey D},
  journal={Nature Reviews Neuroscience},
  volume={2},
  number={1},
  pages={33--42},
  year={2001},
  publisher={Nature Publishing Group}
}

@article{broche2016neural,
  title={Neural substrates of decision-making},
  author={Broche-P{\'e}rez, Y and Jim{\'e}nez, LF Herrera and Omar-Mart{\'\i}nez, E},
  journal={Neurolog{\'\i}a (English Edition)},
  volume={31},
  number={5},
  pages={319--325},
  year={2016},
  publisher={Elsevier}
}

@article{paulus2003increased,
  title={Increased activation in the right insula during risk-taking decision making is related to harm avoidance and neuroticism},
  author={Paulus, Martin P and Rogalsky, Corianne and Simmons, Alan and Feinstein, Justin S and Stein, Murray B},
  journal={Neuroimage},
  volume={19},
  number={4},
  pages={1439--1448},
  year={2003},
  publisher={Elsevier}
}

@article{sanfey2006neuroeconomics,
  title={Neuroeconomics: cross-currents in research on decision-making},
  author={Sanfey, Alan G and Loewenstein, George and McClure, Samuel M and Cohen, Jonathan D},
  journal={Trends in Cognitive Sciences},
  volume={10},
  number={3},
  pages={108--116},
  year={2006},
  publisher={Elsevier}
}

@article{guo2013brain,
  title={Brain areas activated by uncertain reward-based decision-making in healthy volunteers},
  author={Guo, Zongjun and Chen, Juan and Liu, Shien and Li, Yuhuan and Sun, Bo and Gao, Zhenbo},
  journal={Neural Regeneration Research},
  volume={8},
  number={35},
  pages={3344},
  year={2013},
  publisher={Wolters Kluwer--Medknow Publications}
}

@article{albantakis2011changes,
  title={Changes of mind in an attractor network of decision-making},
  author={Albantakis, Larissa and Deco, Gustavo},
  journal={PLoS Comput Biol},
  volume={7},
  number={6},
  pages={e1002086},
  year={2011},
  publisher={Public Library of Science}
}

@article{moreno2007noise,
  title={Noise-induced alternations in an attractor network model of perceptual bistability},
  author={Moreno-Bote, Rub{\'e}n and Rinzel, John and Rubin, Nava},
  journal={Journal of Neurophysiology},
  volume={98},
  number={3},
  pages={1125--1139},
  year={2007},
  publisher={American Physiological Society}
}

@article{hayden2018neuronal,
  title={A neuronal theory of sequential economic choice},
  author={Hayden, Benjamin Y and Moreno-Bote, Rub{\'e}n},
  journal={Brain and Neuroscience Advances},
  volume={2},
  pages={2398212818766675},
  year={2018},
  publisher={SAGE Publications Sage UK: London, England}
}


@article{brunel2001effects,
  title={Effects of neuromodulation in a cortical network model of object working memory dominated by recurrent inhibition},
  author={Brunel, Nicolas and Wang, Xiao-Jing},
  journal={Journal of Computational Neuroscience},
  volume={11},
  number={1},
  pages={63--85},
  year={2001},
  publisher={Springer}
}

@article{wang2002probabilistic,
  title={Probabilistic decision making by slow reverberation in cortical circuits},
  author={Wang, Xiao-Jing},
  journal={Neuron},
  volume={36},
  number={5},
  pages={955--968},
  year={2002},
  publisher={Elsevier}
}


@article{marcos2013neural,
  title={Neural variability in premotor cortex is modulated by trial history and predicts behavioral performance},
  author={Marcos, Encarni and Pani, Pierpaolo and Brunamonti, Emiliano and Deco, Gustavo and Ferraina, Stefano and Verschure, Paul},
  journal={Neuron},
  volume={78},
  number={2},
  pages={249--255},
  year={2013},
  publisher={Elsevier}
}


@article{gilbert1979morphology,
  title={Morphology and intracortical projections of functionally characterised neurones in the cat visual cortex},
  author={Gilbert, Charles D and Wiesel, Torsten N},
  journal={Nature},
  volume={280},
  number={5718},
  pages={120--125},
  year={1979},
  publisher={Nature Publishing Group}
}

@article{rockland1982widespread,
  title={Widespread periodic intrinsic connections in the tree shrew visual cortex},
  author={Rockland, Kathleen S and Lund, Jennifer S},
  journal={Science},
  volume={215},
  number={4539},
  pages={1532--1534},
  year={1982},
  publisher={American Association for the Advancement of Science}
}

@article{livingstone1984specificity,
  title={Specificity of intrinsic connections in primate primary visual cortex},
  author={Livingstone, MARGARET S and Hubel, DAVID H},
  journal={Journal of Neuroscience},
  volume={4},
  number={11},
  pages={2830--2835},
  year={1984},
  publisher={Soc Neuroscience}
}

@article{martin1984form,
  title={Form, function and intracortical projections of spiny neurones in the striate visual cortex of the cat.},
  author={Martin, KA and Whitteridge, David},
  journal={The Journal of Physiology},
  volume={353},
  number={1},
  pages={463--504},
  year={1984},
  publisher={Wiley Online Library}
}

@article{kisvarday1986synaptic,
  title={Synaptic targets of HRP-filled layer III pyramidal cells in the cat striate cortex},
  author={Kisv{\'a}rday, Zolt{\'a}n F and Martin, Kevan AC and Freund, TF and Magloczky, ZS and Whitteridge, D and Somogyi, P},
  journal={Experimental Brain Research},
  volume={64},
  number={3},
  pages={541--552},
  year={1986},
  publisher={Springer}
}

@article{mcguire1991targets,
  title={Targets of horizontal connections in macaque primary visual cortex},
  author={McGuire, Barbara A and Gilbert, Charles D and Rivlin, Patricia K and Wiesel, Torsten N},
  journal={Journal of Comparative Neurology},
  volume={305},
  number={3},
  pages={370--392},
  year={1991},
  publisher={Wiley Online Library}
}

@article{albantakis2009encoding,
  title={The encoding of alternatives in multiple-choice decision making},
  author={Albantakis, Larissa and Deco, Gustavo},
  journal={Proceedings of the National Academy of Sciences},
  volume={106},
  number={25},
  pages={10308--10313},
  year={2009},
  publisher={National Acad Sciences}
}

@article{churchland2008decision,
  title={Decision-making with multiple alternatives},
  author={Churchland, Anne K and Kiani, Roozbeh and Shadlen, Michael N},
  journal={Nature Neuroscience},
  volume={11},
  number={6},
  pages={693--702},
  year={2008},
  publisher={Nature Publishing Group}
}

@article{arnold1974stochastic,
  title={Stochastic differential equations},
  author={Arnold, Ludwig},
  journal={New York},
  year={1974}
}


%@article{goldman2023comprehensive,
%  title={A comprehensive neural simulation of slow-wave sleep and highly responsive wakefulness dynamics},
%  author={Goldman, Jennifer S and Kusch, Lionel and Aquilue, David and Yalcinkaya, Bahar Hazal and Depannemaecker, Damien and Ancourt, Kevin and Nghiem, Trang-Anh E and Jirsa, Viktor and Destexhe, Alain},
%  journal={Frontiers in Computational Neuroscience},
%  volume={16},
%  pages={1058957},
%  year={2023},
%  publisher={Frontiers Media SA}
%}


@article{goldman2020brain,
  title={Brain-scale emergence of slow-wave synchrony and highly responsive asynchronous states based on biologically realistic population models simulated in The Virtual Brain},
  author={Goldman, Jennifer S and Kusch, Lionel and Yalcinkaya, Bahar Hazal and Depannemaecker, Damien and Nghiem, Trang-Anh E and Jirsa, Viktor and Destexhe, Alain},
  journal={bioRxiv},
  year={2020},
  publisher={Cold Spring Harbor Laboratory}
}

@article{drugowitsch2012cost,
  title={The cost of accumulating evidence in perceptual decision making},
  author={Drugowitsch, Jan and Moreno-Bote, Rub{\'e}n and Churchland, Anne K and Shadlen, Michael N and Pouget, Alexandre},
  journal={Journal of Neuroscience},
  volume={32},
  number={11},
  pages={3612--3628},
  year={2012},
  publisher={Soc Neuroscience}
}

@article{mochol2021prefrontal,
  title={Prefrontal cortex represents heuristics that shape choice bias and its integration into future behavior},
  author={Mochol, Gabriela and Kiani, Roozbeh and Moreno-Bote, Rub{\'e}n},
  journal={Current Biology},
  volume={31},
  number={6},
  pages={1234--1244},
  year={2021},
  publisher={Elsevier}
}

@article{shadlen2013decision,
  title={Decision making as a window on cognition},
  author={Shadlen, Michael N and Kiani, Roozbeh},
  journal={Neuron},
  volume={80},
  number={3},
  pages={791--806},
  year={2013},
  publisher={Elsevier}
}

@article{rushworth2011frontal,
  title={Frontal cortex and reward-guided learning and decision-making},
  author={Rushworth, Matthew FS and Noonan, MaryAnn P and Boorman, Erie D and Walton, Mark E and Behrens, Timothy E},
  journal={Neuron},
  volume={70},
  number={6},
  pages={1054--1069},
  year={2011},
  publisher={Elsevier}
}

@article{domenech2015executive,
  title={Executive control and decision-making in the prefrontal cortex},
  author={Domenech, Philippe and Koechlin, Etienne},
  journal={Current Opinion in Behavioral Sciences},
  volume={1},
  pages={101--106},
  year={2015},
  publisher={Elsevier}
}

@article{bechara2000characterization,
  title={Characterization of the decision-making deficit of patients with ventromedial prefrontal cortex lesions},
  author={Bechara, Antoine and Tranel, Daniel and Damasio, Hanna},
  journal={Brain},
  volume={123},
  number={11},
  pages={2189--2202},
  year={2000},
  publisher={Oxford University Press}
}

@article{hampton2006role,
  title={The role of the ventromedial prefrontal cortex in abstract state-based inference during decision making in humans},
  author={Hampton, Alan N and Bossaerts, Peter and O’doherty, John P},
  journal={Journal of Neuroscience},
  volume={26},
  number={32},
  pages={8360--8367},
  year={2006},
  publisher={Soc Neuroscience}
}

@article{sirigu2022reward,
  title={Reward and decision processes in the brains of humans and nonhuman primates},
  author={Sirigu, Angela and Duhamel, Jean-Ren{\'e}},
  journal={Dialogues in Clinical Neuroscience},
  year={2022},
  publisher={Taylor \& Francis}
}

@article{monosov2012regionally,
  title={Regionally distinct processing of rewards and punishments by the primate ventromedial prefrontal cortex},
  author={Monosov, Ilya E and Hikosaka, Okihide},
  journal={Journal of Neuroscience},
  volume={32},
  number={30},
  pages={10318--10330},
  year={2012},
  publisher={Soc Neuroscience}
}

@article{suzuki2021distinct,
  title={Distinct regions of the striatum underlying effort, movement initiation and effort discounting},
  author={Suzuki, Shosuke and Lawlor, Victoria M and Cooper, Jessica A and Arulpragasam, Amanda R and Treadway, Michael T},
  journal={Nature human behaviour},
  volume={5},
  number={3},
  pages={378--388},
  year={2021},
  publisher={Nature Publishing Group UK London}
}

@article{xiao2021data,
  title={A data-informed mean-field approach to mapping of cortical parameter landscapes},
  author={Xiao, Zhuo-Cheng and Lin, Kevin K and Young, Lai-Sang},
  journal={PLoS Computational Biology},
  volume={17},
  number={12},
  pages={e1009718},
  year={2021},
  publisher={Public Library of Science San Francisco, CA USA}
}

@article{wilson1972excitatory,
  title={Excitatory and inhibitory interactions in localized populations of model neurons},
  author={Wilson, Hugh R and Cowan, Jack D},
  journal={Biophysical Journal},
  volume={12},
  number={1},
  pages={1--24},
  year={1972},
  publisher={Elsevier}
}

@article{amari1977dynamics,
  title={Dynamics of pattern formation in lateral-inhibition type neural fields},
  author={Amari, Shun-ichi},
  journal={Biological Cybernetics},
  volume={27},
  number={2},
  pages={77--87},
  year={1977},
  publisher={Springer}
}


@book{doya2007bayesian,
  title={Bayesian brain: Probabilistic approaches to neural coding},
  author={Doya, Kenji and Ishii, Shin and Pouget, Alexandre and Rao, Rajesh PN},
  year={2007},
  publisher={MIT press}
}

@article{giamundo2021neuronal,
  title={Neuronal activity in the premotor cortex of monkeys reflects both cue salience and motivation for action generation and inhibition},
  author={Giamundo, Margherita and Giarrocco, Franco and Brunamonti, Emiliano and Fabbrini, Francesco and Pani, Pierpaolo and Ferraina, Stefano},
  journal={Journal of Neuroscience},
  volume={41},
  number={36},
  pages={7591--7606},
  year={2021},
  publisher={Soc Neuroscience}
}

@article{leotti2010motivational,
  title={Motivational influences on response inhibition measures.},
  author={Leotti, Lauren A and Wager, Tor D},
  journal={Journal of Experimental Psychology: Human Perception and Performance},
  volume={36},
  number={2},
  pages={430},
  year={2010},
  publisher={American Psychological Association}
}

@article{padmala2010interactions,
  title={Interactions between cognition and motivation during response inhibition},
  author={Padmala, Srikanth and Pessoa, Luiz},
  journal={Neuropsychologia},
  volume={48},
  number={2},
  pages={558--565},
  year={2010},
  publisher={Elsevier}
}

@software{baspinarJupyter2023,
  author       = {Emre Baspinar and
                  Gloria Cecchini and
                  Ruben Moreno-Bote and
                  Ignasi Cos and
                  Alain Destexhe},
  title        = {{Jupyter notebook of a biophysically plausible 
                   decision-making model based on interacting
                   cortical columns}},
  month        = {\textit{Zenodo},},
  year         = 2023,
  publisher    = {Zenodo},
  version      = {1.0.0},
  doi          = {10.5281/zenodo.7682309},
  url          = {https://doi.org/10.5281/zenodo.7682309},
  journal       = {Zenodo}
}

@software{baspinarJupyterEbrains2023,
  author       = {Emre Baspinar and
                  Gloria Cecchini and
                  Ruben Moreno-Bote and
                  Ignasi Cos and
                  Alain Destexhe},
  title        = {{Double columnar Adaptive Exponential mean-field model for decision-making }},
  month        = {\textit{EBRAINS},},
  year         = 2023,
  publisher    = {EBRAINS},
  version      = {1.0.0},
  doi          = {},
  url          = {https://search.kg.ebrains.eu/instances/a0cb1207-b6a8-44e9-9cd0-7f98acce2080},
  journal       = {EBRAINS}
}

@article {cecchiniPreprint,
	author = {Gloria Cecchini and Michael DePass and Emre Baspinar and Marta Andujar and Surabhi Ramawat and Pierpaolo Pani and Stefano Ferraina and Alain Destexhe and Rub{\'e}n Moreno-Bote and Ignasi Cos},
	title = {Consequence assessment and behavioral patterns of inhibition in decision-making: modelling its underlying mechanisms},
	elocation-id = {2023.02.14.528595},
	year = {2023},
	doi = {10.1101/2023.02.14.528595},
	publisher = {Cold Spring Harbor Laboratory},
	abstract = {Learning to make adaptive decisions depends on exploring options, experiencing their consequence, and reassessing one{\textquoteright}s strategy for the future. Although several studies have analyzed various aspects of value-based decision-making, most of them have focused on decisions in which gratification is cued and immediate. By contrast, how the brain gauges delayed consequence for decision-making remains poorly understood.To investigate this, we designed a decision-making task in which each decision altered future options. The task was organized in groups of consecutively dependent trials, and the participants were instructed to maximize the cumulative reward value within each group. In the absence of any explicit performance feedback, the participants had to test and internally assess specific criteria to make decisions. This task was designed to specifically study how the assessment of consequence forms and influences decisions as learning progresses. We analyzed behavior results to characterize individual differences in reaction times, decision strategies, and learning rates.We formalized this operation mathematically by means of a multi-layered decision-making model. By using a mean-field approximation, the first layer of the model described the dynamics of two populations of neurons which characterized the binary decision-making process. The other two layers modulated the decision-making policy by dynamically adapting an oversight learning mechanism. The model was validated by fitting each individual participants{\textquoteright} behavior and it faithfully predicted non-trivial patterns of decision-making, regardless of performance level.These findings provided an explanation to how delayed consequence may be computed and incorporated into the neural dynamics of decision-making, and to how learning occurs in the absence of explicit feedback.Competing Interest StatementThe authors have declared no competing interest.},
	URL = {https://www.biorxiv.org/content/early/2023/07/05/2023.02.14.528595},
	eprint = {https://www.biorxiv.org/content/early/2023/07/05/2023.02.14.528595.full.pdf},
	journal = {bioRxiv}
}


@article{funahashi2017prefrontal,
  title={Prefrontal contribution to decision-making under free-choice conditions},
  author={Funahashi, Shintaro},
  journal={Frontiers in neuroscience},
  volume={11},
  pages={431},
  year={2017},
  publisher={Frontiers Media SA}
}


@article{kennerley2011decision,
  title={Decision making and reward in frontal cortex: complementary evidence from neurophysiological and neuropsychological studies.},
  author={Kennerley, Steven W and Walton, Mark E},
  journal={Behavioral neuroscience},
  volume={125},
  number={3},
  pages={297},
  year={2011},
  publisher={American Psychological Association}
}

@article{schultz2022dopamine,
  title={Dopamine reward prediction error coding},
  author={Schultz, Wolfram},
  journal={Dialogues in clinical neuroscience},
  year={2022},
  publisher={Taylor \& Francis}
}

@article{mirenowicz1994importance,
  title={Importance of unpredictability for reward responses in primate dopamine neurons},
  author={Mirenowicz, Jacques and Schultz, Wolfram},
  journal={Journal of neurophysiology},
  volume={72},
  number={2},
  pages={1024--1027},
  year={1994}
}

@article{sanz2013virtual,
  title={The Virtual Brain: a simulator of primate brain network dynamics},
  author={Sanz Leon, Paula and Knock, Stuart A and Woodman, M Marmaduke and Domide, Lia and Mersmann, Jochen and McIntosh, Anthony R and Jirsa, Viktor},
  journal={Frontiers in neuroinformatics},
  volume={7},
  pages={10},
  year={2013},
  publisher={Frontiers Media SA}
}


@article {brzosko2015,
article_type = {journal},
title = {Retroactive modulation of spike timing-dependent plasticity by dopamine},
author = {Brzosko, Zuzanna and Schultz, Wolfram and Paulsen, Ole},
editor = {Bartos, Marlene},
volume = 4,
year = 2015,
month = {oct},
pub_date = {2015-10-30},
pages = {e09685},
citation = {eLife 2015;4:e09685},
doi = {10.7554/eLife.09685},
url = {https://doi.org/10.7554/eLife.09685},
abstract = {Most reinforcement learning models assume that the reward signal arrives after the activity that led to the reward, placing constraints on the possible underlying cellular mechanisms. Here we show that dopamine, a positive reinforcement signal, can retroactively convert hippocampal timing-dependent synaptic depression into potentiation. This effect requires functional NMDA receptors and is mediated in part through the activation of the cAMP/PKA cascade. Collectively, our results support the idea that reward-related signaling can act on a pre-established synaptic eligibility trace, thereby associating specific experiences with behaviorally distant, rewarding outcomes. This finding identifies a biologically plausible mechanism for solving the ‘distal reward problem’.},
keywords = {synaptic plasticity, dopamine, reward},
journal = {eLife},
issn = {2050-084X},
publisher = {eLife Sciences Publications, Ltd},
}


@article {brzosko2017,
article_type = {journal},
title = {Sequential neuromodulation of Hebbian plasticity offers mechanism for effective reward-based navigation},
author = {Brzosko, Zuzanna and Zannone, Sara and Schultz, Wolfram and Clopath, Claudia and Paulsen, Ole},
editor = {Bartos, Marlene},
volume = 6,
year = 2017,
month = {jul},
pub_date = {2017-07-10},
pages = {e27756},
citation = {eLife 2017;6:e27756},
doi = {10.7554/eLife.27756},
url = {https://doi.org/10.7554/eLife.27756},
keywords = {synaptic plasticity, neuromodulation, hippocampus, reward, navigation},
journal = {eLife},
issn = {2050-084X},
publisher = {eLife Sciences Publications, Ltd},
}

@article{turan2023whole,
  title={A whole-brain model of auditory discrimination},
  author={Turan, Abdullah and Baspinar, Emre and Destexhe, Alain},
  journal={bioRxiv},
  pages={2023--09},
  year={2023},
  publisher={Cold Spring Harbor Laboratory}
}

@article{hirata2008functional,
  title={Functional columns in the primate prefrontal cortex revealed by optical imaging in vitro},
  author={Hirata, Yoshihiro and Sawaguchi, Toshiyuki},
  journal={Neuroscience research},
  volume={61},
  number={1},
  pages={1--10},
  year={2008},
  publisher={Elsevier}
}

@misc{humanEbrains,
  doi = {10.25493/A6JP-E63},
  url = {https://search.kg.ebrains.eu/instances/0d145ebe-3ecd-4b3c-9400-913a8cd21a6a},
  author = {DePass, Michael and Cecchini, Gloria and Cos, Ignasi},
  keywords = {Neuroscience},
  title = {Consequence assessment and behavioral patterns of inhibition in decision-making (v1)},
  publisher = {EBRAINS},
  year = {EBRAINS dataset, 2023},
  copyright = {Creative Commons Attribution Non Commercial Share Alike 4.0 International}
}

@misc{macaqueEbrains,
  doi = {10.25493/Z7YX-7W2},
  url = {https://search.kg.ebrains.eu/instances/755b98d5-3212-4220-950e-54d58e809e6e},
  author = {Fontana, Roberto and Andujar, Marta and Marc, Isabel Beatrice and Giuffrida, Valentina and Ramawat, Surabhi and Bardella, Giampiero and Brunamonti, Emiliano and Pani, Pierpaolo and Ferraina, Stefano},
  keywords = {Neuroscience},
  title = {Evaluating consequences in decision-making: behavioral data and neural recordings from monkey premotor and prefrontal cortex (v1)},
  publisher = {EBRAINS},
  year = {EBRAINS dataset, 2022},
  copyright = {Creative Commons Attribution Non Commercial Share Alike 4.0 International}
}

@book{edition2002probability,
  title={Probability, random variables, and stochastic processes},
  author={Edition, Fourth and Papoulis, Athanasios and Pillai, S Unnikrishna},
  year={2002},
  publisher={McGraw-Hill Europe: New York, NY, USA}
}


\end{thebibliography}
