## Supplementary figures and images for "A biologically plausible decision-making model based on interacting neural populations"

### awakeToSleep.png

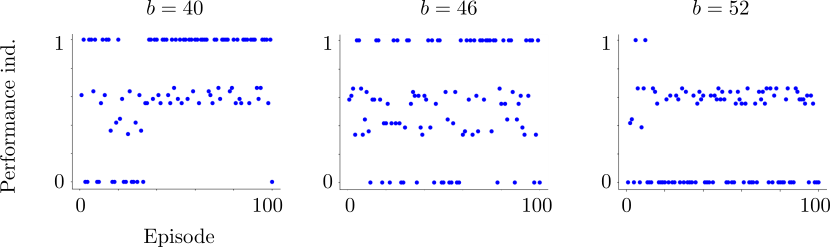

### basicModule.pdf

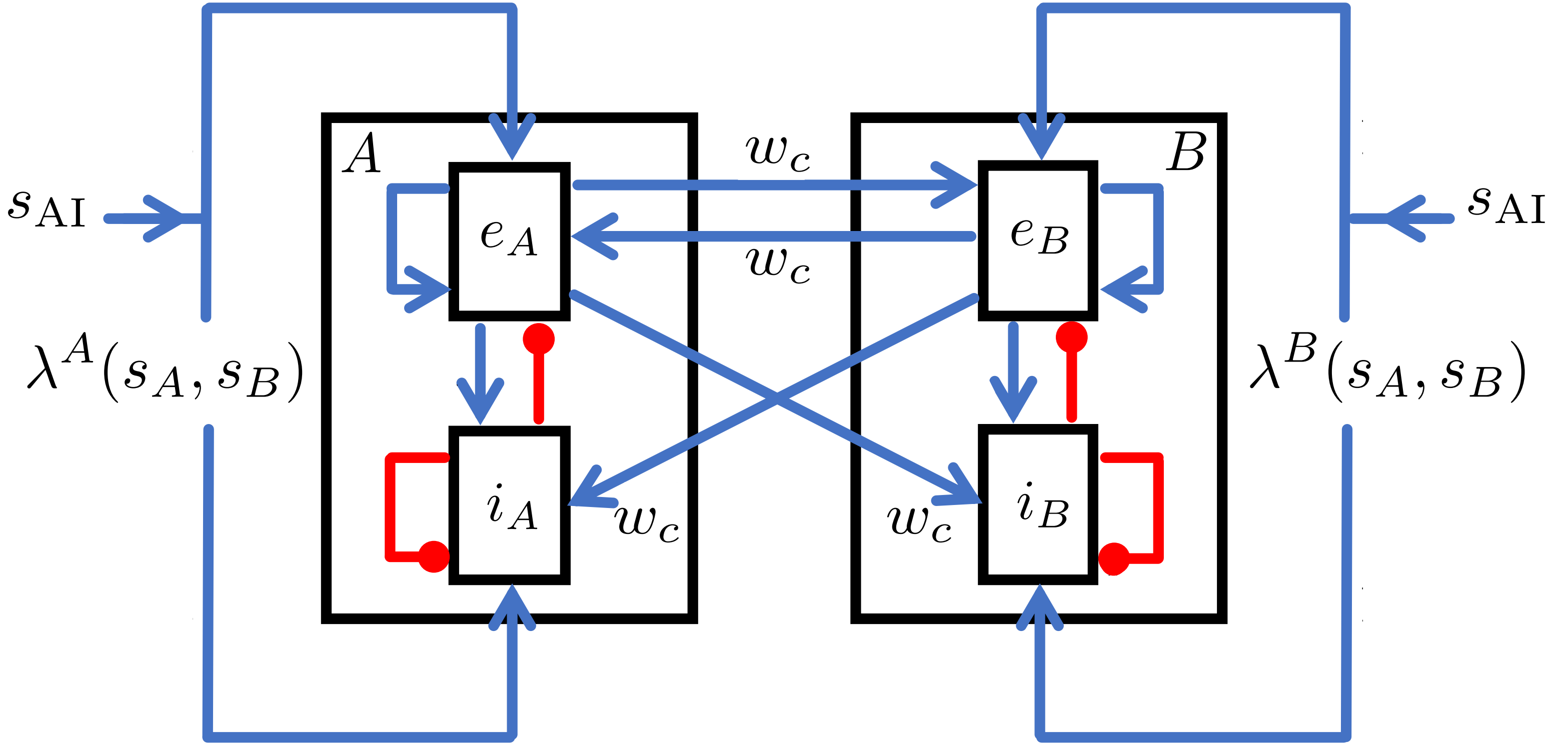

### c0EffectV.png

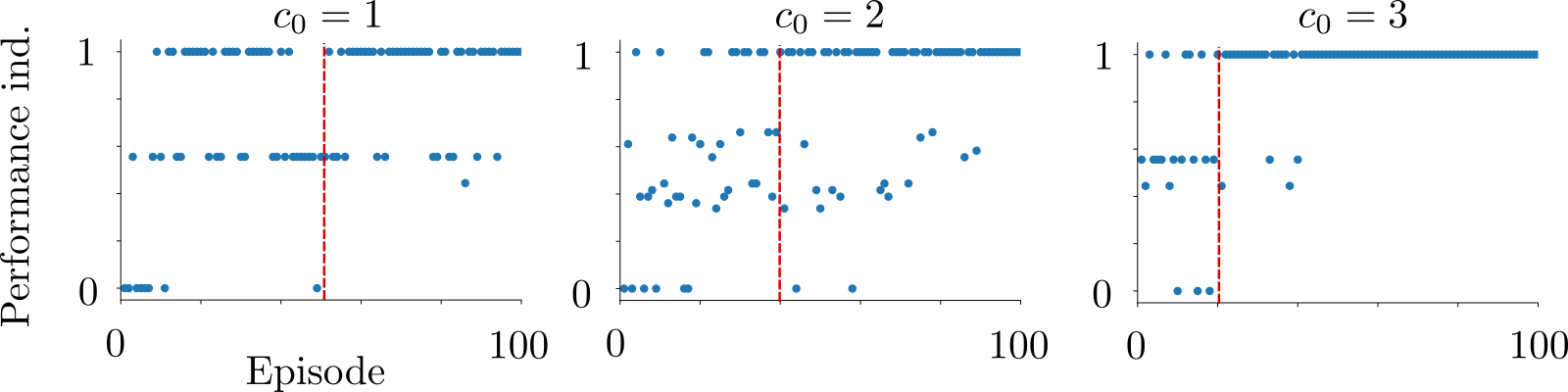

### differentA_B_Stimuli.png

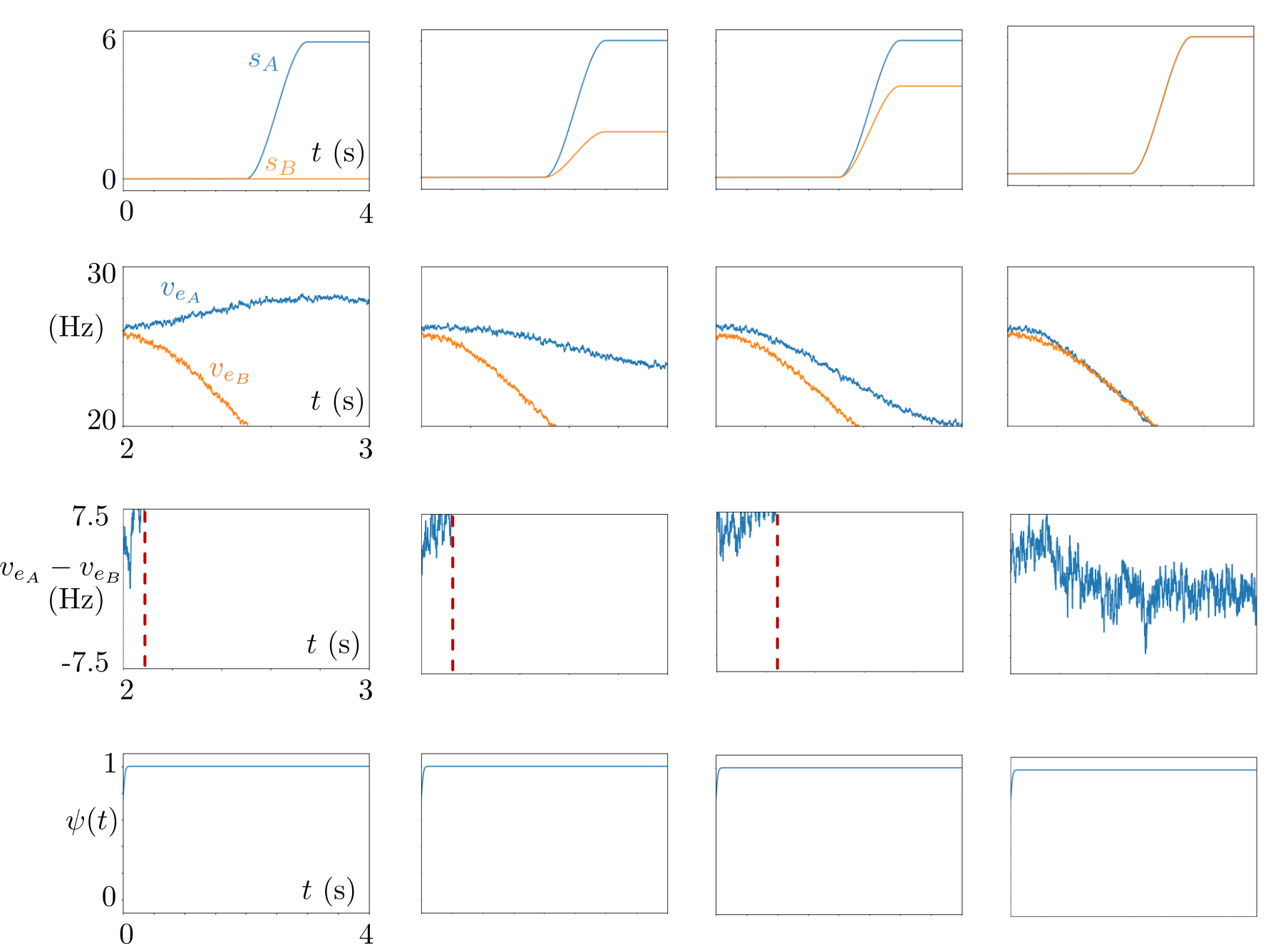

### energyFunctionalConverge.png

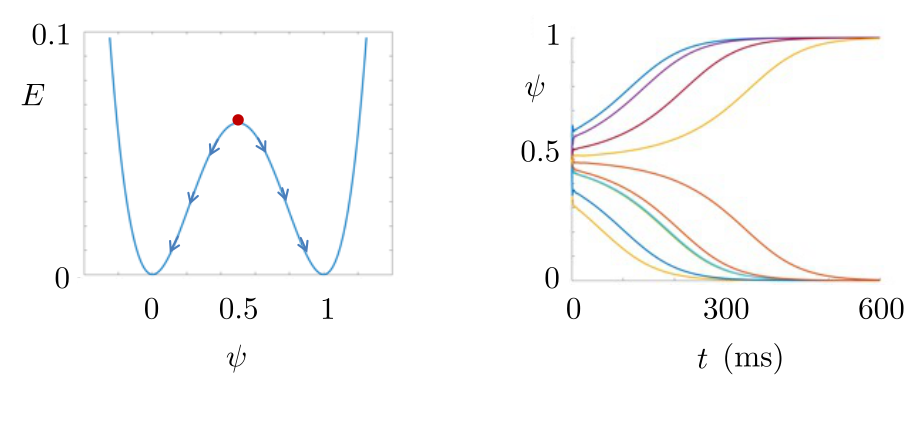

### examplePsiEffect.png

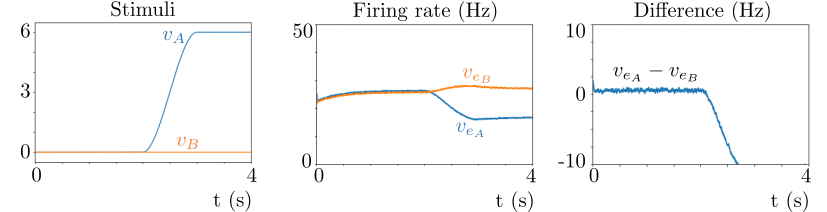

### gainEffect.png

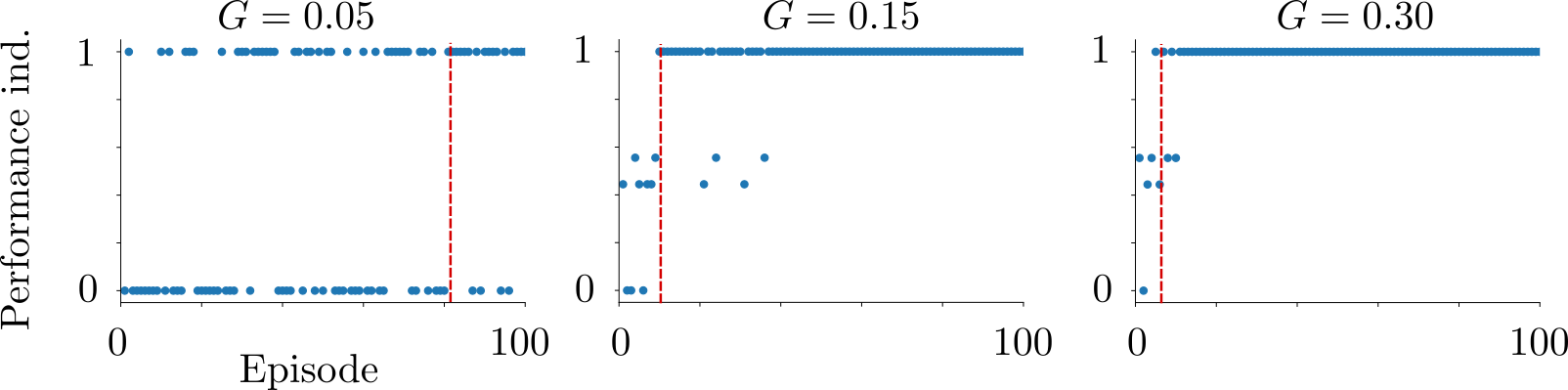

### globalMeasures.png

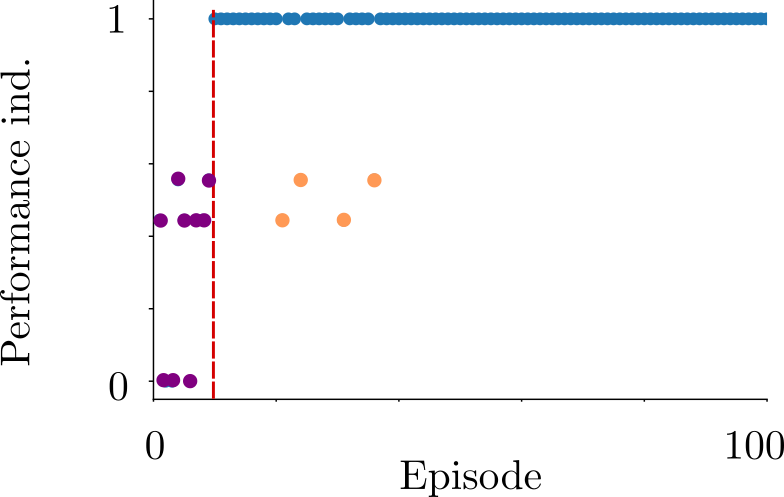

### H0_RT_Histogram.png

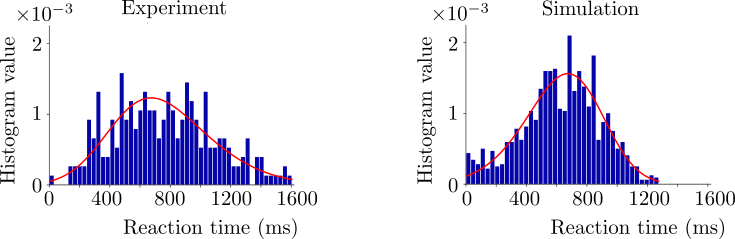

### H1_Monkey_RT_Histogram.png

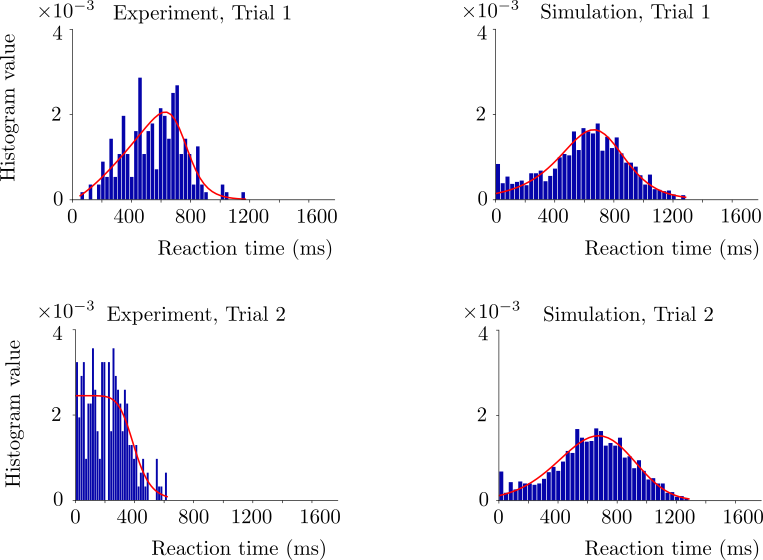

### H1_RT_Histogram.png

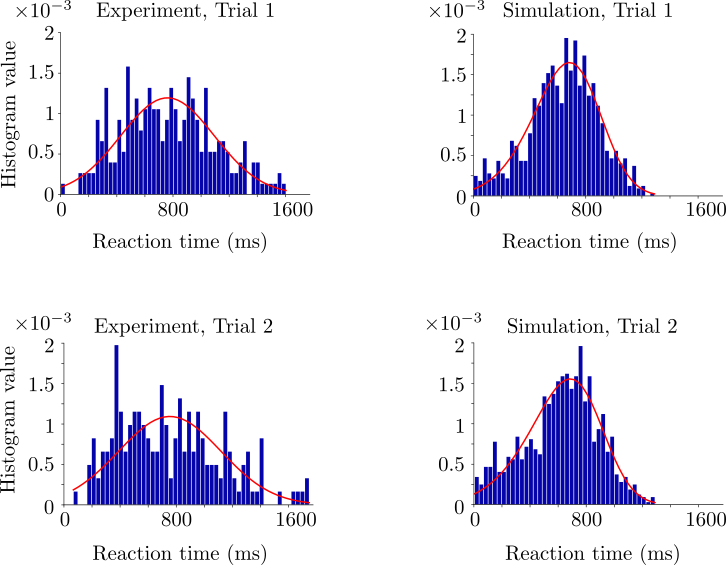

### humanExpScenario.png

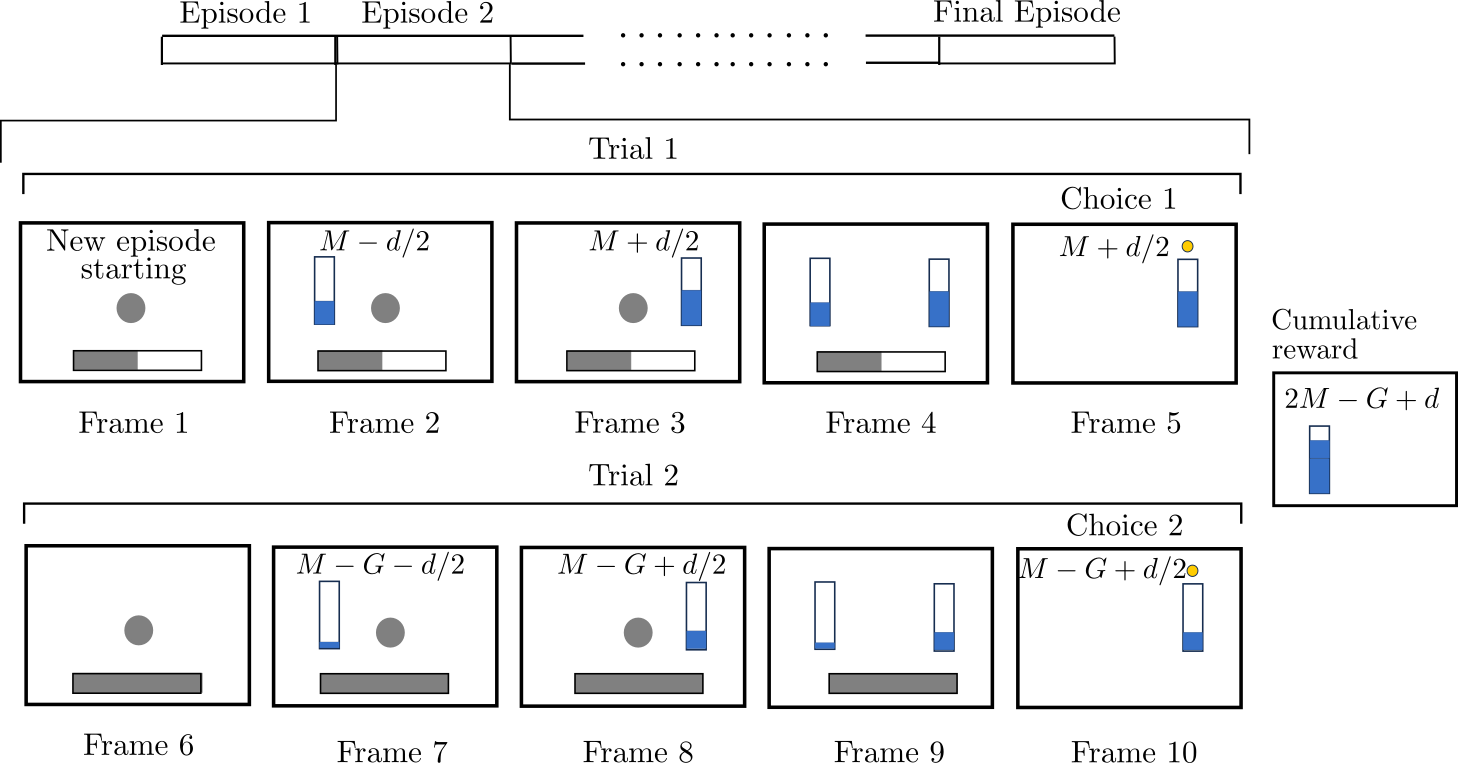

### learningSpeedEffect.png

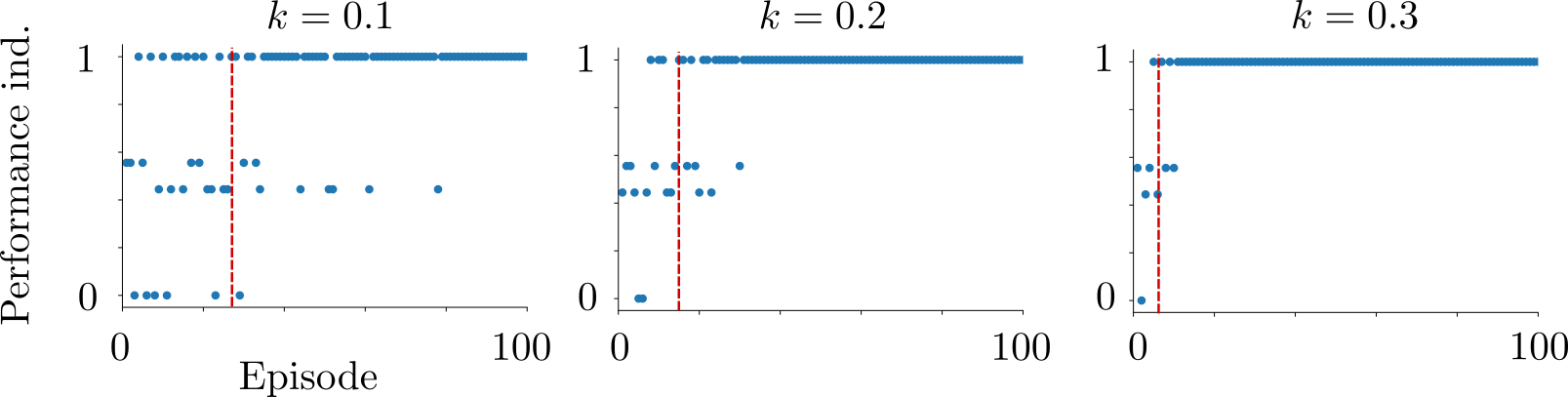

### learningTime.png

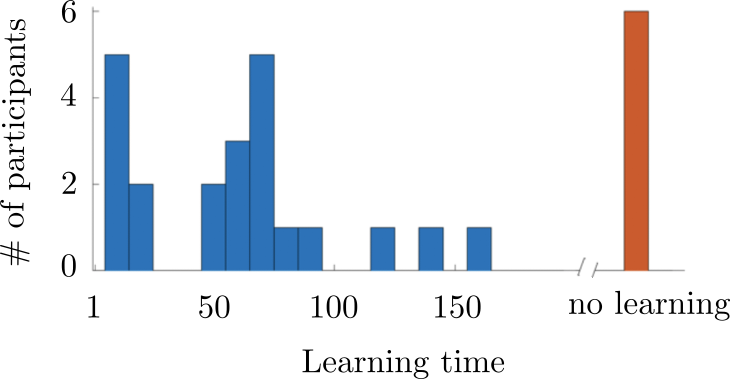

### monkeyExpScenario.png

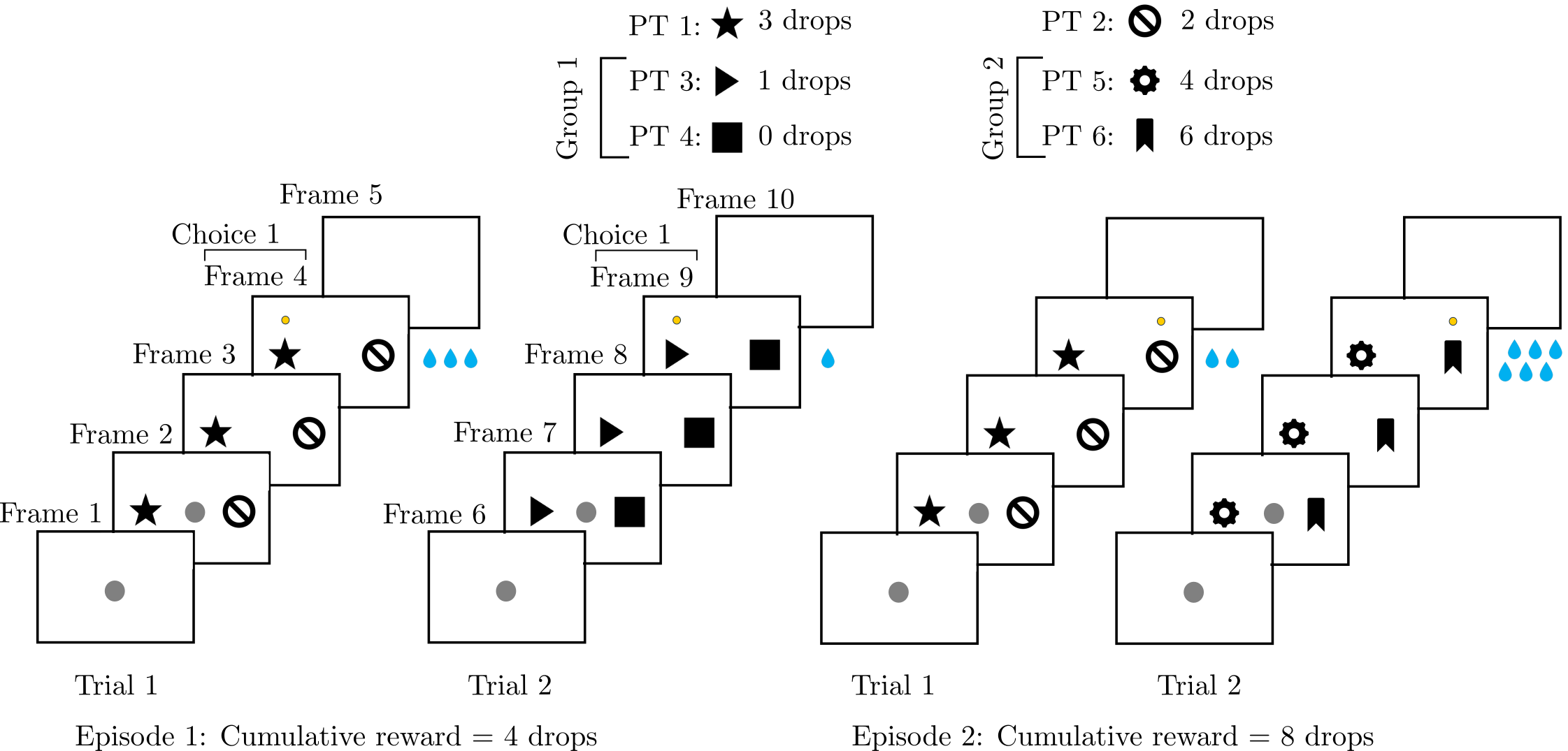

### performanceCDF.png

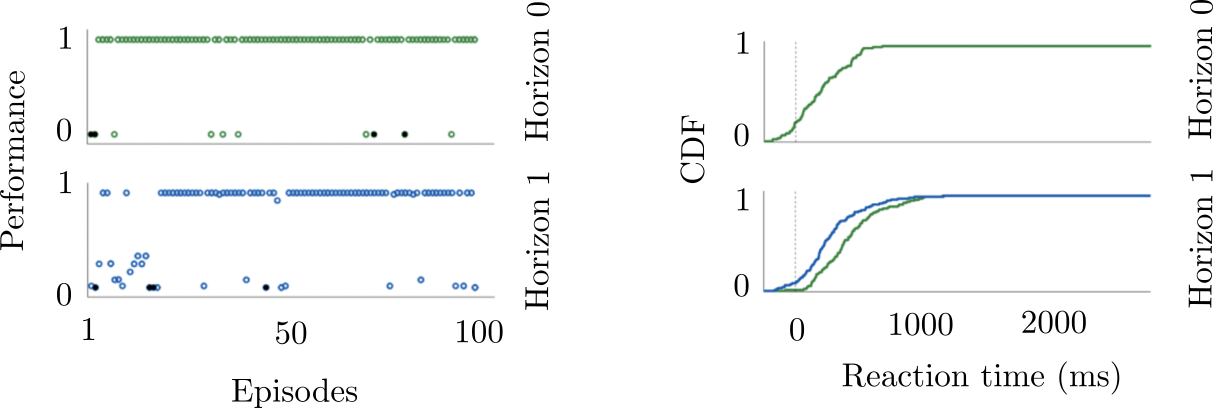

### PsiEffectFullSys.png

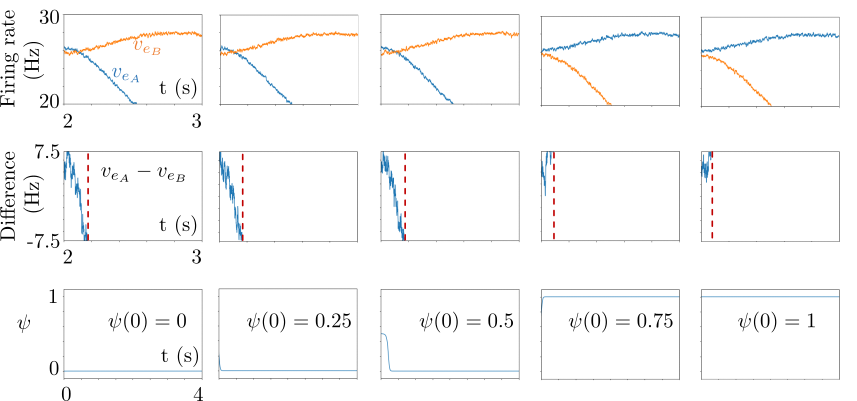

### RT_H0.png

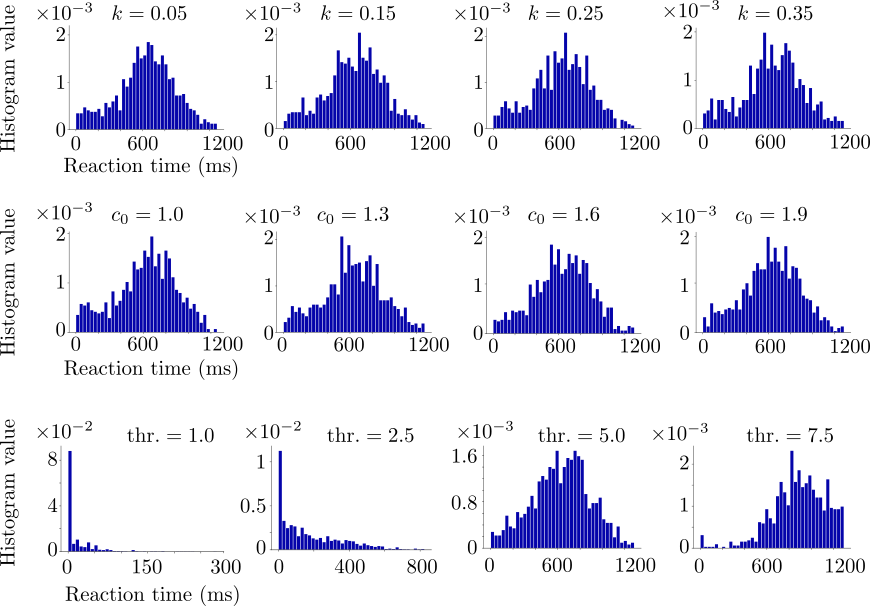

### RT_H1_T1_T2.png

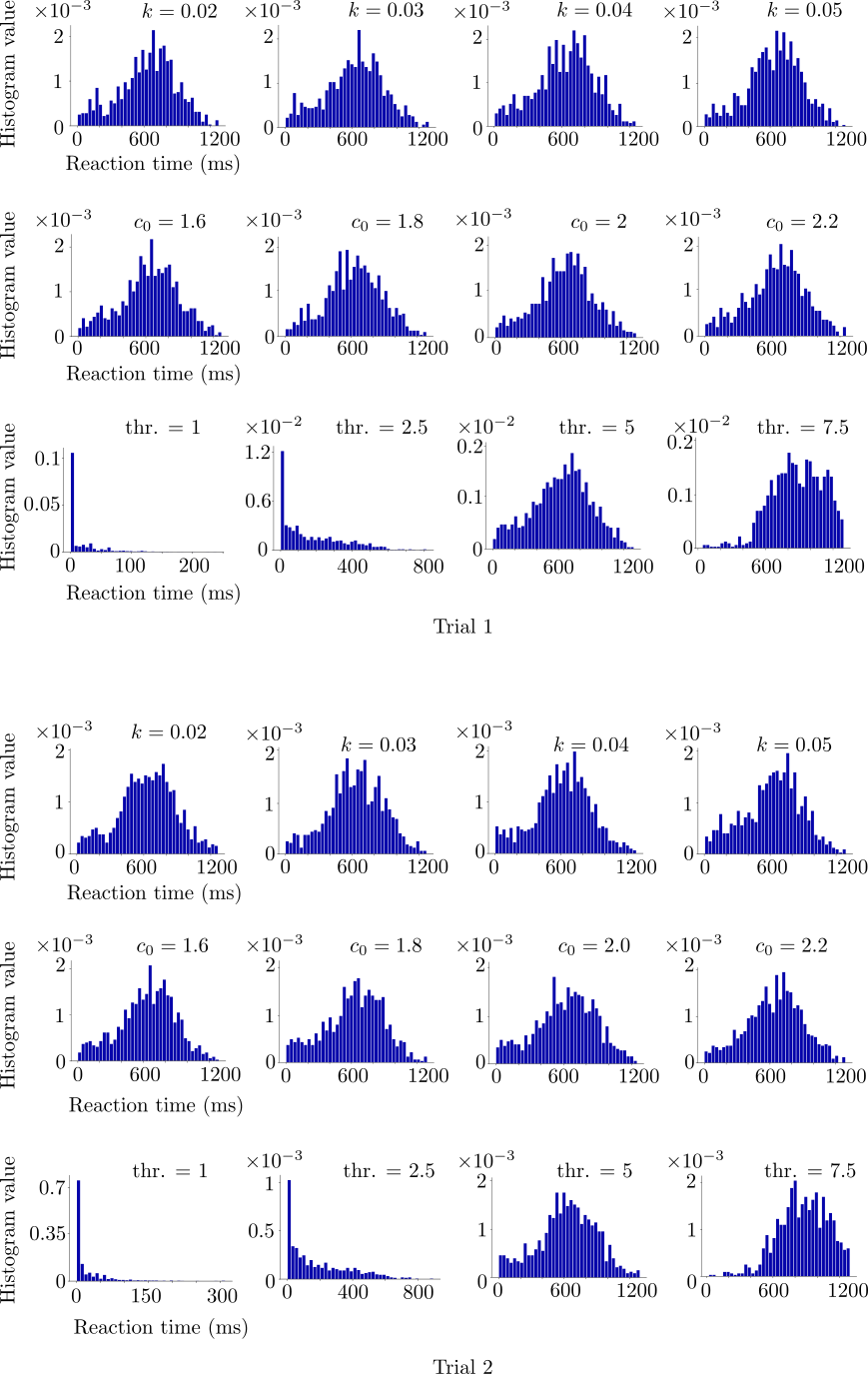

### statisticalH0.png

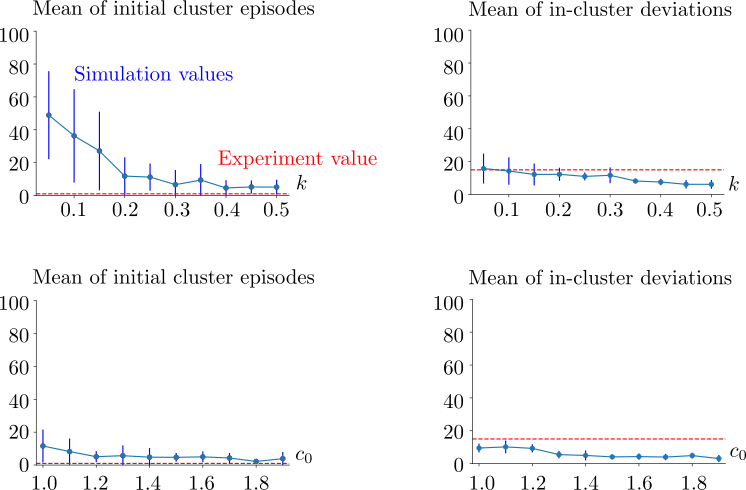

### statisticalH1.png

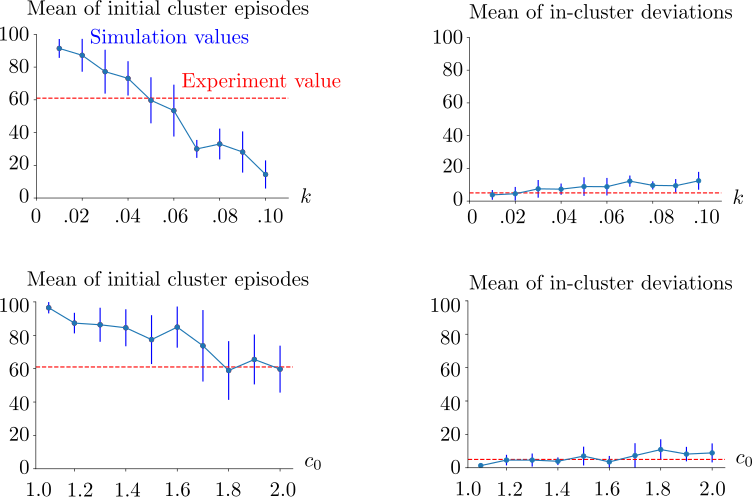

### statisticalH1Monkey.png

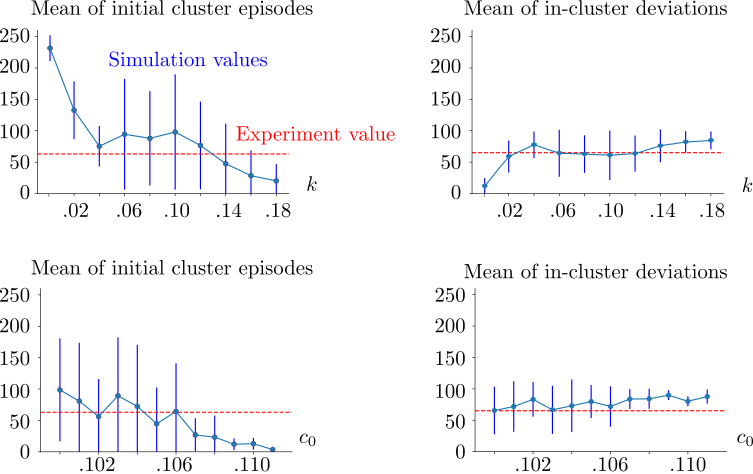

### withRewardFullSys1.png

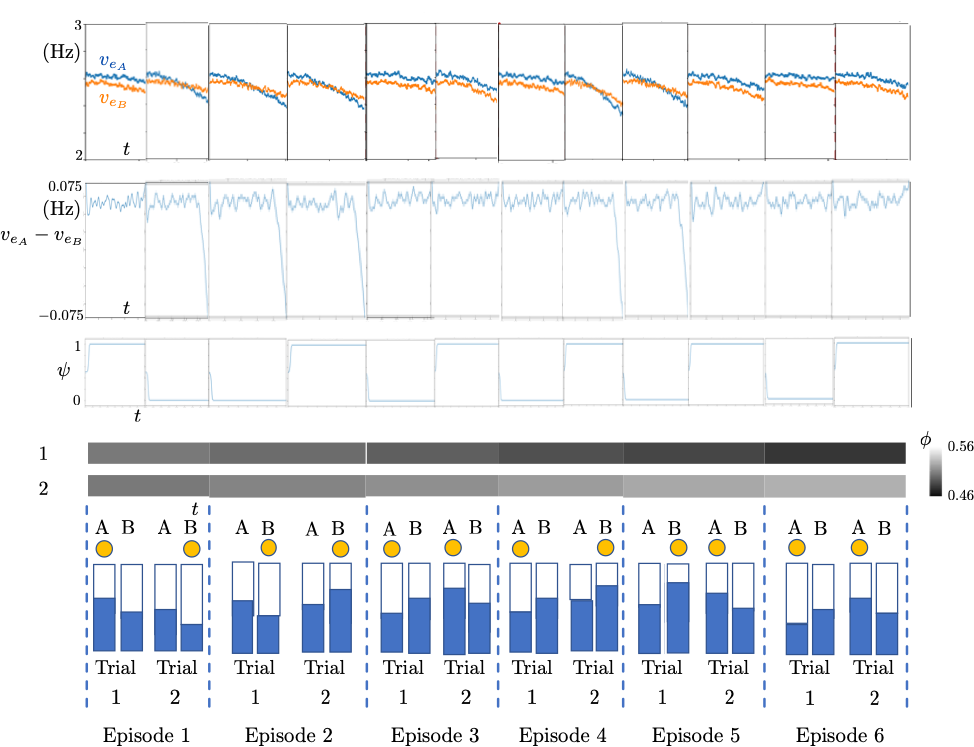
